# Climate change will redefine taxonomic, functional, and phylogenetic diversity patterns of Odonata in space and time

**DOI:** 10.1101/2022.04.04.486993

**Authors:** Tommaso Cancellario, Rafael Miranda, Enrique Baquero, Diego Fontaneto, Alejandro Martínez, Stefano Mammola

## Abstract

Climate change is rearranging the mosaic of biodiversity on our planet. These broad-scale species re-distributions will affect the structure of communities across multiple biodiversity facets (taxonomic, phylogenetic, and functional diversity). The current challenges to understand such effects involve focusing on organisms other than vertebrates and considering the signature of species redistribution on phylogenetic and functional diversity in addition to species composition. Using European dragonflies and damselflies (Odonata), we asked: i) how climate change will redefine taxonomic, phylogenetic, and functional diversity at continental scales; ii) which traits will mediate species’ response to global change; and iii) whether this response will be conserved across the phylogeny. First, we constructed stacked species distribution models for 107 species of Odonata under current and future climate conditions. Then, we quantified the temporal variation of taxonomic, functional and phylogenetic components, forecasting alpha and beta diversity changes through our geographical grid. Lastly, we used phylogenetic comparative models to test the influence of phylogeny and traits on range shifts. We observed broad latitudinal and altitudinal rearrangements in community composition driven by climate change. Given the high dispersal ability of Odonata, changes are predicted to be rapid, especially in areas experiencing faster climate change rates. According to our predictions, changes in species composition cascade to affect functional and phylogenetic diversity, determining broad turnovers in traits and evolutionary lineages. There was no clear phylogenetic signal in the range-shift response of European Odonata to climate change. According to our phylogenetic regression models, only body size and flight period can be partly correlated with observed range shifts. By considering all three primary facets of biodiversity, our results support the design of inclusive management and conservation strategies, accounting not only for the diversity of species, but also the services they provide and the phylogenetic heritage they carry in a targeted ecosystem.

## Introduction

Recent climate change is driving the reshuffling of the biodiversity patchwork on the Earth (Pecl *el al*., 2017). Upon those abrupt global changes, few species can survive *in situ* by adapting to the novel environmental conditions, whereas many more are forced to shift their ranges tracking their eco-physiological optima for growth and survival (Bellard *et al*., 2012; Diamond, 2018). Never before a single human generation witnessed such a rapid and massive biological migration induced by the increase of temperature, with terrestrial species rising towards higher latitudes and elevations and marine life sinking at greater depths (Perry *et al*., 2005; Chen *et al*., 2011; Lenoir *et al*., 2020). Inevitably, these rapid readjustments in species ranges are leaving a considerable imprint on the structure of local communities, which has cascading effects on ecosystem functioning and the provisioning of nature’s contribution to human societies (Nelson *et al*., 2013; Prather *et al*., 2013). The ecological and economic impacts of these changes will be unprecedented (Ripple *et al*., 2021).

Climate changes will lead to cumulative non-linear responses in the biological assemblages, which are expected to permeate through all biodiversity facets. This is because as climate changes, so does the distribution of certain species, with a ripple effect on species richness, trait composition, and evolutionary heritage of local communities (Saladin *et al*., 2020; Gallagher *et al*., 2013; Stewart *et al*., 2022). Therefore, the impact of climate change can be quantified by looking at predicted changes in the number of species that are present in an ecosystem (hereinafter “taxonomic diversity”), as well as in the diversity of function (“functional diversity”) and evolutionary lineages (“phylogenetic diversity”) represented therein. As approximative as the approach might be, a quantification of the rearrangement of these metrics is paramount to understand causally the mechanisms that drive the evolution of biodiversity across its multiple facets. Given that taxonomic, functional, and phylogenetic biodiversity are linked with ecosystem functioning and stability, ecologists and conservation biologists are increasingly considering these three facets when designing conservation plans (Pollock et al., 2020).

Historically, ecologists to assess the potential effects of environmental constraints on the biological communities have focused mostly on the variation of alpha diversity, which summarises the structure of a biological community as the total richness of taxa, traits, and evolutionary history, but does not incorporate information about their identity (Mammola *et al*., 2021a; Tucker *et al*., 2017; Pavoine & Bonsall, 2011; Petchey & Gaston, 2002). This may be problematic. Even if experimental studies have matched higher alpha diversity with greater resilience to perturbations, the lack of information on community composition prevents causal understanding of the mechanisms that may regulate this relationship insofar as the identity of the interplaying elements is lost (Wang & Loreau, 2014). Conversely, this information is retained in the calculation of beta diversity, which traces the individual elements that change across biological communities. To better understand these mechanisms, beta diversity can further be decomposed into its replacement and richness components (*sensu* Cardoso *et al*., 2014). In particular, the replacement component measures turnover between species across two sample units as a consequence of abiotic or dispersal processes (Fontana *et al*., 2020); whereas the richness component (*sensu* Cardoso *et al*., 2014) measures the gain or loss of species due to colonisation and extinction events (Fontana *et al*., 2020).

Here, we described the spatio-temporal effect produced by the shift of habitat suitability induced by climate changes on three biodiversity facets, incorporating both alpha and beta diversity metrics. We chose dragonflies and damselflies (Odonata) because they are well-established model organisms to address general macroecological questions in global change biology (Hassall, 2015; Grewe *et al*., 2013) and thermal physiology (Moore *et al*., 2021), being even regarded as “barometers” for climate change (Hassall, 2015). First, we modelled how global warming will affect the habitat suitability of each European species of Odonata. Next, we evaluated how the predicted changes in species habitat suitability will influence the Odonata communities in space, approximated using taxonomic, phylogenetic, and functional diversity. Finally, we used the predicted range shift to assess whether the response of Odonata to climate change is driven mainly by their evolutionary history or by distinctive biological and ecological traits. Under the assumption that Odonata species will track their ecological optima with dispersal, we expect to observe species redistributing poleward along the latitudinal gradient and upward along the altitudinal gradient. Furthermore, we predict that changes in community composition will permeate phylogenetic and functional components, such that alpha diversity will increase in areas with more conservative climates. In contrast, we predict that beta diversity change will be greater in areas experiencing faster climate change rates, especially so in the beta richness component given the high dispersal ability of Odonata. Lastly, we expect that the response of Odonata to climate change will also be explained by a shared evolutionary history, since phylogenetically related species may have similar patterns of distribution change and similar biological and ecological traits related to their dispersal ability.

## Materials and Methods

### 1 Rationale

To model species distribution, we used Species Distribution Models (SDMs), mainstream analytical tools in ecological and biogeographical research (Peterson *et al*., 2011; Franklin, 2010; Guisan and Thuiller, 2005), including to predict arthropod distributions (Mammola *et al*., 2021b). In short, distribution modelling refers to the practice of using an algorithm to infer a relationship between the occurrences for a given species (e.g., georeferenced points) and environmental predictors (e.g., climatic variables, topographic parameters, habitat type), forecasting its potential distribution in space and/or time. Due to the easy implementation and the often accessible interpretation of results (but see Ryo *et al*., 2021), species distribution models are routinely used in disciplines as diverse as conservation planning (Guisan *et al*., 2013), habitat restoration (Adams *et al*., 2016), invasion biology (Ficetola *et al*., 2009; Wang *et al*., 2007), and climate change biology (Santini *et al*., 2021; Guyennon *et al*., 2022).

As a model organism, we selected Odonata, an order of insects with tropical evolutionary origin (Pritchard and Leggott, 1987) and including species with contrasting thermal preferences. Odonata are well-established model organisms in ecology and behaviour (Clausnitzer *et al*., 2009; Córdoba-Aguilar, 2008; Corbet *et al*., 1999), and have been successfully used for tracking climate change using species distribution models (Hassall, 2015). These insects have an amphibiotic life with benthic vagile larvae living in freshwater habitats, whereas the adults are excellent fliers with high dispersibility compared to other freshwater invertebrates (Troast *et al*., 2016).

### 2 Taxonomic checklist and assembly of distribution data

We produced a complete checklist of all 169 European Odonata by merging the information of the “Atlas of the European dragonflies and damselflies” (Boudot and Kalkman, 2015) and the field guide “Dragonflies of Britain and Europe’’ (Dijkstra and Schröter, 2020) (Supplementary material S1). These are the most comprehensive references for European Odonata available today. We focused on the European continent because it has been intensively studied compared to other areas of the world (Titley *et al*., 2017). We excluded European Russia (including Kaliningrad) due to the scarcity of Odonata occurrences therein. We included Turkey to account for the entire arch of northern Mediterranean countries.

We downloaded all georeferenced occurrences of Odonata available at the Global Biodiversity Information Facility (GBIF, 09 January 2021; DOI: 10.15468/dl.kvrqug). Despite its biases (Beck *et al*., 2014), GBIF remains one of the most extensive global biodiversity databases (Zizka *et al*., 2020). The coverage provided by GBIF (highest coverage in UK, France, the Netherlands, Austria and Germany; lowest in southern and eastern Europe) for Odonata is congruent with the current expert-based knowledge about European odonates (Grewe *et al*., 2013; Kalkman *et al*., 2018).

We discarded data for fossil, non-European species, records before 1970, and occurrences falling outside the study area. We also removed duplicates and records with spatial uncertainty greater than the resolution of our predictor variables (∼10 km; see section 4). We minimised the effects of uneven sampling effort *via* spatial thinning with the function *reduceSpatialCorrelation* from the pack SDMworkshop (https://github.com/BlasBenito/sdmflow), setting the minimum.distance parameter to 1 (∼10 km) to match the resolution of our predictors.

### 3 Accessible area delimitation

For each species, we calibrated models within an accessible area, namely the geographical space that an organism has hypothetically occupied across its evolutionary history (Barve *et al*., 2011). In multi-species analyses, when lacking detailed information on species biogeographic history and dispersal ability, the simplest way to limit the boundary of the accessible area is by constructing a continuous border where most of the occurrences of a taxon are contained. For this, we used a Minimum Convex Polygon, the smallest area surrounding the points in which every internal angle does not exceed 180° (Burgman & Fox, 2003). We estimated a conservative Minimum Convex Polygon for each species using the R function *mcp* from the package *adehabitatHR* version 0.4.19 (Calenge, 2006), setting the percentage of outliers to be omitted at 1%. Finally, as a proxy of potential dispersal, we created an external buffer around each accessible area, weighting the distance with the flight period of each species [100 000distance in meters * (Flight period in months/10)], assuming that the flight ability across the species stays constant.

### 4 Selection of environmental predictors

We downloaded four variables from WorldClim 2 (Fick & Hijmans, 2017): monthly minimum and maximum temperature (°C), monthly precipitation (mm), and Digital Elevation Model (m a.s.l.). Current climatic data are the average for the period 1970–2000. We retrieved the water bodies’ map from the FAO’s GeoNetwork data portal. We adjusted the resolution of the water bodies’ map to 5 minutes using the function *resample* from the R package *raster* version 3.5-2 setting ‘bilinear’ method (Hijmans, 2020). Starting from the three climate variables (min/max temperature and precipitation), we calculated 19 bioclimatic variables using the function *biovars* from the R package *dismo* version 1.3-3 (Hijmans, 2020) and 16 environmental variables using the function *layerCreation* from the package *envirem* version 2.3 (Title & Bemmels, 2018). More information about the latter variables can be retrieved at https://www.worldclim.org/data/bioclim.html and https://envirem.github.io.

We visualise the multicollinearity effect amongst our 37 predictors variables (19 bioclimatic, 16 environmental, elevation, water bodies) via pairwise Pearson’s *r* correlation and a dendrogram based on variables’ distance matrix (Dormann *et al*., 2013). We extracted the final set of predictor variables at |*r*| < 0.5 (Mukaka, 2012) and then we removed variables with a Variance Inflation Factor (VIF) > 3 (Zuur et al., 2010).

We downloaded the same predictors for three future climate scenarios (Global Circulation Models: BCC-CSM1; MIROC-ESM-CHEM; NorESM1-M) and two time periods, 2050 (average for 2041–2060) and 2070 (average for 2061–2080). We chose a moderate Representative Concentration Pathway (RCP 4.5), namely a scenario that accounts for the greenhouse emission according to the current green policies (Hausfather & Peters, 2020). We assumed elevation and water bodies to remain constant in the future.

### 5 Modelling procedure

To model the distribution, we selected one algorithm for each main family of modelling algorithms (regression, maximum entropy, and decision trees) (Mammola *et al*., 2019; Mammola *et al*., 2018). We opted for Generalized Additive Model (GAM; Hastie & Tibshirani, 2017), MaxEnt (Phillips *et al*., 2006; Phillips *et al*., 2004), and Boosted Regression Trees (BRT; Elith *et al*., 2008), respectively, given their high performance (Elith *et al*., 2006). Furthermore, we compared the performance of each individual algorithm with an ensemble model, computed with the function *calc* in the package *raster*, since the aggregation of forecasts of different models (ensemble model) may improve the prediction habitat suitability of a given species (Araújo & New, 2007; Grenouillet *et al*., 2011). Specific settings and parameters for each algorithm are available in Supplementary material S2. To discriminate the areas where each species was more likely to be absent, we contrasted the presence data against a set of background points generated within their buffered accessible area. The number of background points doubled the number of presences (Phillips *et al*., 2009).

We evaluated the model performance using a holdout approach, whereby we used 75% of the occurrences of each species as a “train” dataset and the remaining 25% as “test” dataset to evaluate their predictive power. We calculated two performance metrics: Area Under the Receiver Operator Curve (AUC) and Boyce index (Hirzel *et al*., 2006). The AUC values range from 0 to 1, with higher values indicating better model discrimination. Whereas this metric is problematic for determining the absolute performance ability of SDMs, it is acceptable to use it for relative comparisons across models fitted with the same data (Zhang *et al*., 2021). The Boyce index is considered one of the most appropriate model evaluation metrics when absence data are lacking (Hirzel *et al*., 2006), and thus we chose it as a *proxy* measure of absolute model performance. The continuous Boyce index varies from –1 to 1: values above zero indicate model predictions consistent with distribution data, values around zero indicate performance no better than random, and values below zero refer to incorrect model predictions (Hirzel *et al*., 2006). We considered predictions with AUC < 0.7 and/or Boyce < 0.4 as low-quality performance.

After their evaluation, we fitted a final model for each species with the complete set of occurrences and used it to project potential distribution ranges under current and future climates. We converted the continuous habitat suitability projections into binary maps by using a threshold maximising the sensitivity (True Positive Rate) and specificity (True Negative Rate) (Liu *et al*., 2005; Martín-Vélez & Abellán, 2022). We calculated both spatial (e.g., suitable range size, mean elevation, and centroid) and biodiversity measures (see the next paragraph for biodiversity measures) only on the binary maps obtained from the best-performing modelling method (Qiao *et al*., 2015).

### 6 Estimation of taxonomic, functional, and phylogenetic diversity metrics

We calculated three diversity metrics for the predicted community of Odonata occurring within a cell of each raster map. We first stacked SDM projection for all the analysed species. We estimated taxonomic diversity as the number of species predicted to occur in each cell. We calculated functional and phylogenetic diversity as the total branch length entailed by the species predicted to occupy each cell, based on a functional and phylogenetic tree (Faith, 1992; Petchey and Gaston, 2002, 2006; Cadotte *et al*., 2010; see next sections). We chose tree-based descriptors of relationships to make the formulation of functional and phylogenetic diversity more comparable (Mammola *et al*., 2021a).

#### Estimation of the functional dendrogram

We calculated the functional tree for European Odonata using six traits broadly related to dispersal and species response to climate change, namely: total body size (mm), abdomen length (mm), wings length (mm), abdomen pigmentation (in RGB), habitat (lentic or lotic), and flight season time (in months) (Table 1). We focused on the adult stage because they disperse at large spatial scales via morphological (e.g., wings) and behavioural (e.g., reversible polarotaxis, repulsion/attraction of polarised light; Mitchell, 2018). In contrast, larva might disperse as well, but its ability is limited to the aquatic environment. Therefore, we expect that immigration promoted by climate change will involve mainly adults.

**Table 1.** Traits considered in the analyses with an indication of their expected functional meaning and the number of Gower distance groups (*sensu* de Bello et al., 2021).

| Trait | Trait type | Expected functional meaning | Bibliography | Gower group |
| --- | --- | --- | --- | --- |
| Body size | Biological<br>[Continuous] | Body size is tightly linked to temperature. Body size of assemblages of odonates is mainly driven by temperature. | (1; 2) | Group 1 |
| Abdomen length | Biological<br>[Continuous] | As for body size. |  | Group 1 |
| Wings length | Biological<br>[Continuous] | Proxy for dispersal. | (3; 4) | Group 1 |
| Habitat | Ecological<br>[Categorical] | Freshwater habitats (lentic/lotic) are among the most threatened ecosystems by climate change. | (5) | Group 2 |
| Flight season time | Ecological<br>[Continuous] | Indirect measure of dispersal potential. | (6) | Group 3 |
| Abdomen pigmentation | Biological<br>[Continuous] | Pigmentation and colour patterns are directly related with thermoregulatory mechanisms. For example, melanism is linked to greater absorption of solar radiation heat in cooler regions. | (1; 2; 7; 8; 9) | Group 4 |

(1) Hassall & Thompson, 2008; (2) Acquah-Lampsey *et al.*, 2020; (3) Outomuro & Johansson, 2019; (4) Rundle *et al.*, 2007; (5) Finlayson *et al.*, 2019; (6) Grewe *et al.*, 2013; (7) Okude & Futahashi, 2021; (8) Mani, 2013; (9) Suárez-Tovar *et al.*, 2022

**Table 2.**
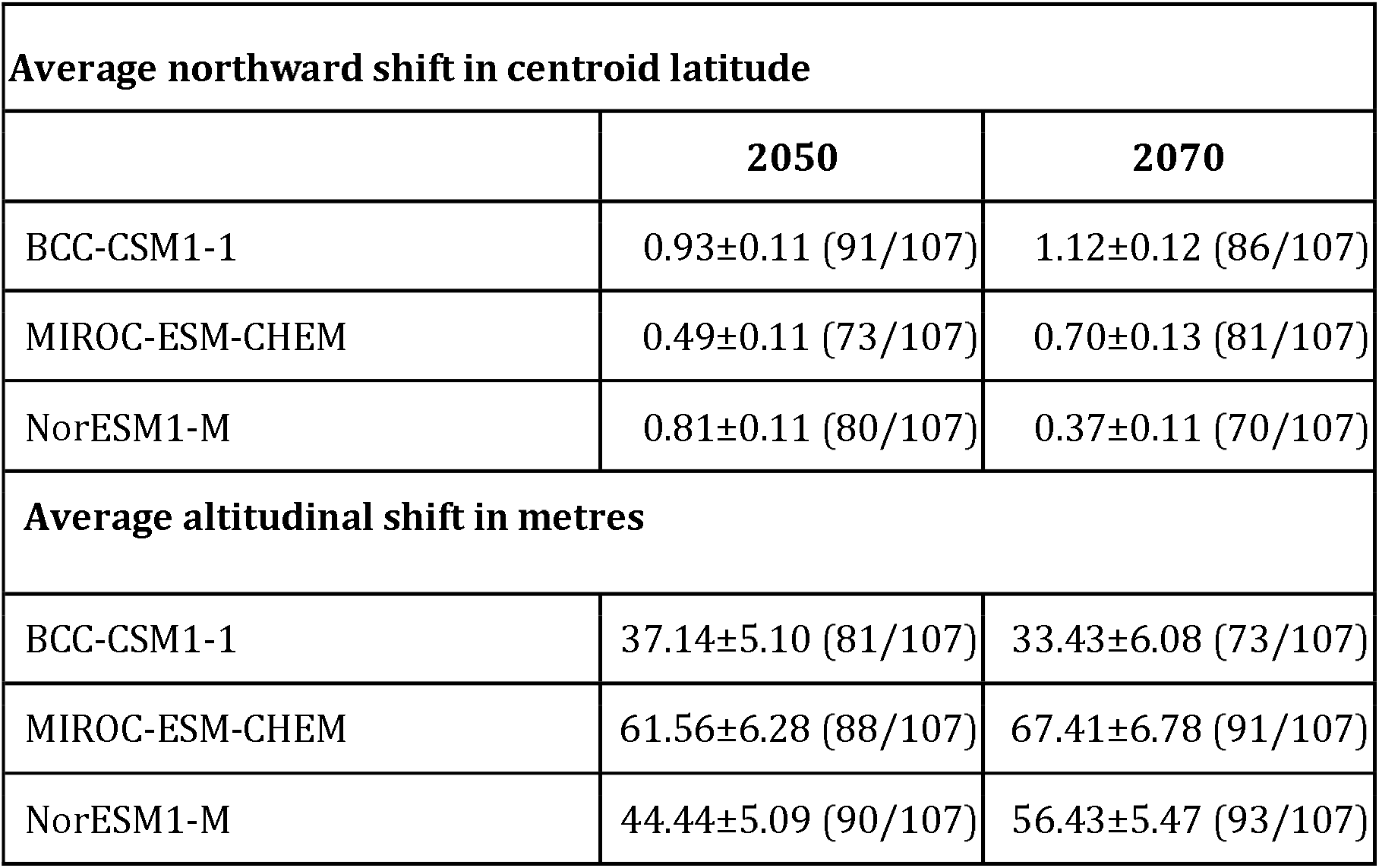
Magnitude and number of species shifting toward Northward latitudes and upper altitudes.

**Table 3.** Results of PGLS models with significative response variables (in bold). Total table containing PGLS results is in Supplementary material S11.

|  | Estimate | Std. Error | t value | Pr(>\|t\|) | Res. variable | Scenario | Time period |
| --- | --- | --- | --- | --- | --- | --- | --- |
| (Intercept) | 207228.608 | 52990.1971 | 3.91069705 | 0.00016874 | Centroid difference | MIROC | 2050 |
| Body length | -1828.0532 | 606.952165 | -3.0118571 | **0.00329576** | Centroid difference | MIROC | 2050 |
| Flight Season | 2777.48566 | 5808.89903 | 0.47814321 | 0.63360243 | Centroid difference | MIROC | 2050 |
| Habitat Lentic | 12534.6293 | 31484.3892 | 0.39812204 | 0.69139748 | Centroid difference | MIROC | 2050 |
| Habitat Lotic | 17994.3776 | 34691.3793 | 0.51869882 | 0.60512828 | Centroid difference | MIROC | 2050 |
| (Intercept) | 253527.566 | 63388.1236 | 3.99960673 | 0.00012243 | Centroid difference | MIROC | 2070 |
| Body length | -1767.575 | 709.51574 | -2.4912414 | **0.01439255** | Centroid difference | MIROC | 2070 |
| Flight Season | 1037.68713 | 6738.33589 | 0.15399754 | 0.87792517 | Centroid difference | MIROC | 2070 |
| Habitat Lentic | 7731.64985 | 39206.2133 | 0.19720471 | 0.84407166 | Centroid difference | MIROC | 2070 |
| Habitat Lotic | 8406.52507 | 42340.684 | 0.19854486 | 0.84302595 | Centroid difference | MIROC | 2070 |
| (Intercept) | 1.87163593 | 0.32124278 | 5.826235 | 7.23E-08 | Relative area change | BCC | 2070 |
| Body length | 0.00159414 | 0.00359034 | 0.44400809 | 0.6580156 | Relative area change | BCC | 2070 |
| Flight Season | -0.1305059 | 0.03658227 | -3.5674636 | **0.00055972** | Relative area change | BCC | 2070 |
| Habitat Lentic | 0.28382835 | 0.18772987 | 1.51189769 | 0.13377797 | Relative area change | BCC | 2070 |
| Habitat Lotic | 0.22166274 | 0.20737374 | 1.06890458 | 0.2877385 | Relative area change | BCC | 2070 |
| (Intercept) | 1.24519775 | 0.33799621 | 3.68405838 | 0.00037442 | Relative area change | MIROC | 2050 |
| Body length | 0.00813323 | 0.00379381 | 2.14381522 | **0.03449684** | Relative area change | MIROC | 2050 |
| Flight Season | -0.0631417 | 0.03622388 | -1.7430959 | 0.08442164 | Relative area change | MIROC | 2050 |
| Habitat Lentic | 0.26332324 | 0.20705918 | 1.27172934 | 0.20644872 | Relative area change | MIROC | 2050 |
| Habitat Lotic | 0.02818103 | 0.22588042 | 0.12476082 | 0.90096589 | Relative area change | MIROC | 2050 |
| (Intercept) | 1.67191708 | 0.25146401 | 6.64873314 | 1.63E-09 | Relative area change | NOR | 2050 |
| Body length | 0.00223091 | 0.00292457 | 0.76281493 | 0.44738758 | Relative area change | NOR | 2050 |
| Flight Season | -0.0984925 | 0.02816582 | -3.4968784 | **0.00070638** | Relative area change | NOR | 2050 |
| Habitat Lentic | 0.2071339 | 0.14418534 | 1.43658081 | 0.15398984 | Relative area<br>change | NOR | 2050 |
| Habitat Lotic | 0.23521455 | 0.1625593 | 1.44694616 | 0.151071 | Relative area<br>change | NOR | 2050 |

We determined the abdomen pigmentation from three pictures of each species, preferably downloaded from Dragonflypix (http://www.dragonflypix.com/index.html). We clipped the image around the abdomen using the software Gimp (GIMP Development Team 2019) and extracted the RGB colorspace using the function *getImageHist* (*colordistance* version 1.1.2; Weller, 2020). We obtained the mean value of the abdomen colour for each species as the average of the two predominant colours on the three photos (data available at https://osf.io/swnu4/download).

We calculated functional dendrograms (Petchey & Gaston 2002) with the *hclust* function in the R package *stats* version 4.1.0 (R Core Team 2020) and a Gower’s dissimilarity matrix constructed with the package *gawdis* version 0.1.3 (de Bello *et al*., 2021). This function is an extension of the classical Gower’s distance that provides a solution to limit unequal traits contribution when different traits are combined in a multi-trait dissimilarity matrix (de Bello *et al*., 2021) (functional dendrogram: Supplementary material S3). The Gower’s distance groups are reported in Table 1.

#### 6.2 Estimation of the phylogenetic tree

We calculated phylogenetic diversity from a tree calculated with sequences available in GenBank for the analysed species. We retained the five molecular markers (16S rRNA gene; 18S rRNA gene; Cytochrome c oxidase subunit I, COI; Histone H3; NADH dehydrogenase subunit 1, NADH) with the highest taxonomic coverage. We aligned each marker separately using the E-INS-i algorithm implemented in MAFFT v.7 (Katoh & Standley, 2013). We translated alignments of protein-coding genes into amino acids and checked them for indels and stop codons. When multiple sequences were available for the same species, we chose the one with the greatest quality and length. Our final alignment included a 1996 base pair for the 16S rRNA gene (number of aligned sequences 87), 1772 base pairs for the 18S rRNA gene (37), 658 base pairs for COI (101), 329 base pairs for H3 (17), and 340 base pairs for NADH (31). We concatenated gene fragments with SequenceMatrix (Vaidya *et al*., 2011) and selected the optimal partition scheme using the Akaike Information Criterion calculated in PartitionFinder (Lanfear *et al*., 2017). We calculated ultrametric phylogenetic trees using BEAST 2 (Bouckaert *et al*., 2019), setting a relaxed molecular clock model for each partition and a Yule model for the estimation of the topology. Our four Markov Chain Monte Carlo were allowed to run for 100 000 000 generations and sampled every 10 000 generations. The 10% of initial trees were discarded. We used Tracer version 1.7.1 (Rambaut *et al*., 2018) to confirm the correct mixing of all the parameters and TreeAnnotator version 2.6.0 (Bouckaert *et al*., 2019) to calculate the consensus tree (Supplementary material S4).

#### 6.3 Elaboration of the taxonomic, functional, and phylogenetic diversity maps

We assembled taxonomic, functional, and phylogenetic diversity maps using modified versions of the functions *alpha* (temporalAlpha) and *beta* (temporalBeta) from the package *BAT* version 2.7.1 (Cardoso *et al*., 2021). First, we stacked the binary maps obtained from the best-performing SDMs models of each species. Then, we calculate alpha diversity across the three biodiversity facets for present and future stacked maps. We quantified variations in alpha diversity between present and future scenarios by subtracting the alpha diversity values in the future and the present. We calculated beta diversity in the same way, estimating replacement and richness components of beta diversity (Cardoso *et al*., 2014) for each cell comparing future and present communities. To calculate the alpha/beta functional and phylogenetic diversity, we used the functional or phylogenetic tree as an additional parameter into the functions.

### 7 Testing for phylogenetic signal and trait influence on species response to climate change

We used phylogenetic comparative methods to examine the influence of phylogeny and traits on the responses of Odonata to climate change. We characterised the response to climate change of each species using three response variables: i) the proportional variation in habitat suitability, calculated as the ratio between future and current predicted area; ii) the altitudinal shift in the distribution, estimated as the difference between future and current mean altitude; and iii) the centroid shift in the distribution, measured as the linear distance between the position of future and current centroid. We used the function *distGeo* from the r package *geosphere* version 1.5-14 to estimate the centroid position.

We investigated whether closely related species experience similar responses to climate change using Pagel’s λ and Blomberg’s K, as implemented in the function *phylosig* from the R package *phytools* version 0.7-80 (Revell, 2012). Values close to 0 indicate a weak phylogenetic signal, whereas values close to 1 or higher suggest the presence of phylogenetic signal. We then visualised the phylogenetic signal of each trait using ancestral character reconstruction as implemented the function *contMap* of the R package *phytools*.

Finally, we explored the relationship between traits and the species response to climate change (approximated with the three response variables above) using Phylogenetic Generalized Least Squares (PGLS), using the function *pgls* from the package *caper* version 1.0.1 (Orme *et al*., 2018). We used three functions of branch transformation (lambda, kappa, and delta) to adjust the covariance matrix to the data selecting the best transformation through a maximum likelihood procedure. Prior to model fitting, we performed data exploration, visually inspecting for the presence of outliers in the predictor and response variables with dotcharts and verifying multicollinearity among predictor variables (Zuur et al., 2010).

### 8. Reproducibility

In constructing and reporting SDMs, we followed the ODMAP (Overview, Data, Model, Assessment and Prediction) protocol (Zurrell *et al*., 2020), designed to maximise reproducibility and transparency of distribution modelling exercises. The ODMAP for this study is available as Supplementary material S2.

We stored all data, raw predictor variables and detailed model outputs in the OSF repository (https://osf.io/4rjuc/). All code used to perform analyses and produce plots is available in GitHub (https://github.com/TommasoCanc/Odonata_SDM_2022).

## Results

### 1 Species distribution models

#### 1.1 Predictor variables and model performance

We successfully calculated the habitat suitability for 107 species of European Odonata out of 169 contained in our checklist. We omitted 62 species due to the low number of occurrences available in GBIF. Our models incorporated seven non-collinear predictors: waterbodies, elevation, Emberger’s pluviometric quotient (embergerQ), temperature annual range (bio 7), mean temperature of the wettest quarter (bio 8), mean temperature of the warmest quarter (bio 10), and precipitation seasonality (bio 15). Multicollinearity and Variance Inflation Factor analyses are reported in Supplementary material S5. Boosted Regression Trees favoured the AUC in 96 species, the ensemble of models in six species, the Maximum Entropy in four, and the Generalized Additive Model only in one (Supplementary materials S6).

#### 1.2 Species distribution model future predictions

In accordance with our first hypothesis, climate projections consistently predicted an increase in habitat availability for the majority of species towards northern regions of Europe and at upper elevations (Tab. 2). These shifts were coupled with a contraction of suitable areas in the Mediterranean regions. We found only minor variations among the predictions under different future Global Circulation Models. The increased habitat availability in the northern areas is highlighted from a centroid’s shift towards northern latitudes for the majority of Odonata species (Tab. 2). The model outcomes also revealed a rise of mean elevation occupied by many species of Odonata in the future climate scenarios (Tab. 2). Example model projections for one of the species is available in Figure 2 (see Supplementary material S7 for the entire set of species).

**Fig. 1.**
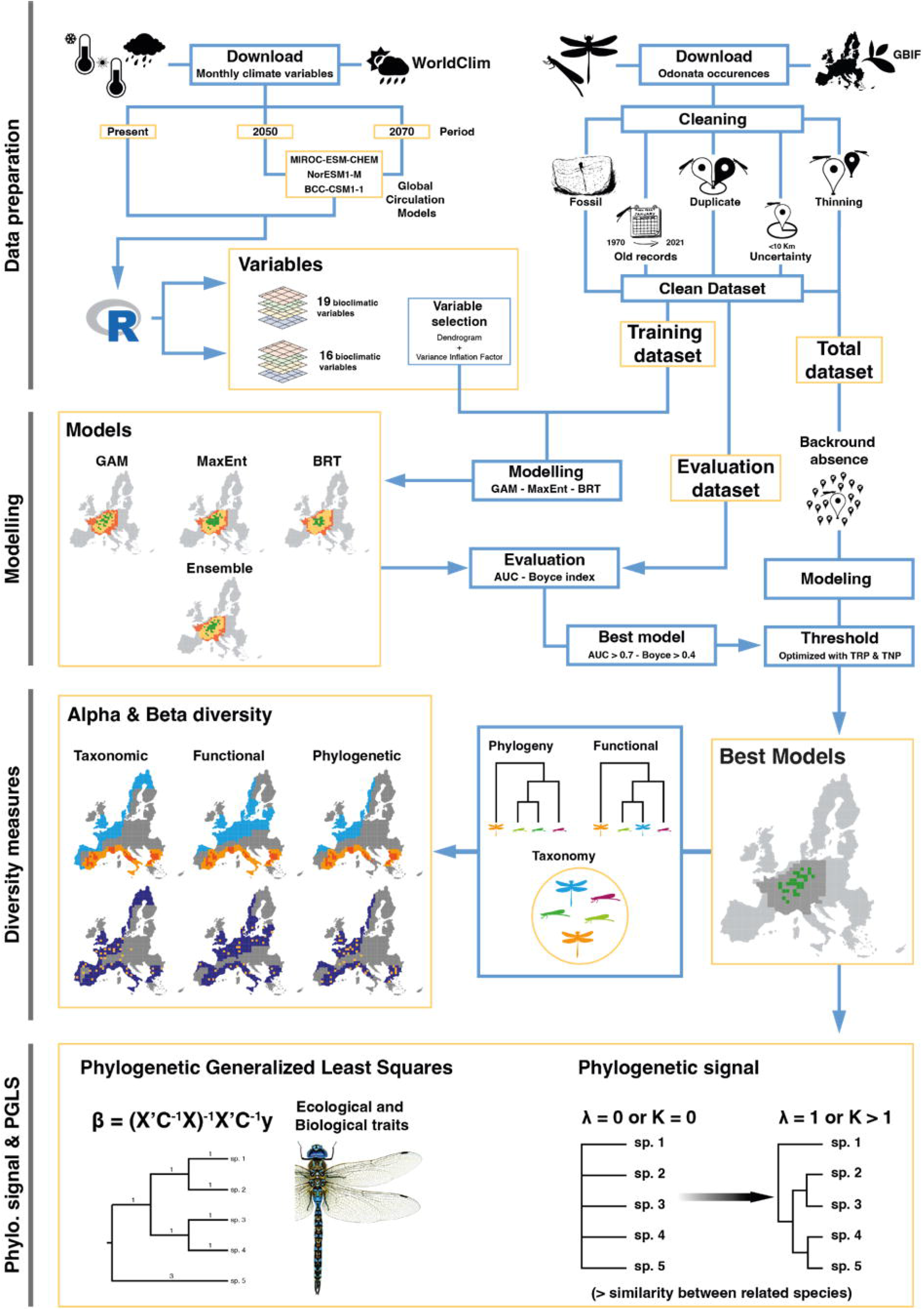
Infographic summarising the study workflow. In this work, we first constructed a species distribution model for each species of European Odonata to predict their current and future habitat suitability. Then, we stacked the model projections and used community-level data to quantify the temporal variation of taxonomic, functional, and phylogenetic diversity via estimating alpha and beta diversity. Finally, we used the predicted range shift to assess whether the response of Odonata to climate change is driven mainly by their evolutionary history or by distinctive biological and ecological traits.

**Fig. 2.**
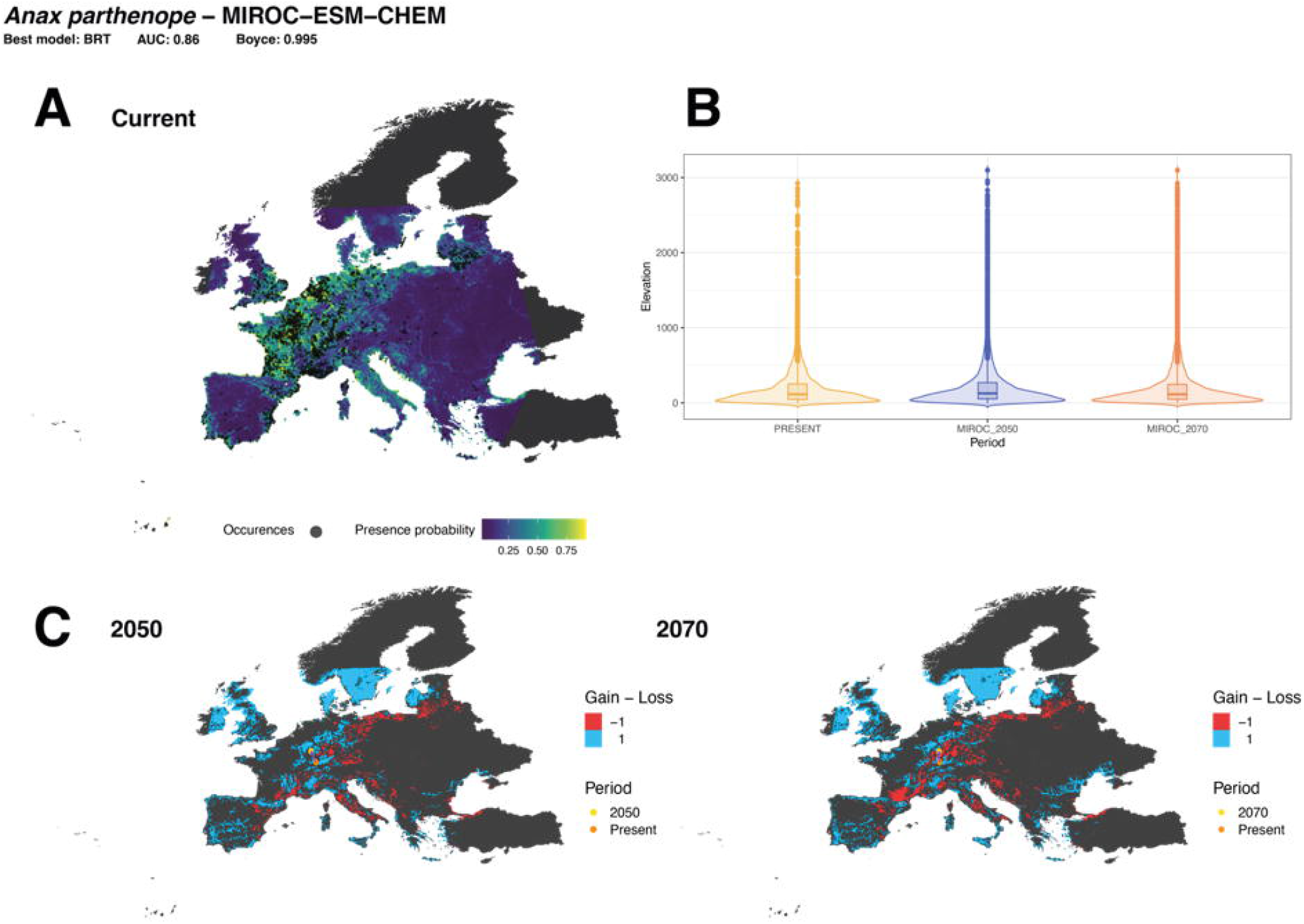
Example of summarised species distribution model projections for an individual odonate species. A) Best model prediction map for the current time period. B) Extent of elevation shift across time periods. C) Variation of habitat availability between future and current time periods. Habitat gain and loss are depicted with blue and red colours respectively. Centroid shift is represented by the variation among the orange (present) and yellow point (future). Summarised SDM outcomes for all species are available in Supplementary material S7.

### 2 Quantification of change of biodiversity measures

We calculated taxonomic, functional, and phylogenetic diversity for 105 of the 107 species, since we lack genetic data for *Orthetrum taeniolatum* (Schneider, 1845) and *Sympetrum sinaiticum* Dumont, 1977.

#### 2.1 Alpha diversity patterns

Alpha diversity revealed congruent patterns across its three facets under the current climate scenario. The highest values of taxonomic diversity concentrated around the Central-Atlantic European region. Compared to taxonomic diversity, higher values for functional and phylogenetic diversity were attained in Italy, Ireland, and the North of the United Kingdom (Fig. 3).

**Fig. 3.**
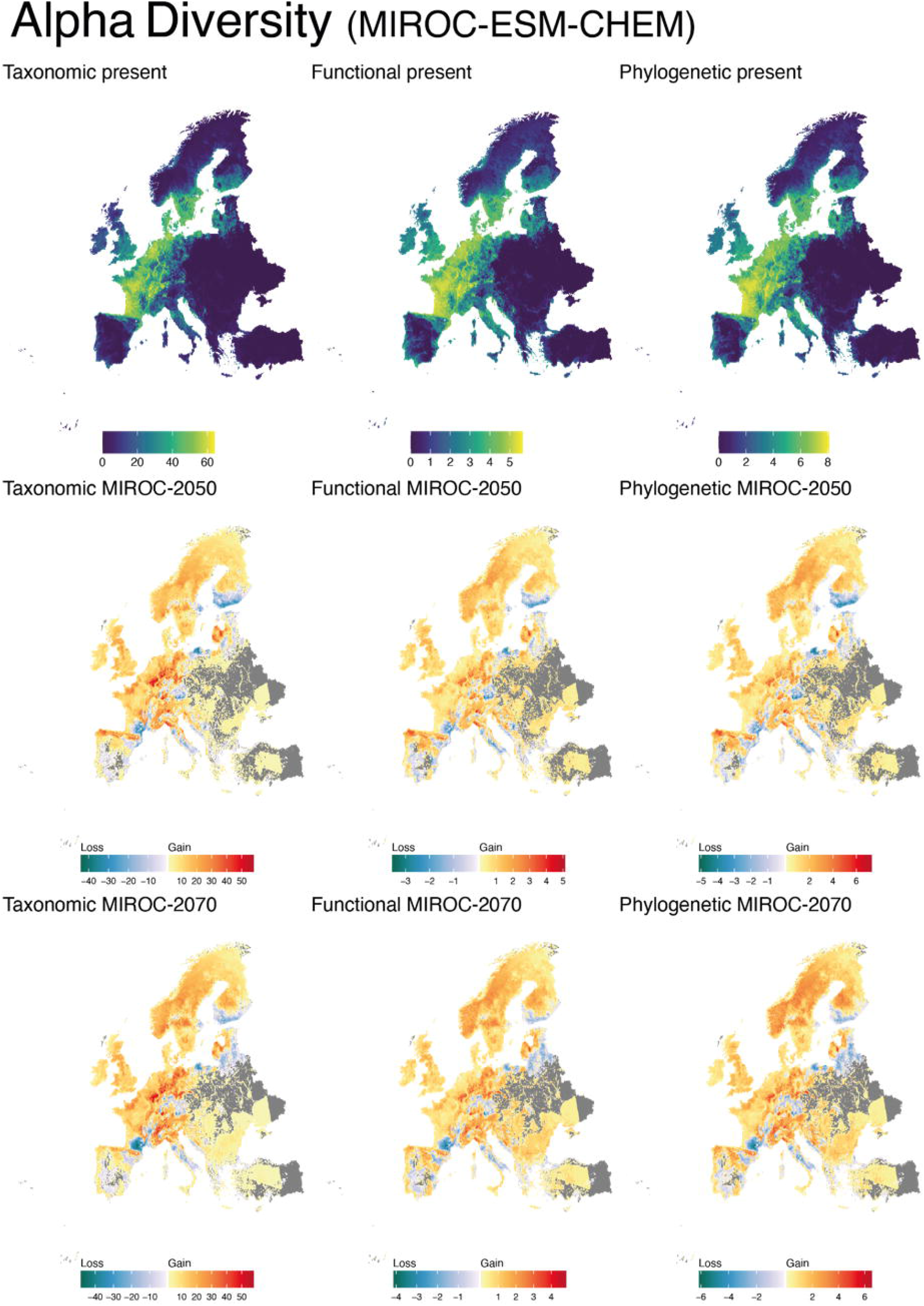
Quantification of alpha diversity per different climate scenarios (BCC-CSM1-1; MIROC-ESM-CHEM; NorESM1-M) and time periods (current; 2050; 2070). For future scenarios, the cold-colour gradient indicates the extent of species loss, whereas the warm-colour gradient indicates the species gain.

Alpha diversity projections towards the future revealed a substantial geographical re-arrangement over time again across the three biodiversity facets, with main increments in taxonomic, functional, and phylogenetic diversity recorded in the northern and eastern regions of Europe, particularly in the Scandinavian Peninsula, the British Isles, and the Black Sea region. In contrast, a decrease in the three facets of alpha diversity was predicted in Central Europe and Mediterranean areas, particularly in France, Germany, and the Baltic countries, as well as the Hellenic, Italian, and Iberian Peninsulas. Furthermore, the shift towards higher altitude predicted for many species was visible as an overall increase of species richness, together with functional and phylogenetic diversity, in the main European mountain range such as the Alps, Cantabrian Mountains, and the Pyrenees (Fig. 3; Supplementary material S8).

#### 2.2 Beta diversity patterns

We observed greater beta diversity in the Iberian Peninsula, Scandinavia, and scattered areas across western Europe. Although with different values, this pattern was congruent across the three biodiversity metrics and future Global Circulation Models, not differing substantially between 2050 and 2070 predictions (Fig. 4). These changes were primarily explained by changes in the richness component of beta diversity, rather than replacement. The highest values of beta richness were predicted for the Iberian Peninsula, Turkey, Scandinavia, and Eastern European countries. The highest values of beta replacements were registered for the Iberian Peninsula, the Balkans, and the Baltic countries (Fig. 4; Supplementary material S8).

**Fig. 4.**
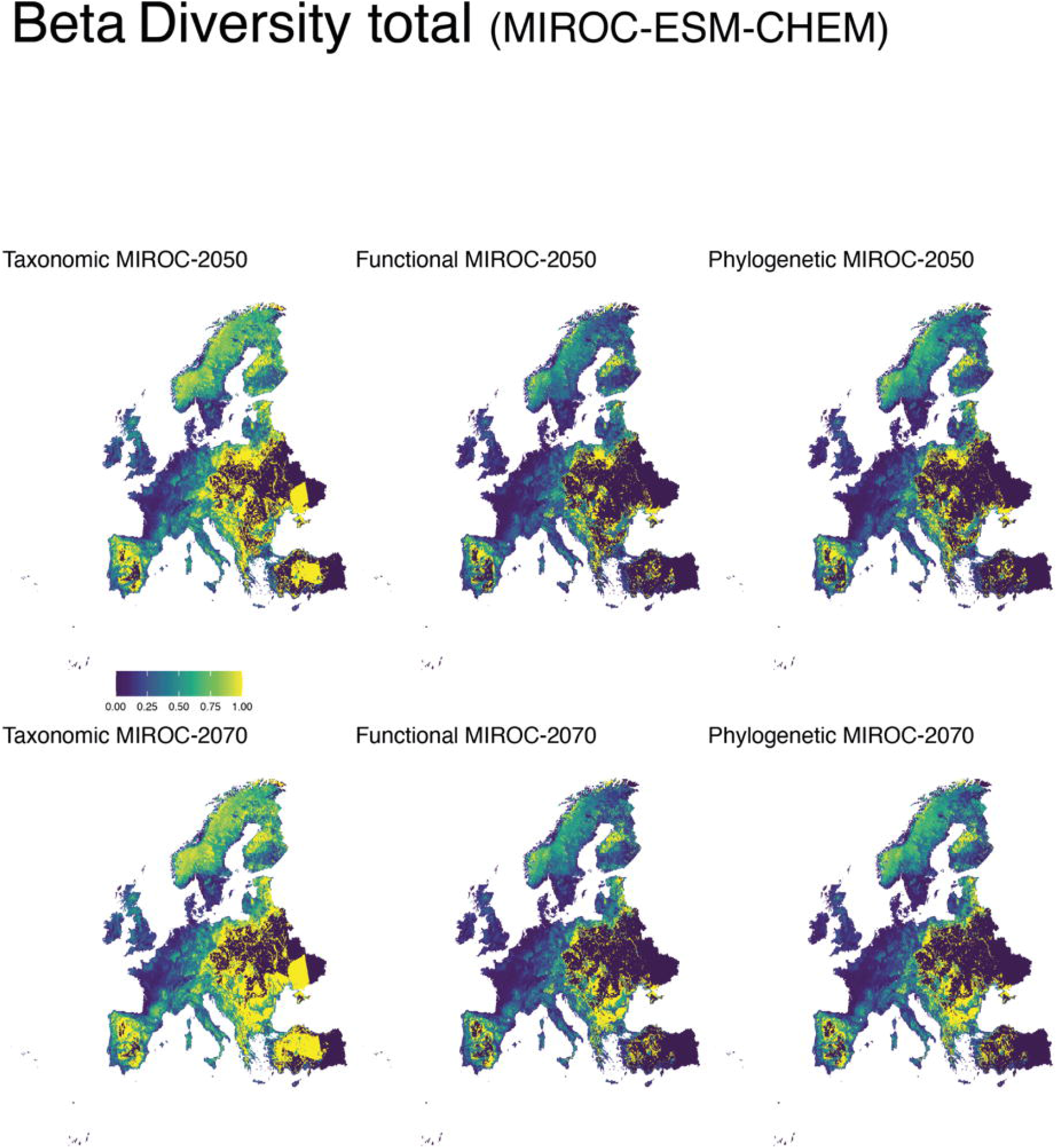
Quantification of total-beta diversity (beta-replacement + beta-richness *sensu* Cardoso *et al*., 2014) per different climate change scenarios (BCC-CSM1-1; MIROC-ESM-CHEM; NorESM1-M) and time periods (current; 2050; 2070).

### 3 Phylogenetic signal and Phylogenetic Generalized Least Squares

We observed a negligible phylogenetic signal in both Pagel’s λ and Blomberg’s K for all predictor variables (proportional variation in habitat suitability; altitudinal shift; centroid shift), climate scenarios (BCC-CSM1; MIROC-ESM-CHEM; NorESM1-M) and time periods (2050; 2070). The only exception was centroid shift (MIROC-ESM-CHEM 2050), for which Pagel’s λ revealed a significant phylogenetic signal (λ = 0.73; p = 0.003; Supplementary material S9). The lack of phylogenetic signals is further confirmed by the ancestral character reconstruction analysis, revealing no clustering pattern nested into the phylogeny (an example in Fig. 5, Supplementary material S10).

**Fig. 5.**
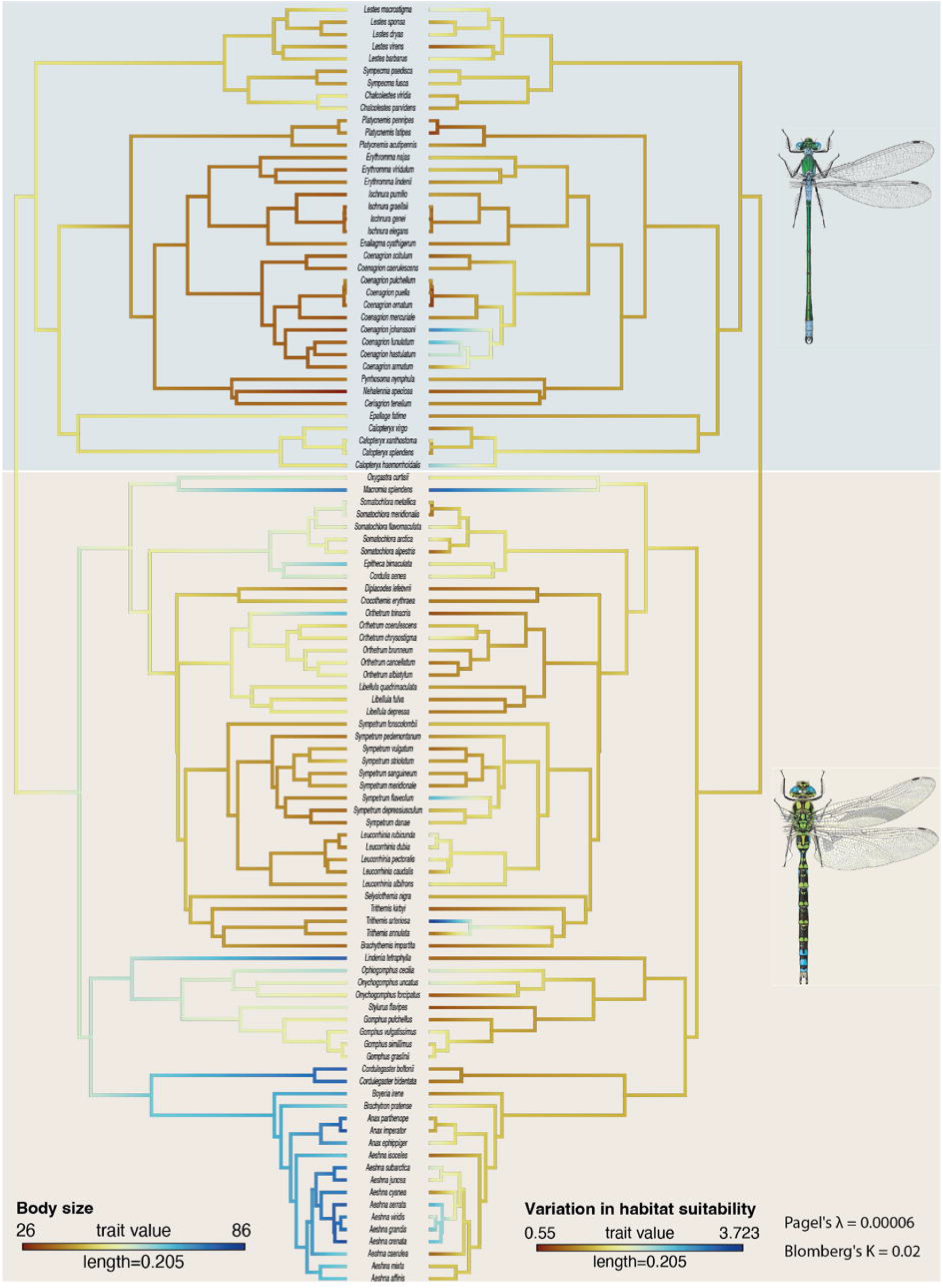
Reconstruction of ancestral character states for the variables body size (left) and variation in habitat suitability (right). Pagel’s λ and Blomberg’s K indicate the estimated values for the response variables “Variation of habitat suitability” (see Supplementary material S10 for the other tree of ancestral character reconstructions). “Length” in the legend provides the scale for the branch lengths of the phylogenetic tree. The grey box delimits the Zygoptera clade whereas the brown one the Anisoptera clades.

For most climate scenarios and time periods, the outcomes of PGLS only partly support the effect of traits (body length, flight period, and habitat) on the response of Odonata to climate change. Body size and flight period returned significant effects on the proportional variation in habitat suitability and the centroid shift for different global circulation models and time periods (Tab. 3). No other traits revealed significant effects (Supplementary Material S11).

## Discussion

In this study, we forecasted variations in future habitat availability for 107 species of European Odonata. Specifically, we quantified the impact of those changes as regional changes of alpha and beta diversity, and explored the role played by their evolutionary history or specific traits in promoting such changes. Overall, our results predict conspicuous readjustments in the Odonata communities following climate change; these changes permeate through all facets of biodiversity. Conversely, we did not find evidence that closely related species respond in a similar way to climate change, since there was no clear phylogenetic signal associated with the magnitude of range shift across the evolutionary tree of European Odonata. After accounting for phylogenetic effects, one biological (body size) and one ecological (flight period) trait affected the extent of change in distribution range induced by climate change.

Foremost, our projections consistently predicted an increase in habitat suitability towards northern latitudes and upper elevations coupled with a contraction of suitable areas in the Mediterranean regions for most species of dragonflies and damselflies. This result was largely expected, since shifts induced by climate change are well-documented for many freshwater invertebrates (Hickling *et al*., 2005, 2006; Heino *et al*., 2009; Mustonen et al., 2018), although their effects on the structure and composition of natural communities are still poorly examined. More precisely, our results hint that biological communities will not reshuffle randomly in the future. Indeed, our predictions show that odonate communities will face a future taxonomic rearrangement, paralleled by a readjustment of provided functional services and of the amount of evolutionary heritage enclosed in their aggregations. This has important implications for species management (Samways *et al*., 2020) and societal consequences since invertebrates represent irreplaceable nodes into ecological networks providing uncountable ecosystem services to human communities (Eisenhauer *et al*., 2019; Cardoso *et al*., 2020).

Further gains or losses of biological diversity in its all three facets are challenging to predict using correlative methods, since they will depend on non-linear species-interactions branching throughout the system. For instance, increasing taxonomic, functional, and phylogenetic diversity might import new evolutionary lineages or improve the resilience of natural systems with novel ecosystem functions (Thomas, 2020). Also, the new possibilities of interaction might increase the competitive pressure among the species (Krosby *et al*., 2015), and boost the chances of hybridisation of previously isolated taxa (Bybee et al., 2016). For example, the rise of hybridisation events between two European damselflies *Ischnura elegans* (Vander Linden, 1820) and *Ischnura graellsii* (Rambur, 1842) have been documented and attributed to climate-driven range expansion of *I. elegans* to areas formerly occupied exclusively by *I. graellsii* (Sánchez-Guillén *et al*., 2011). In contrast, the decrease of biodiversity components could reduce ecosystem’ stability and resilience due to narrowing possible species-specific responses to environmental fluctuations leading to functional homogenization (Tobias & Monika, 2012) and reducing genetic diversity (Pauls *et al*., 2013). A homogenisation of the odonate communities driven by climate change and urbanisation has already been documented for the North American populations. Indeed, Ball-Damerow *et al*., (2014) demonstrated that changes in environmental conditions led to a homogenization of odonates community favouring the expansion of highly mobile habitat generalists species and a parallel loss of habitat specialist or species with the peculiar physiological state as diapause.

Not without caveats (e.g., abundance patterns or fitness; Lee-Yaw *et al*., 2021, species coexistence; Pichler & Hartig, 2021), species distribution models are considered robust and reliable correlative approaches to map the species’ potential habitat preference across space and time. Despite evaluation metrics suggesting our models being robust, we acknowledge that their outcome is unavoidably coupled with the goodness of the variables selected (Fourcade *et al*., 2018). Therefore, due to the lack of specific habitat variables (e.g., intermittent freshwater habitats or future water extension) and the main use of climatic variables, our projected ranges must be interpreted as general indications of future trends of biodiversity change, rather than precise descriptions of species range boundaries. Another potential criticism to our approach is the inference of the habitat availability of odonate using species occurrence retrieved from GBIF. We are conscious about the limitations of GBIF data (e.g., samples collected opportunistically and spatially distorted). However, we consider that the occurrences used to perform the model are in line with the actual knowledge about the current distribution of Odonata. Moreover, the availability of high-quality field guides and the facility to recognize the adult stage of these insects (compared to other freshwater insects) allows limiting taxonomical errors. Finally, our models do not consider potential immigration events of non-European species (for example, *Trithemis kirbyi* Sélys, 1891 is one of the most recent species that arrived in Europe from Africa due to climate changes; Boudot and Kalkman, 2015). Therefore, future estimates of biodiversity facets values might be slightly underestimated. Despite these limitations, we still consider our results of the current alpha taxonomy plausible since they agree with the distribution map proposed by Kalkman *et al*. (2018).

Despite existing evidence that supports the tendency of the species to migrate toward poleward latitudes and upper altitudes in response to climate change (Freeman *et al*., 2018; Parmesan, 2006; Chen *et al*., 2011), sometimes the observed movements may follow unexpected directions compared to those predicted by models (Diamond, 2018). Therefore, forecasting how the habitat availability might shift across species in the near future, exploring how such change may affect biological communities and uncovering the role played by biological and ecological traits in the organism range shift, is critical to designing effective management and conservation plans (Guisan and Thuiller, 2005). In the present work, we did not find any significant relation between the predicted range shift and phylogenetic relatedness of European Odonata coupled with the lack of clusterization in the reconstruction of the ancestral character states. These results highlight how the effect of climate change will be pervasive across the entire phylogenetic tree of odonate with responses species-specific to climate variation. Moreover, PGLS models’ outcomes did not show a solid and consistent effect of traits on range shifts. These results are in line with those proposed by Grewe *et al*., 2013, where neither biological (e.g., abdomen length and wing size) nor ecological (e.g., flight period) traits have returned significant relation with observed range shift.

Therefore, further research about ecology, physiology, and behaviour can benefit our knowledge about these freshwater insects and favour the design of efficient and effective conservation strategies. Further investigations based on mechanistic models (Chichorro *et al*., 2022) or high-resolution physiological and dispersal traits (Buckley & Kingsolver, 2012; Mammola *et al*., 2021b) could be useful to better identify key traits associated with climate-induced species range shifts and potentially even extinction risk. Our results might be substantially improved by including into the models traits directly linked with the dispersal ability (e.g., GPS-tracking, flight muscle mass, wing loading and shape) as well as traits and distribution of the larval stages, since most of the life of these insects is spent underwater [e.g., in *Anax imperator* (Leach, 1815) the life span is two years in larvae and eight to nine weeks in adults (Corbet, 1957)]. Unfortunately, this kind of information is still scarce for most odonate species. Therefore, further basic biological research about ecology, physiology, and behaviour can benefit our knowledge about these freshwater insects and favour the design of efficient and effective conservation strategies.

## Supporting information

S1_Odonata_Species_List

S2_ODMAP_CancellarioEtAl__2021-09-29-2

S3_Functional_dendrogram

S4_Phylogenetic_tree

S5_Dendrogram_collinearity

S6_Model_performances

S7_Supplementary_material_Odonata_SDM

S8_Alpha and Beta_v2

S9_Phylogenetic_signal

S10_Ancestral_character_recostruction

S11_Total table_PGLS

## ACKNOWLEDGEMENTS

SM acknowledges support from the European Commission through Horizon 2020 Marie Skłodowska-Curie Actions (MSCA) individual fellowship (grant no. 882221).

## AUTHOR CONTRIBUTIONS

TC, AM, and SM conceptualised the study and prepared the original draft. TC led the modelling analysis and functional trait calculation, with support and advice by SM. AM and TC analysed phylogenetic data. RM, EB, and DF contributed to conceptualization and planning. All authors contributed to writing, reviewing, and editing.

