## Supplementary material for "Climate change will redefine taxonomic, functional, and phylogenetic diversity patterns of Odonata in space and time": S2_ODMAP_CancellarioEtAl__2021-09-29-2

– ODMAP Protocol –

Tommaso Cancellario, Rafael Miranda, Enrique Baquero, Diego Fontaneto, Alejandro Martínez, Stefano Mammola

2021-09-29

### Overview

##### Authorship

Contact :

##### Model objective

Model objective: Forecast and transfer

Target output:

- continuous occurrence probabilities
- binary maps of gain/loss habitat availability
- alpha and beta taxonomy, functional, and phylogenetic diversity

##### Focal Taxon

Focal Taxon: Odonata

##### Location

Location: Europe

##### Scale of Analysis

Spatial extent: -31.25, 44.83333, 27.66667, 71.16667 (xmin, xmax, ymin, ymax)

Spatial resolution: ~10 km grid squares (0.08333333)

Temporal extent:

- average for the period 1970-2000 (Present)
- average for 2041–2060 (2050)
- average for 2061–2080 (2070)

Boundary: rectangle

##### Biodiversity data

Observation type: field survey, citizen science, standardised monitoring data

Response data type: presence-only

##### Predictors

Predictor types: climatic, topographic, habitat

##### Hypotheses

Hypothesis: We hypothesized that the future changes of habitat suitability of odonates under the press ion of climate change would cause alteration in their communities with ripple effects on the most significant biodiversity components as taxonomic, functional, and phylogenetic diversity.

##### Assumptions

Model assumptions:

- We assumed that species are at equilibrium with the environment
- We assumed that elevation and waterbodies do not change across the time

##### Algorithms

Modelling techniques: gam, brt, maxent

Model complexity: We optimized each statistical technique choosing different modelling parameters. Model parameters were chosen to obtain good robustness from the models avoiding overfitting. See the scripts about the models at GitHub for further details.

Model averaging: We combined the results of the three models into an ensemble. The ensemble was obtained averaging the results of the three models.

##### Workflow

Model workflow:

- To estimate the potential habitat suitability, we confronted the performance of GAM, MaxEnt, and BRT.
- We computed the ensemble model calculating the average between the three models previously mentioned.
- To evaluate model performance, we used a holdout approach. We split the occurrences of each species into a train (75%) and test (25%) dataset.
- In order to compare the model performances, we used: Area Under the Receiver Operator Curve (AUC) and Boyce index.
- Once we evaluated the models, we determined the potential distribution range of the species under current and future climates using the complete dataset.
- We converted the continuous habitat suitability projections into binary maps by using a threshold maximising the sensitivity (True Positive Rate) and specificity (True Negative Rate).
- After selecting only the binary maps deriving from the best-performing modelling method, we calculated both spatial and biodiversity measures among present and future climate conditions.

##### Software

Software: All the analysis were run in R version 4.1.0.

Package: adehabitatHR (Calenge, 2020), dismo (Hijmans, 2020), envirem (Title and Bemmels, 2018), gbm (Greenwell *et al.,* 2020), mgcv (Wood, 2021), raster (Hijmans, 2020), SDMworkshop (<https://github.com/BlasBenito/sdmflow>)

Code availability: <https://github.com/TommasoCanc/Odonata_SDM_2022>

Data availability: <https://doi.org/10.15468/dl.kvrqug>

### Data

##### Biodiversity data

Taxon names: The full list of species is available at Supplementary material S1.

Taxonomic reference system: We follow the taxonomy contained in the Atlas of the European dragonflies and damselflies (Boudot and Kalkman, 2015) and the field guide Dragonflies of Britain and Europe (Dijkstra and Schröter, 2020).

Ecological level: species, communities

Data sources: We used the data from GBIF. Accession date: 09 January 2021. DOI: <https://doi.org/10.15468/dl.kvrqug>

Sampling design: miscellaneous

Sample size: The full list with the number of species occurrences used to perform the final models is available at Supplementary material S1.

Clipping: All data were included within the boundary of our predictor raster variables.

Scaling: To define the accessible area for each species we estimated a conservative Minimum Convex Polygon setting the percentage of outliers to be omitted from the computation corresponding to 1%. Then, to consider the potential dispersion for each species, we created an external buffer around each accessible area, weighting the distance with the flight period of each species.

##### Cleaning

To enhance GBIF’s spatial information data quality:

- We removed the records classified as ‘fossil’ and those before 1970.

- We excluded non-European species, the occurrences falling outside the study area, the records with identical coordinates, and those with spatial uncertainty greater than our predictor variables resolution (~10 km).

- We performed spatial thinning with the function *reduceSpatialCorrelation* from the pack SDMworkshop (https://github.com/BlasBenito/sdmflow).

##### Background data

We generated background points for each species equal to twice the number of presences within the accessible area to characterise ranges of low presence probability.

##### Data partitioning

Training data: 75% of total occurrences.

Validation data: We used a holdout approach.

##### Predictor variables

Predictor variables:

- Topography: Digital Elevation Model
- Climatic: monthly minimum and maximum temperature; precipitation; 19 bioclimatic predictors (calculated); 16 environmental variables (calculated).
- Habitat: waterbodies from FAO’s GeoNetwork

Data sources:

- Topography: WorldClim 2 (<https://www.worldclim.org>)
- Climatic: WorldClim 2 (<https://www.worldclim.org>)
- Habitat: waterbodies from FAO’s GeoNetwork (<http://www.fao.org/geonetwork/srv/en/main.home?uuid=ba4526fd-cdbf-4028-a1bd-5a559c4bff38>)

Spatial extent: -180, 180, -90, 90 (xmin, xmax, ymin, ymax)

Spatial resolution:

- Topography: ~10 km (0.08333333)

- Habitat: ~1 km (0.008333333)

Coordinate reference system: EPSG:4326 - WGS 84 - Geographic

Temporal extent: Average for the period 1970-2000

Temporal resolution: Monthly minimum and maximum temperature; precipitation

Data processing: We adjusted the resolution of spatial layers waterbodies to 5 minutes using the function *resample* from the package *raster* setting ‘bilinear’ method.

##### Transfer data

Equal to predictor variable section**.**

### Model

##### Multicollinearity

We estimated pairwise Pearson’s r correlation and calculated a dendrogram to visualize the multicollinearity effect among our 35 predictor variables (Dormann et al., 2013). We selected the final predictor variables extracting the layers with a |r| < 0.5 from the dendrogram and then removing the variables with a Variance Inflation Factor (VIF) > 3.

##### Model settings

**GAM:**

- Formula: presencia ~ s(embergerQ) + s(Waterbodies_5_null) + s(elevation_5_null) +

s(bio7) + s(bio8) + s(bio10) + s(bio15)

- Example weight: "Presence points = 1137"

"Weight for presences = 0.000879507475813544"

"Background points = 3032"

"Weight for background = 0.000329815303430079"

gam(formula.gam, family=binomial(link=logit), data=sp, weights= weight)

**BRT:**

gbm.step(data=sp, gbm.x = 5:11, gbm.y = 4, family = "bernoulli", tree.complexity = 5,

learning.rate = 0.001, bag.fraction = 0.5)

**MAXENT:**

maxent(variables.maxent, presence.background.maxent, path = dir.path)

##### Analysis and Correction of non-independence

##### To reduce the spatial autocorrelation, we performed spatial thinning with the function *reduceSpatialCorrelation*.

##### Threshold selection

### We converted the continuous habitat suitability projections into binary maps by using a threshold maximizing the sensitivity (True Positive Rate) and specificity (True Negative Rate).

### Assessment

##### Performance statistics

We compared the model performances using: Area Under the Receiver Operator Curve (AUC) and Boyce index (Hirzel et al., 2006).

We considered predictions with a Boyce < 0.4 as low-quality performance, and we excluded algorithms with AUC < 0.7 in the selection of the best performing model

##### Plausibility check

- Expert judgement

- Outcomes of current alpha taxonomy agree with the distribution map proposed by in Kalkman et al., (2018).

### Prediction

##### Prediction output

Prediction unit: presence and absence
