## Supplementary figures and images for "Climate change will redefine taxonomic, functional, and phylogenetic diversity patterns of Odonata in space and time"

### S5_Dendrogram_collinearity

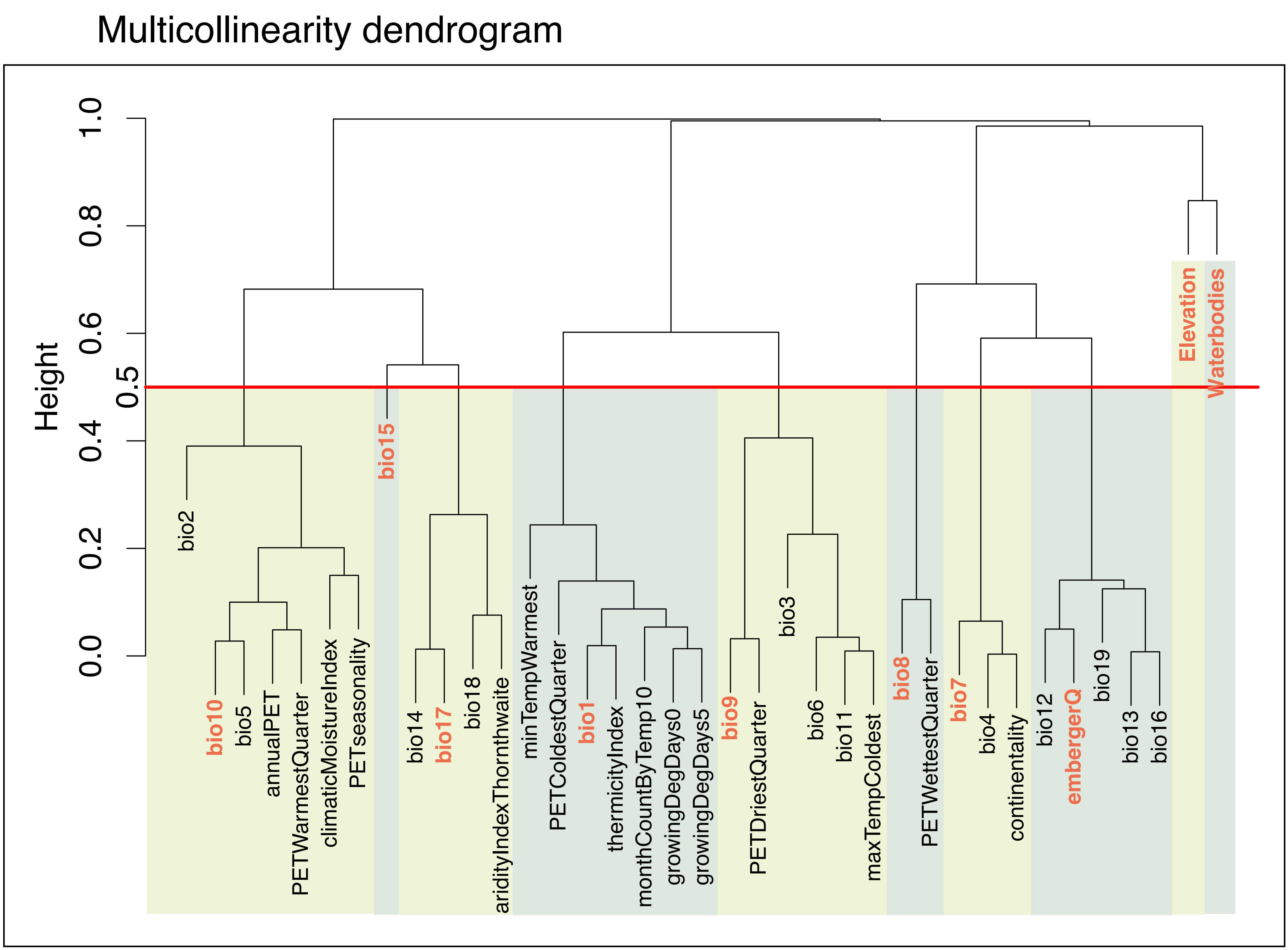


#### Variance Inflation Factor

| | |
|---|---|
| Elevation | 1.59 |
| embergerQ | 2.48 |
| Waterbodies | 1.07 |

| | |
|---|---|
| bio7 | 2.52 |
| bio8 | 1.80 |
| bio10 | 1.41 |
| bio15 | 1.46 |
