## Supplementary material for "Climate change will redefine taxonomic, functional, and phylogenetic diversity patterns of Odonata in space and time": S7_Supplementary_material_Odonata_SDM

#### **Supplementary material B**

Main scheme of the following pictures:

- **Top-left map**: Best model prediction map for the current condition.

- **Top-right maps**: Habitat availability gain (light blue), loss (red) maps and centroid shift future (yellow point) vs present (orange point); time period 2050 (up) and 2070 (down).

- **Bottom-left table**: Lon: Longitude centroid; Lat: Latitude centroid; period: Time period; avgAltitude: average altitude of the predicted binary map; cellNumber: Number of cells of the predicted binary map; cellArea: total cell area of the predicted binary map; cellDifArea: Difference of area between future and present prediction; direction: direction of centroid shift (in degree, for further details see the R function XXX); dist_km: Shift of centroid comparing future and present prediction (in kilometres; for further detail see the R function XXX).

- **Bottom-right plot**: Violin plot shows the elevation shift across time periods.

### **Anisoptera**

##### **Family: Aeshnidae**

##### *Aeshna affinis* Vander Linden, 1820

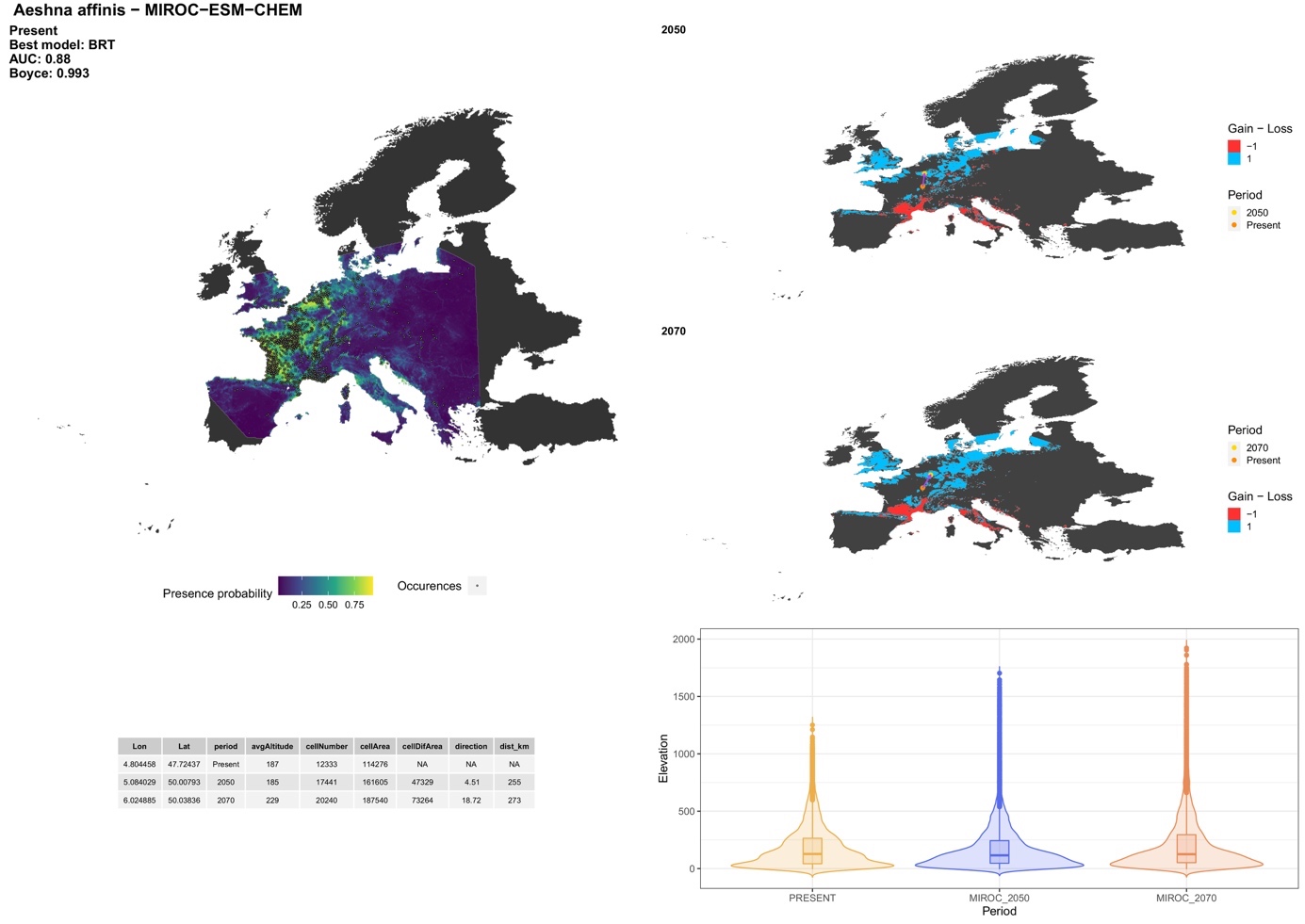

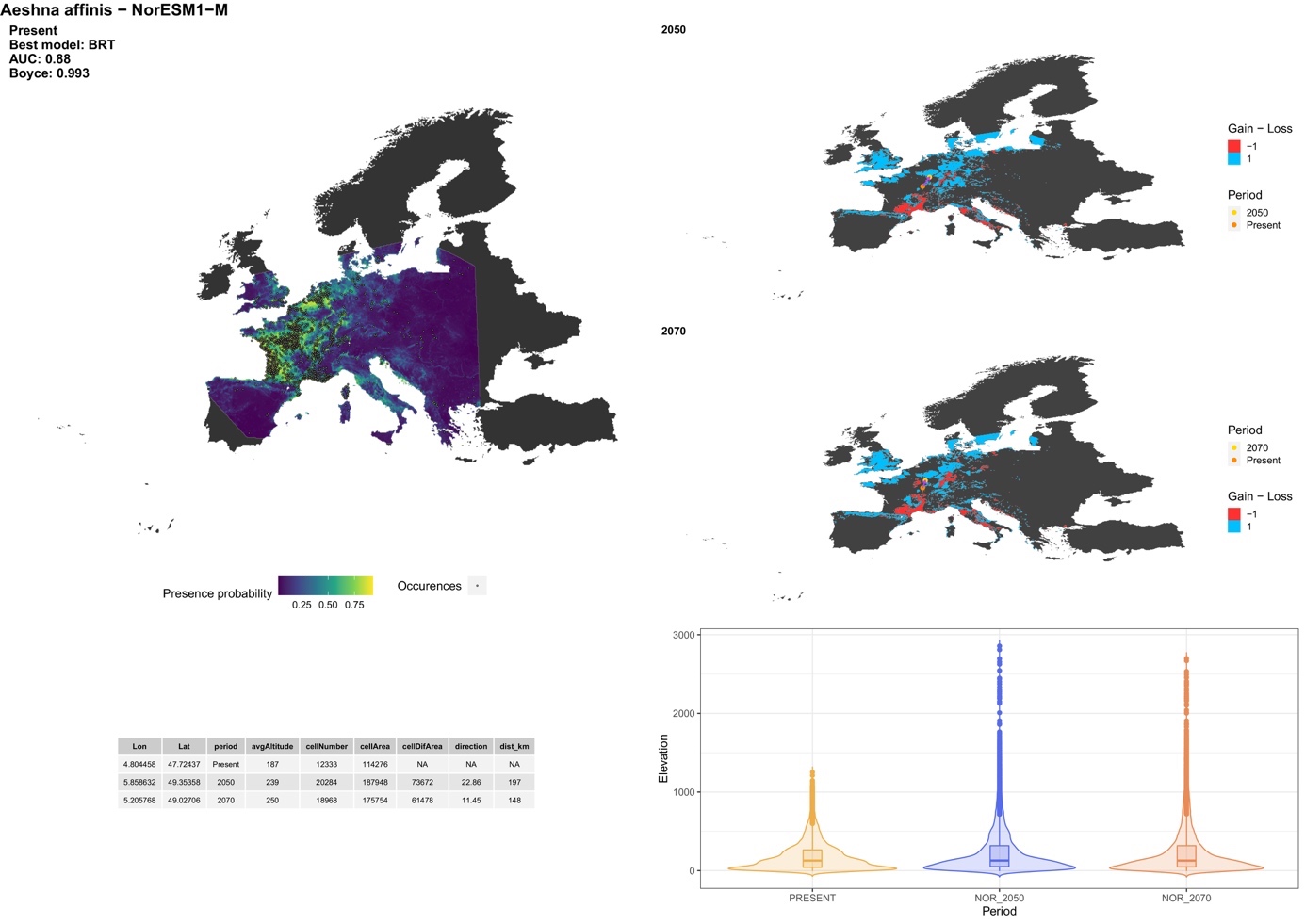

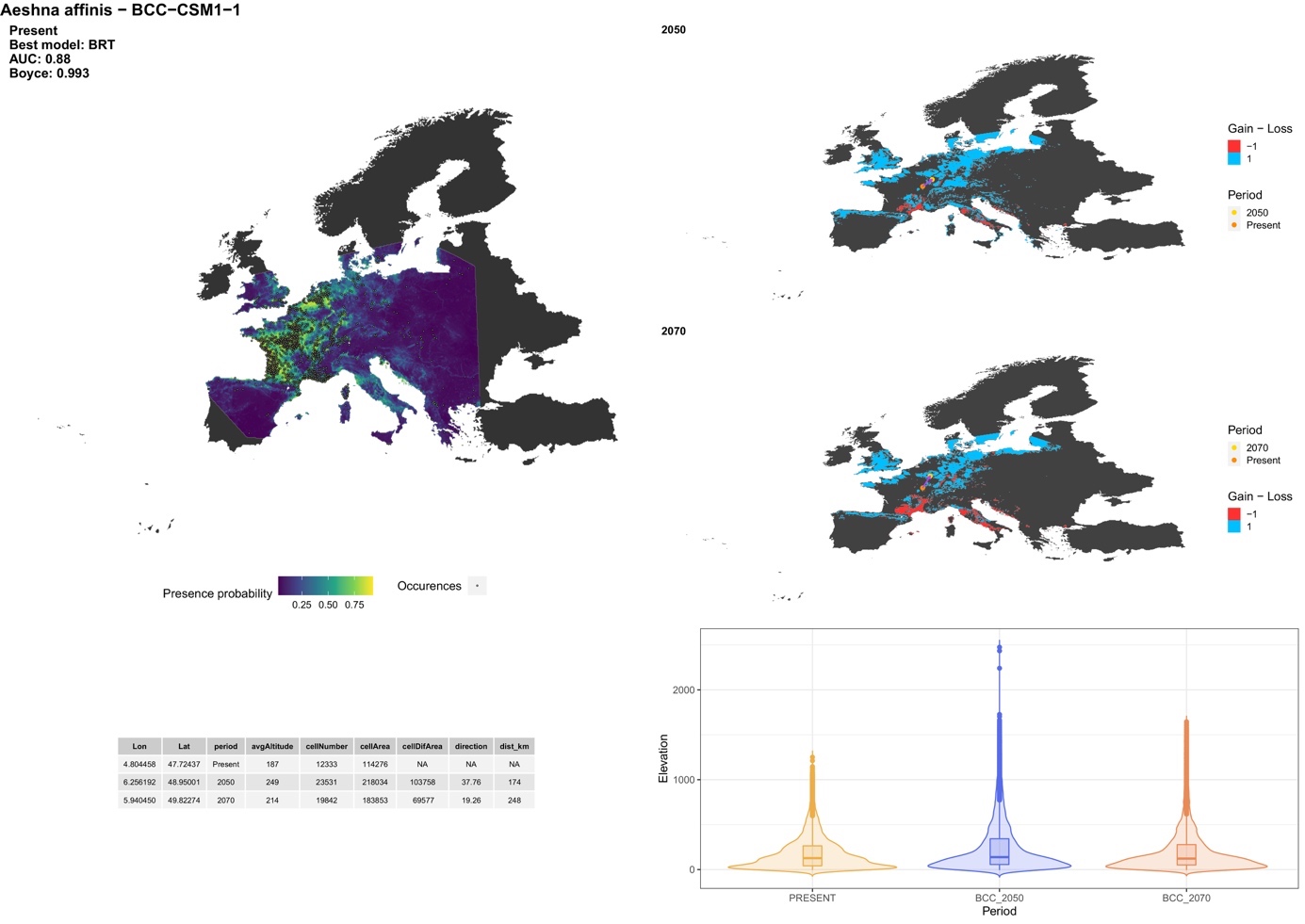

##### *Aeshna caerulea* (Ström, 1783)

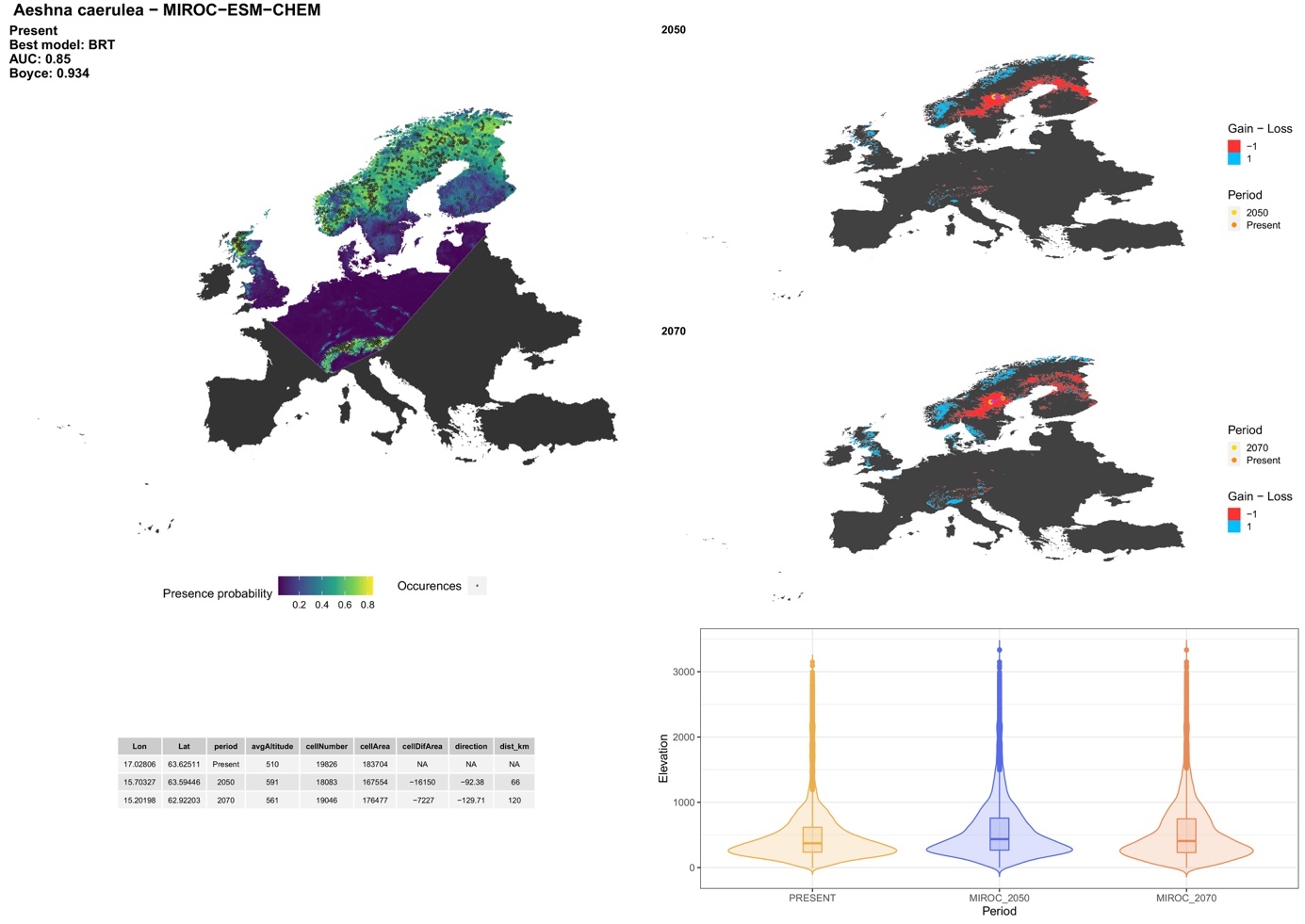

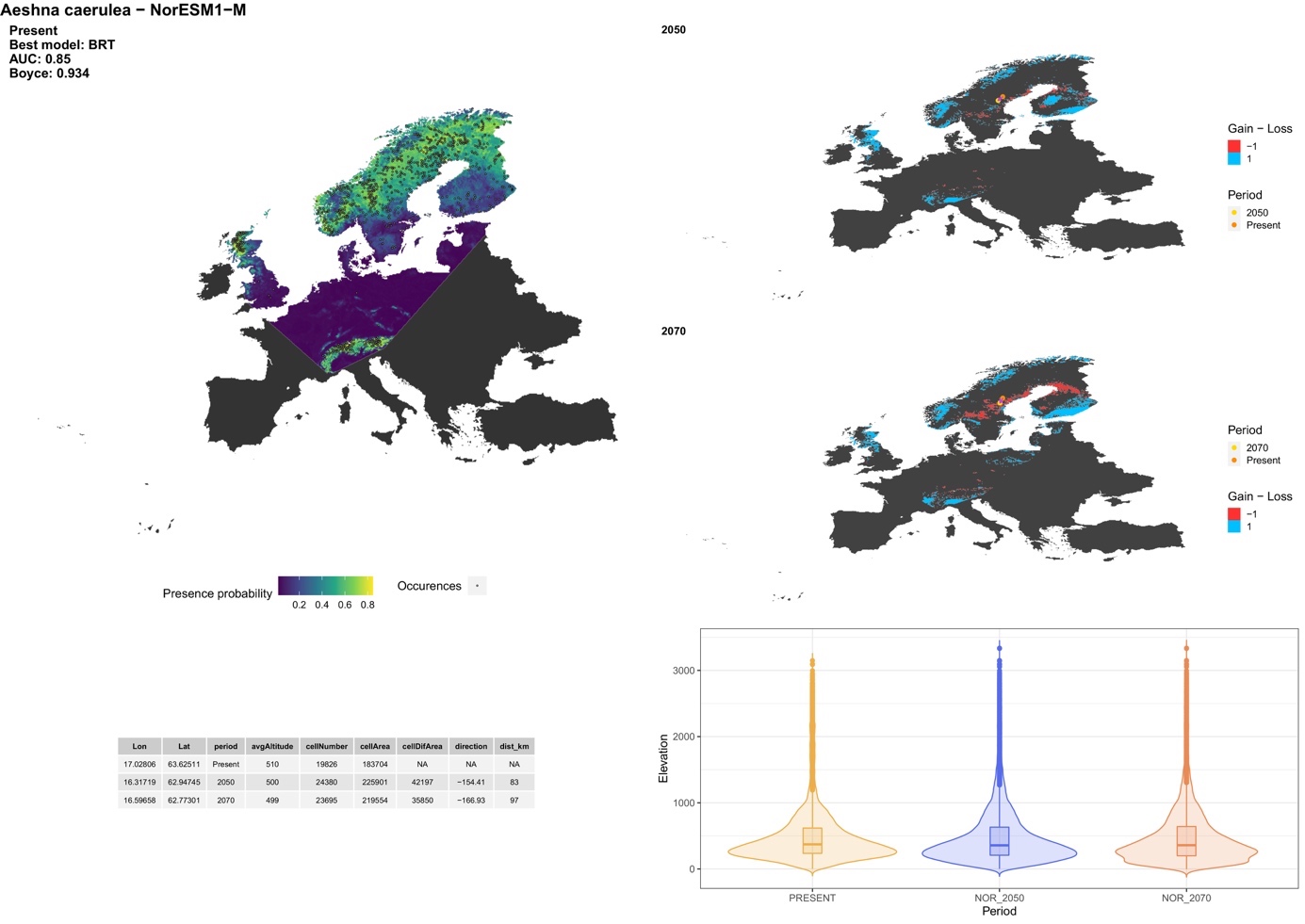

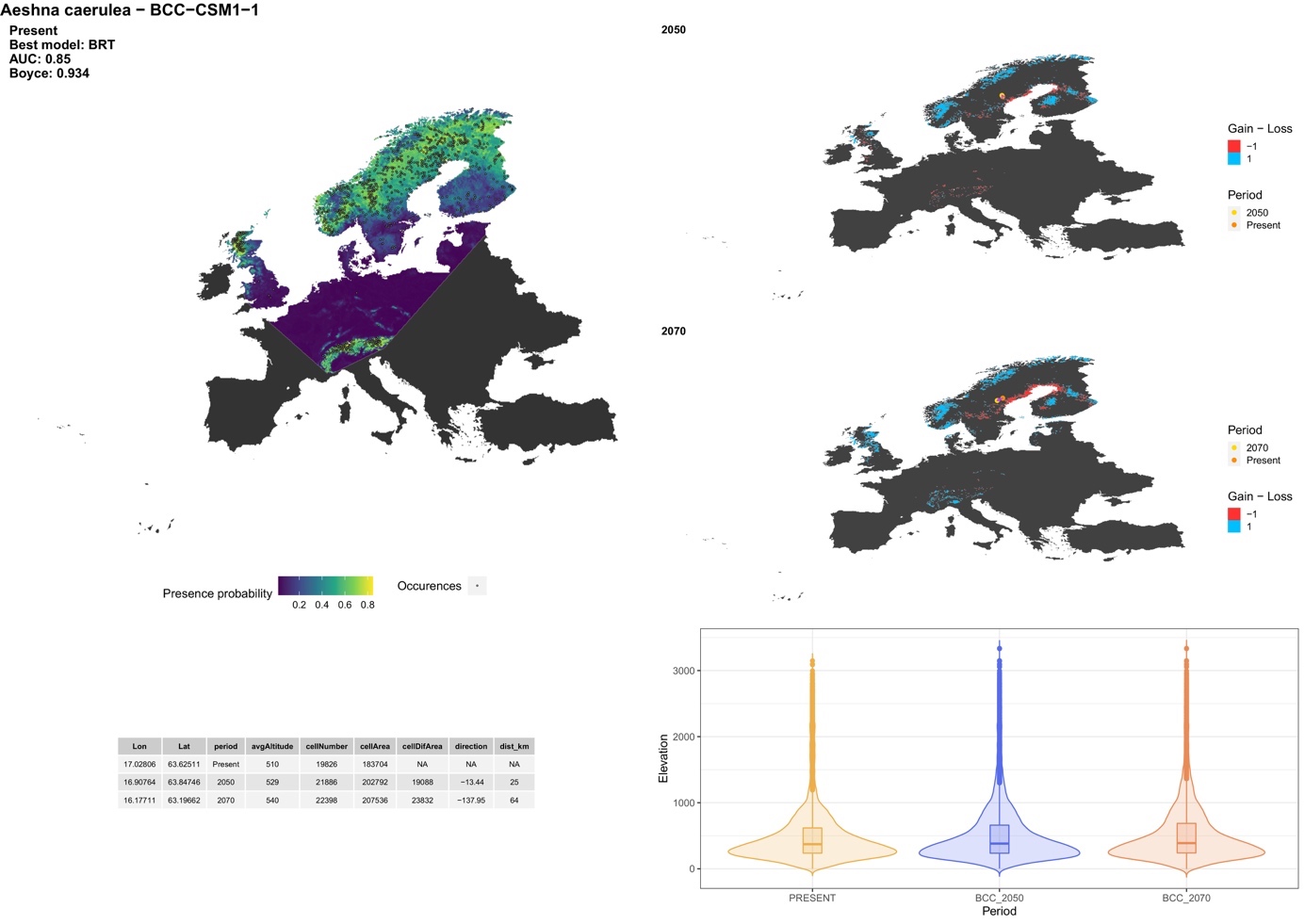

##### *Aeshna crenata* Hagen, 1856

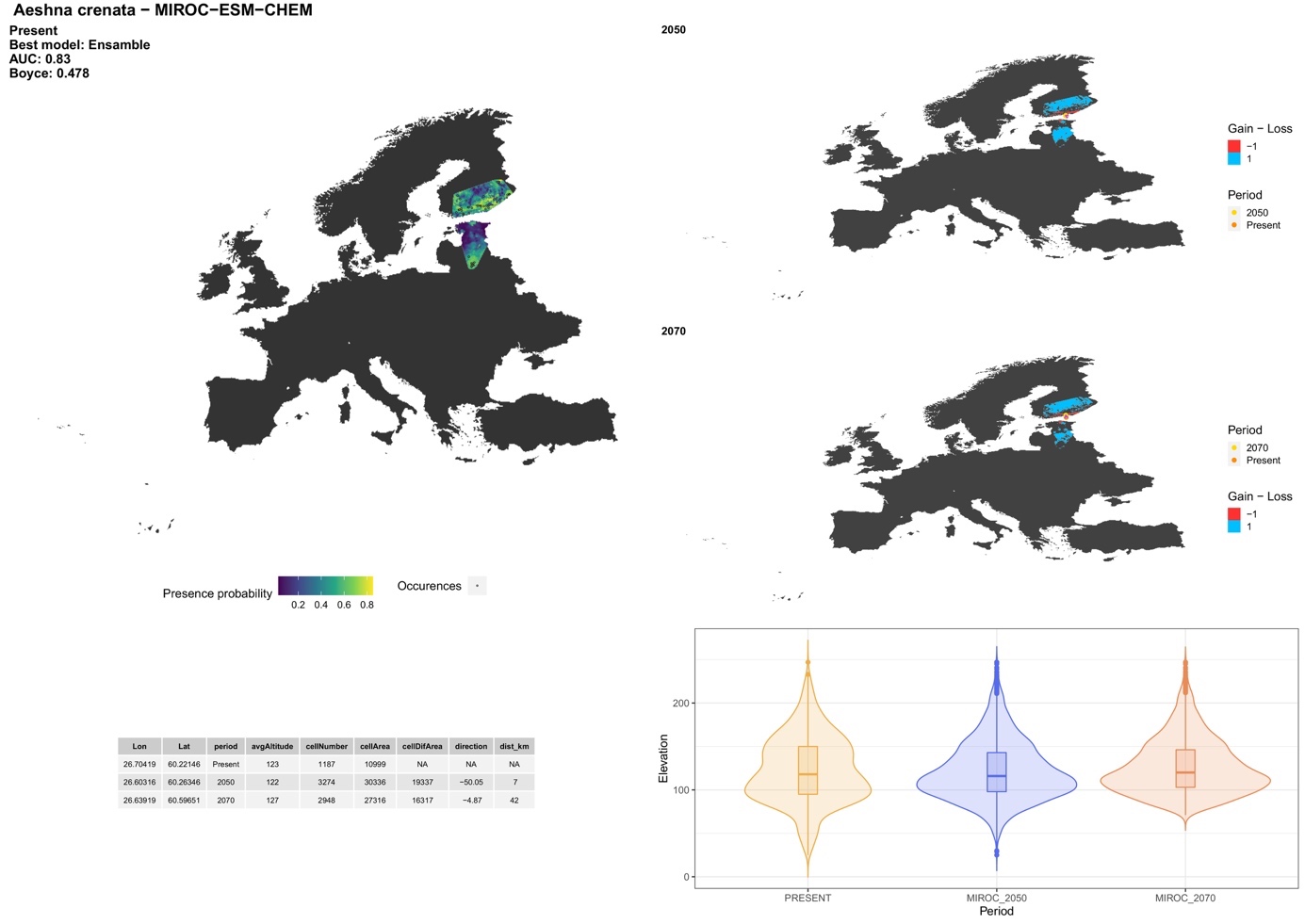

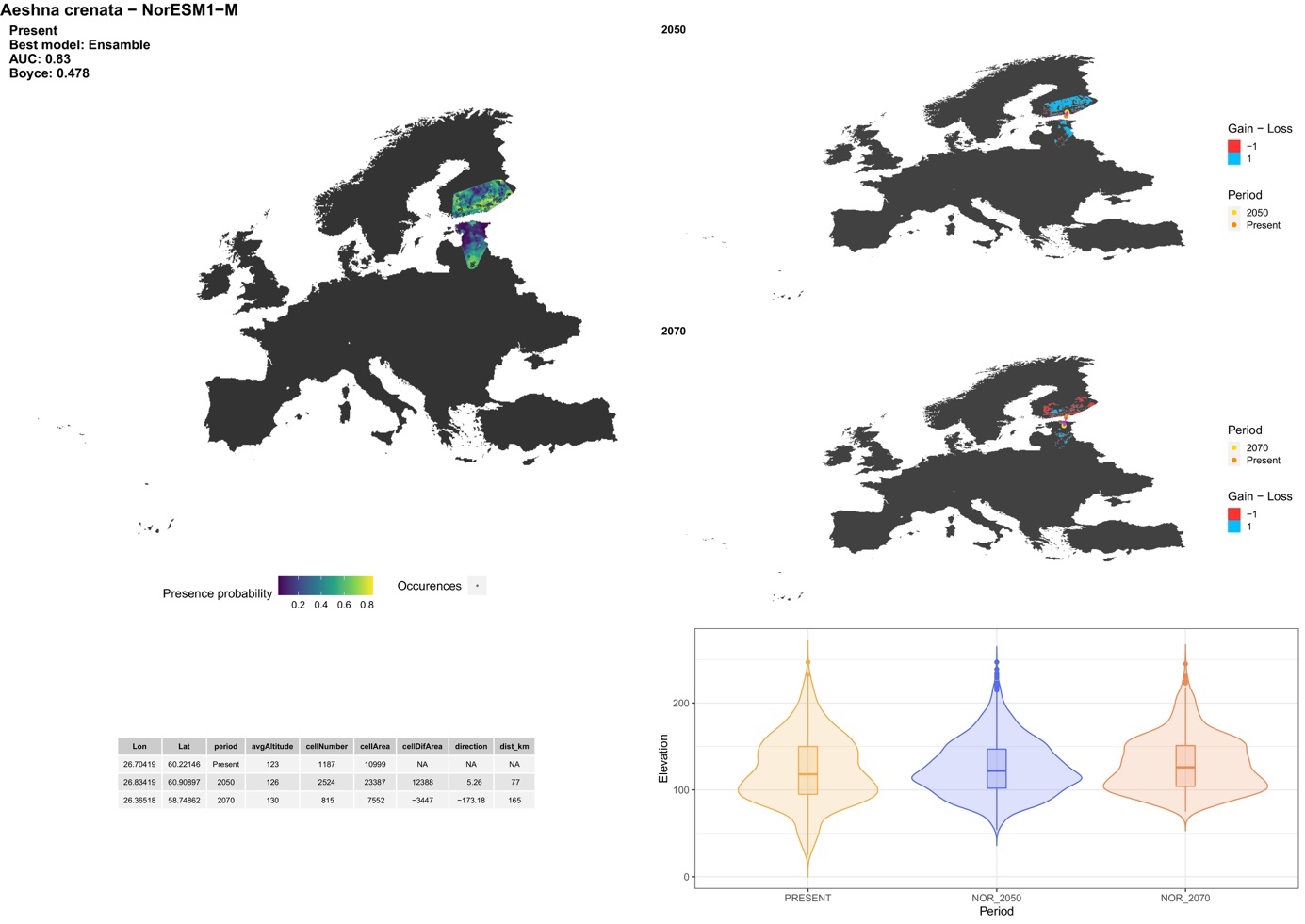

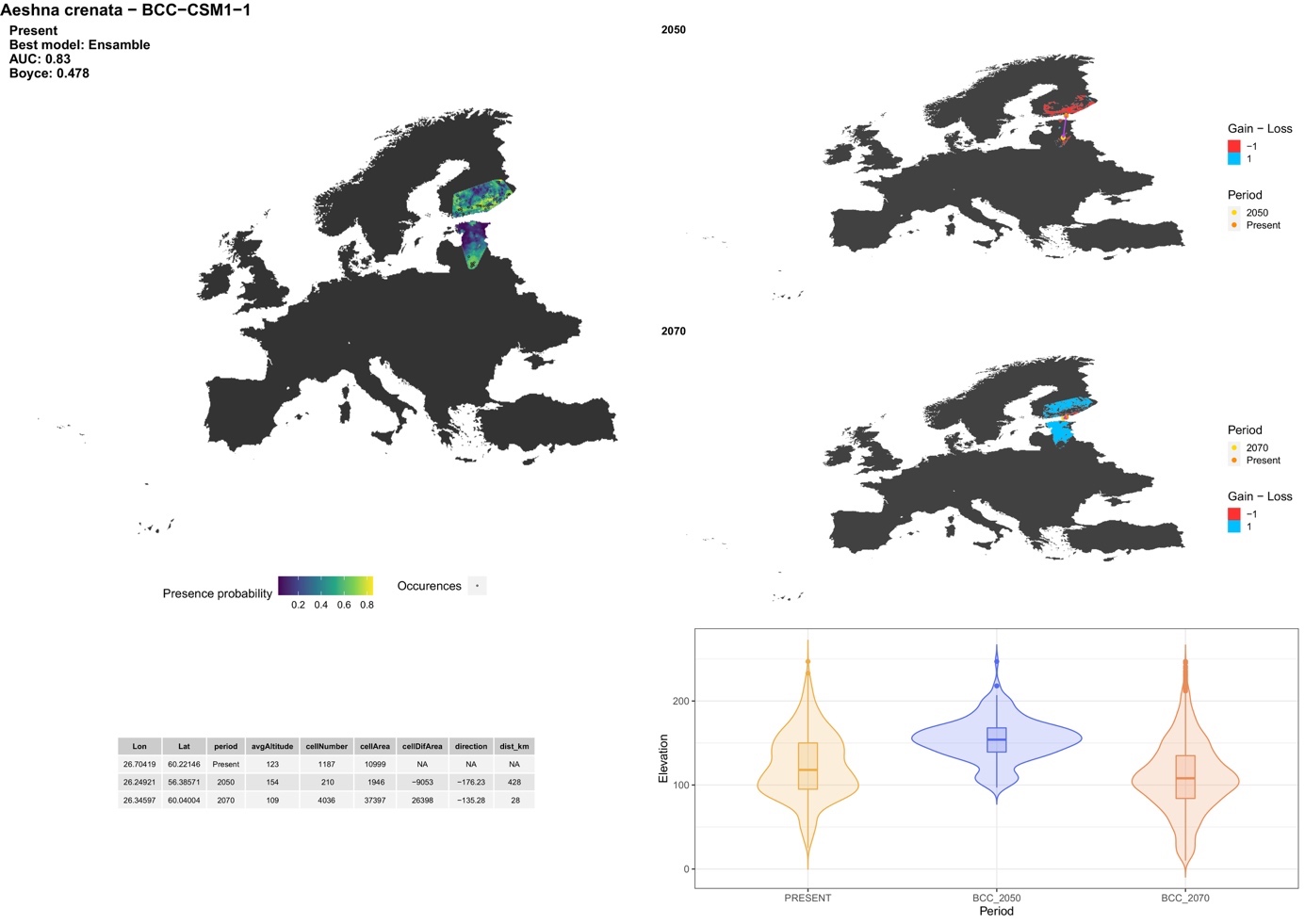

##### *Aeshna cyanea* (Müller, 1764)

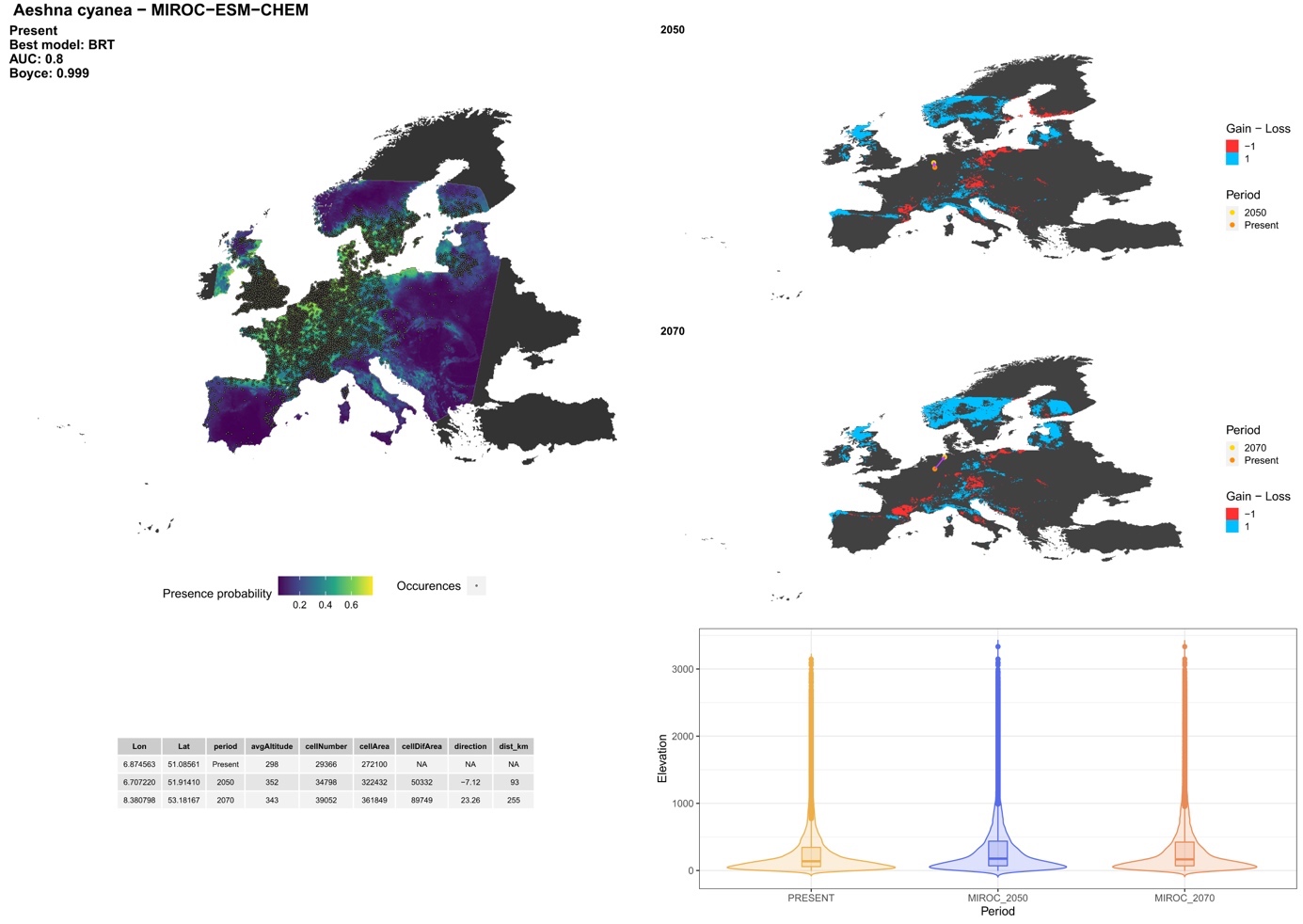

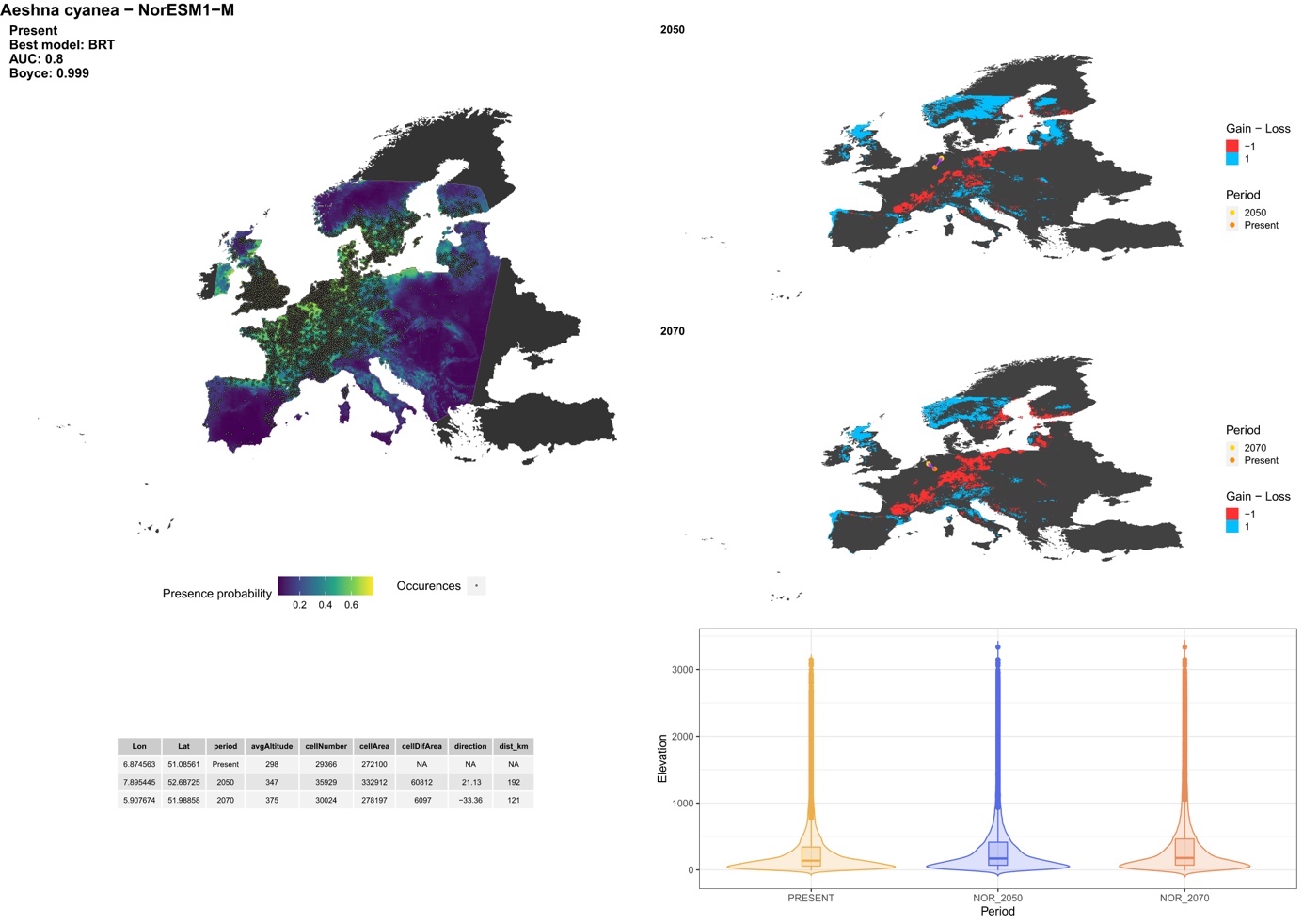

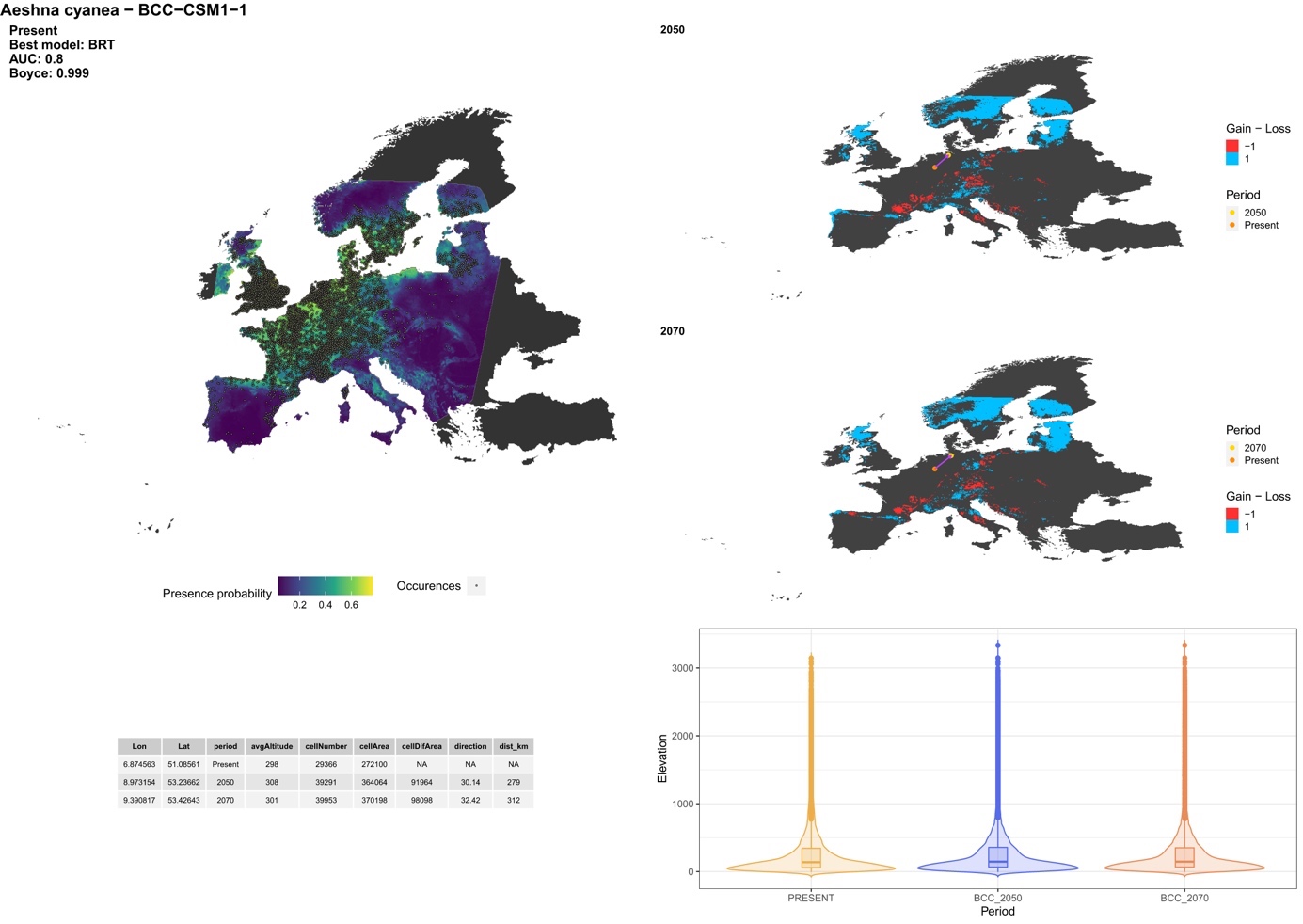

##### *Aeshna grandis* (Linnaeus, 1758)

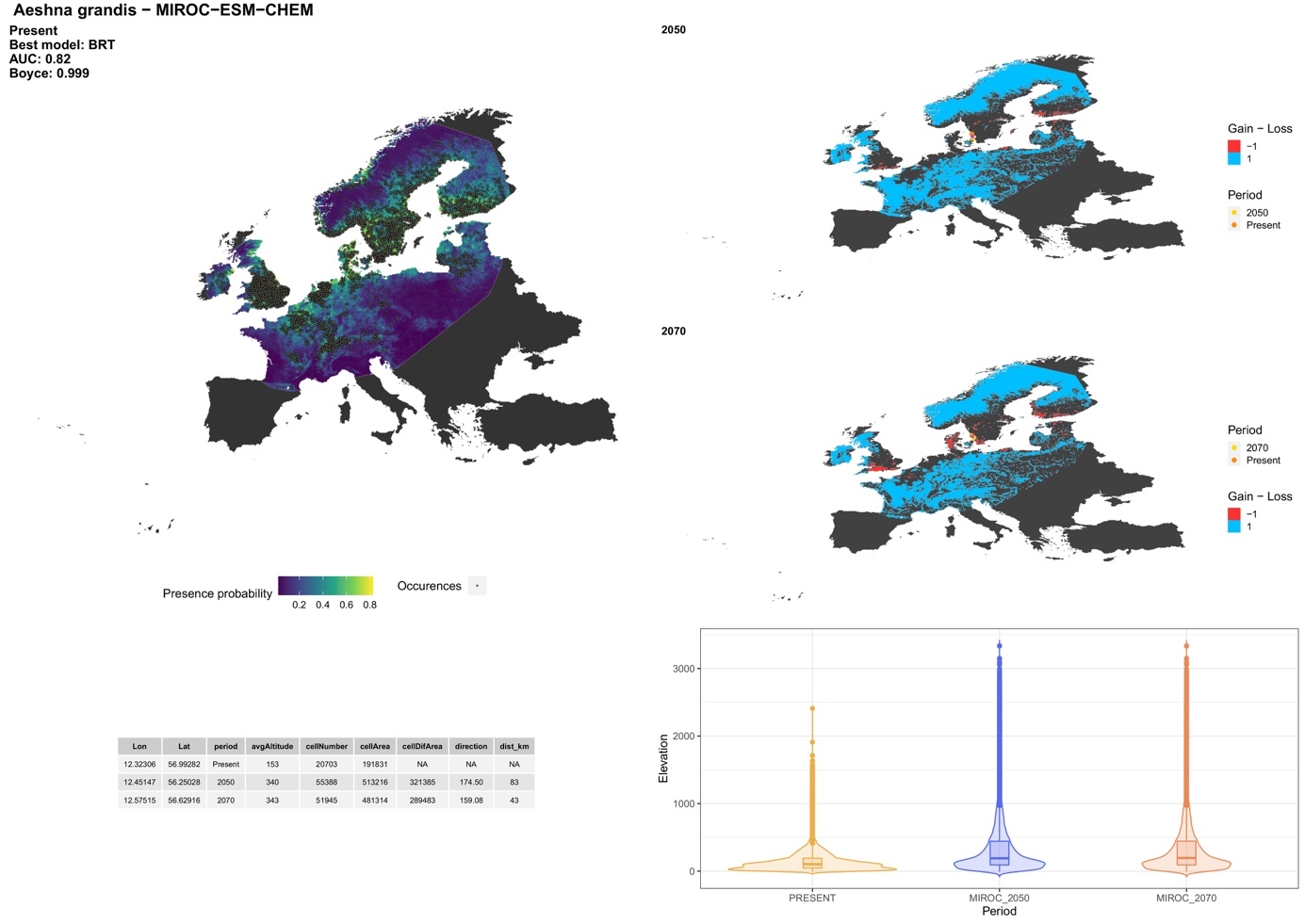

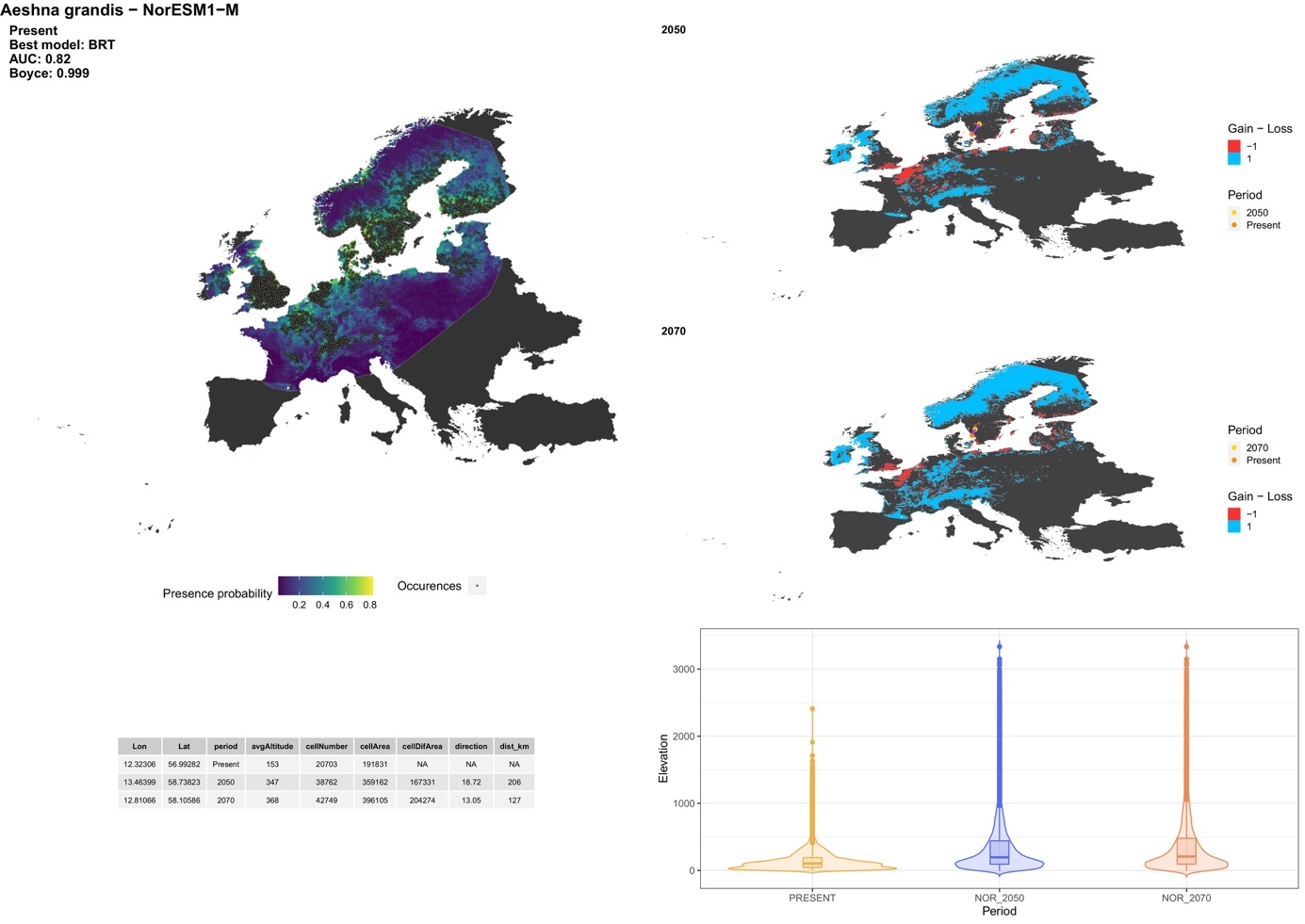

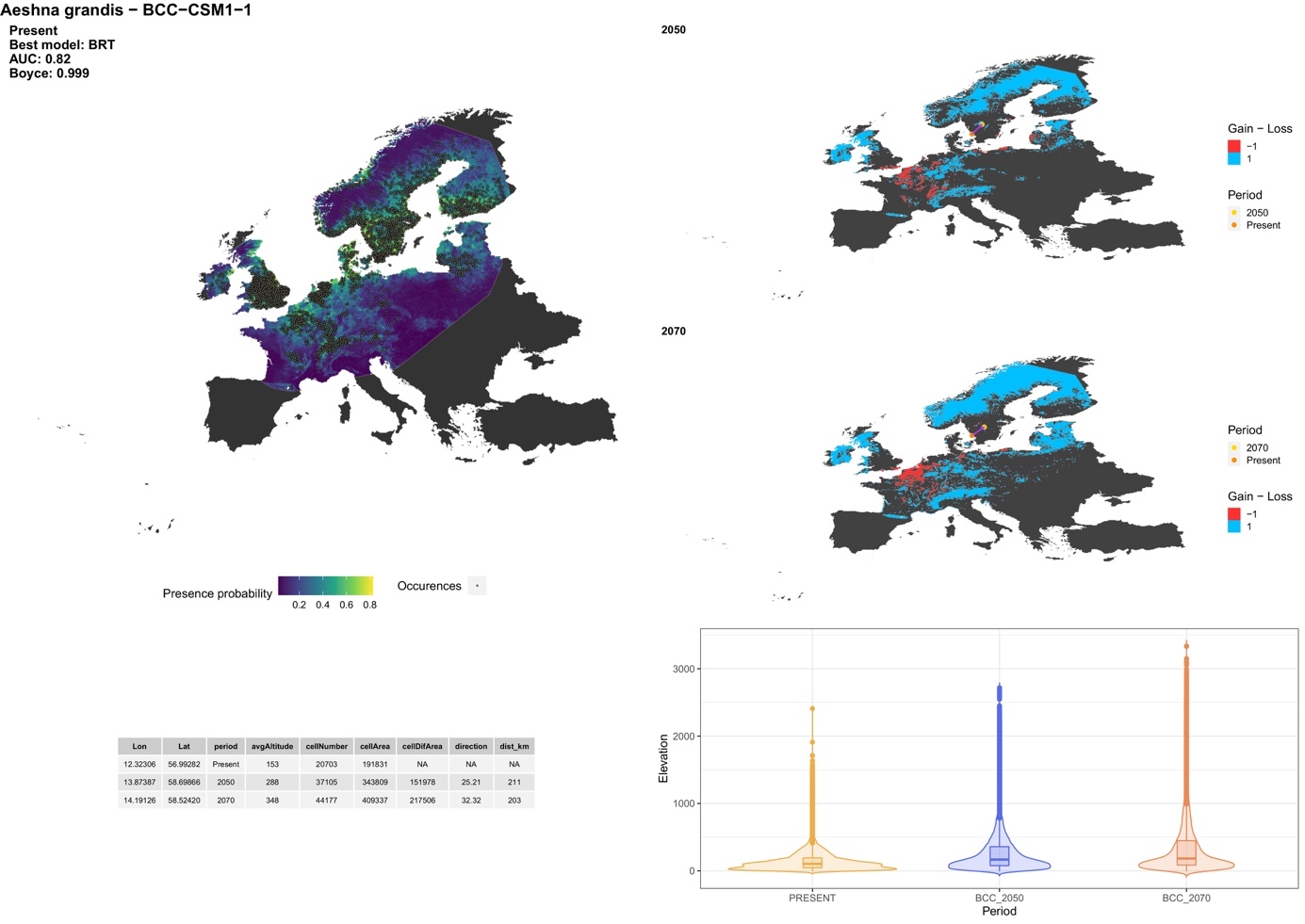

##### *Aeshna isosceles* (Müller, 1767)

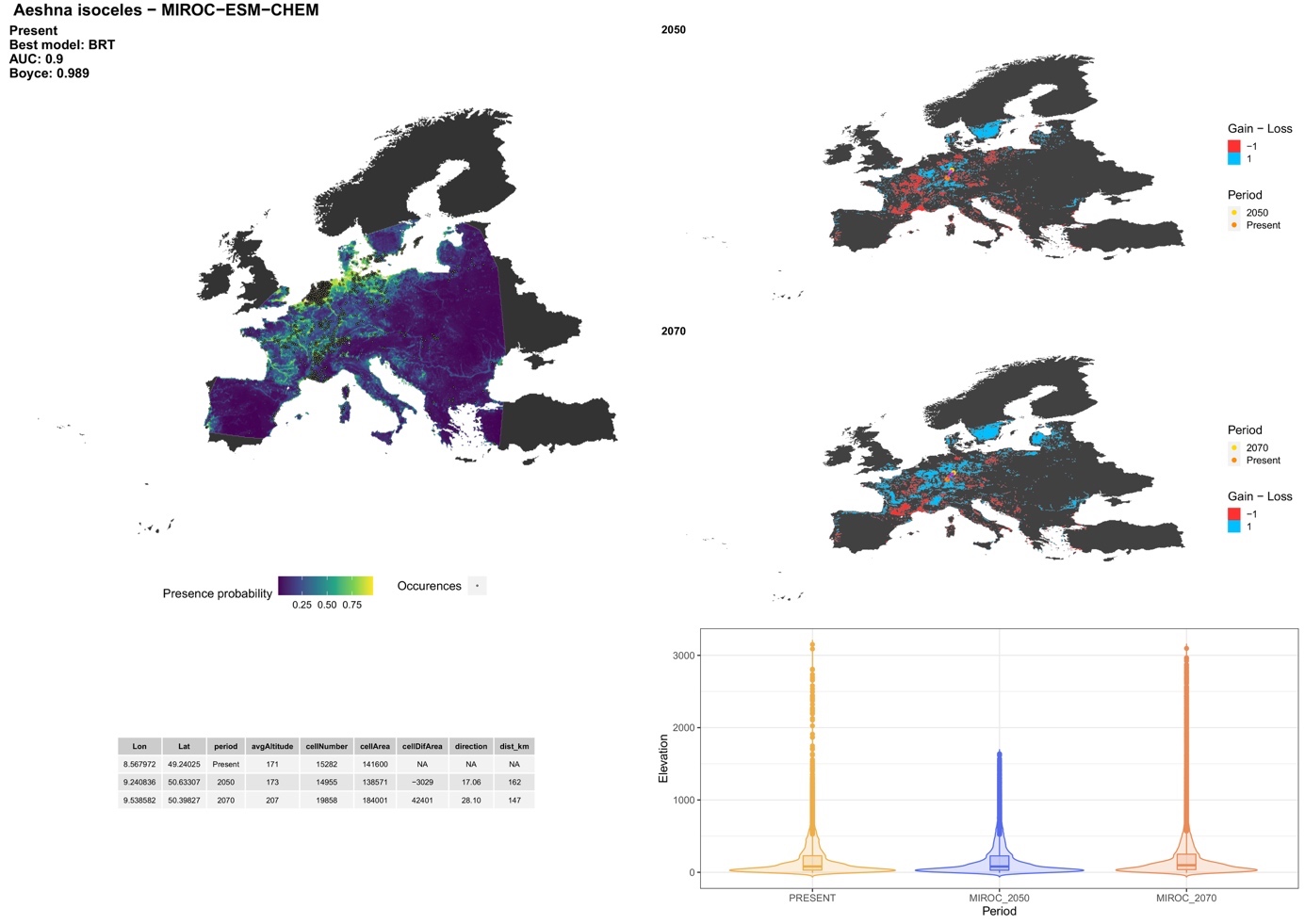

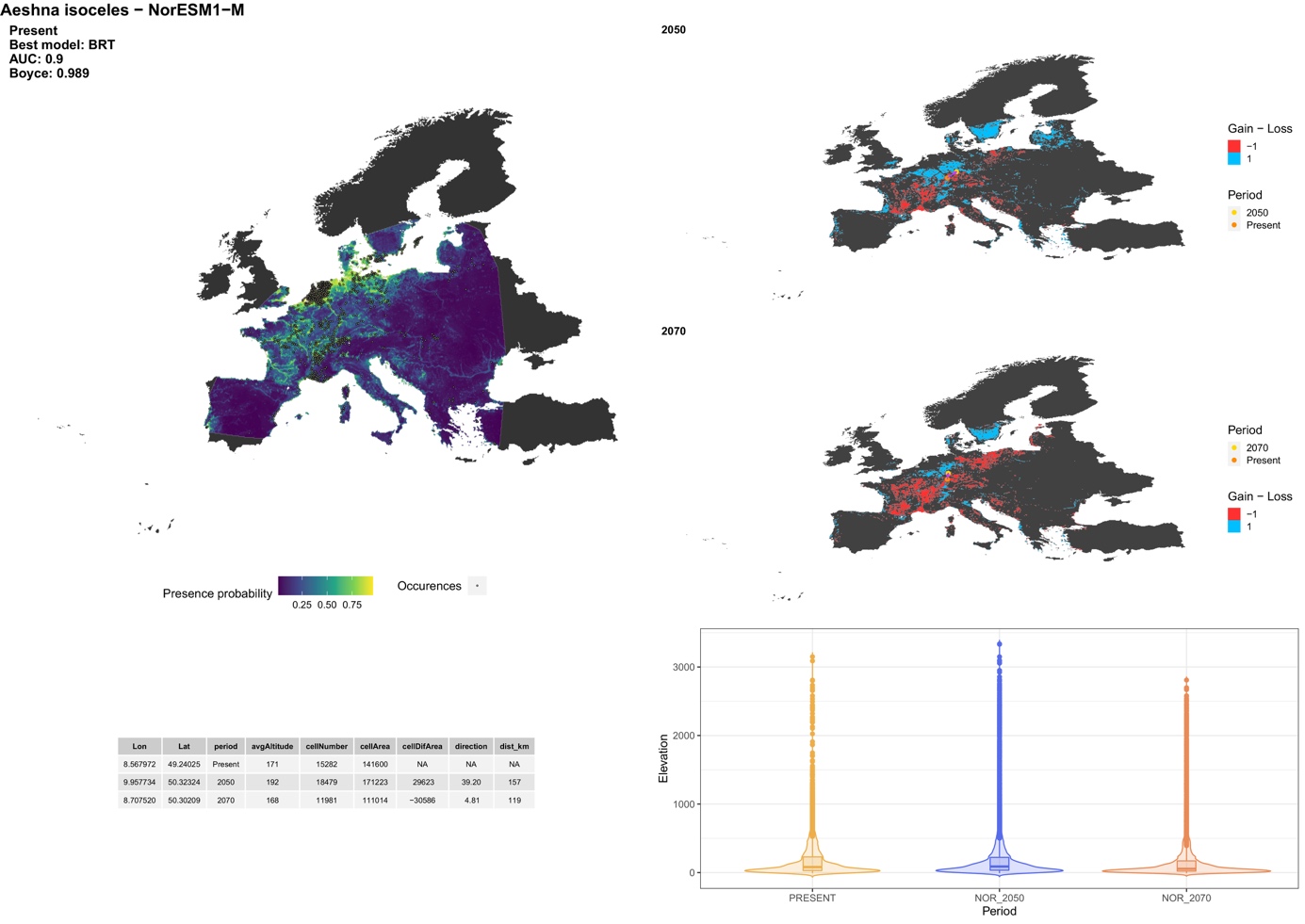

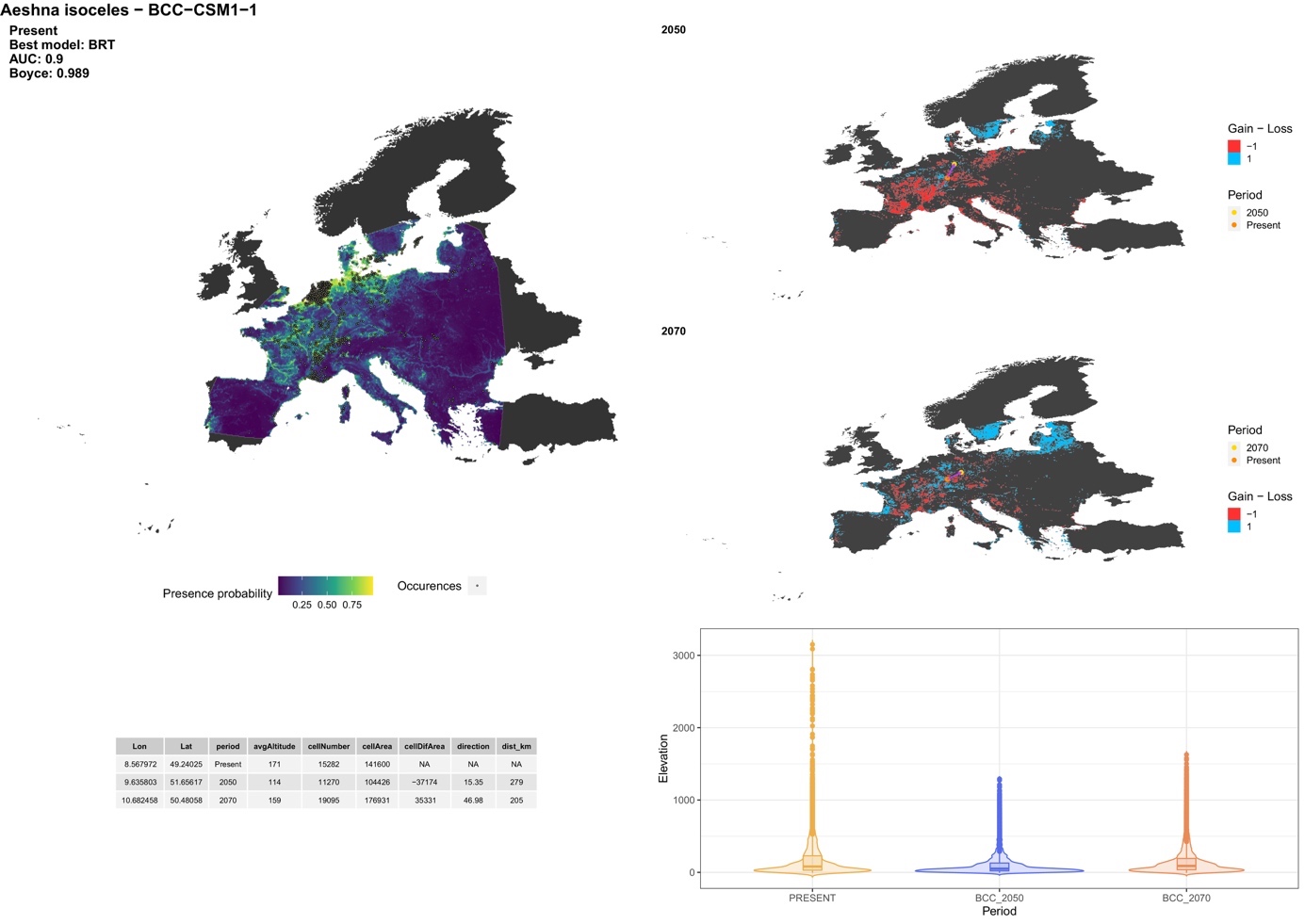

##### *Aeshna juncea* (Linnaeus, 1758)

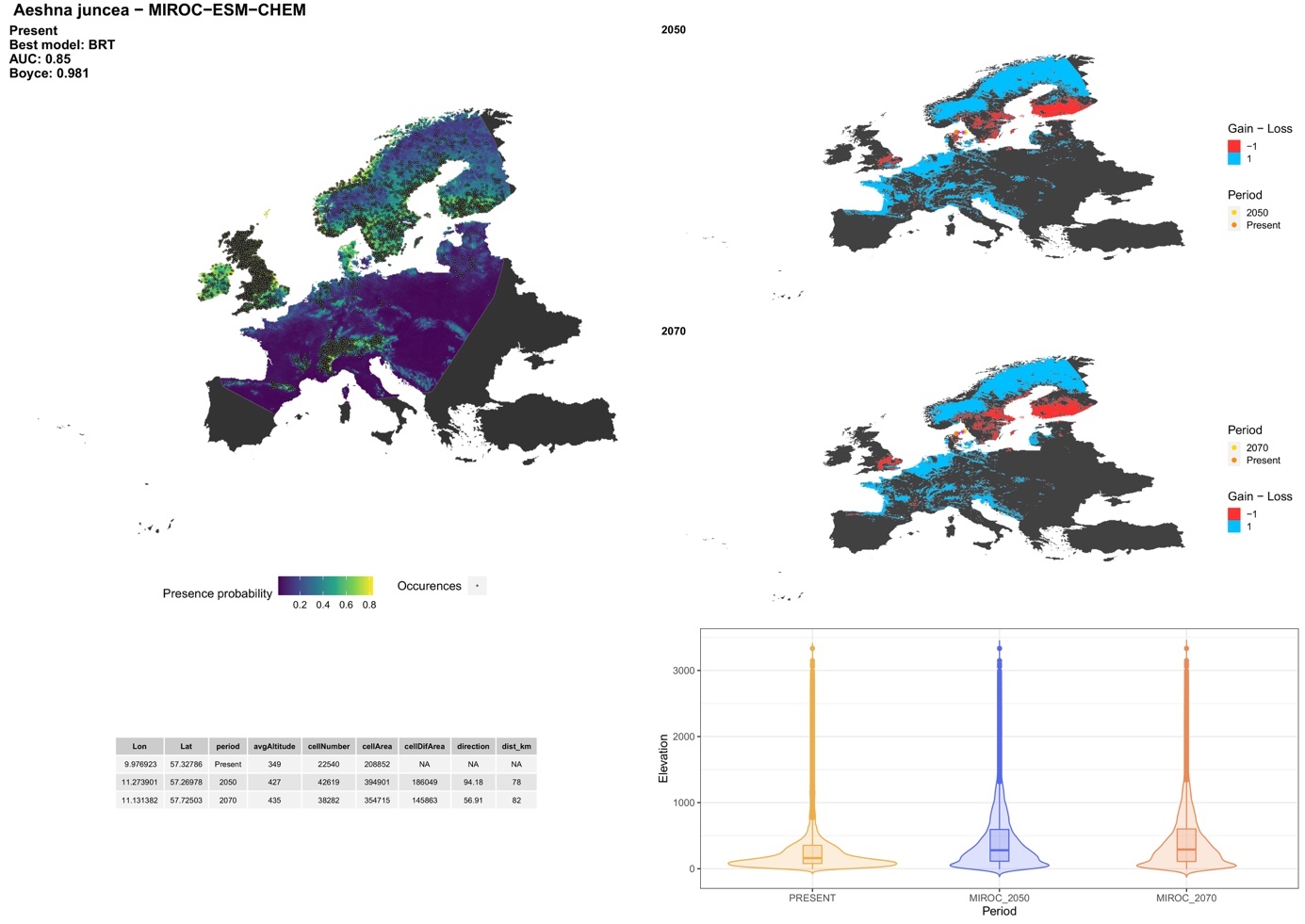

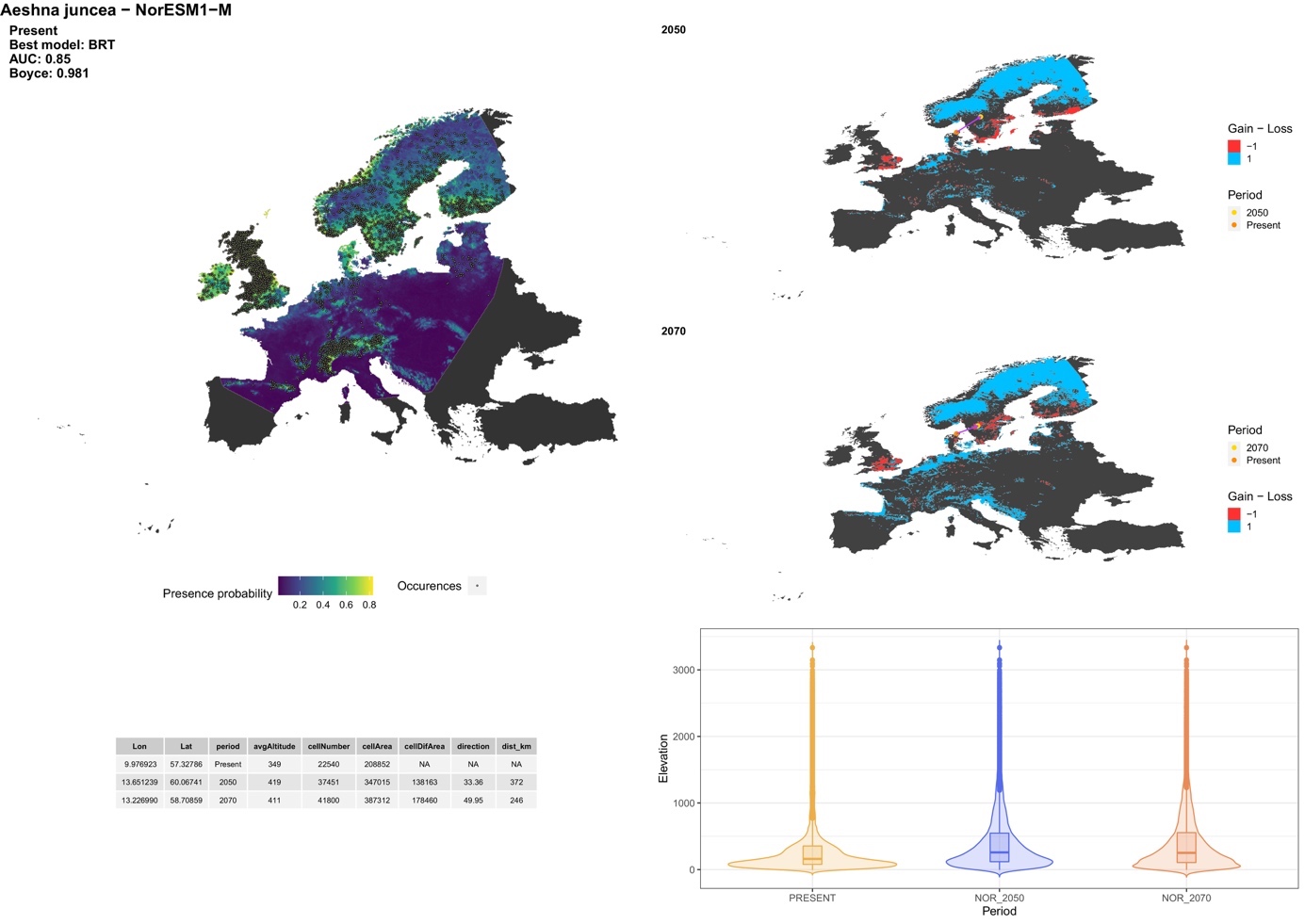

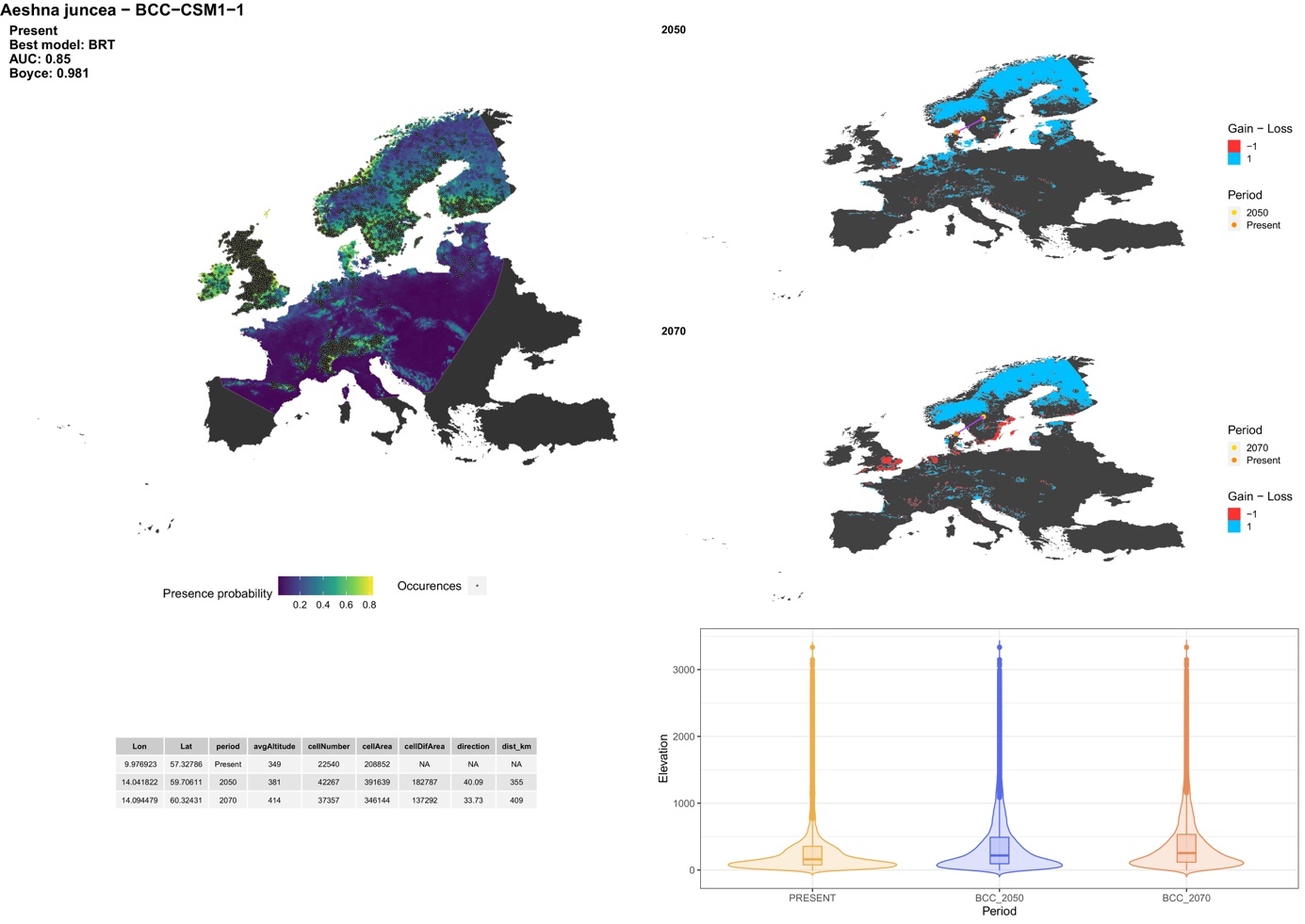

##### *Aeshna mixta* Latreille, 1805

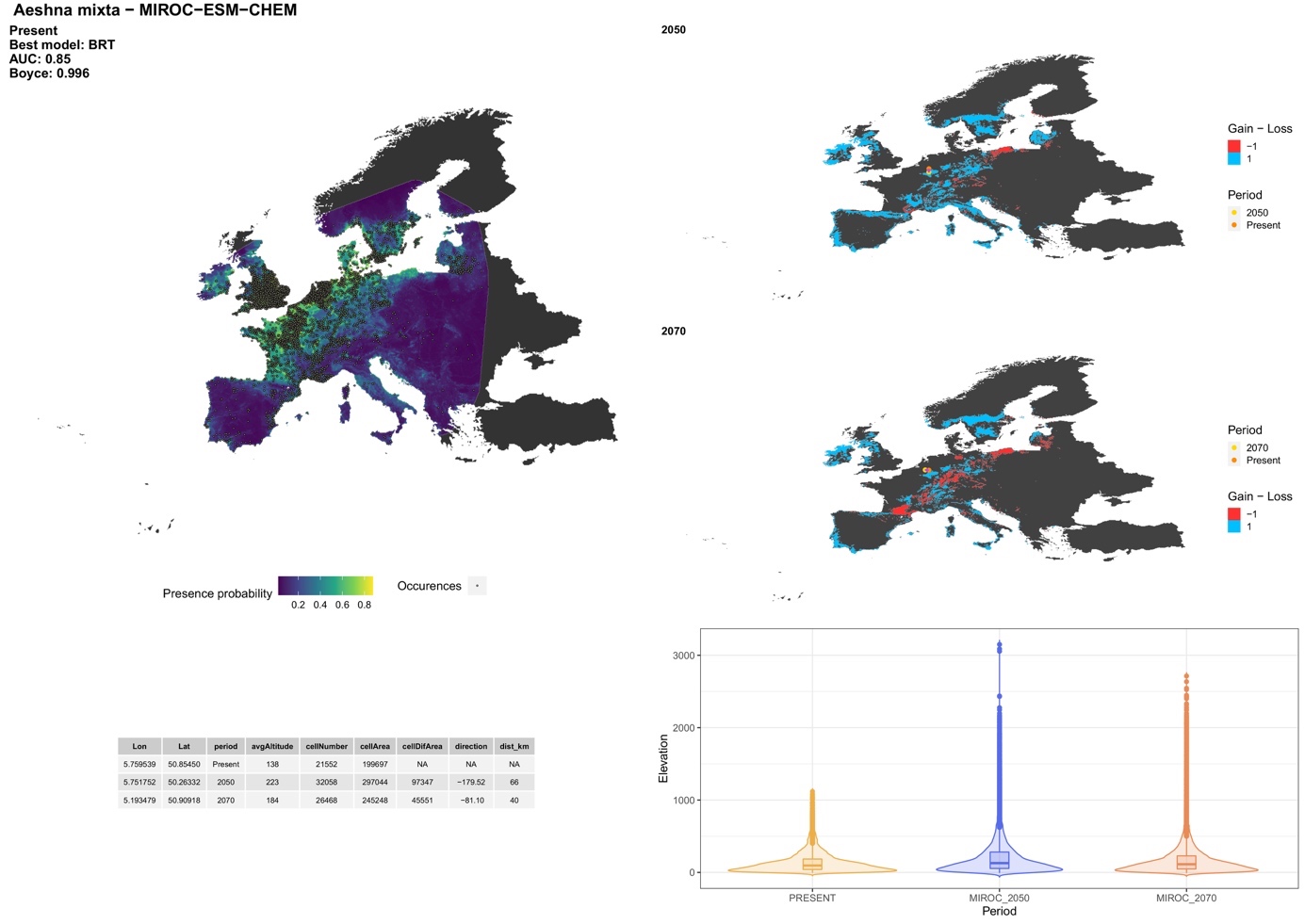

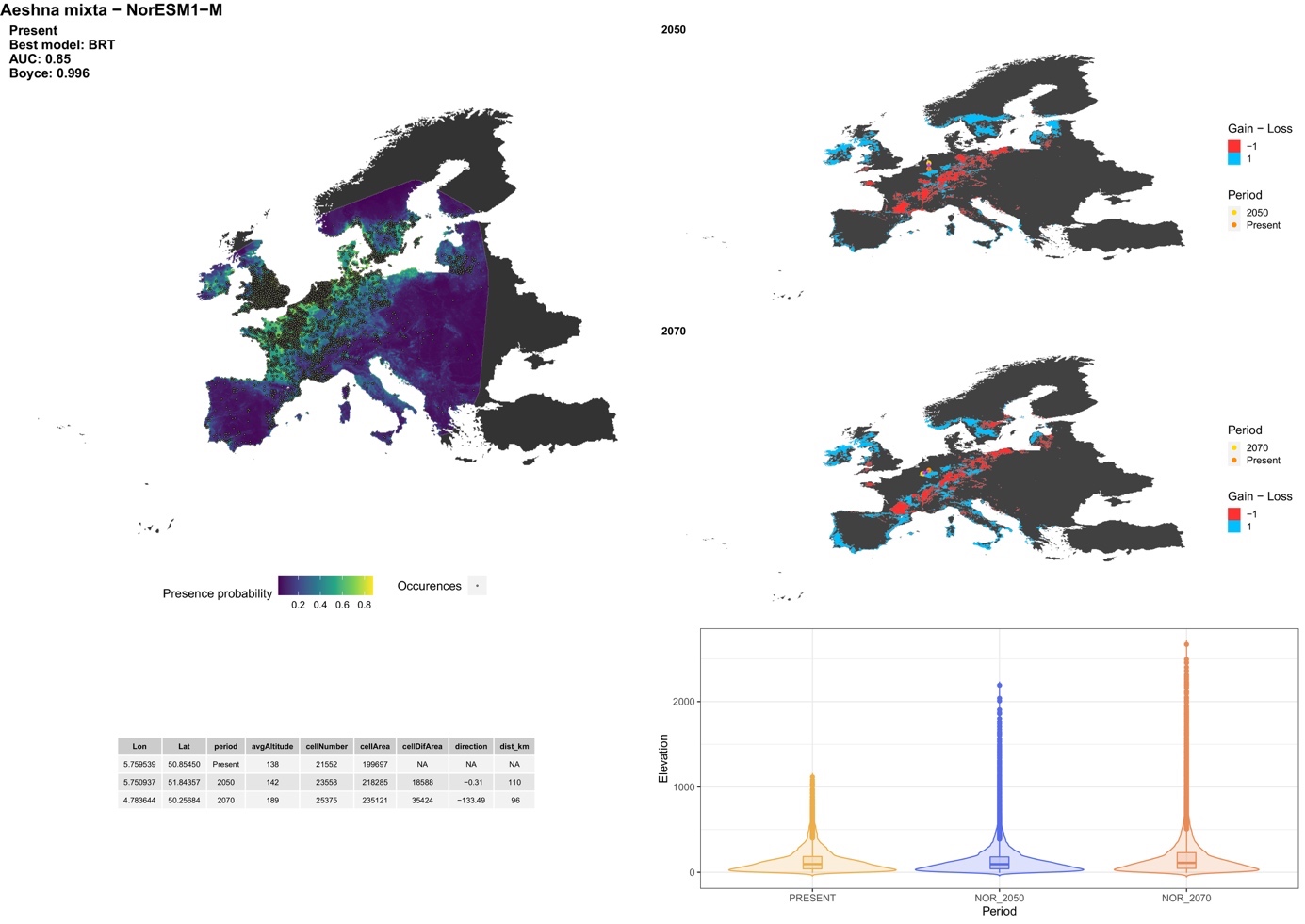

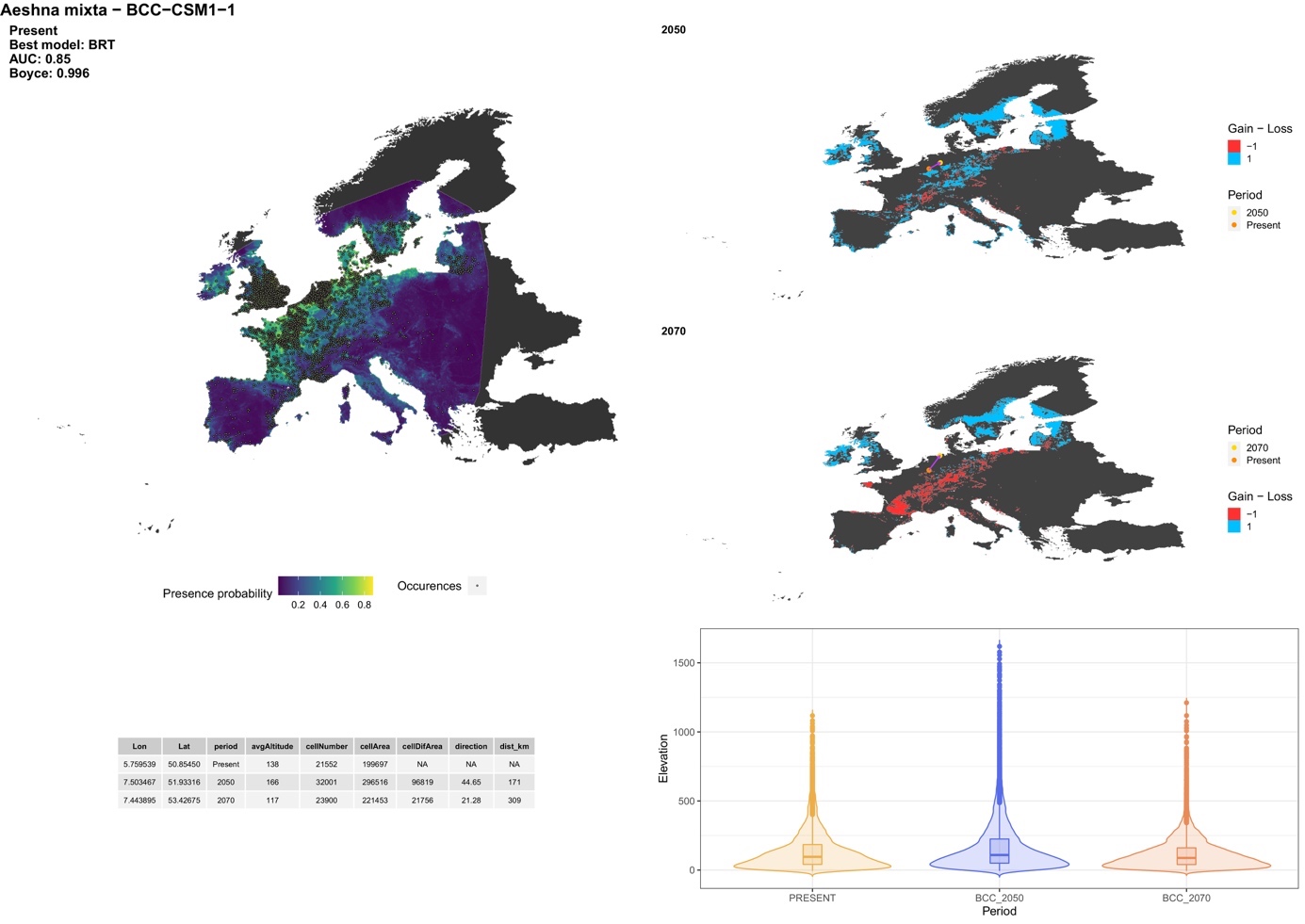

##### *Aeshna serrata* Hagen, 1856

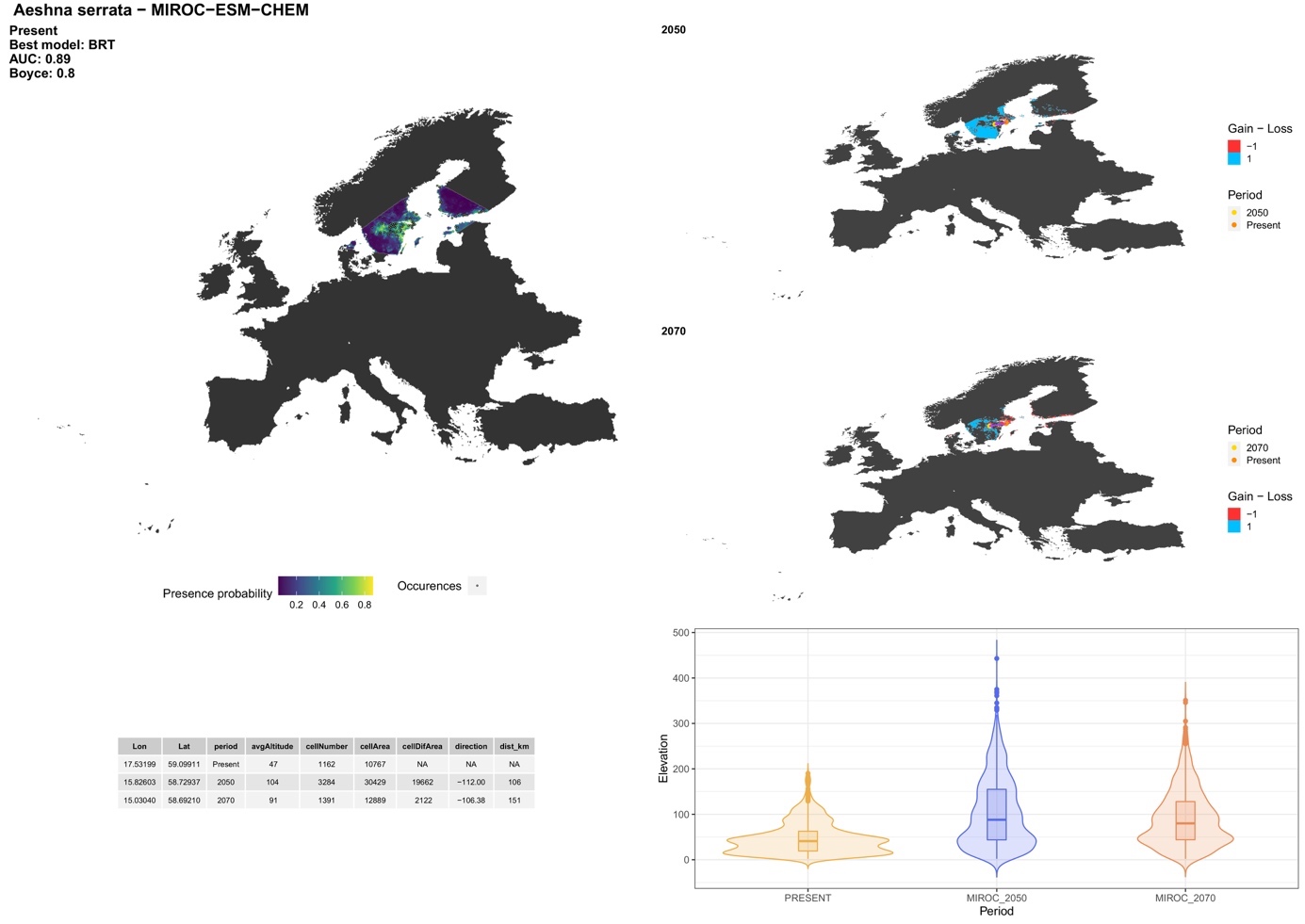

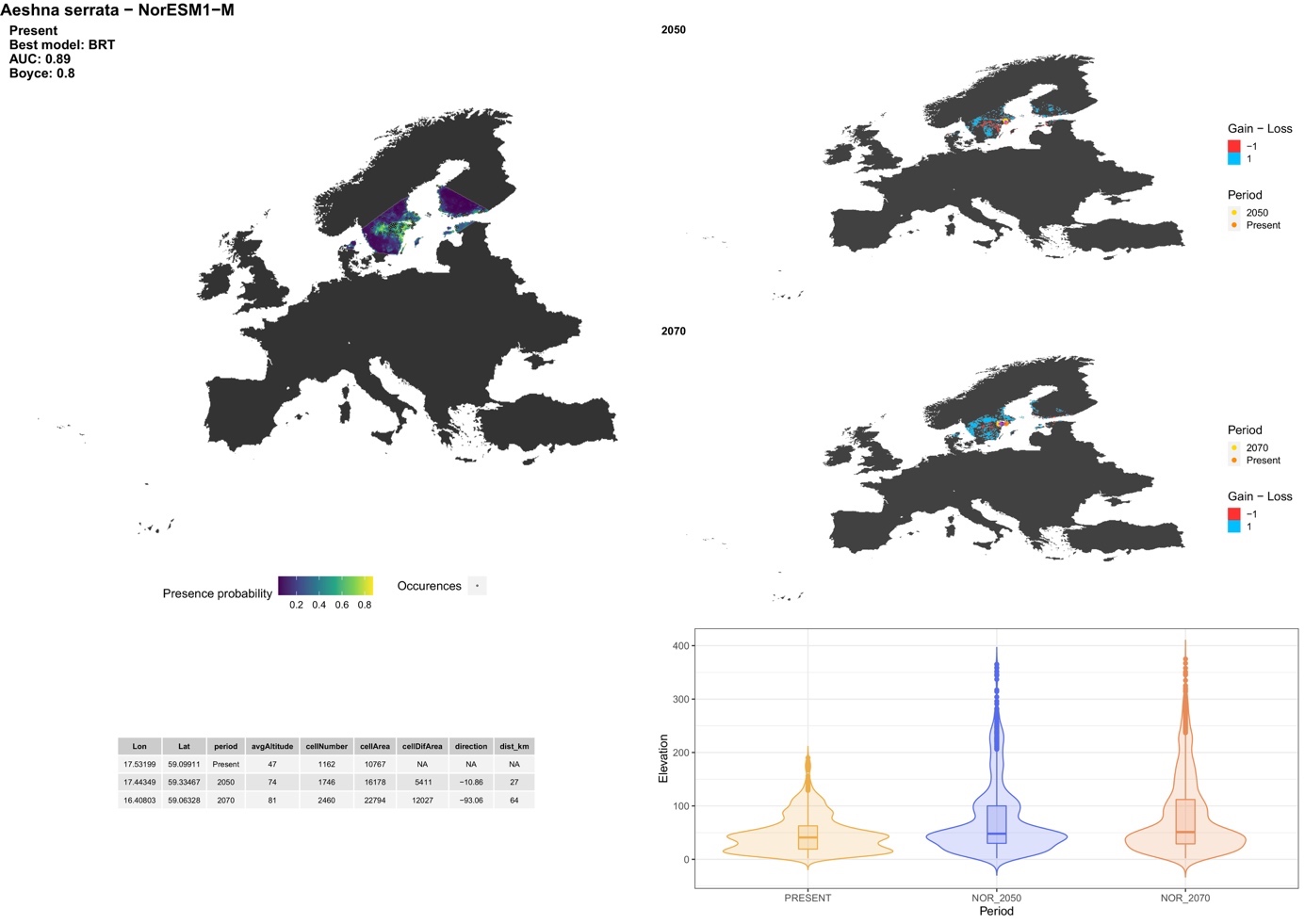

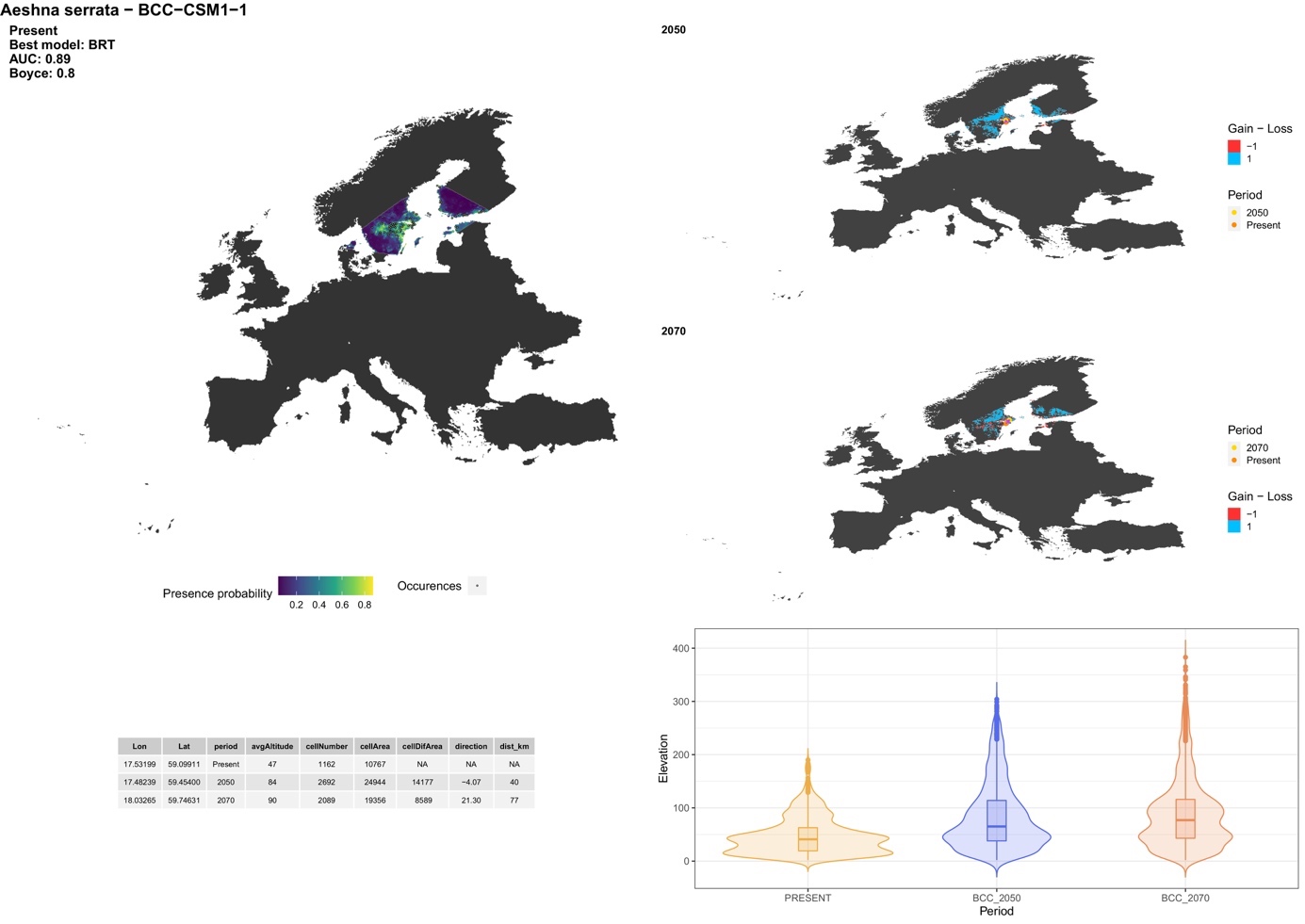

##### *Aeshna subarctica* Walker, 1908

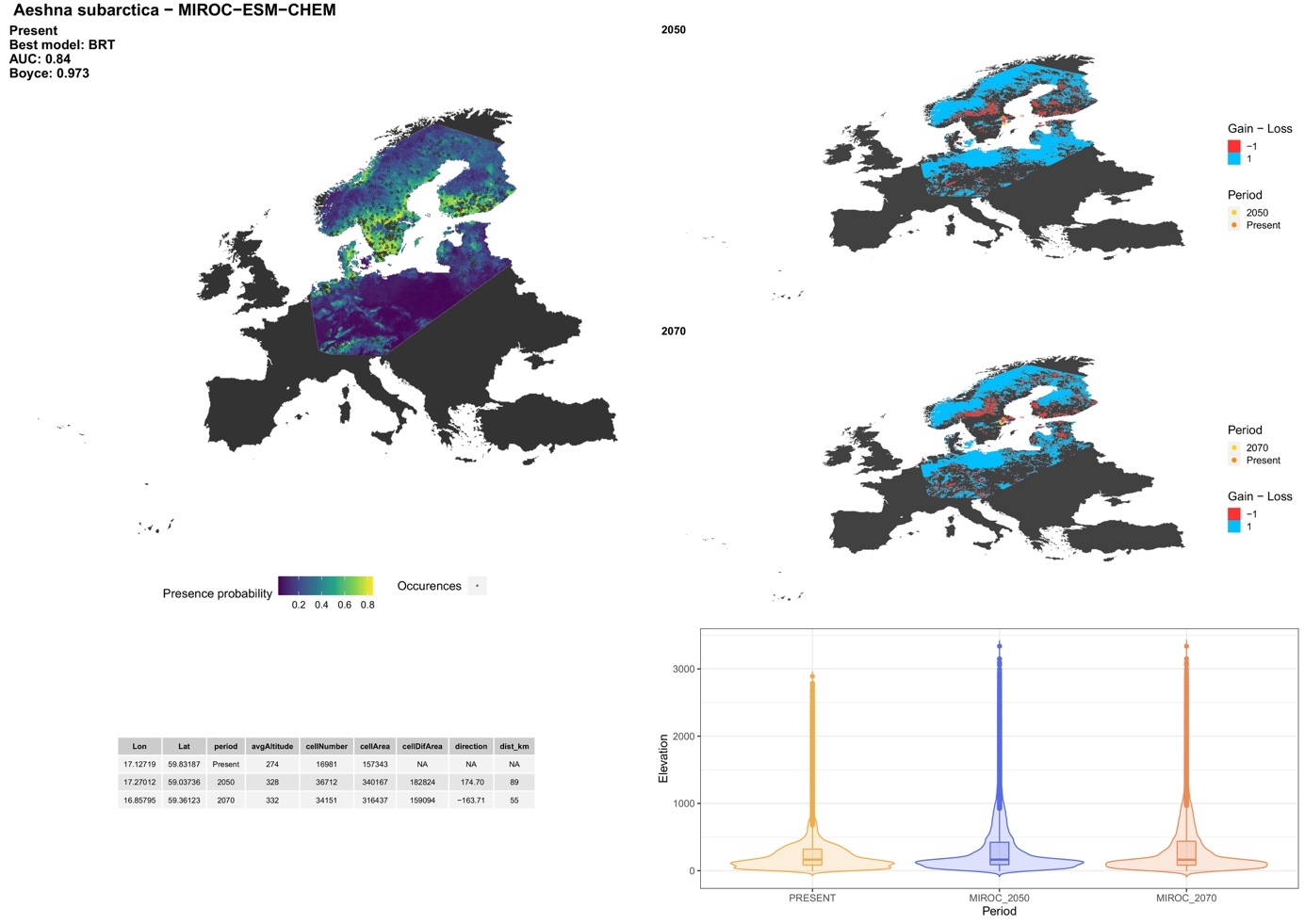

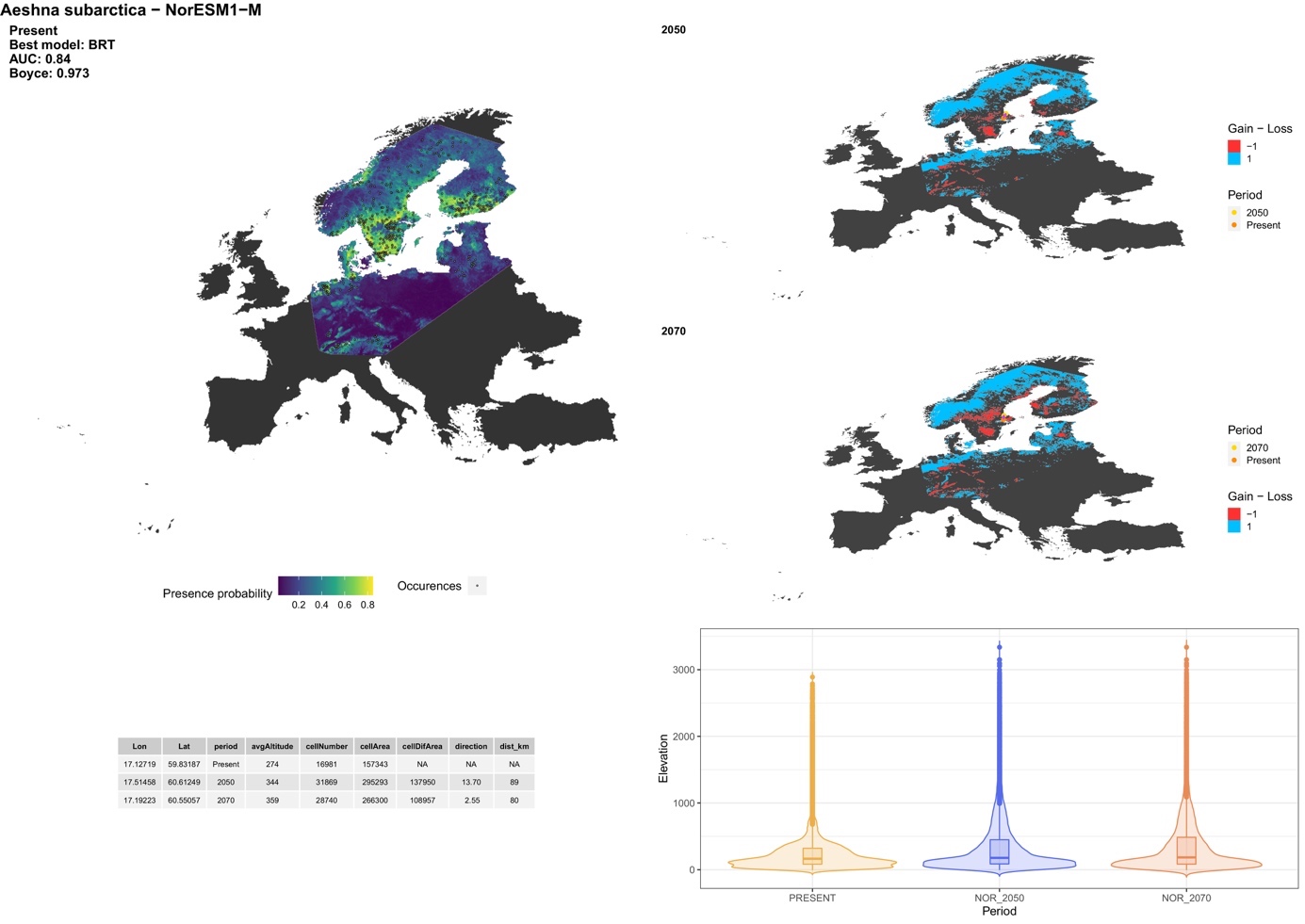

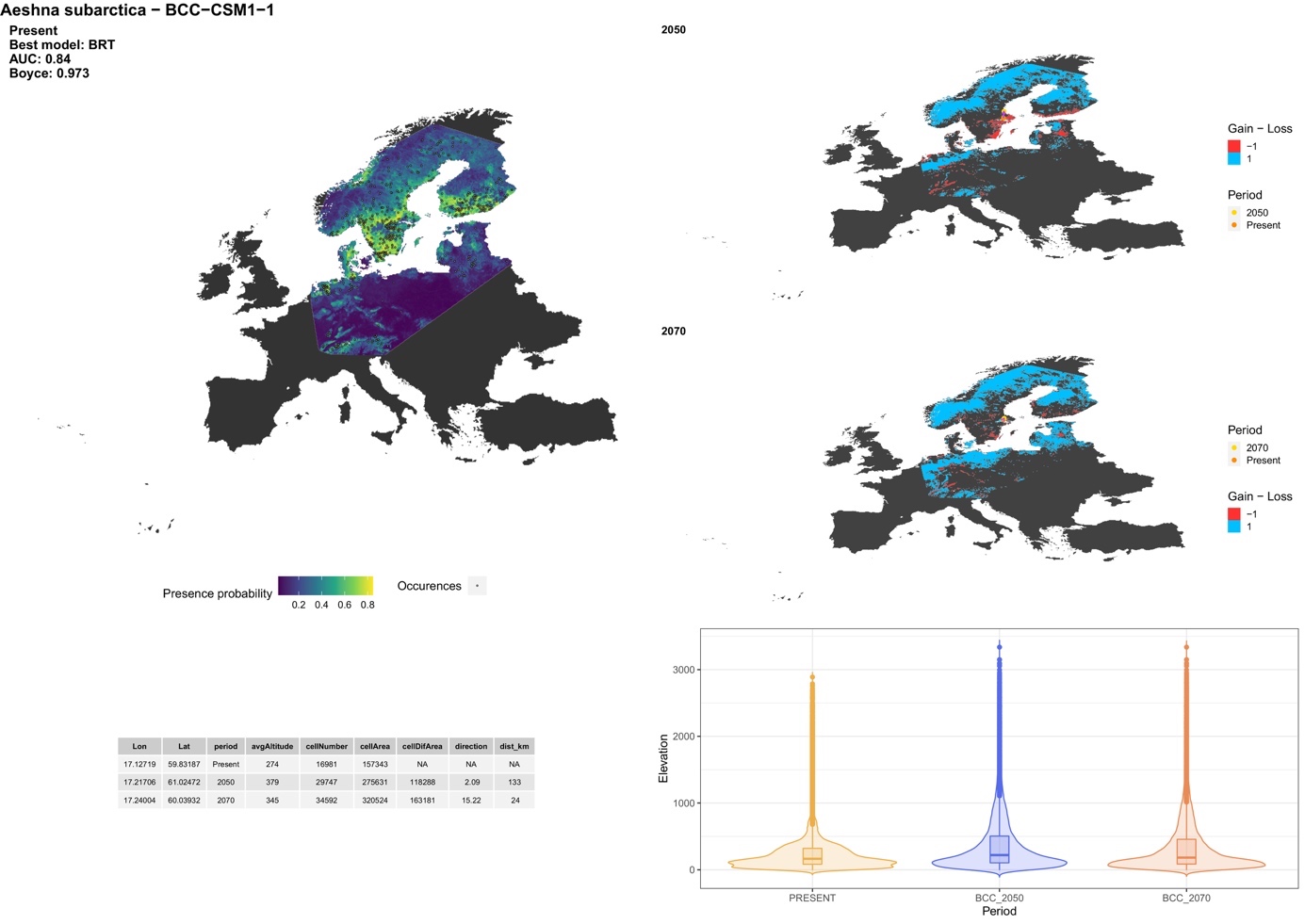

##### *Aeshna viridis* Eversmann, 1836

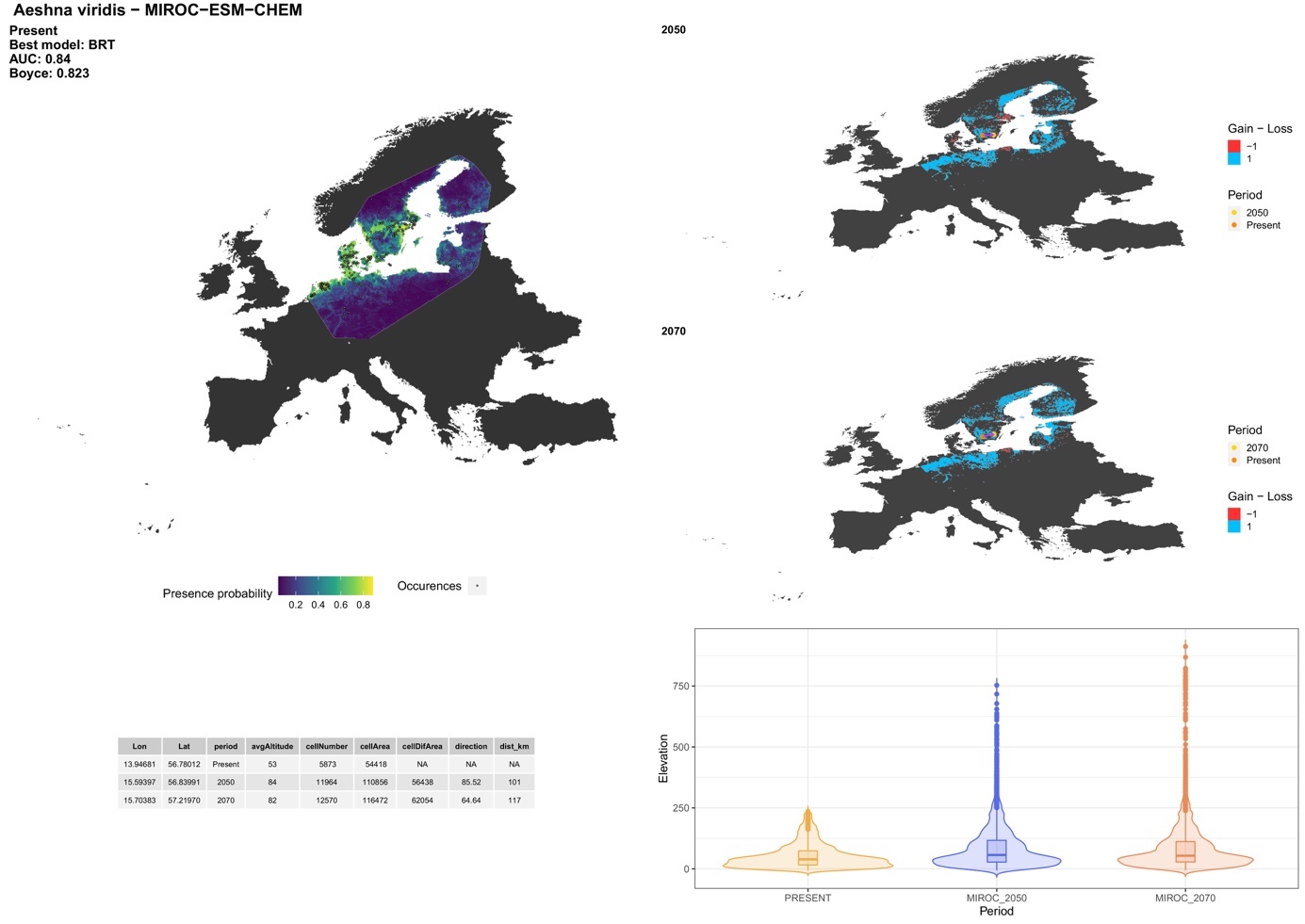

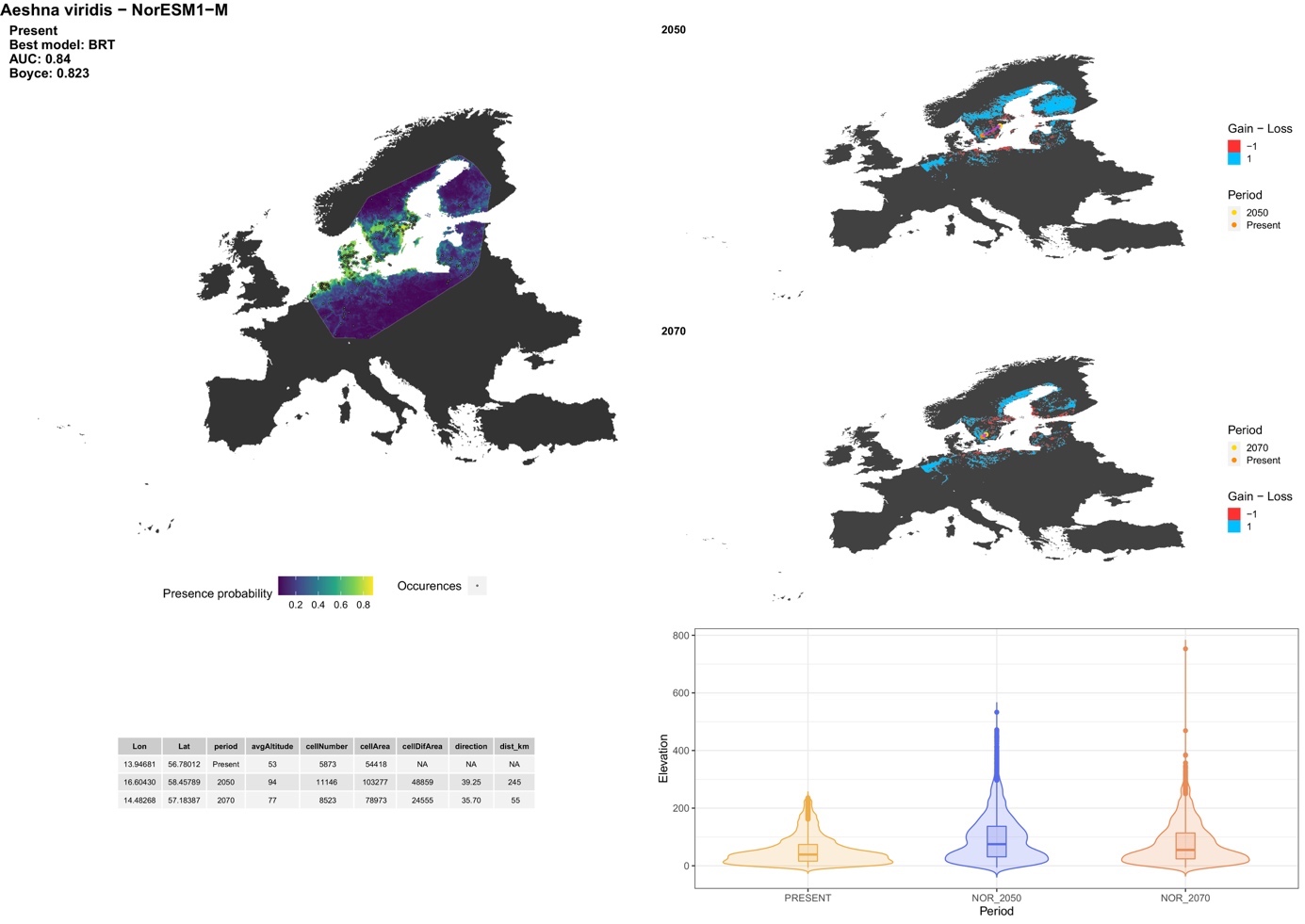

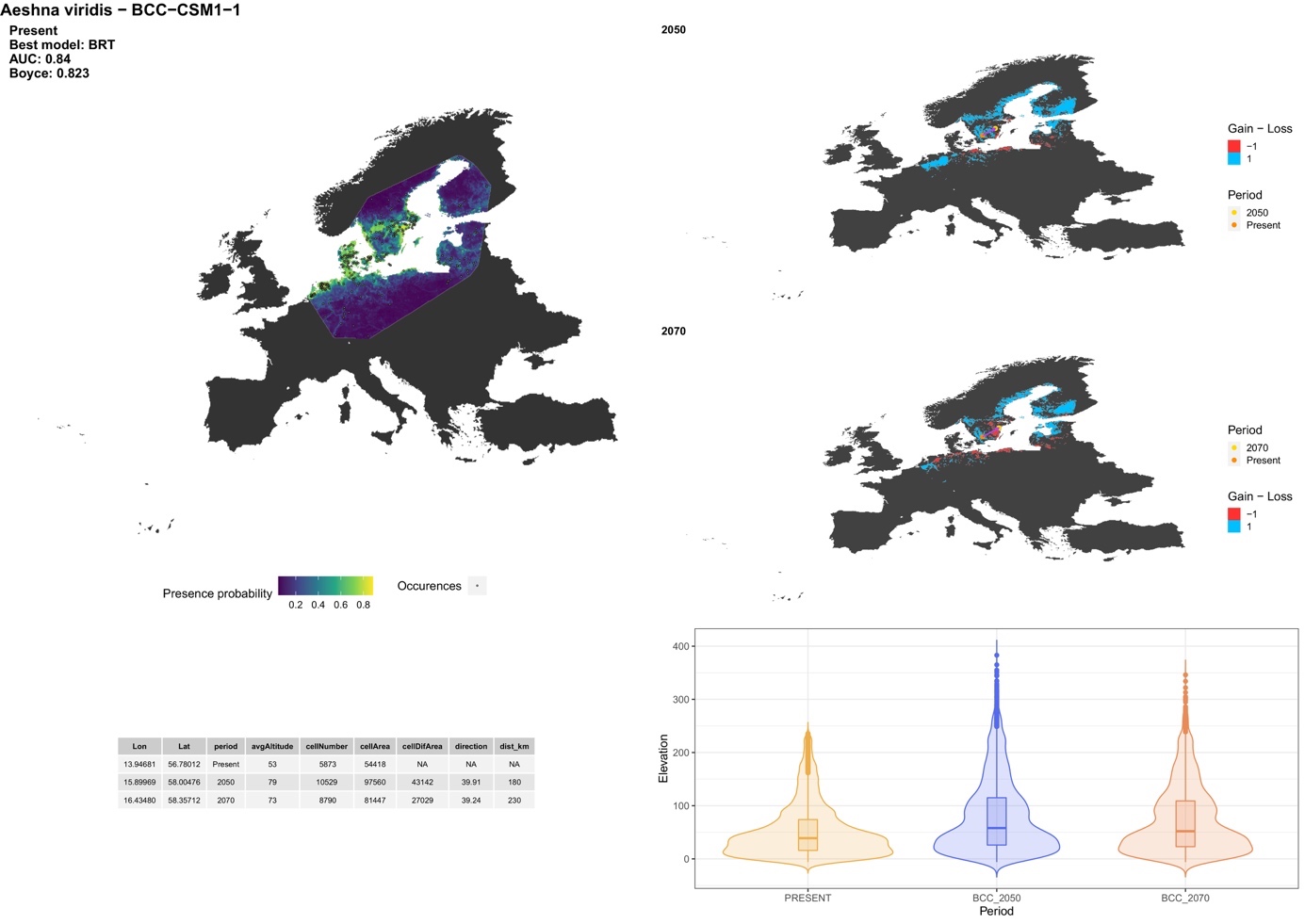

##### *Anax ephippiger* (Burmeister, 1839)

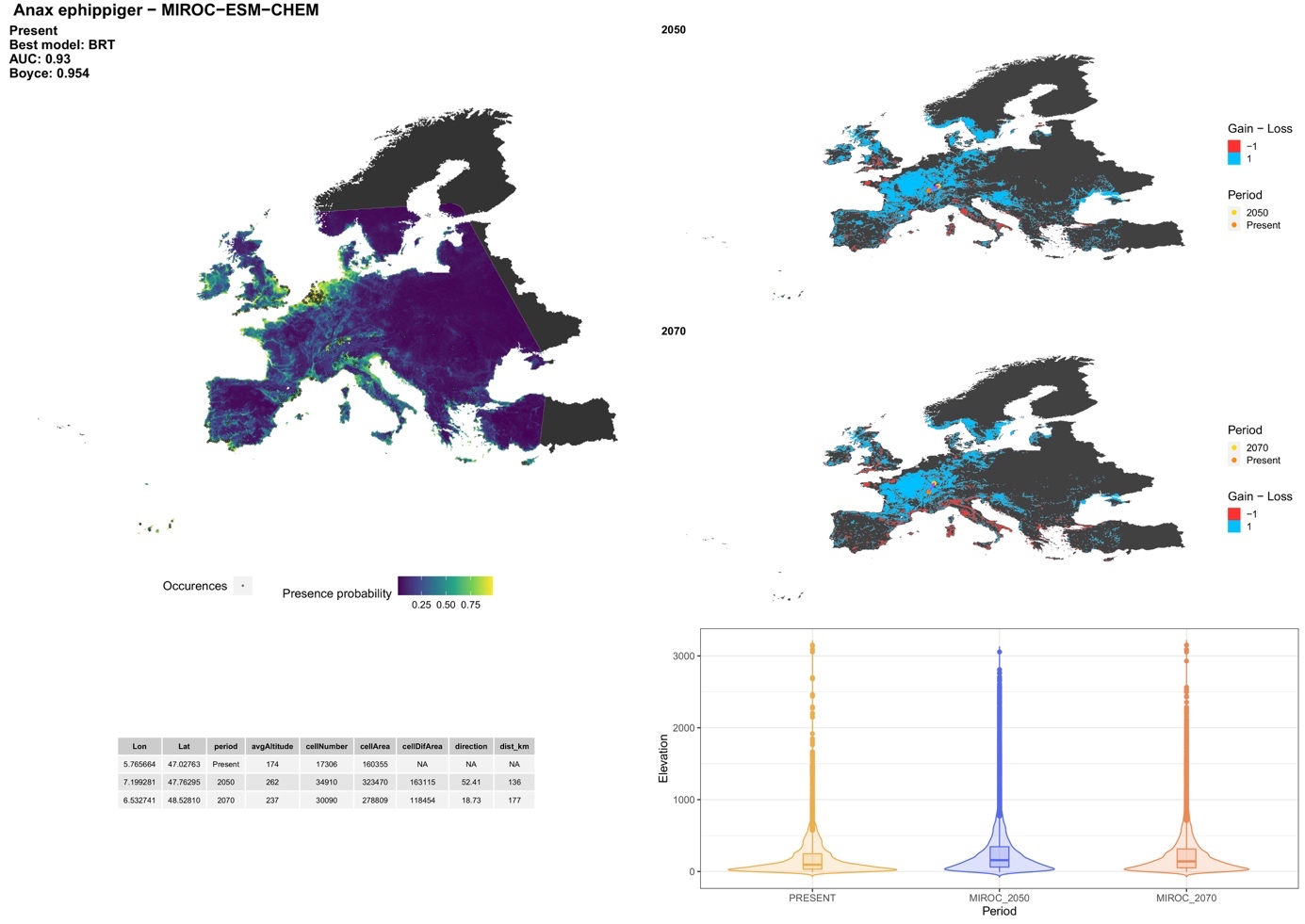

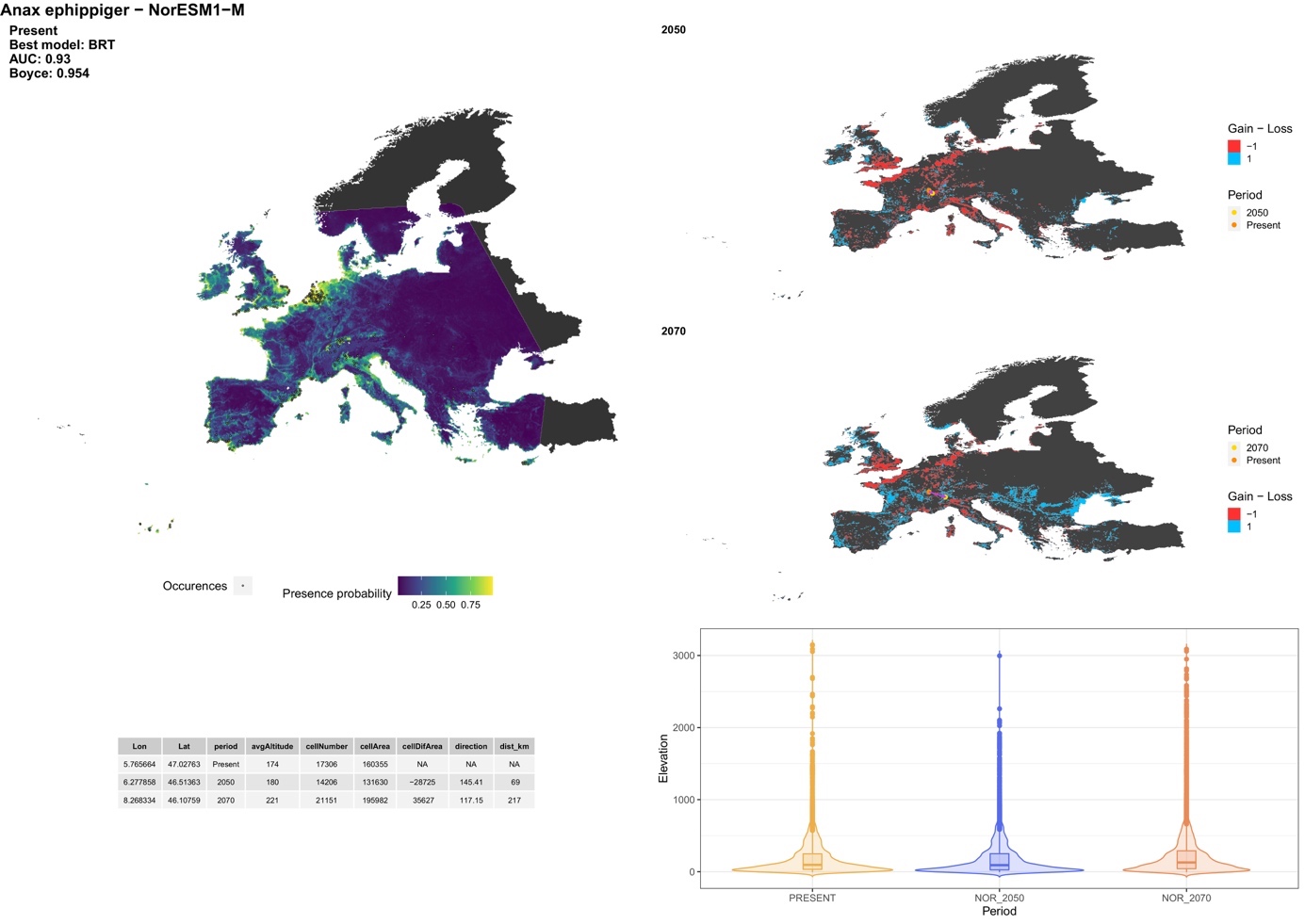

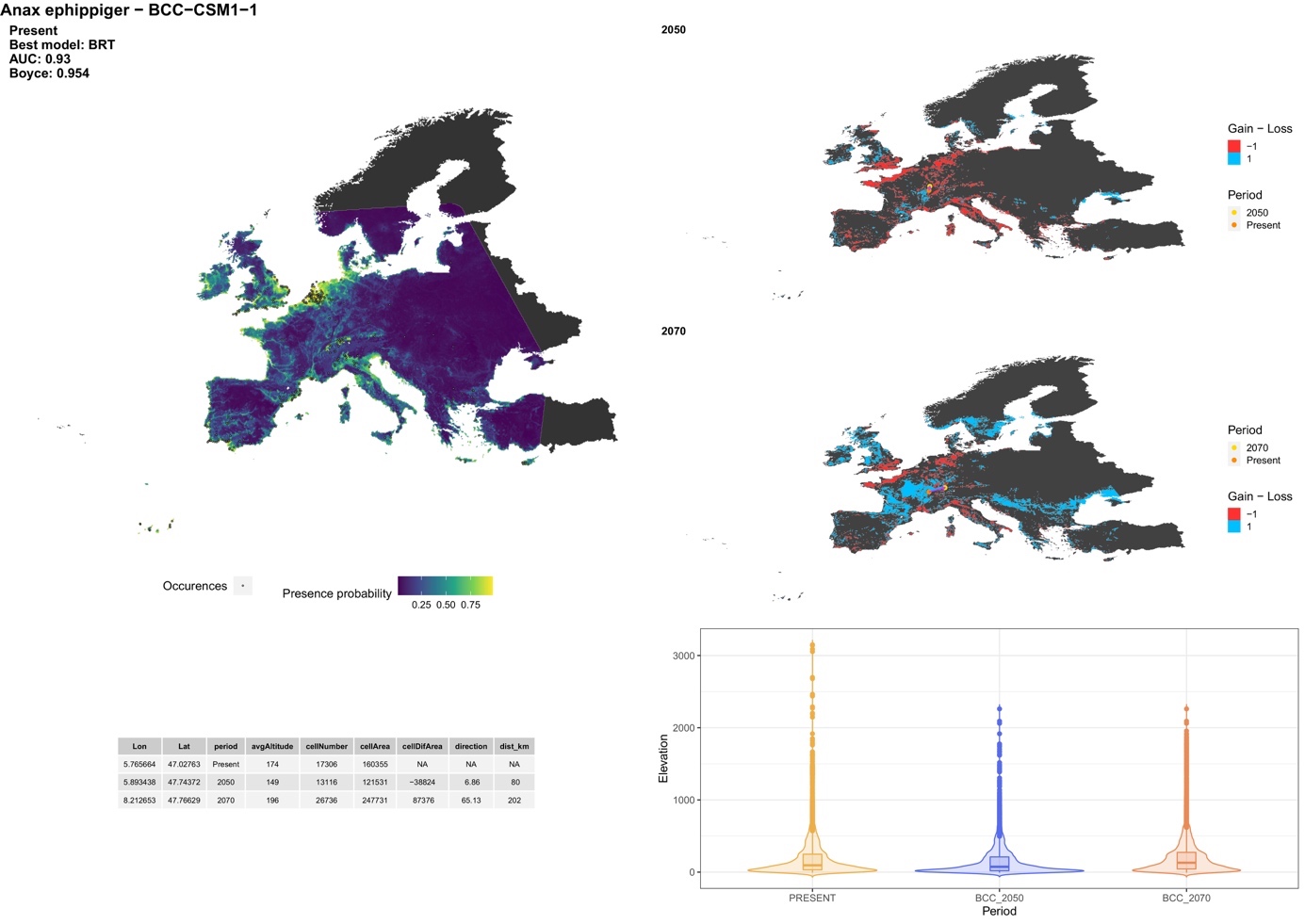

##### *Anax imperator* Leach, 1815

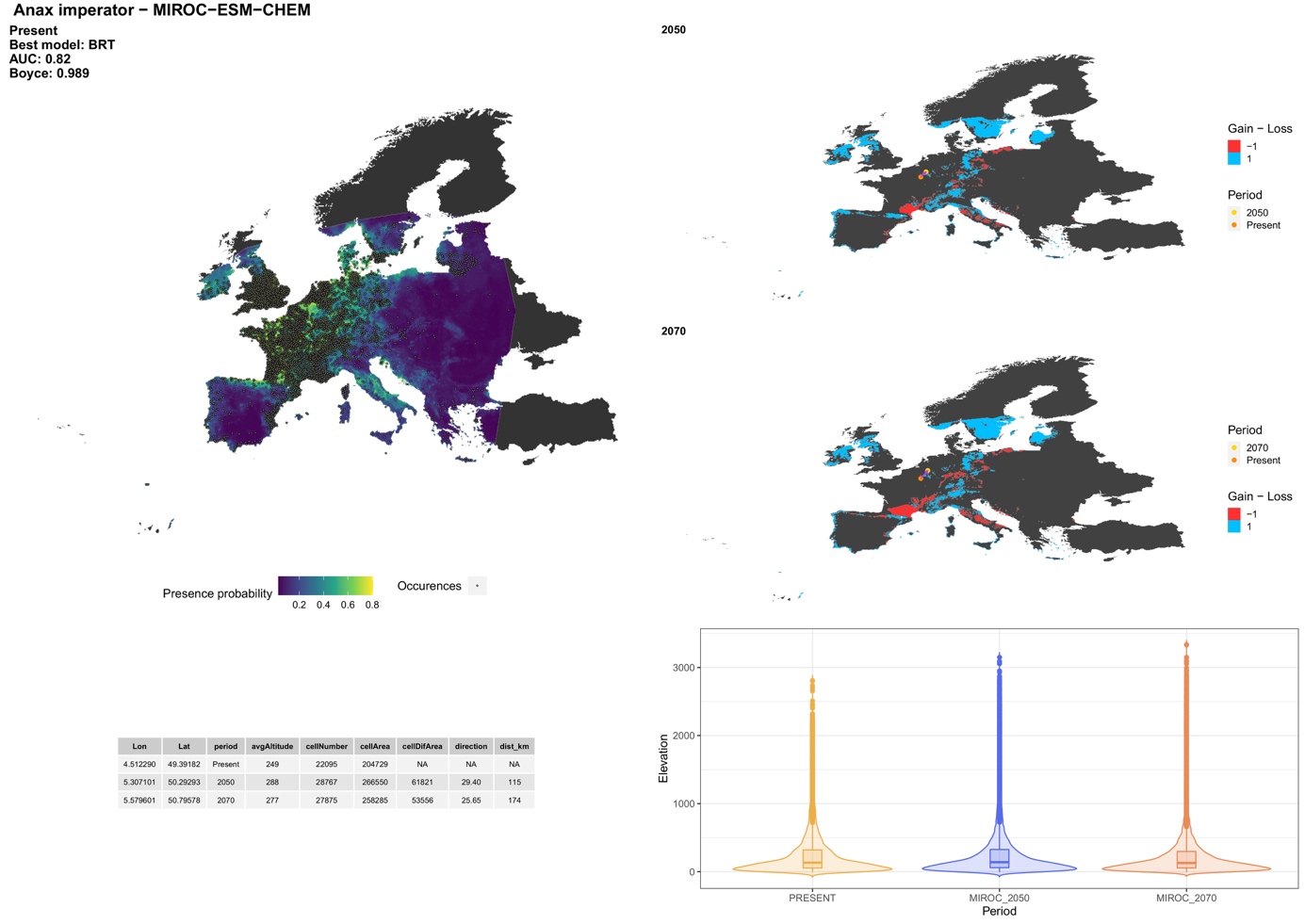

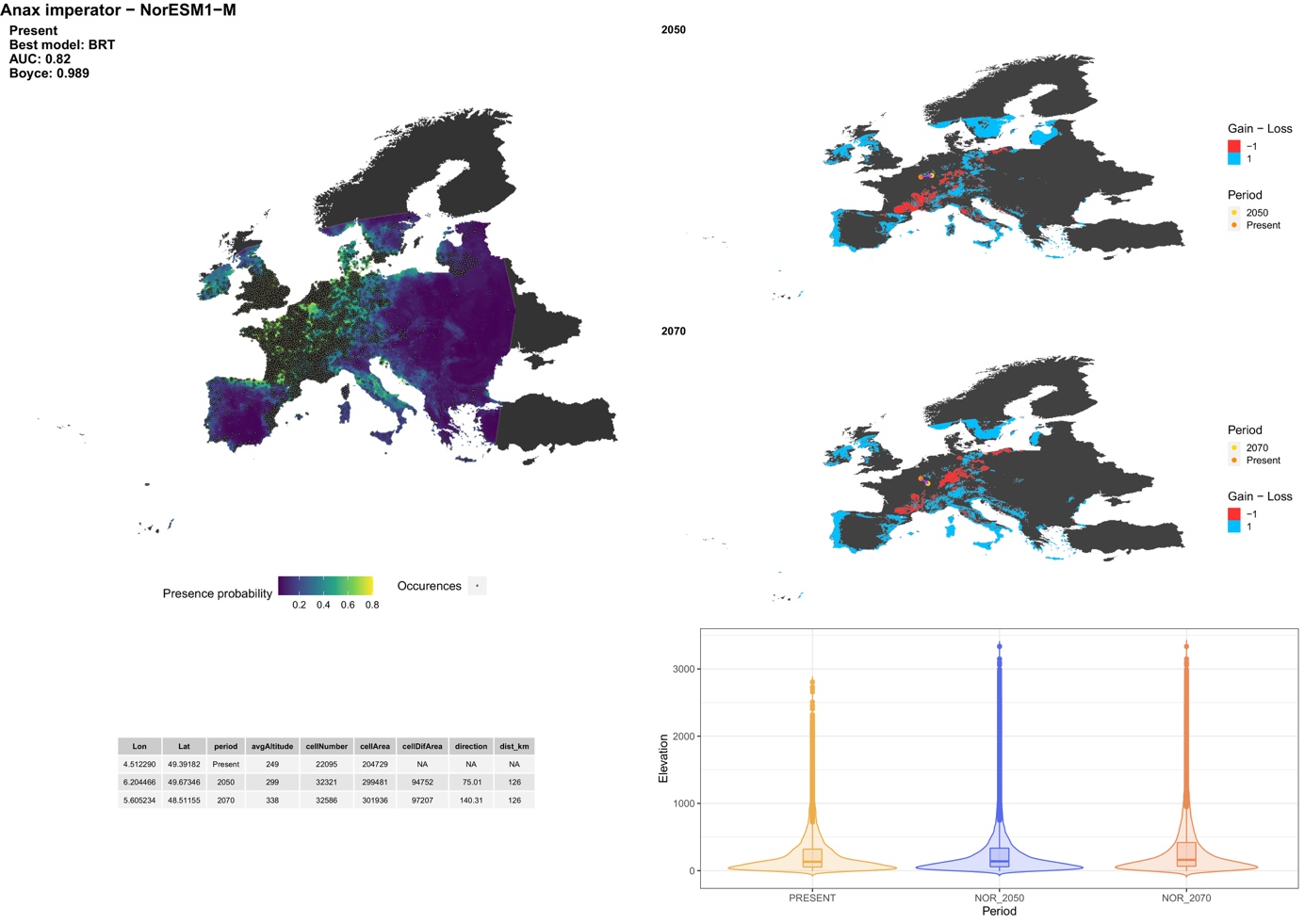

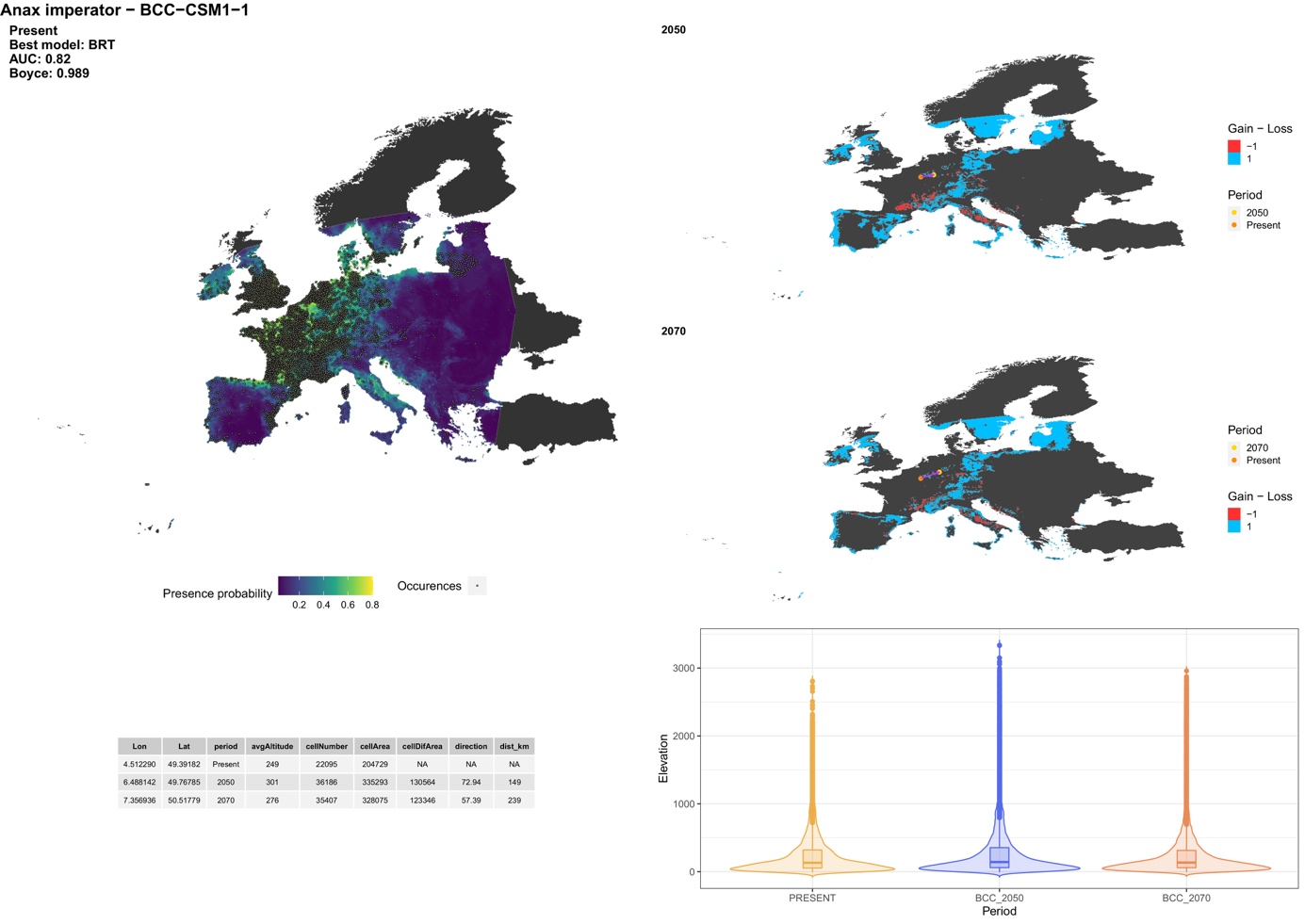

##### *Anax parthenope* (Selys, 1839)

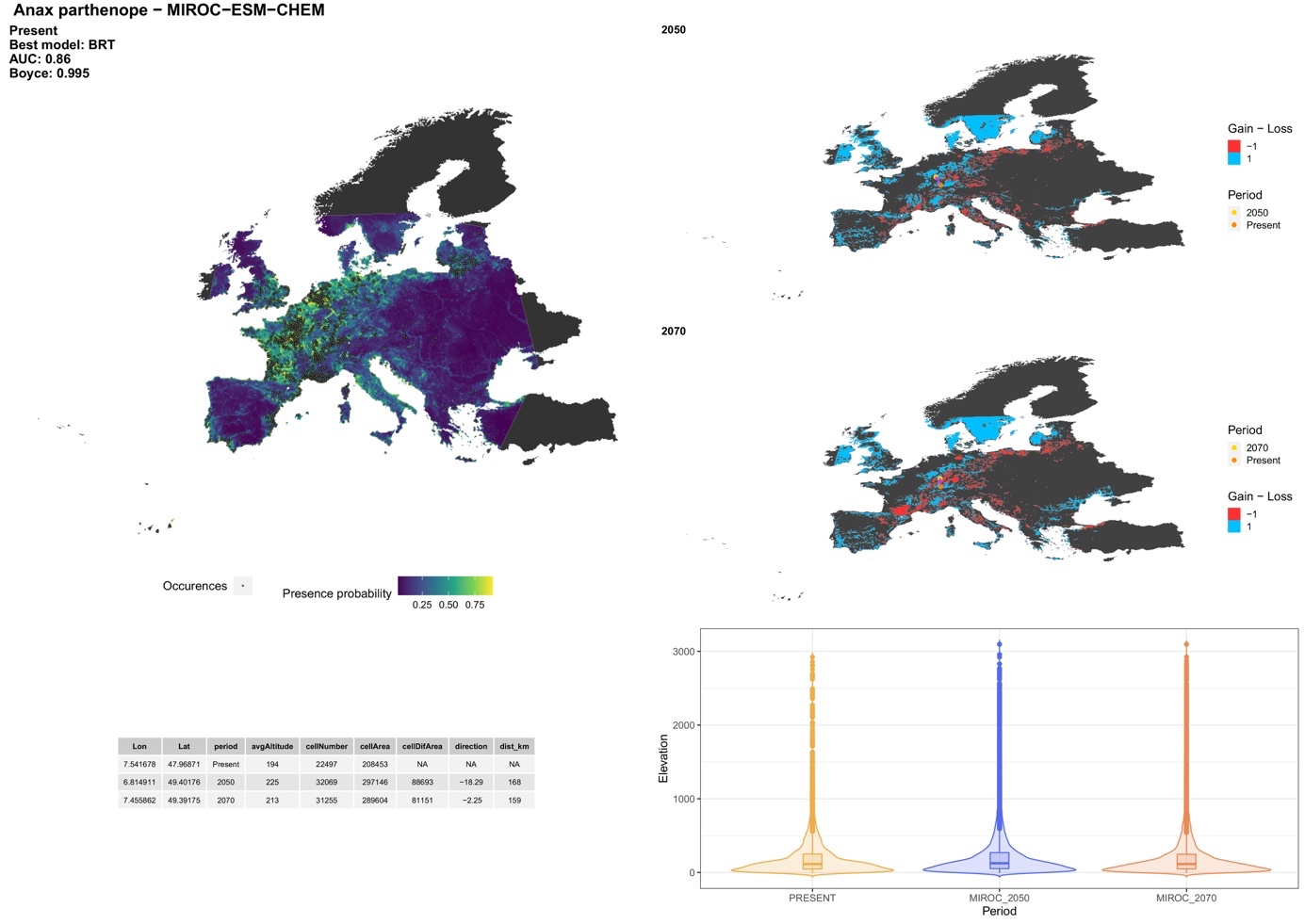

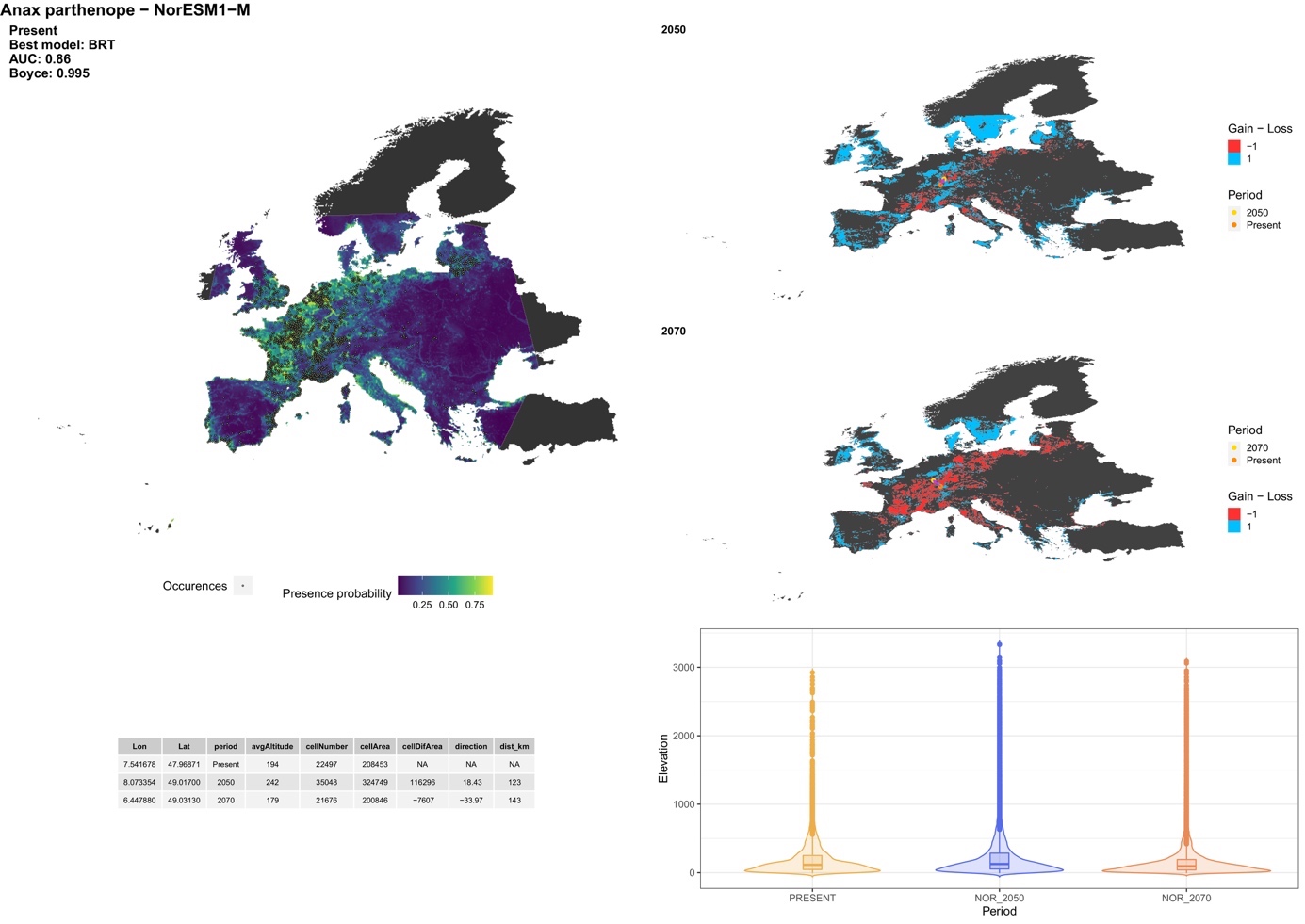

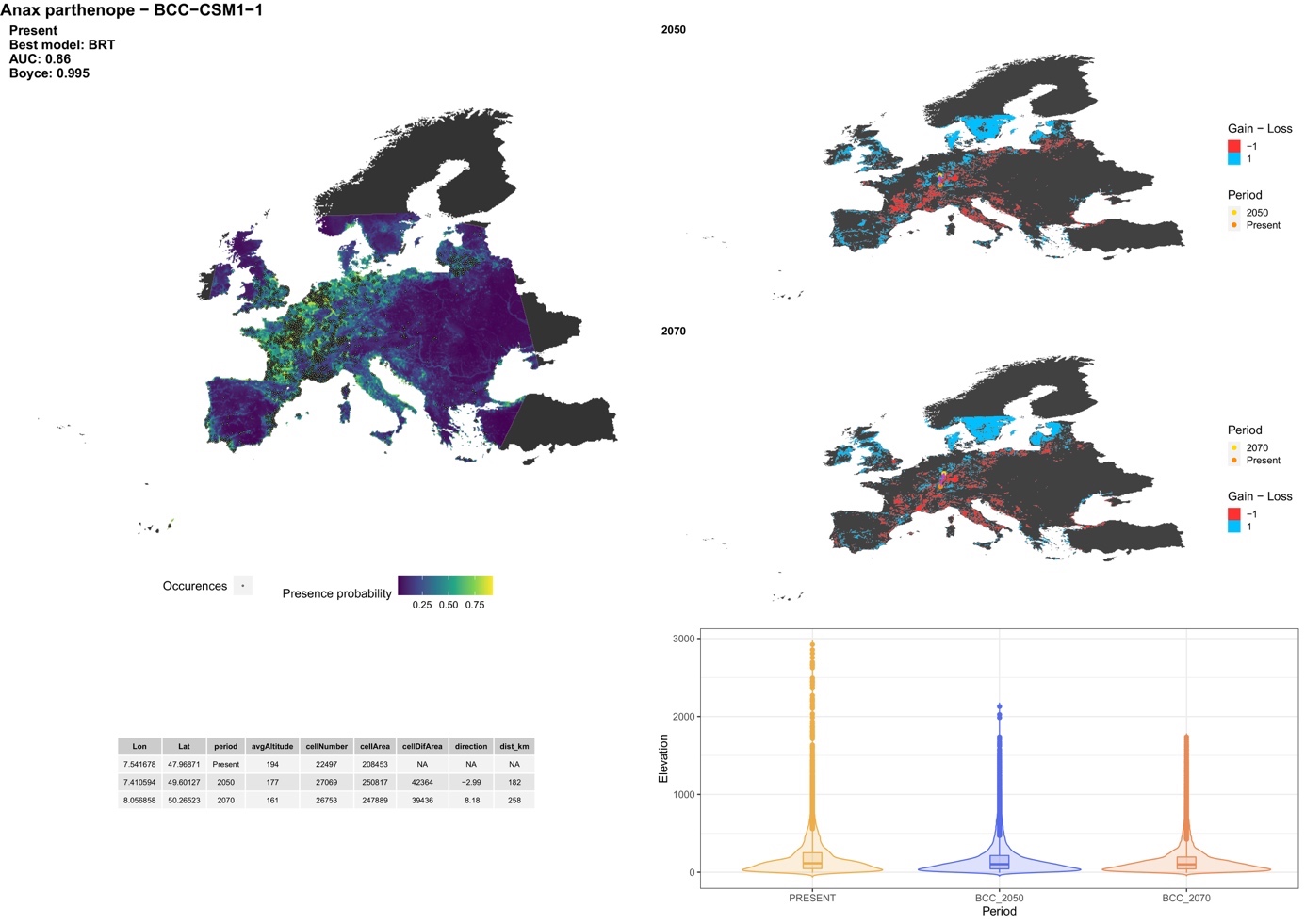

##### *Boyeria irene* (Fonscolombe, 1838)

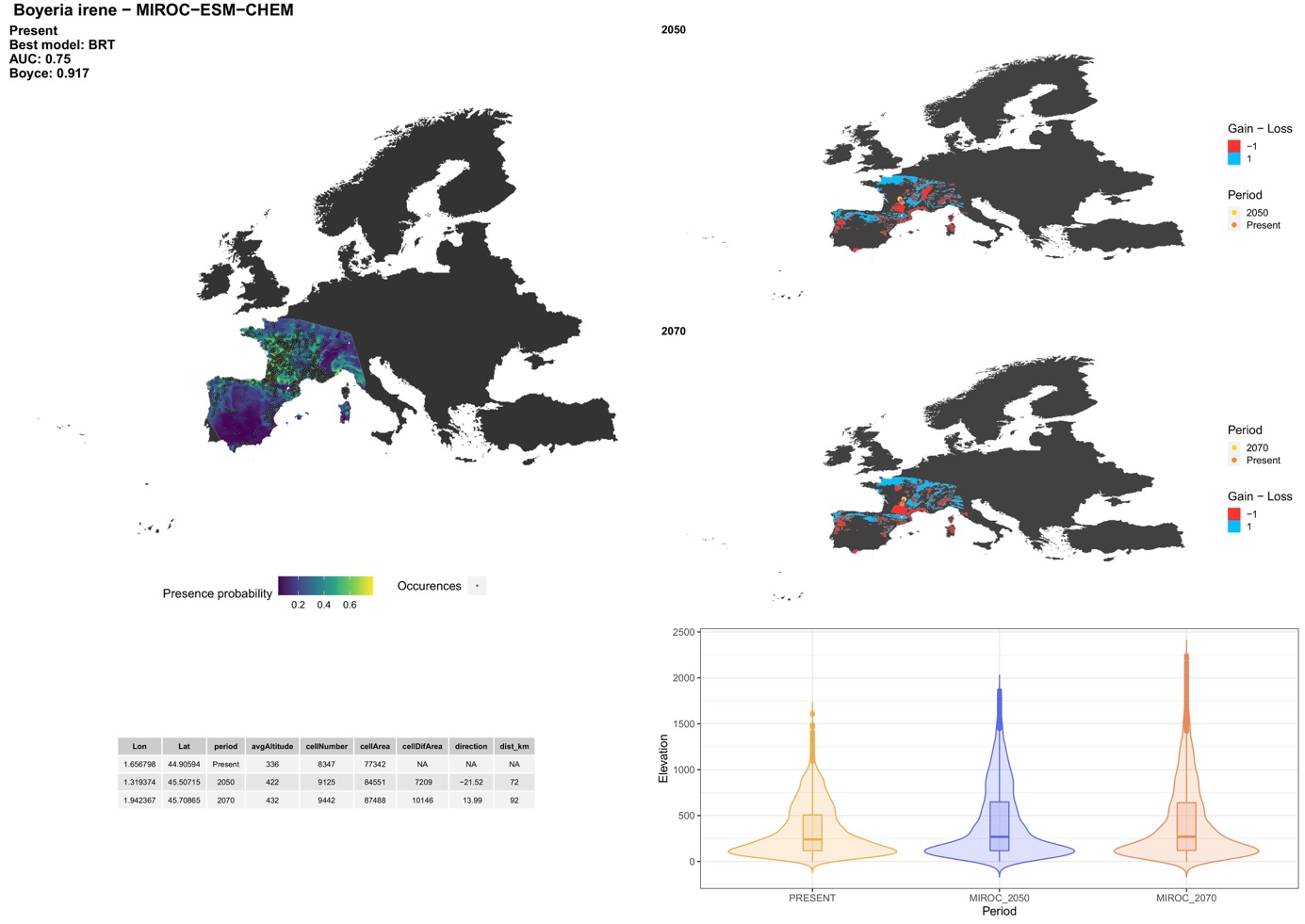

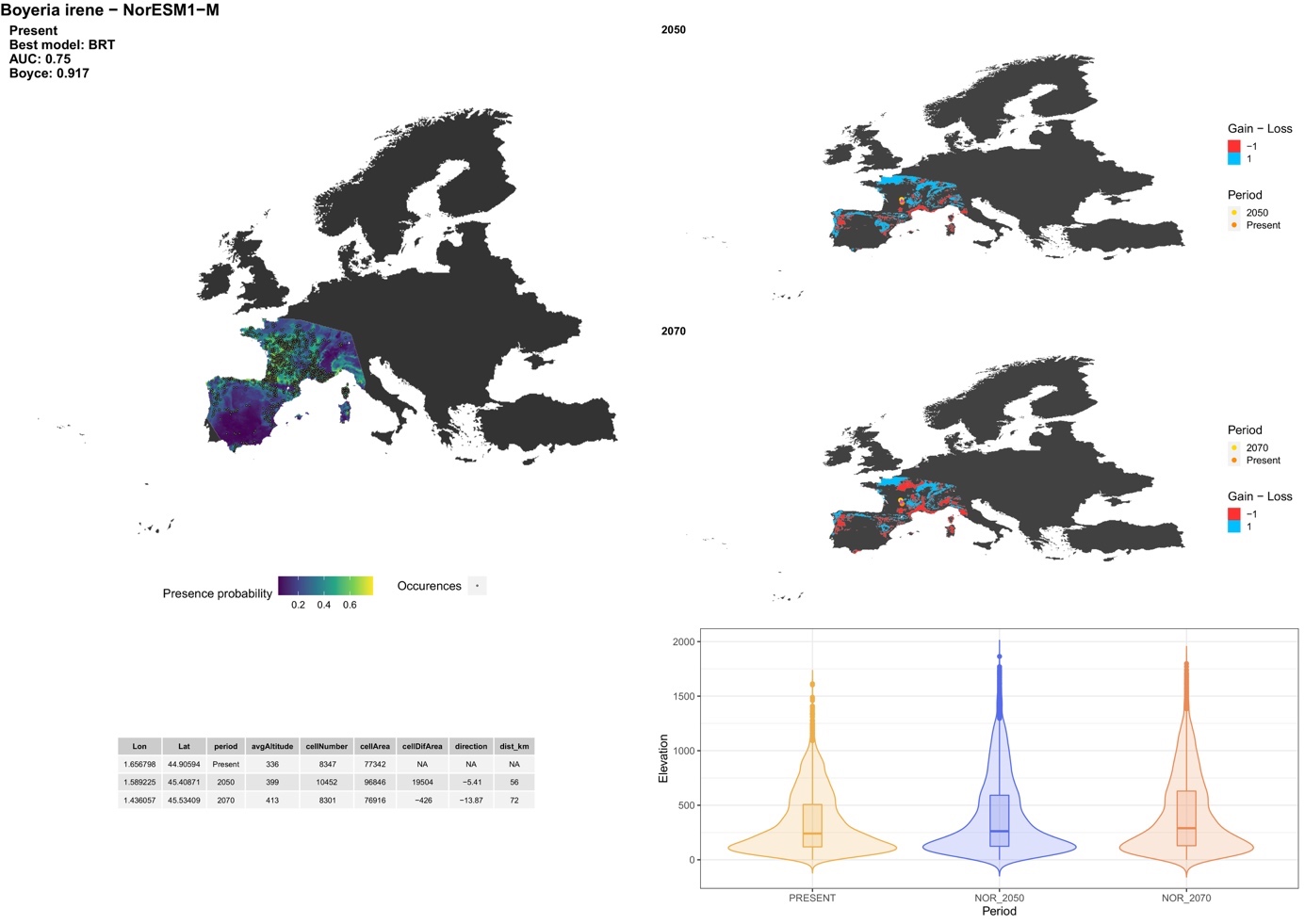

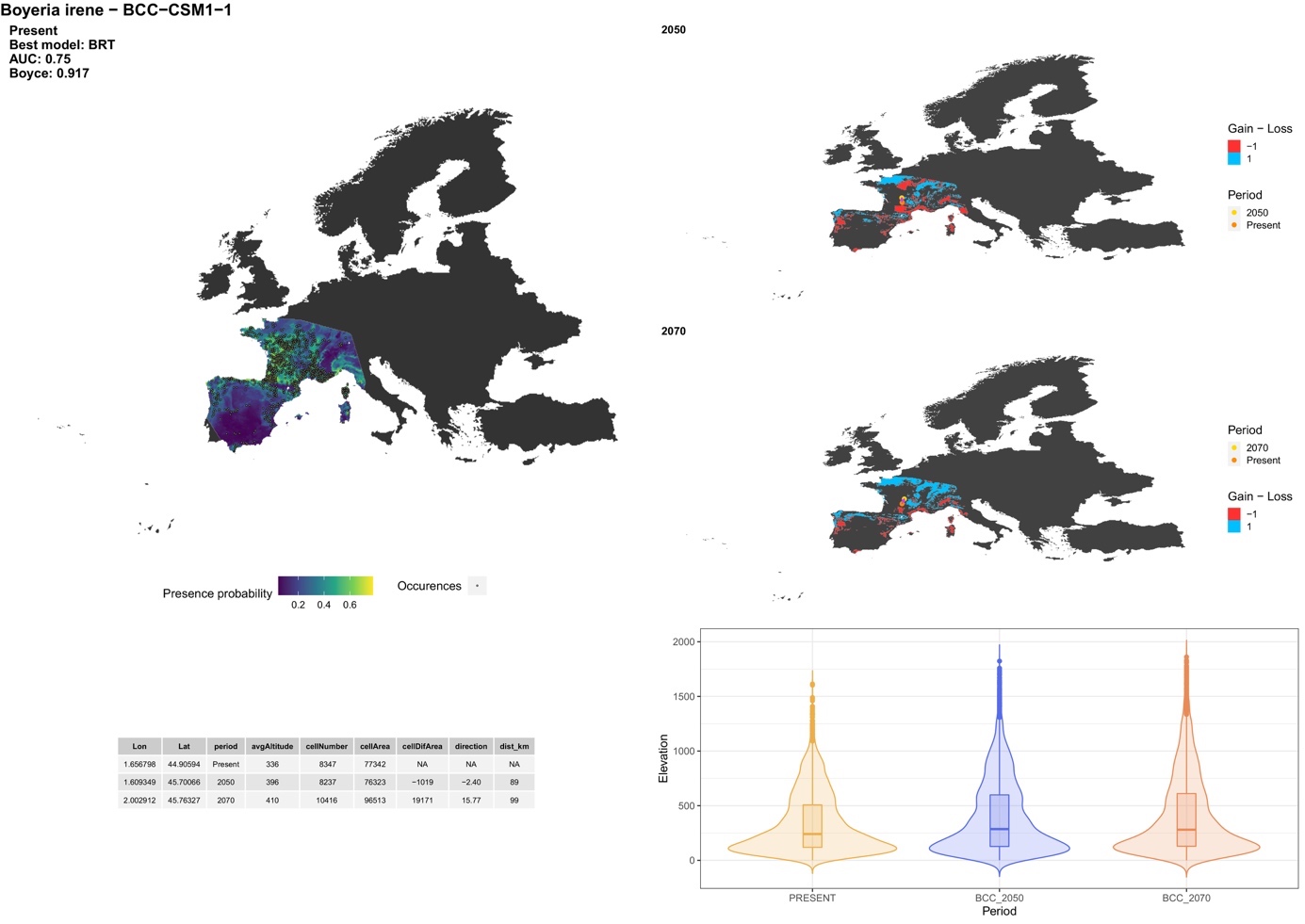

##### *Brachytron pratense* (Müller, 1764)

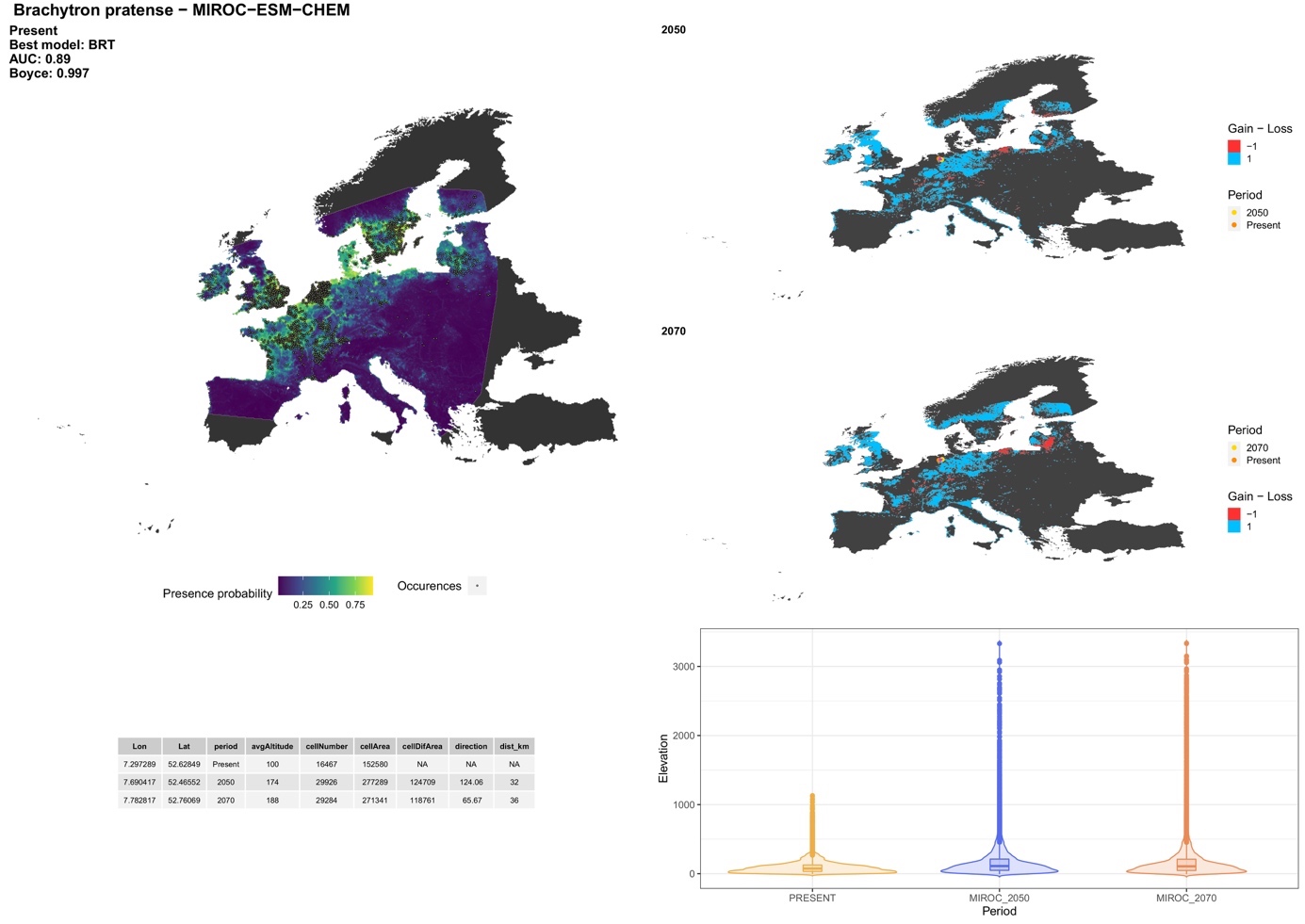

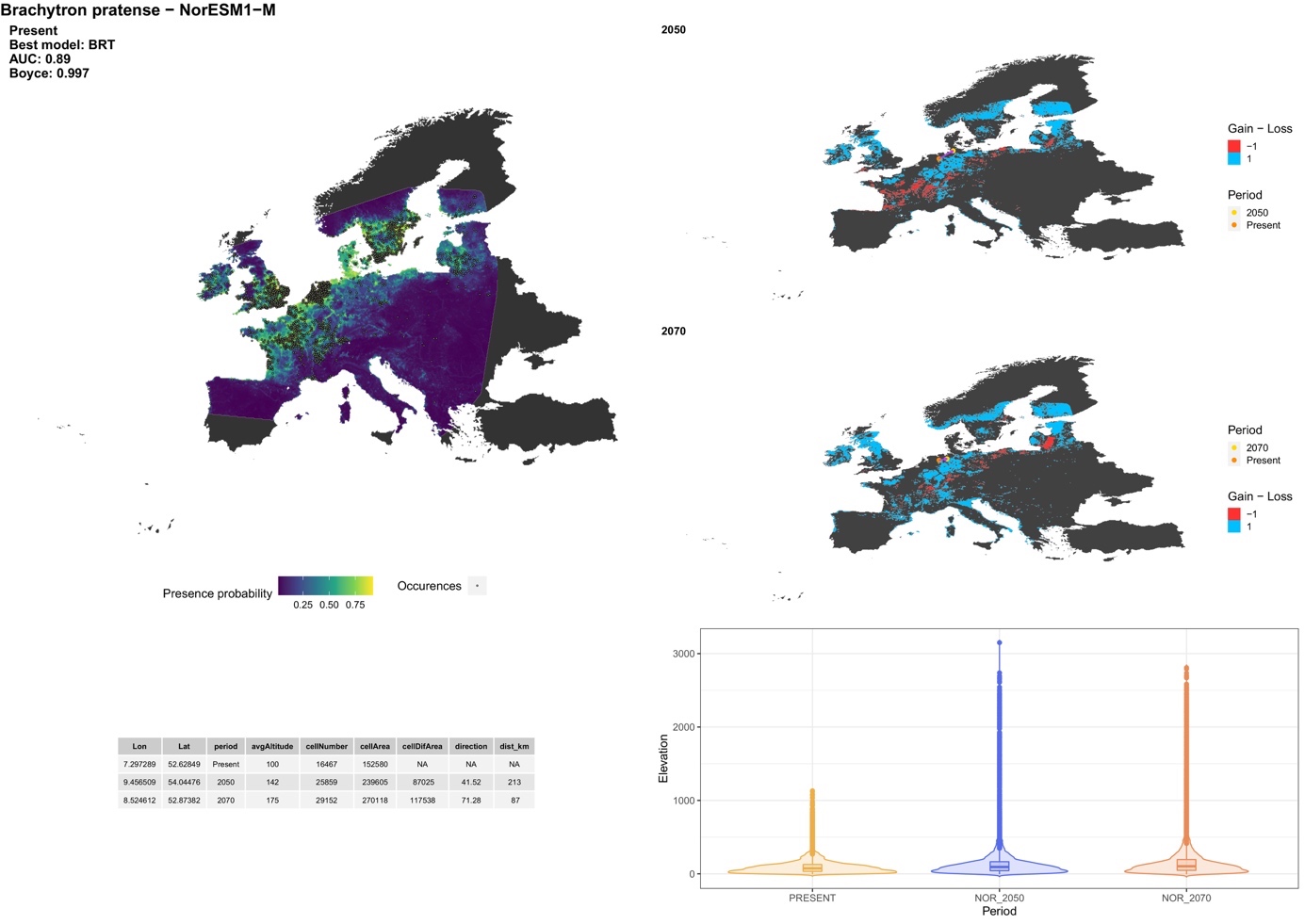

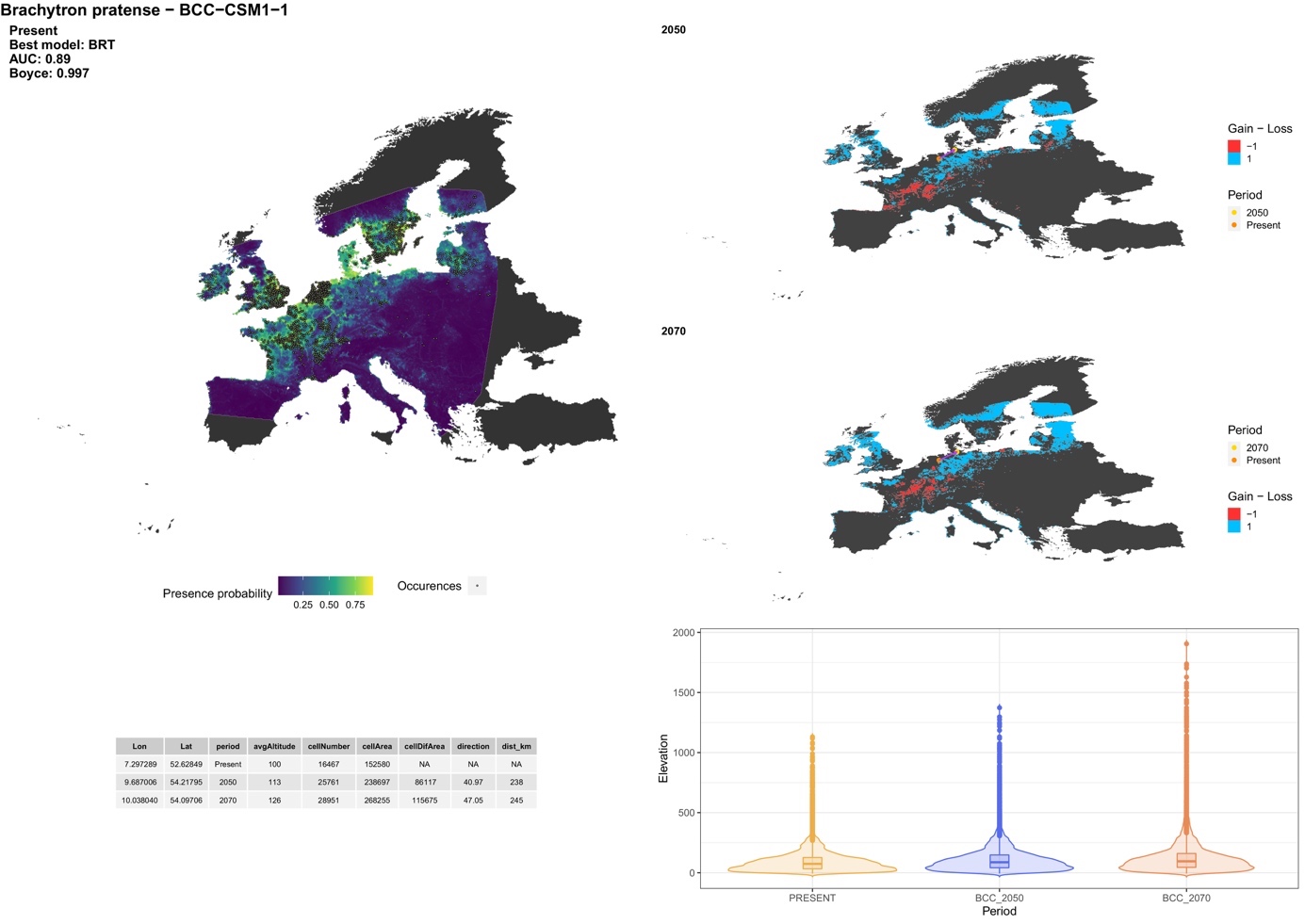

##### *Caliaeschna microstigma* (Schneider, 1845)

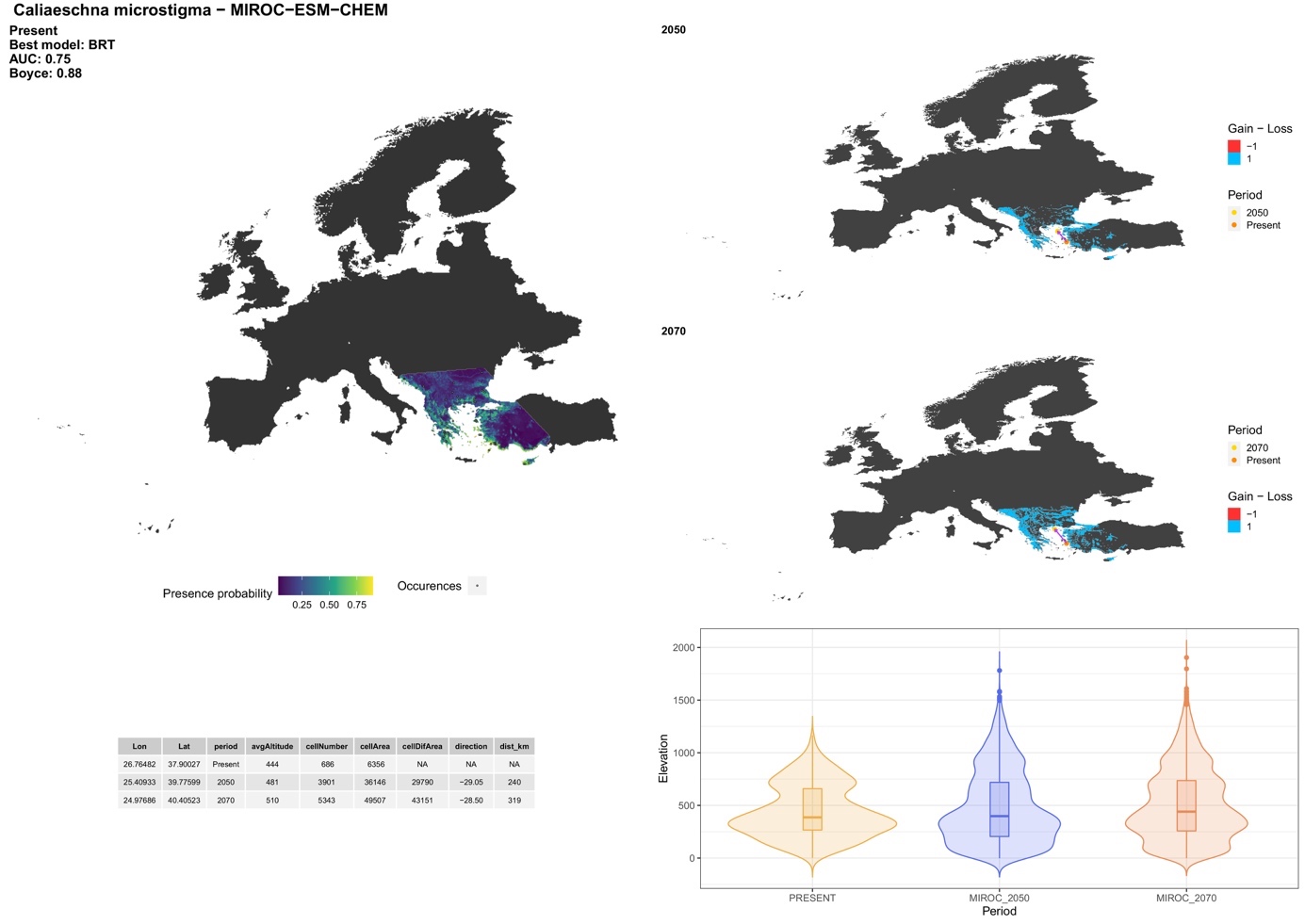

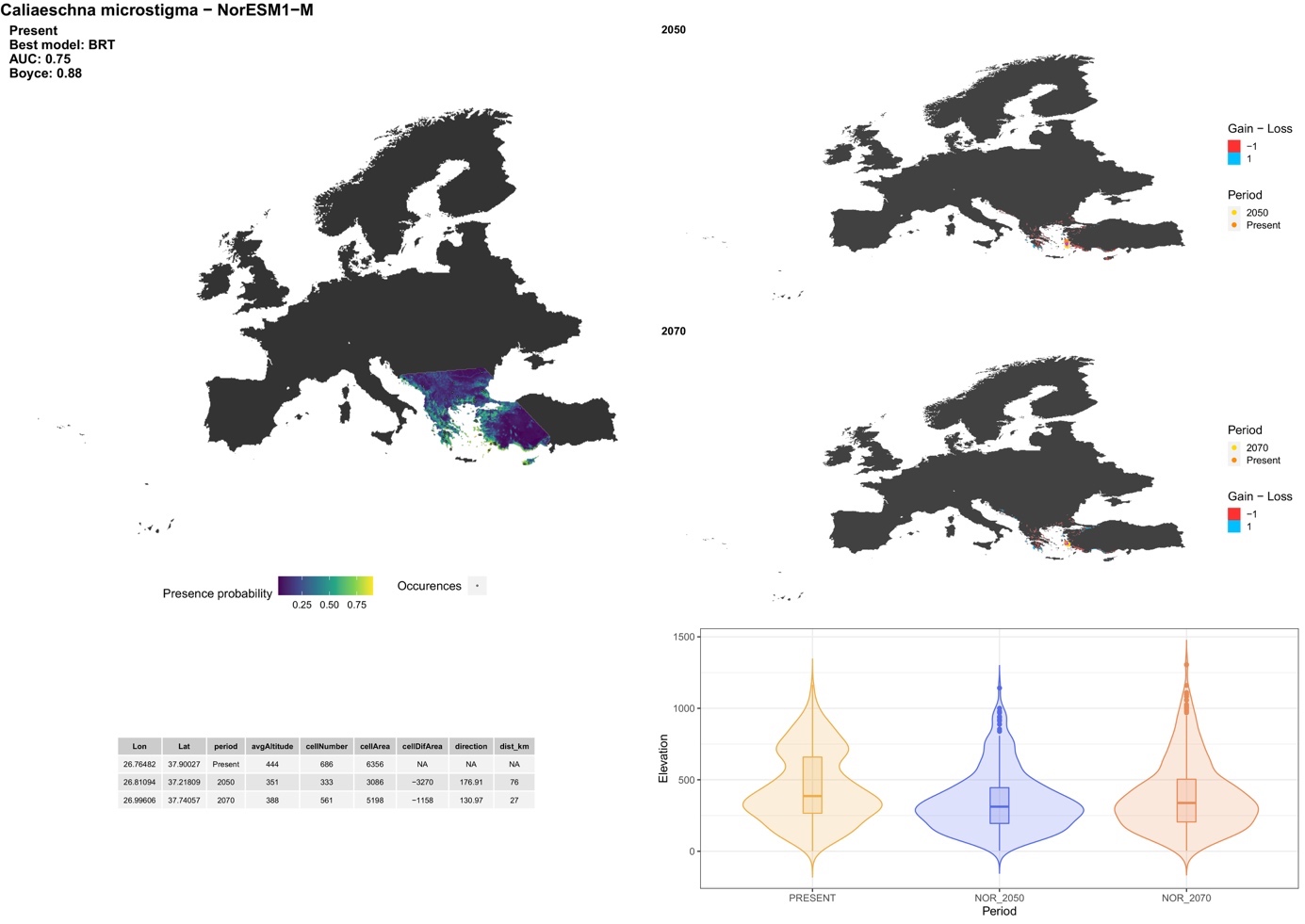

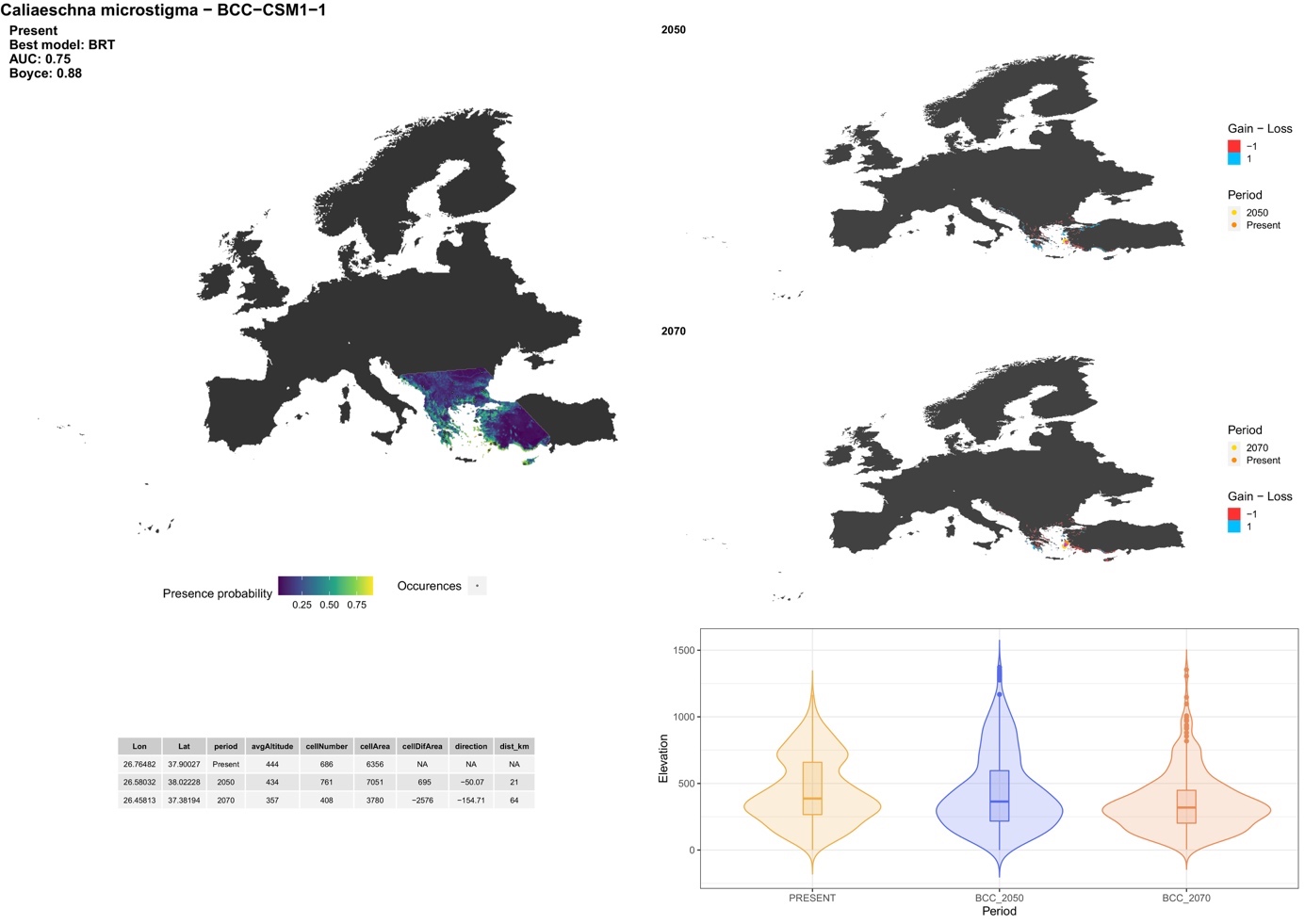

##### **Family: Gomphidae**

##### *Gomphus graslinii* Rambur, 1842

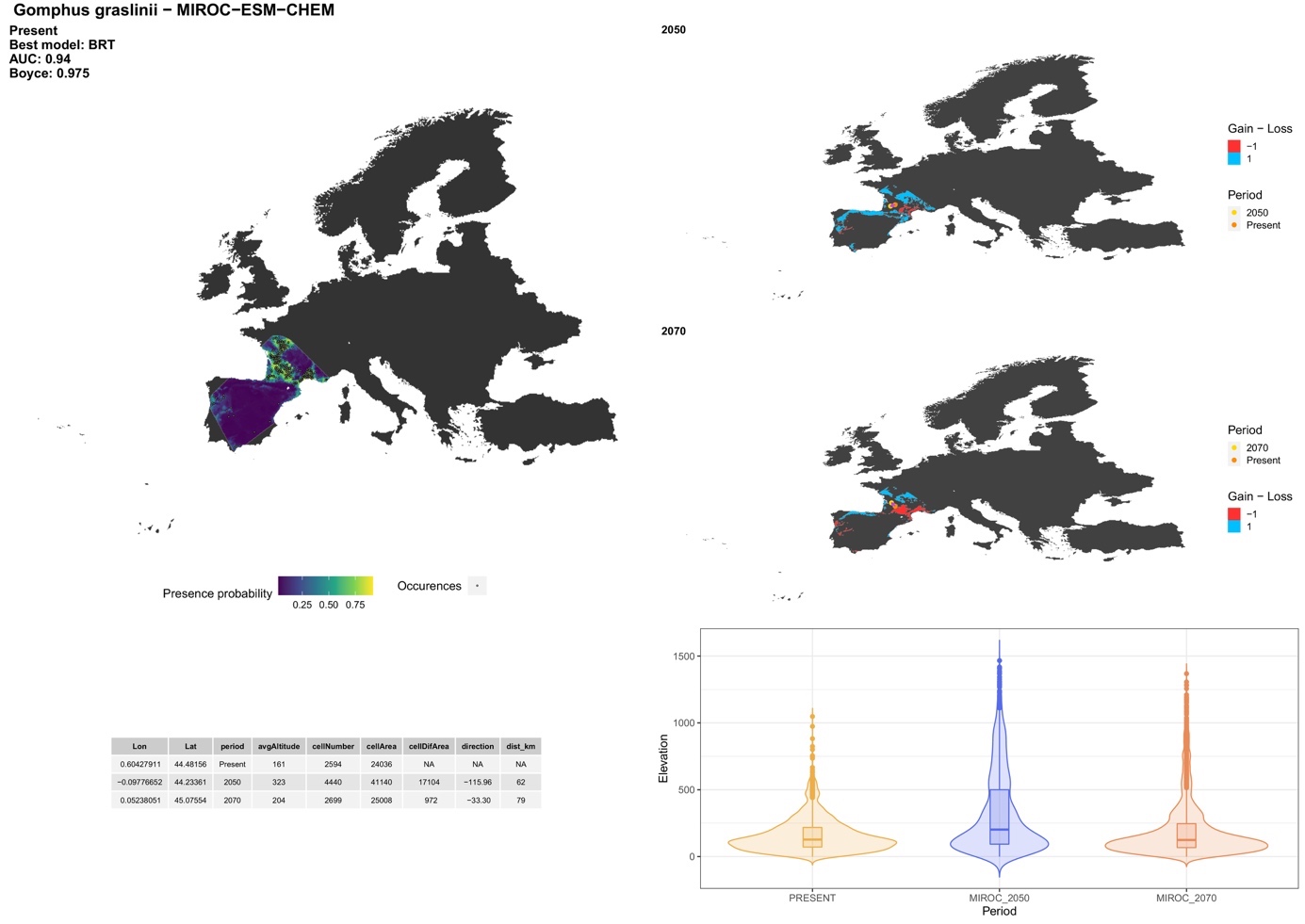

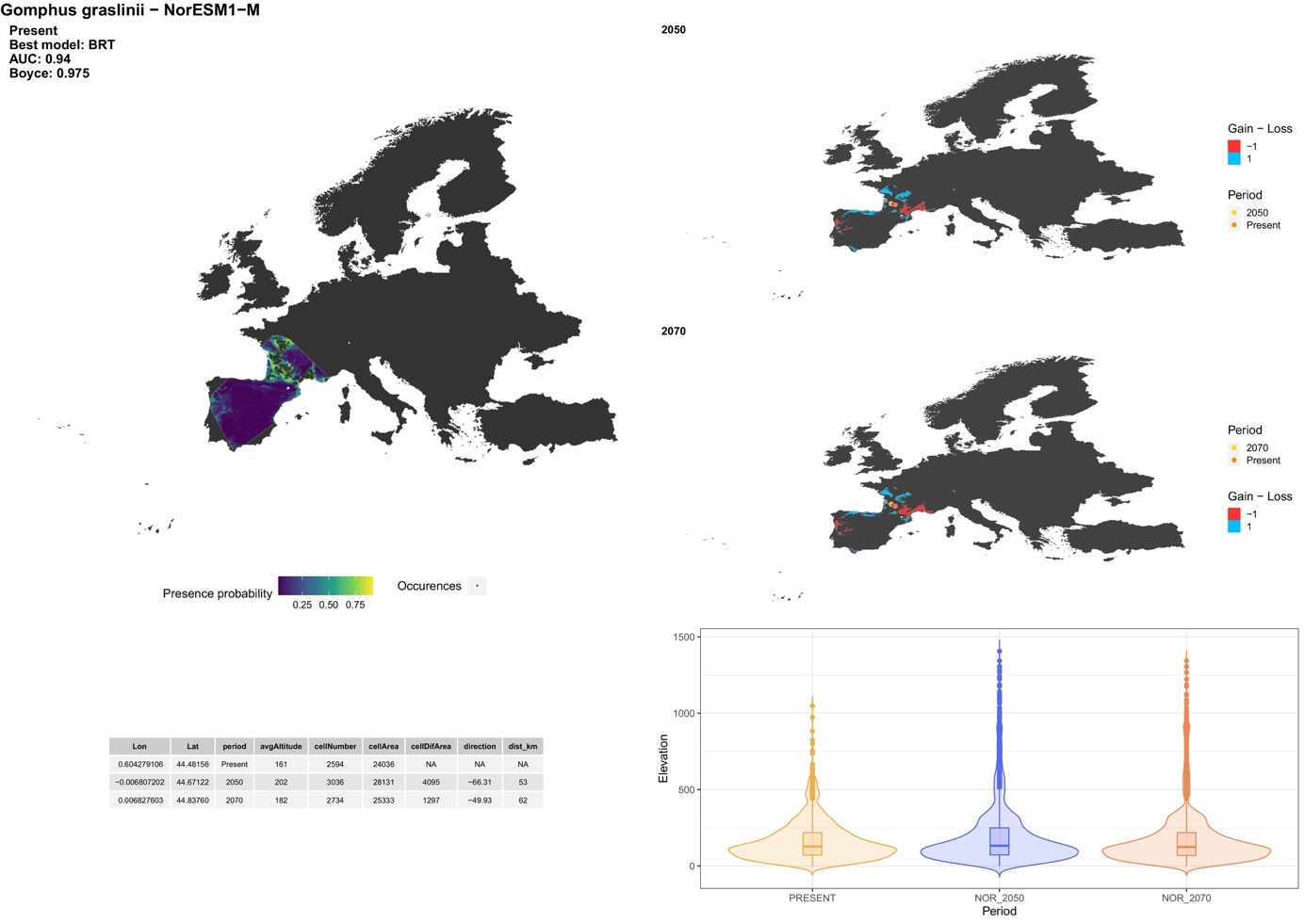

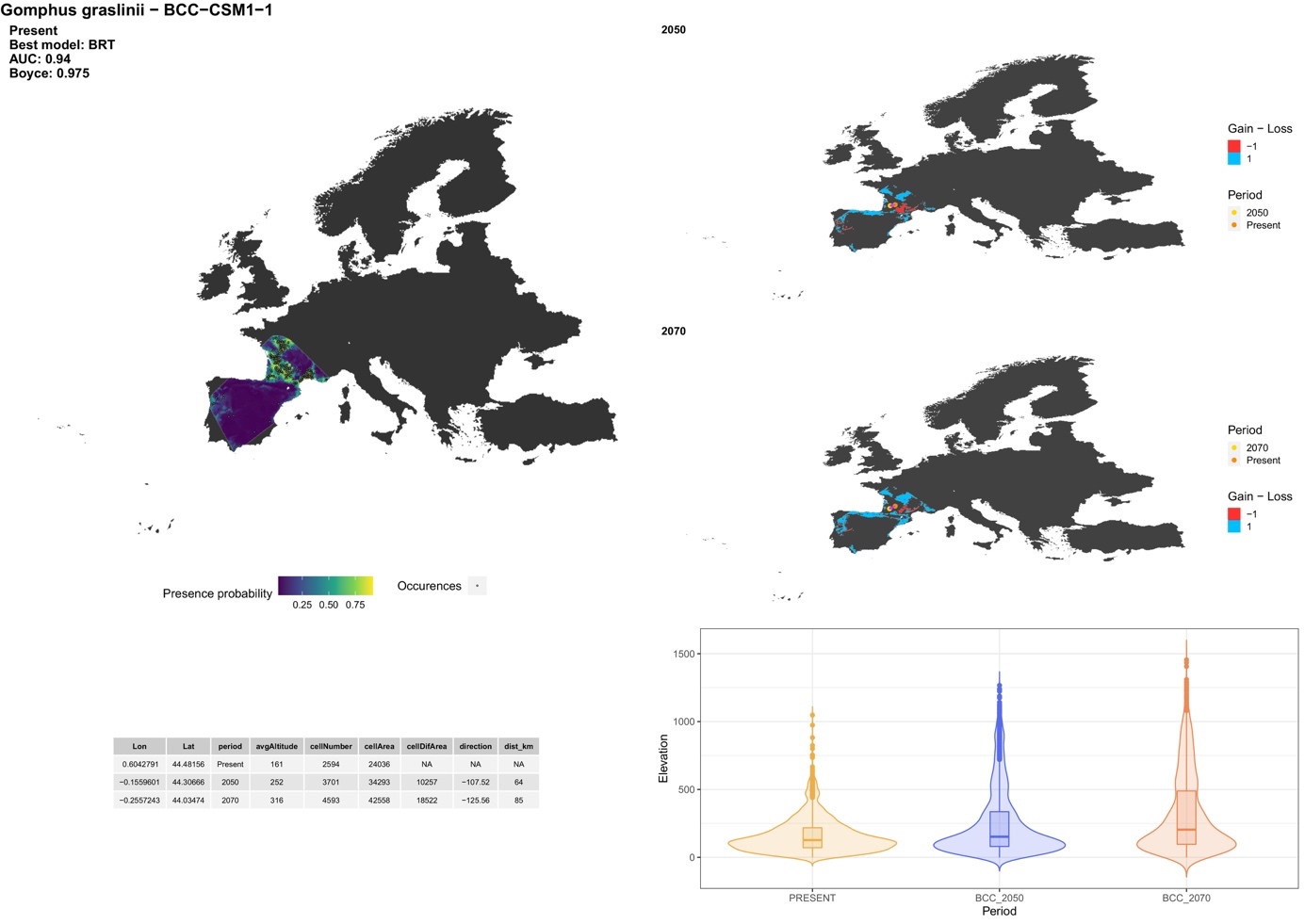

##### *Gomphus pulchellus* Selys, 1840

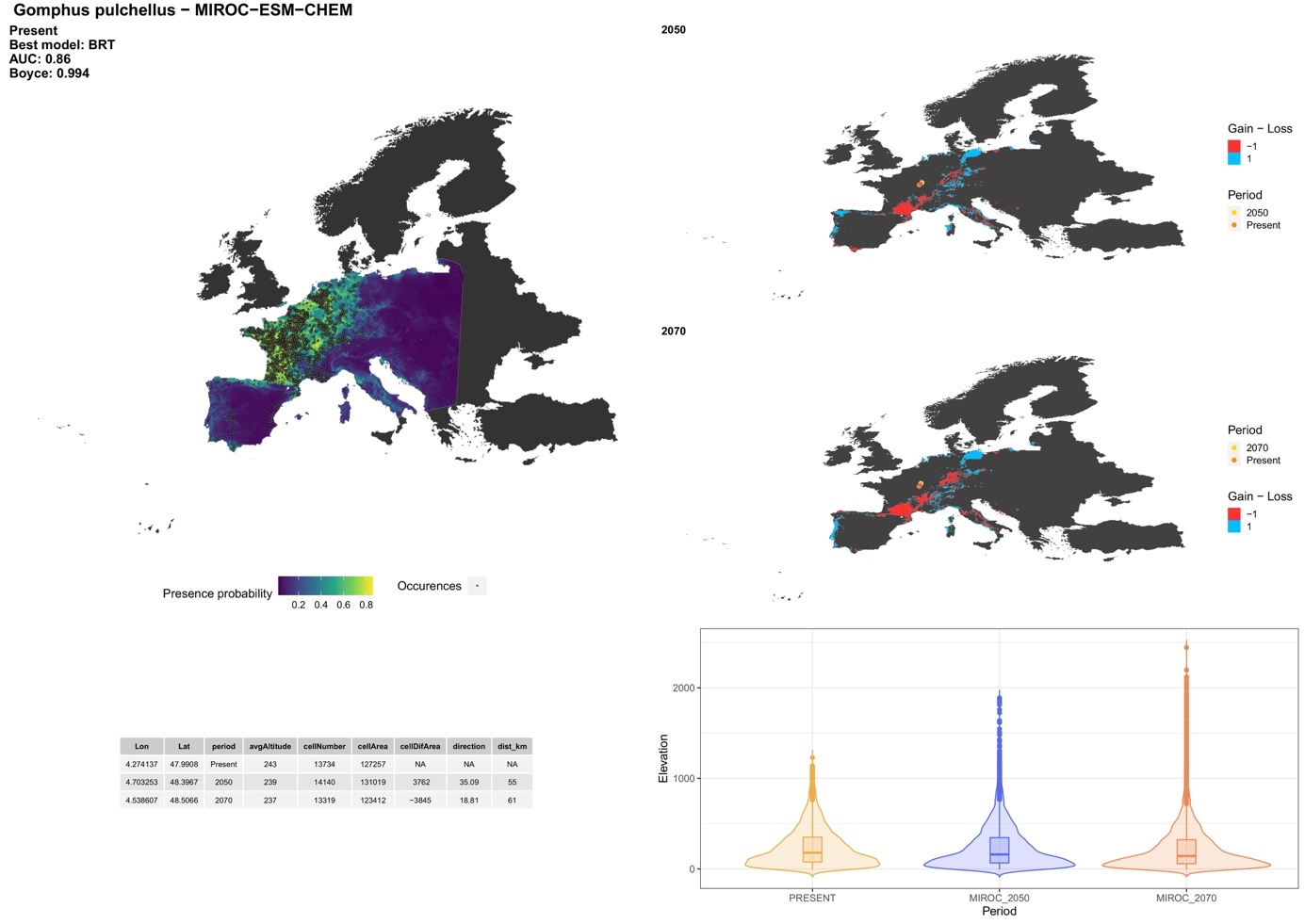

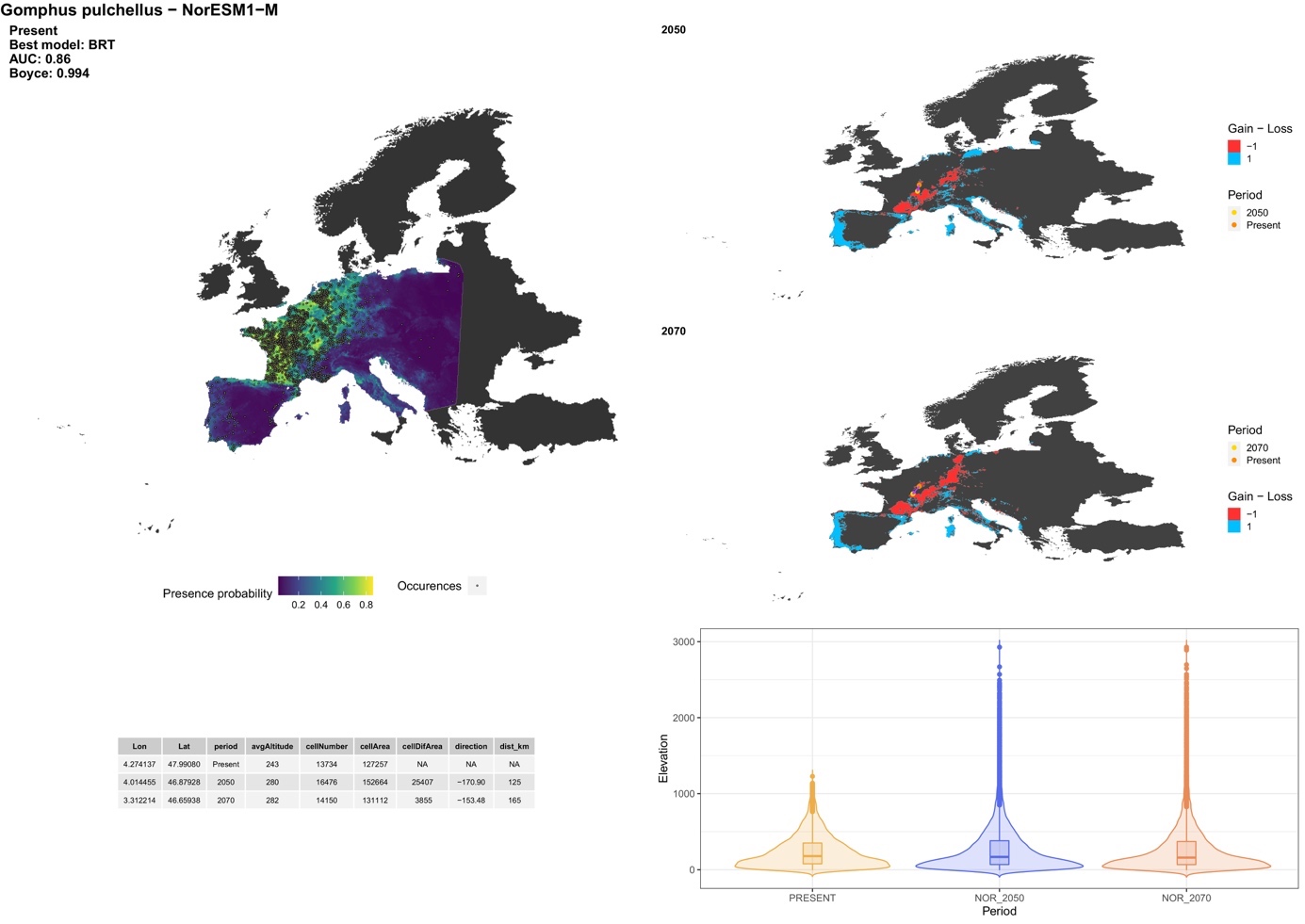

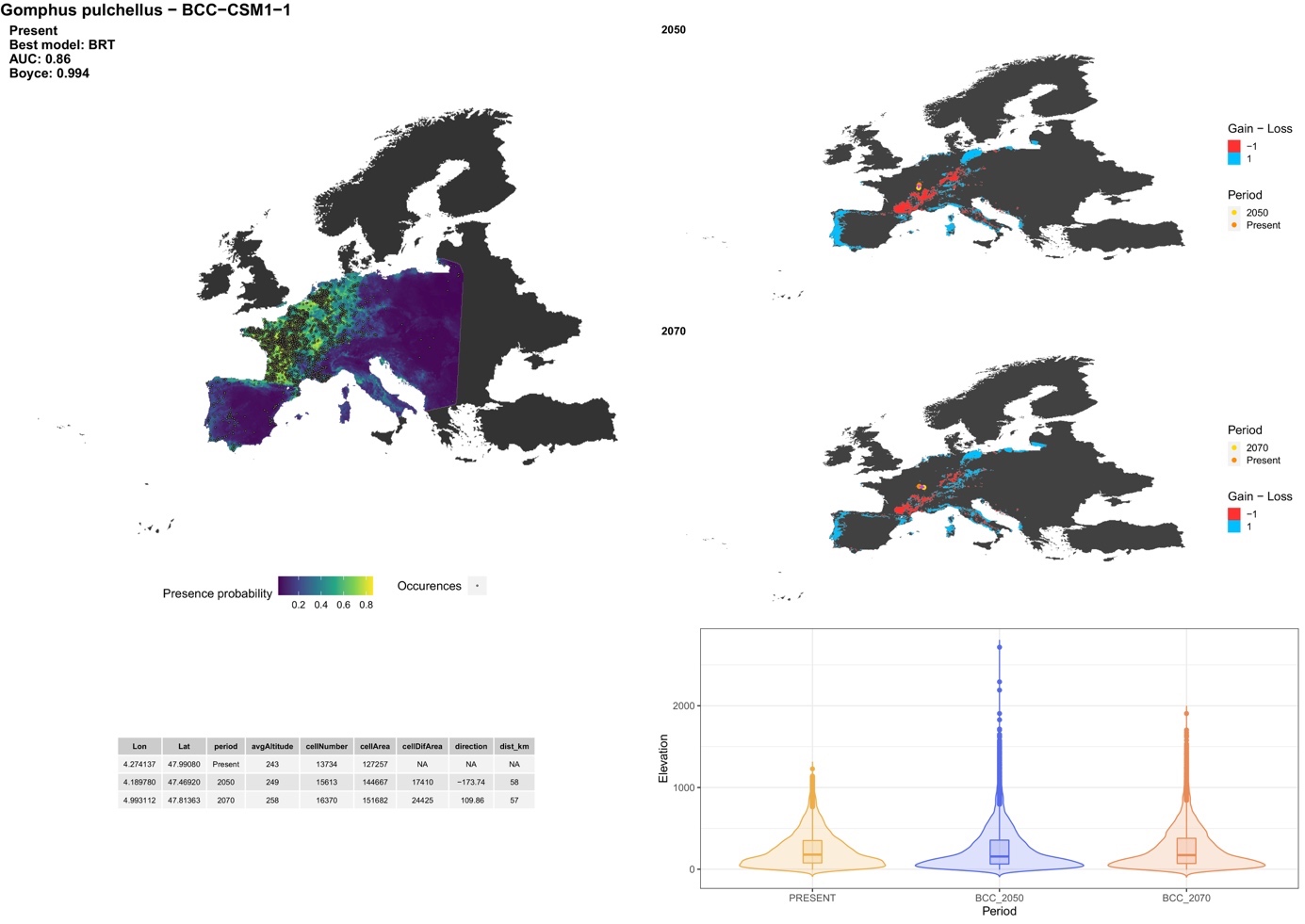

##### *Gomphus simillimus* Selys, 1840

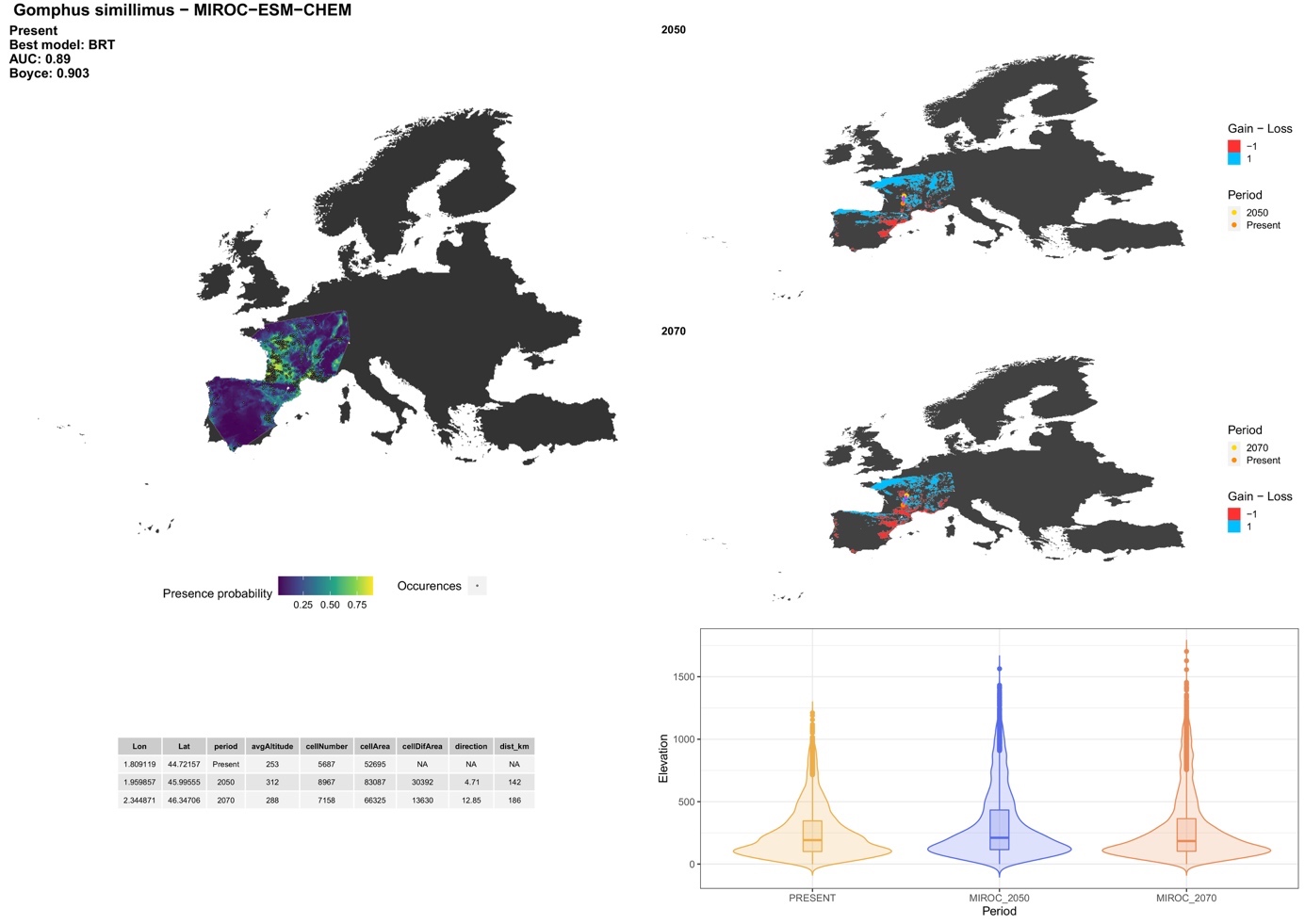

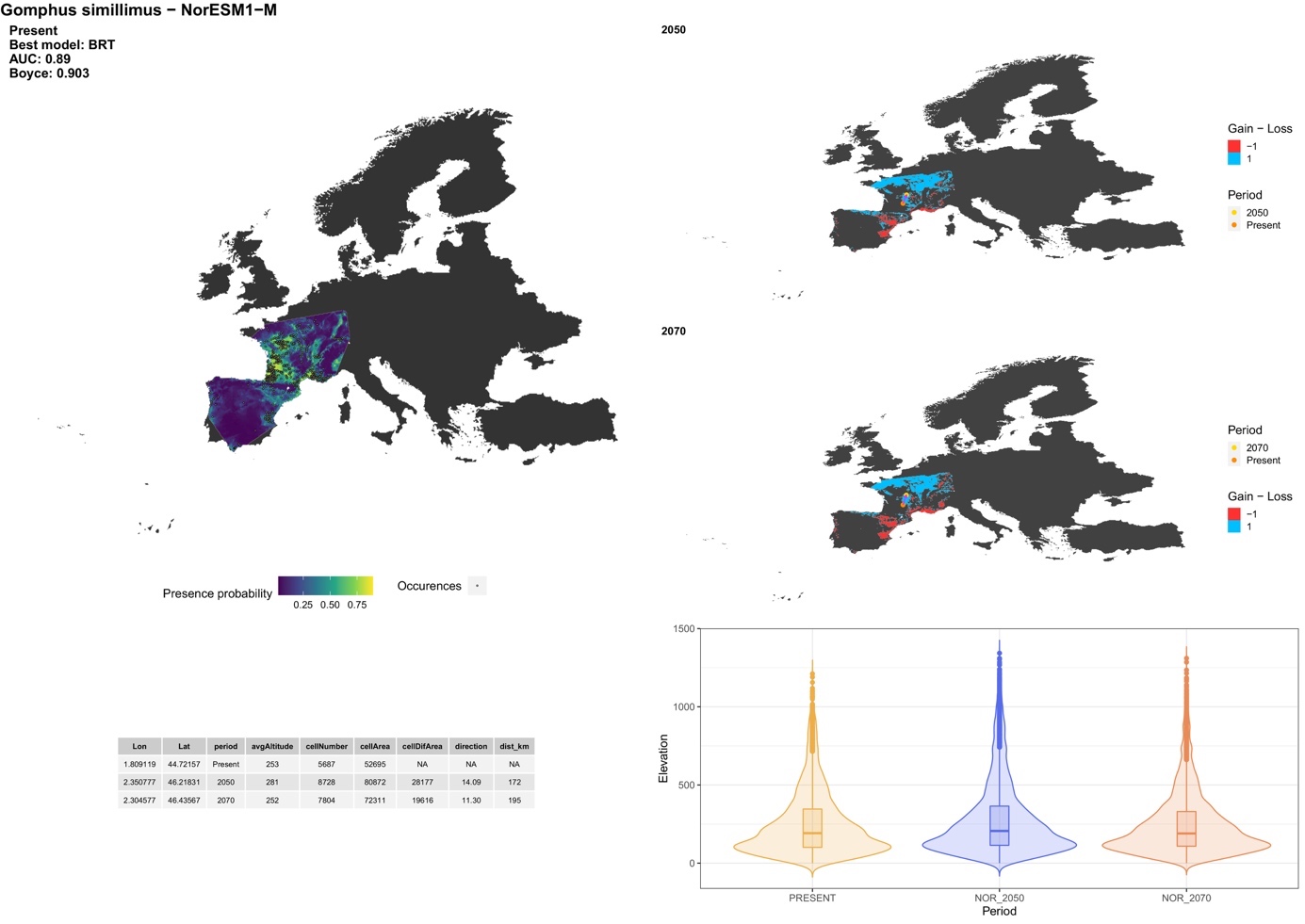

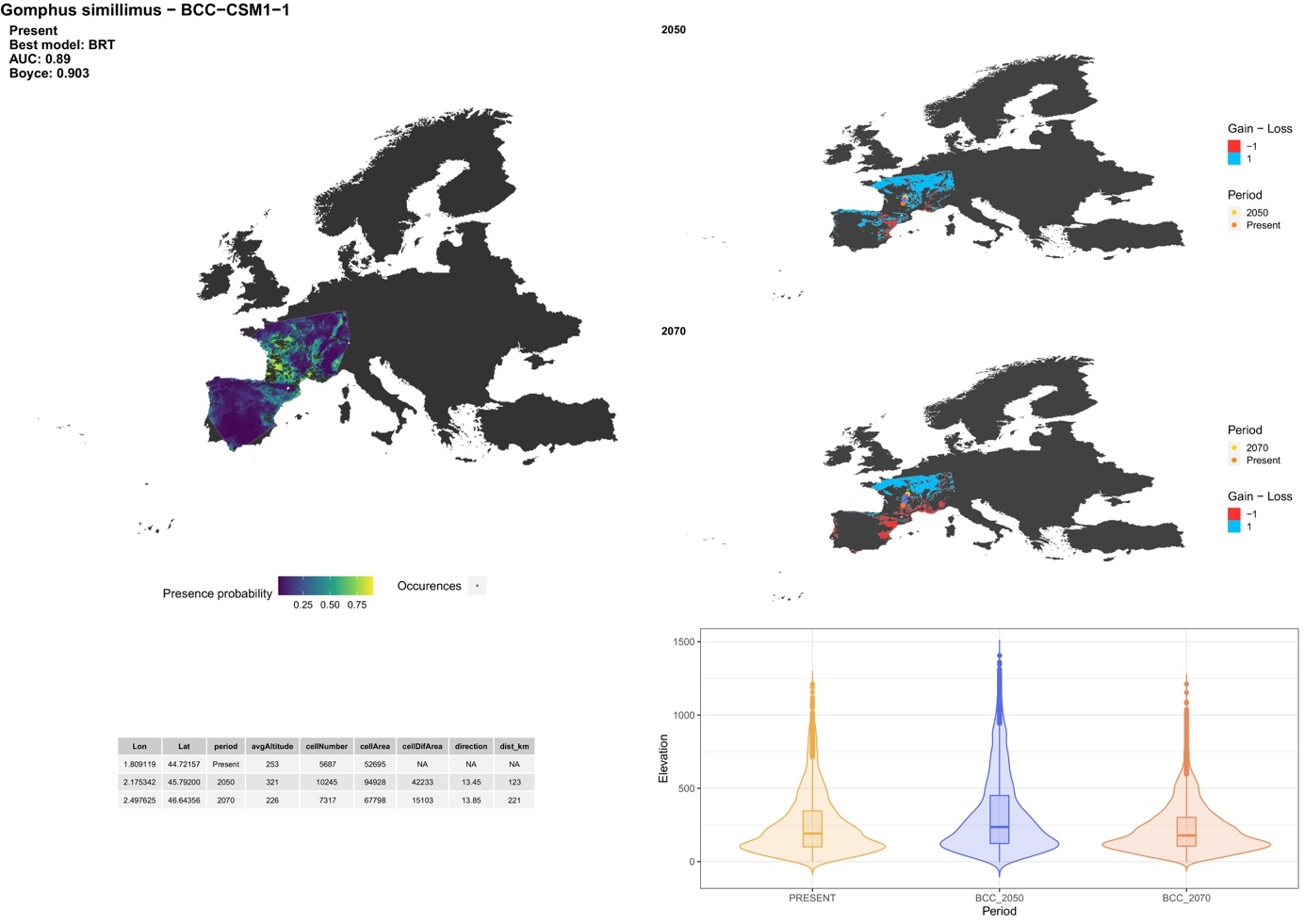

##### *Gomphus vulgatissimus* (Linnaeus, 1758)

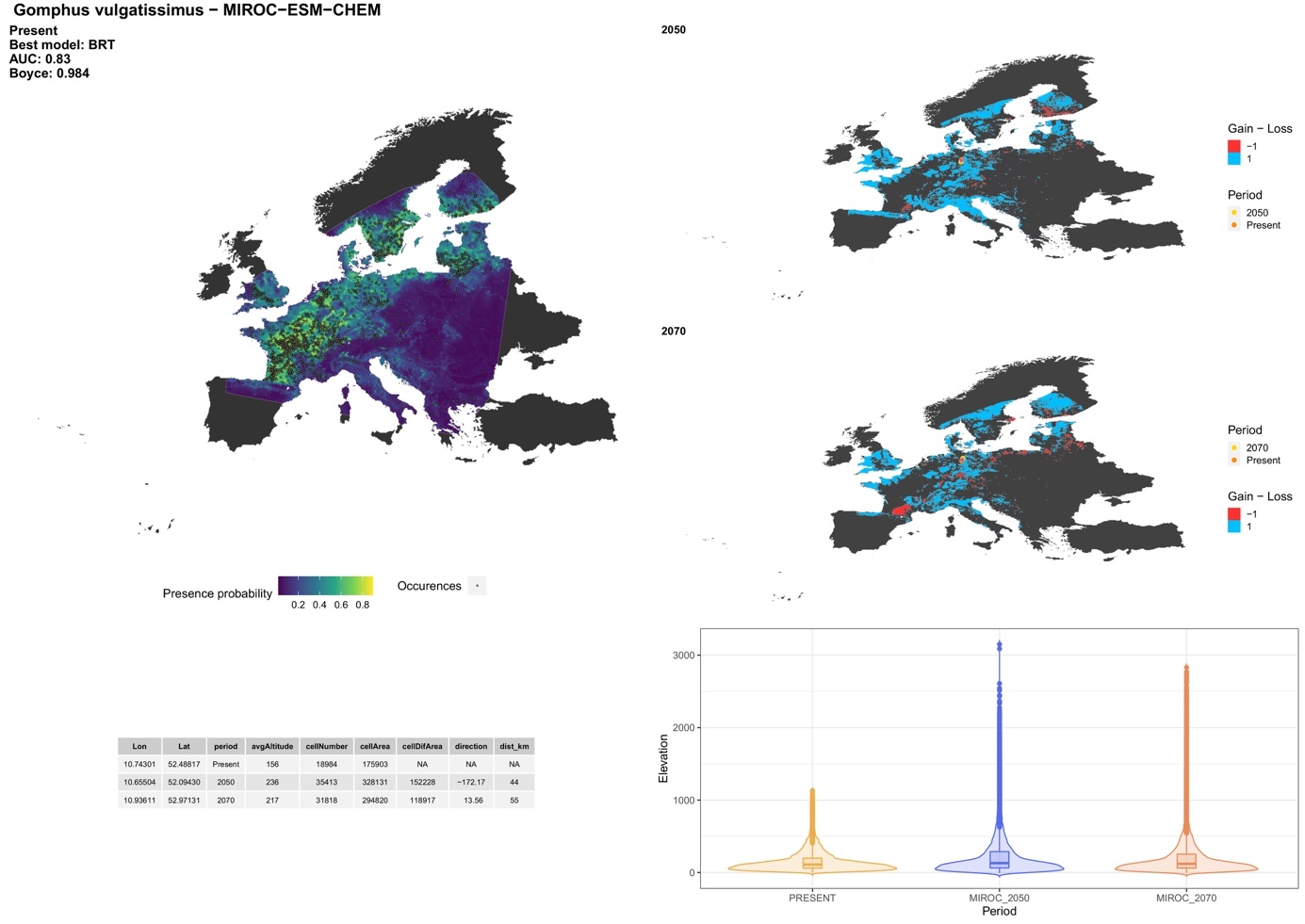

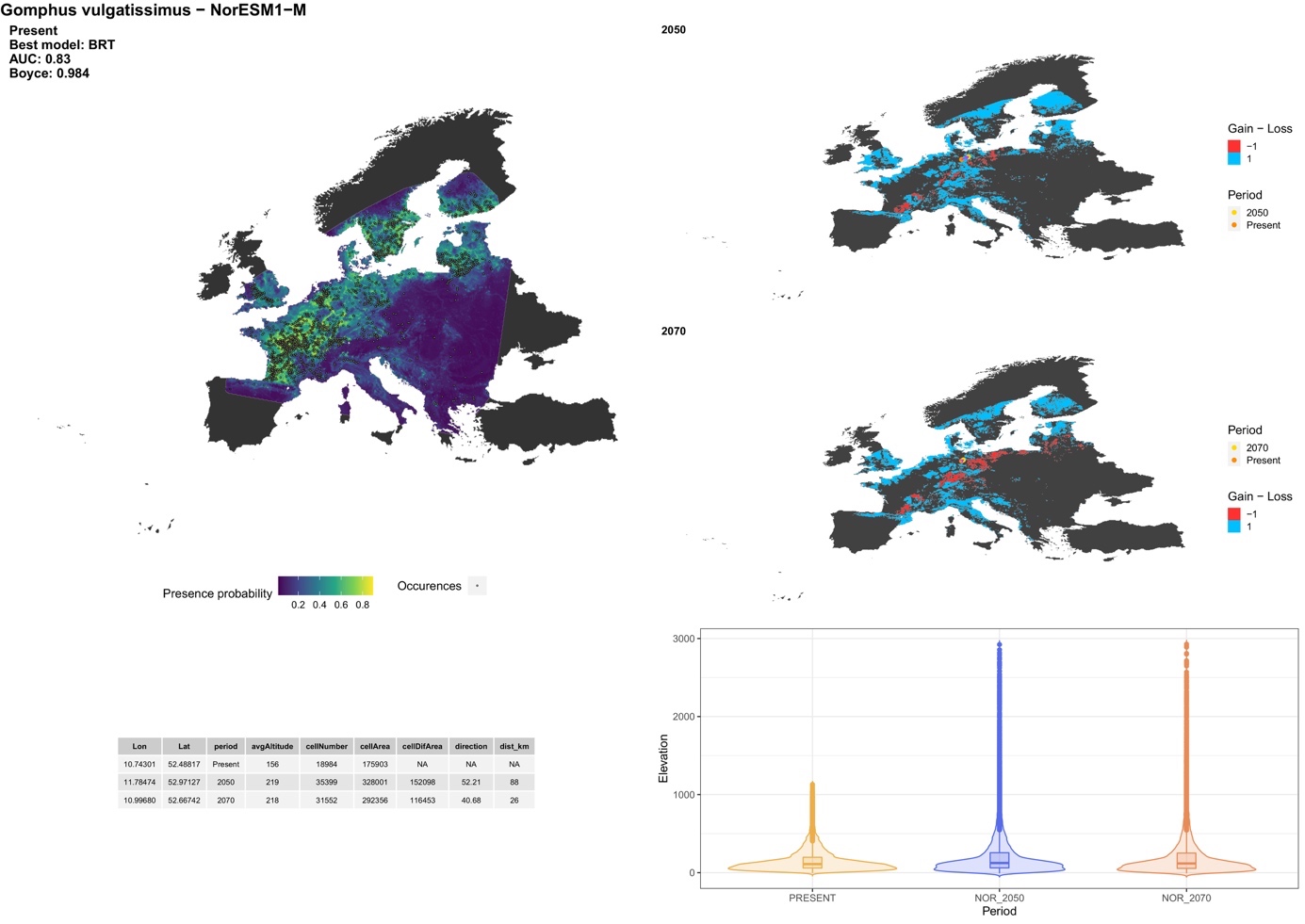

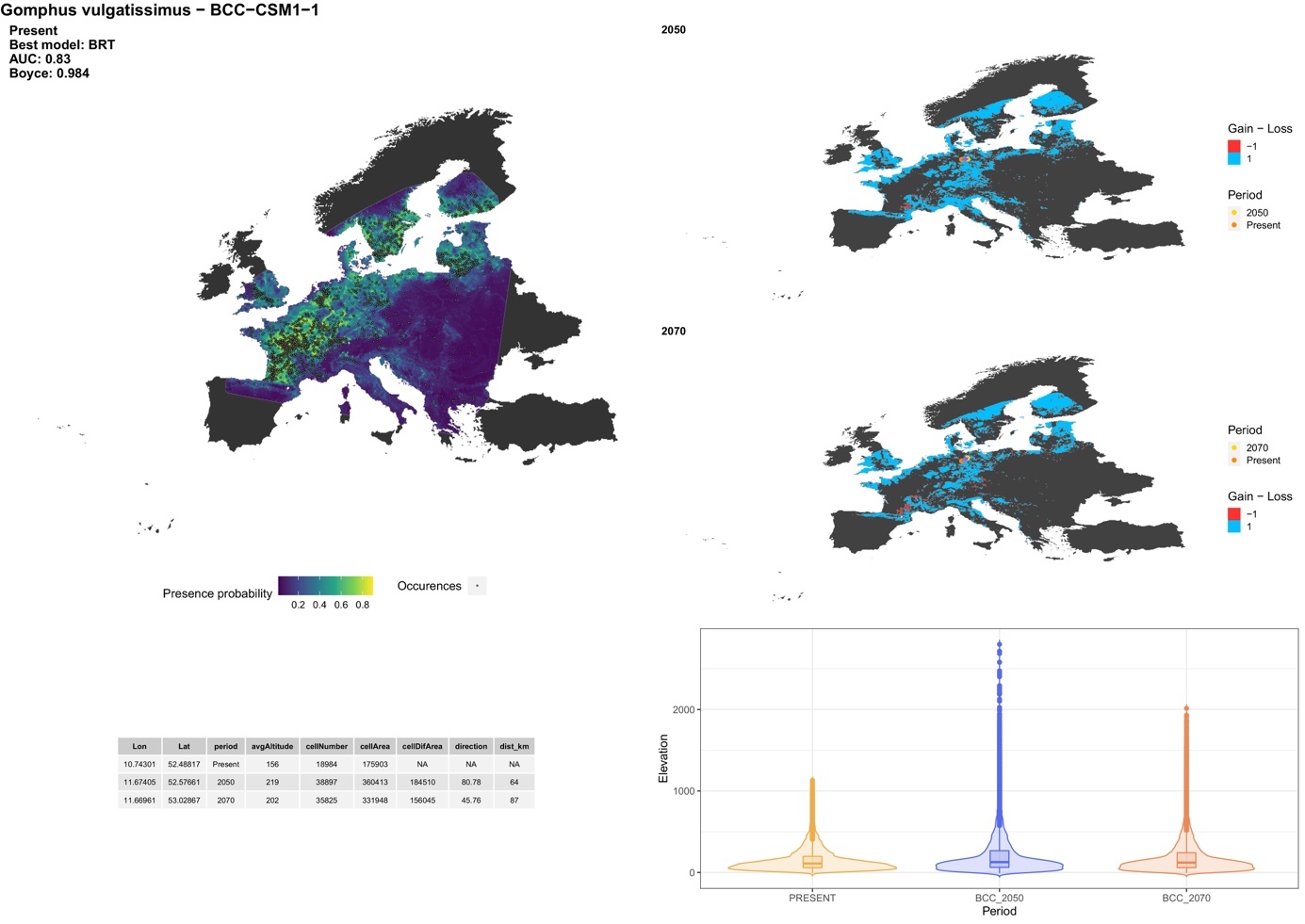

##### *Lindenia tetraphylla* (Vander Linden, 1825)

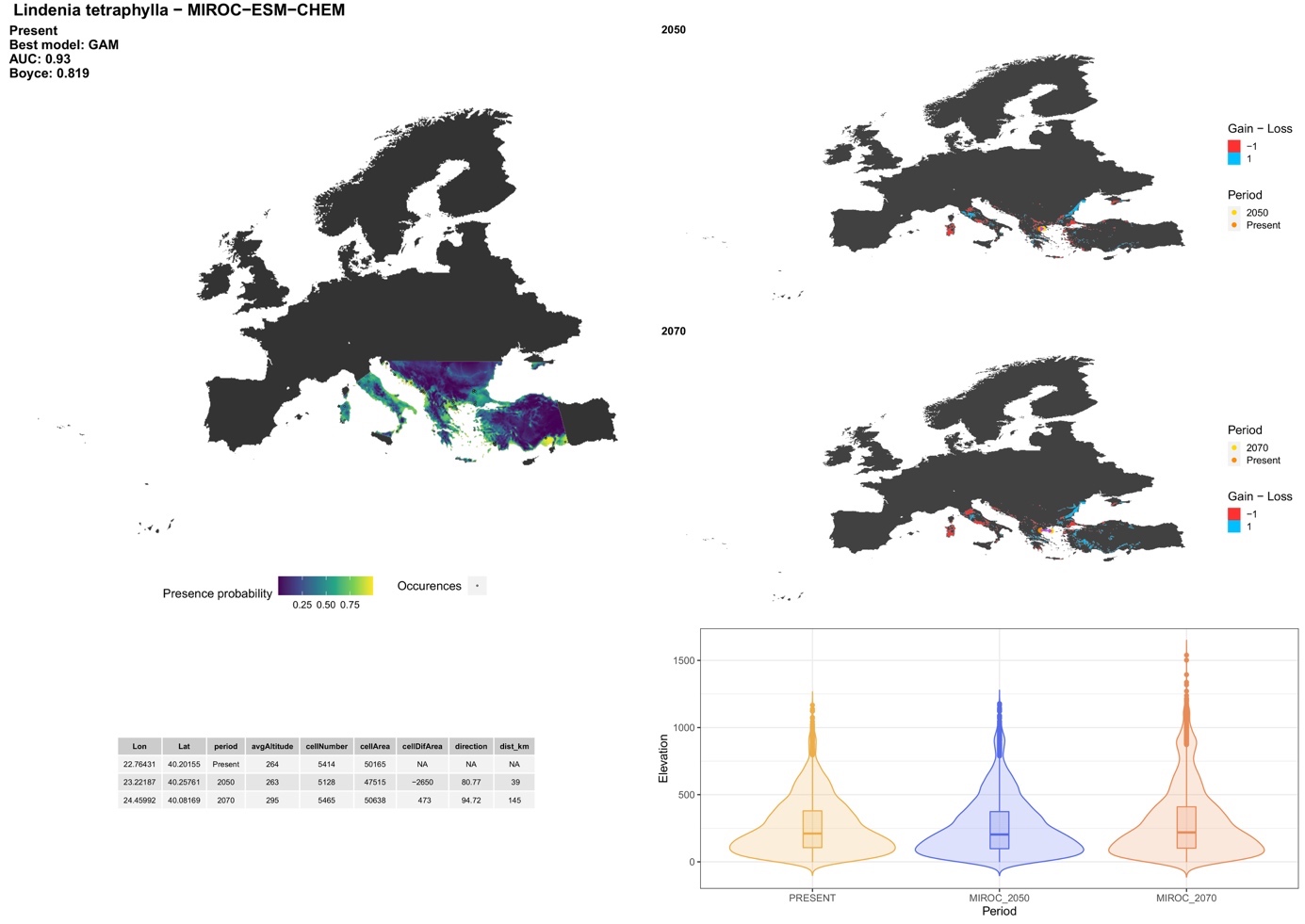

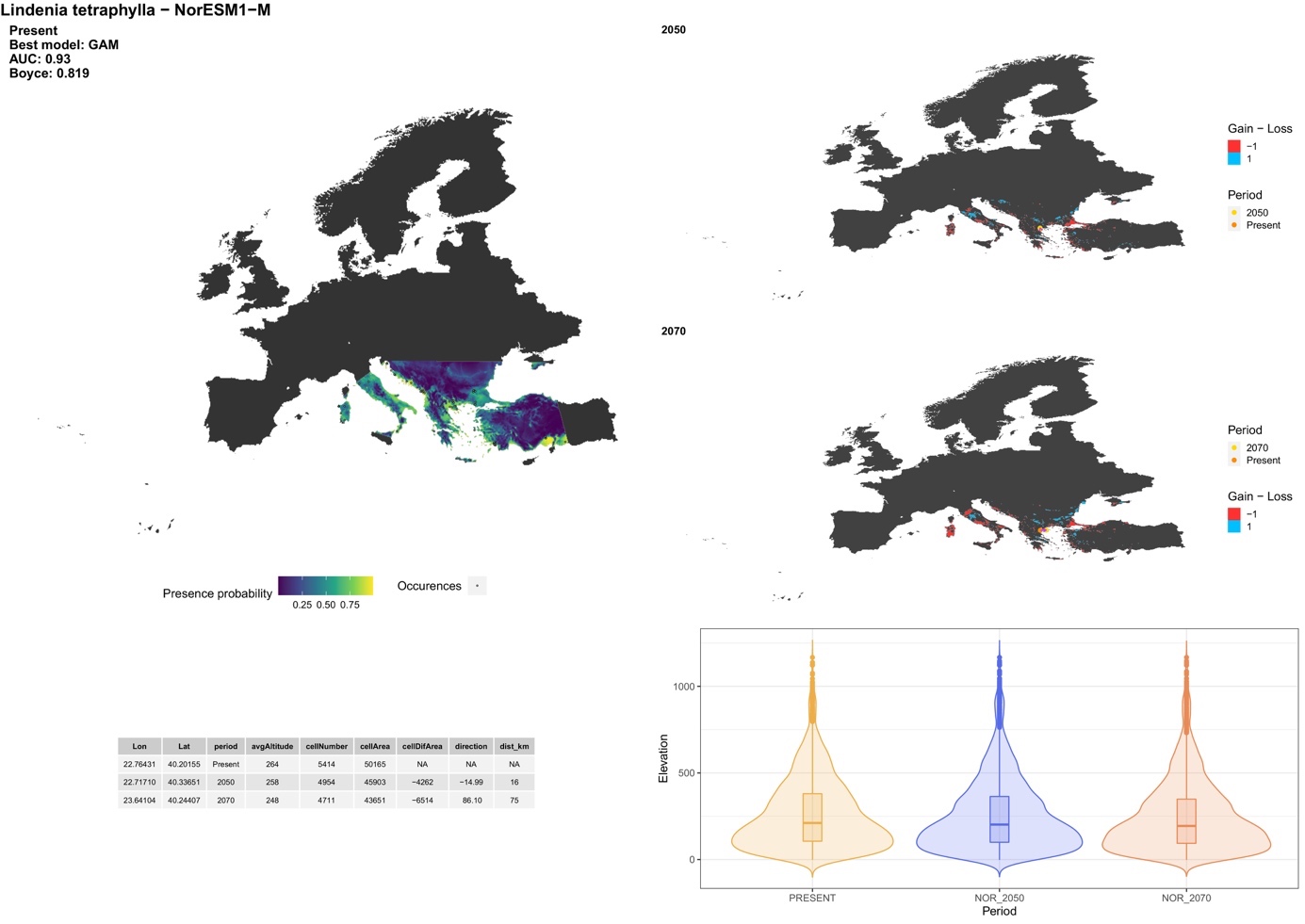

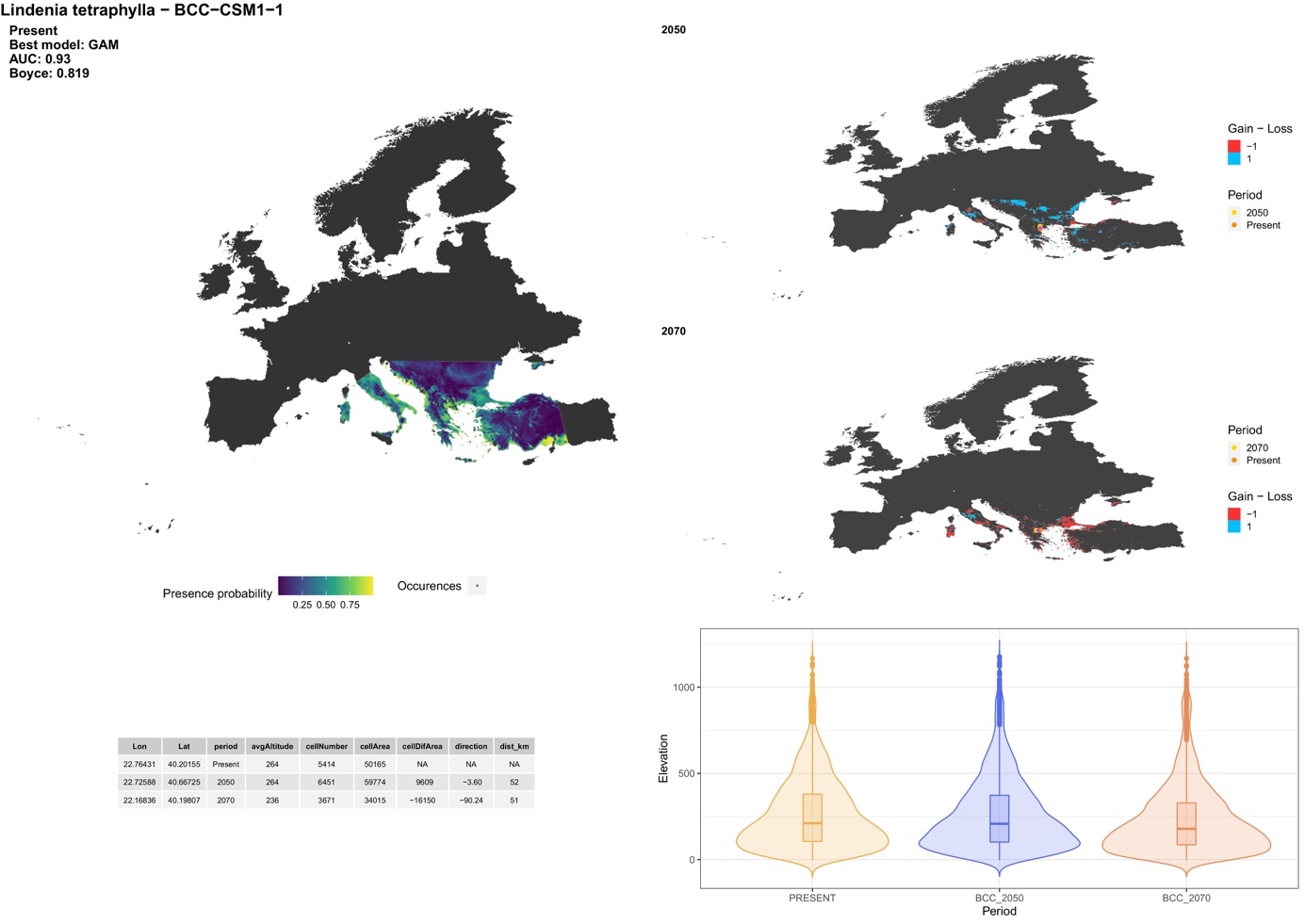

##### *Onychogomphus forcipatus* (Linnaeus, 1758)

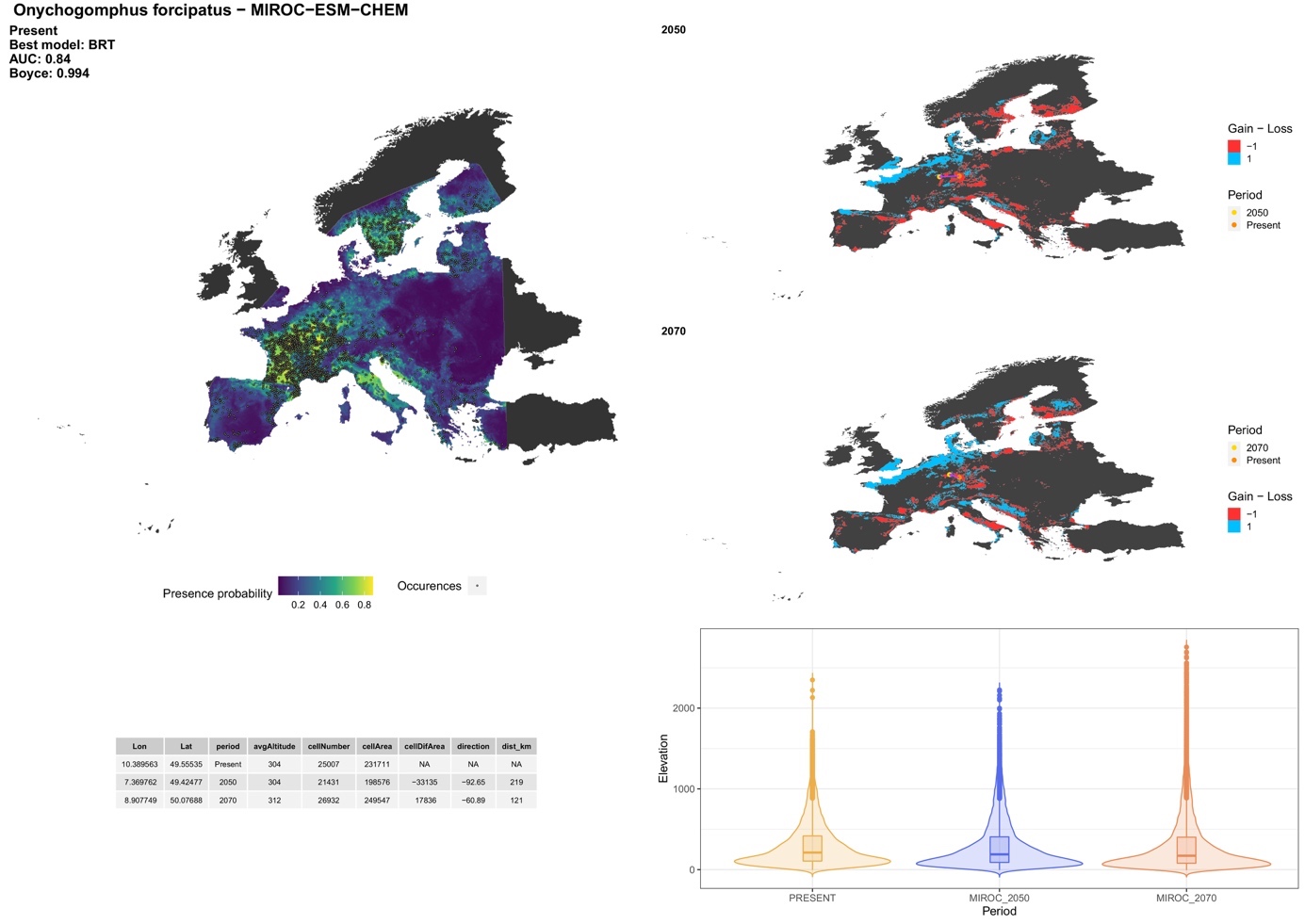

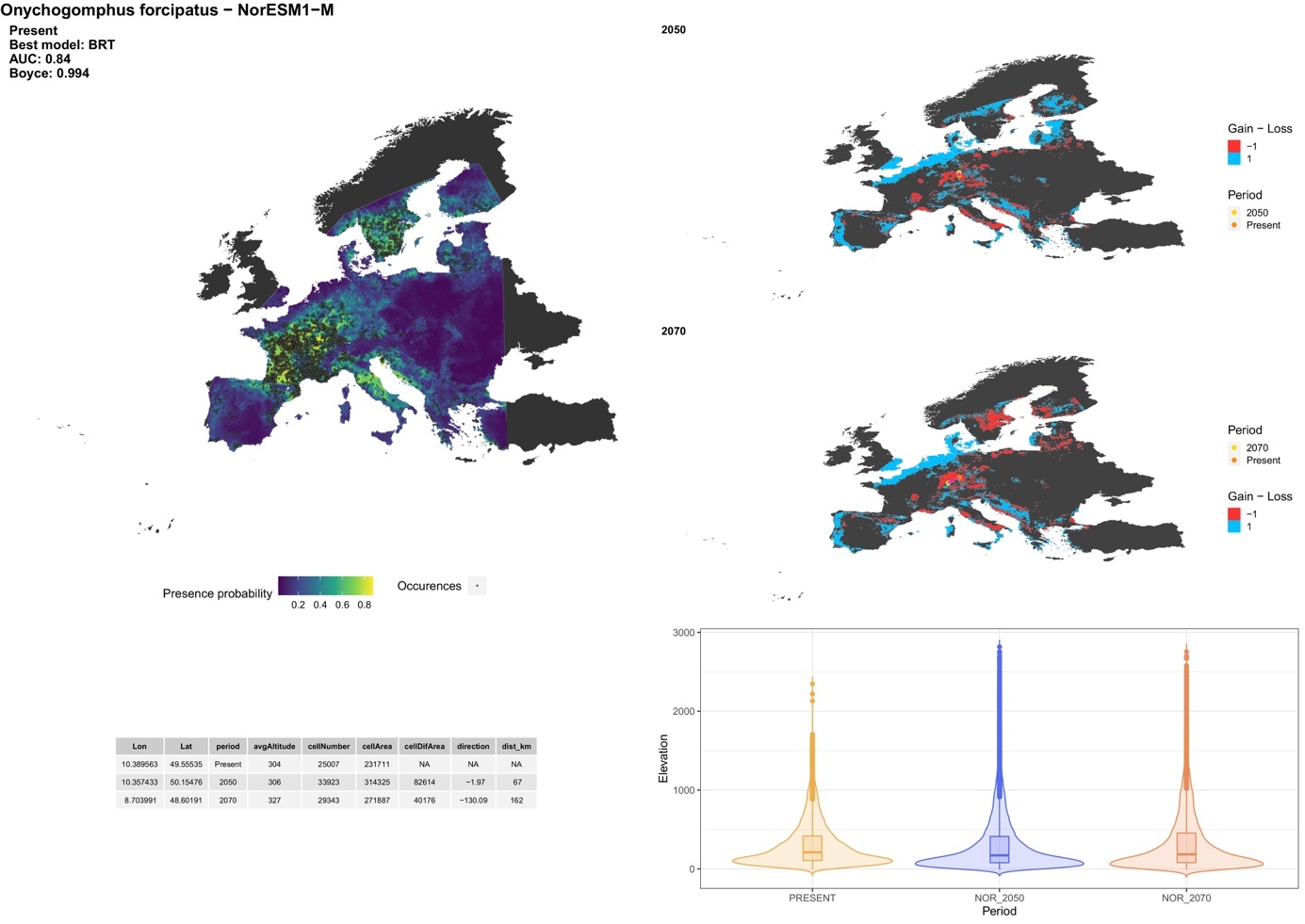

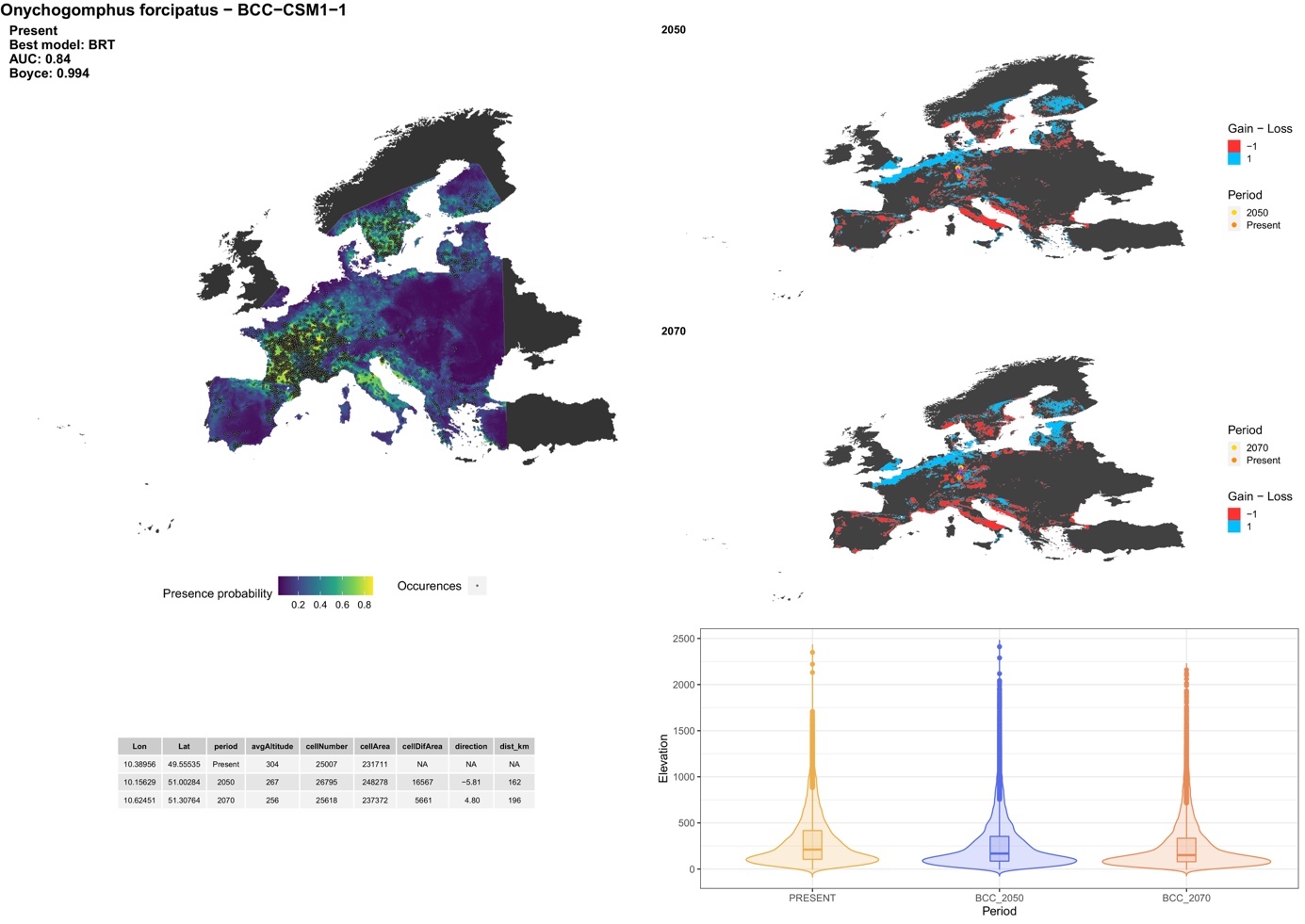

##### *Onychogomphus uncatus* (Charpentier, 1840)

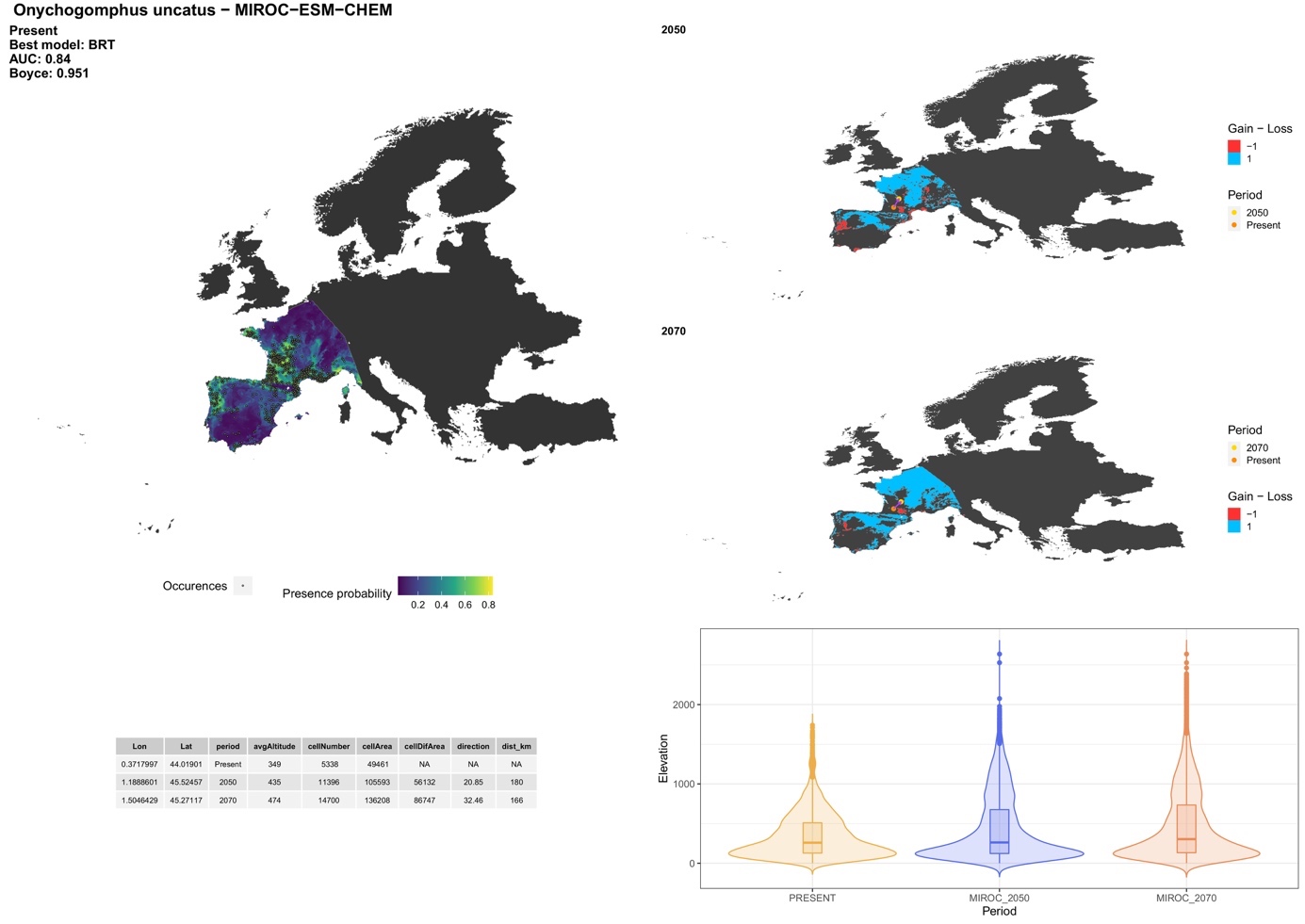

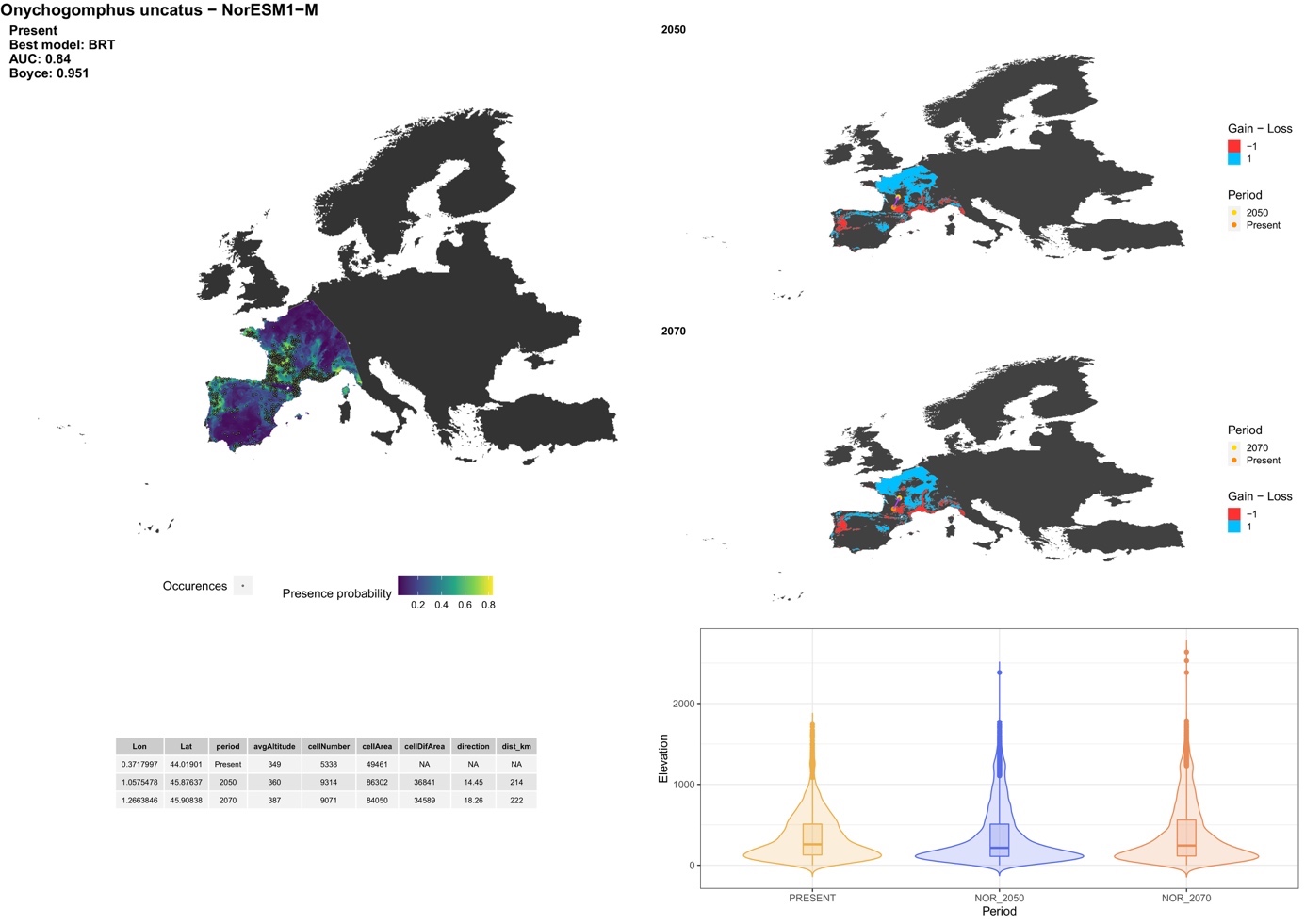

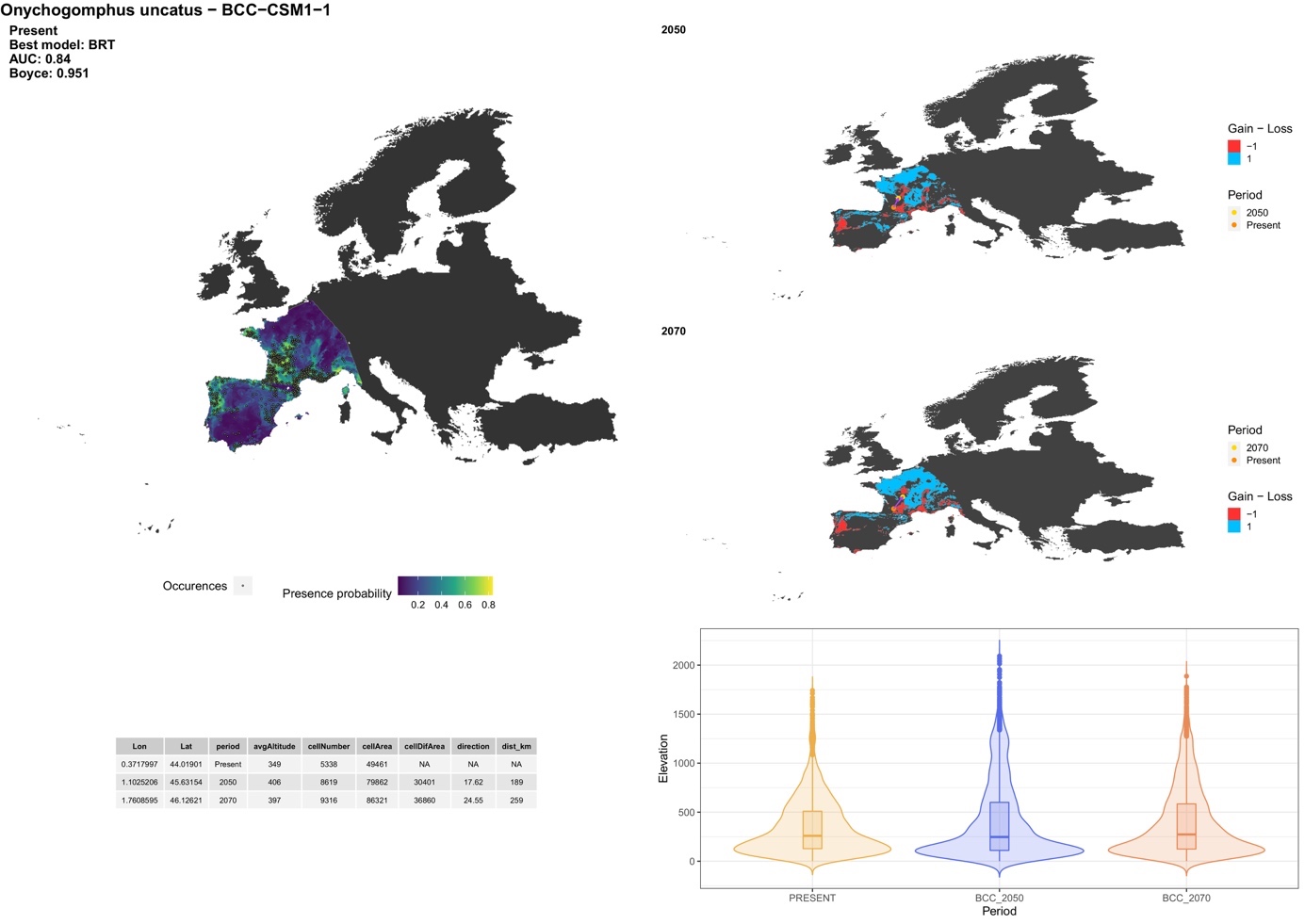

##### *Ophiogomphus cecilia* (Geoffroy in Fourcroy, 1785)

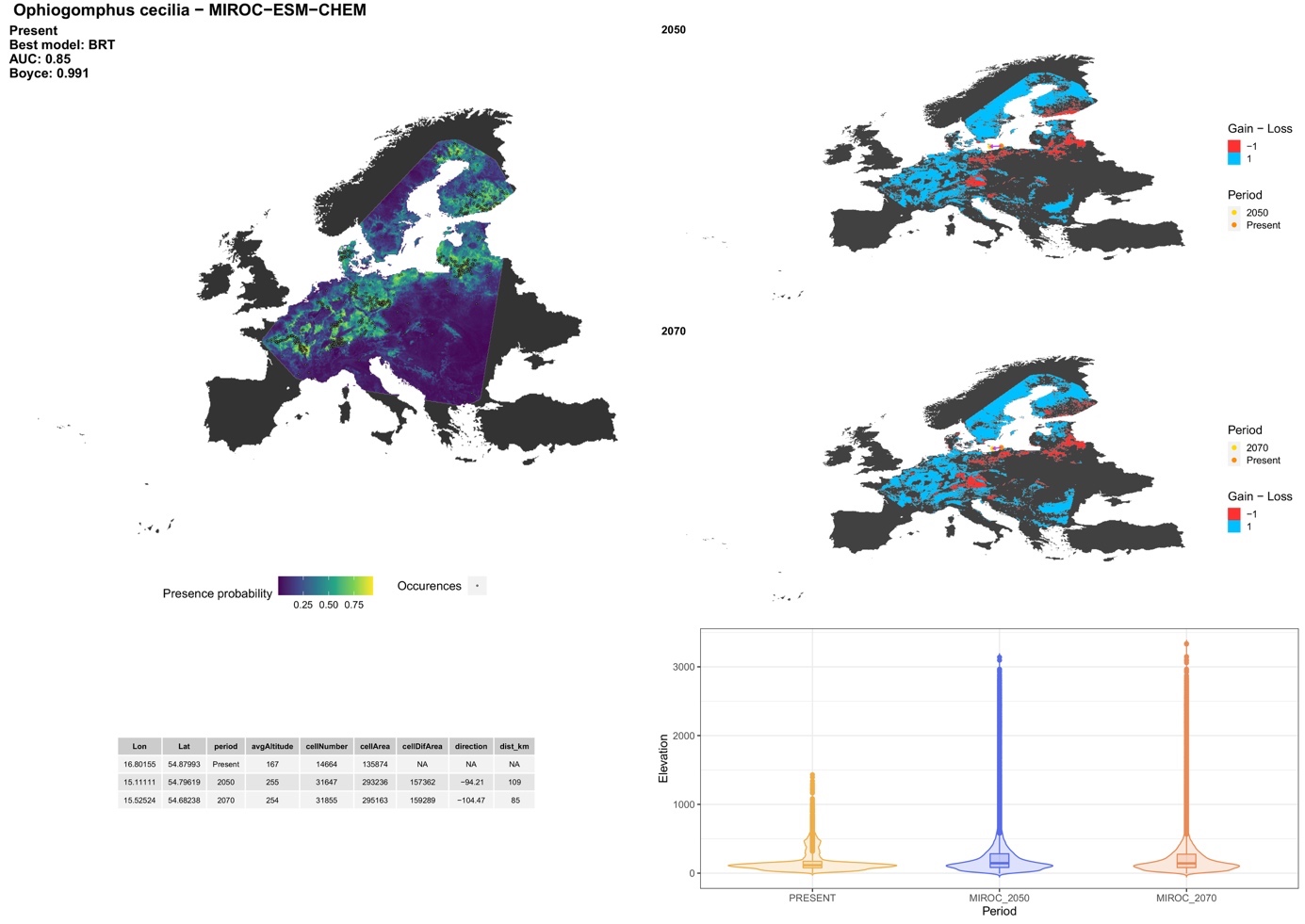

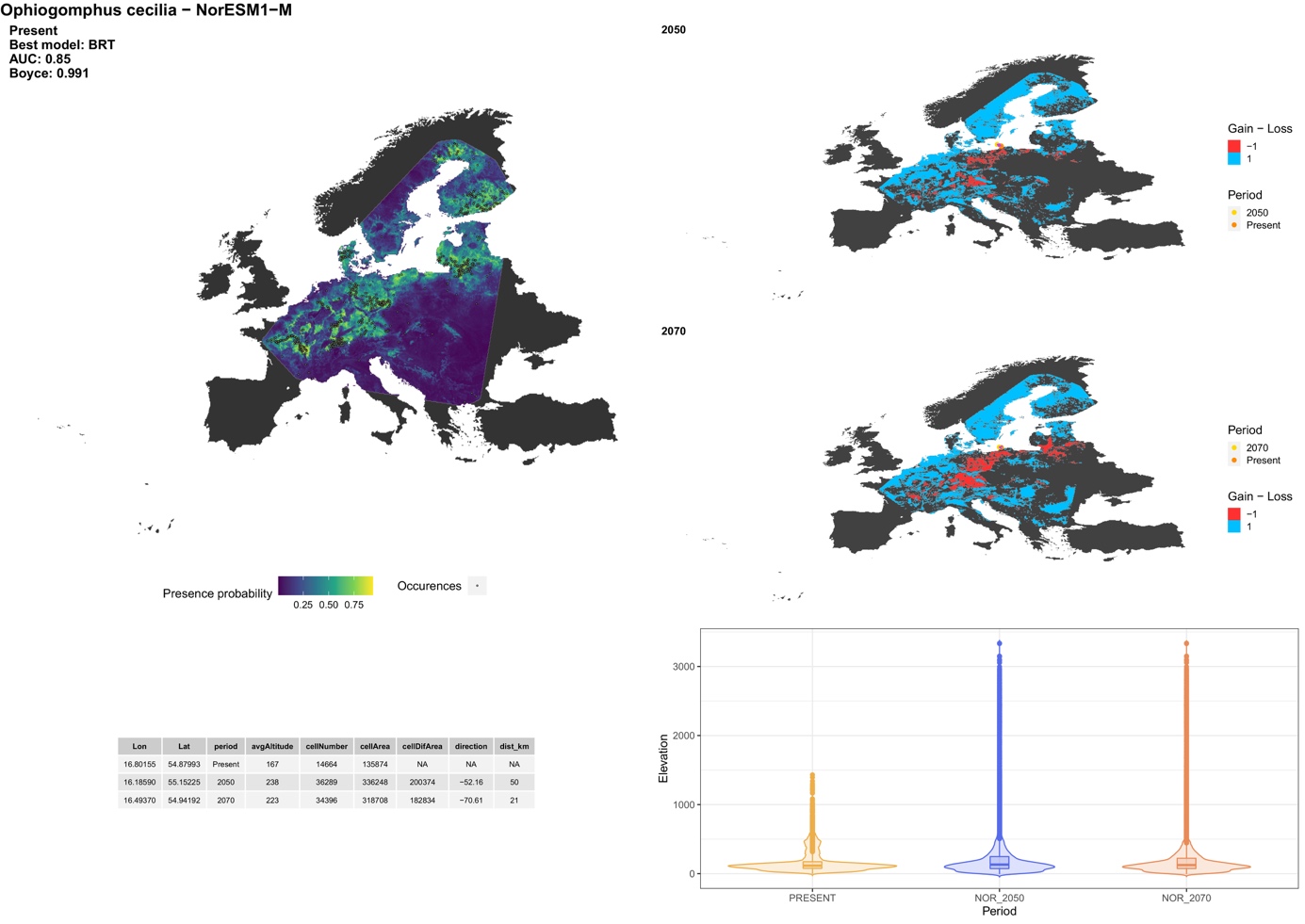

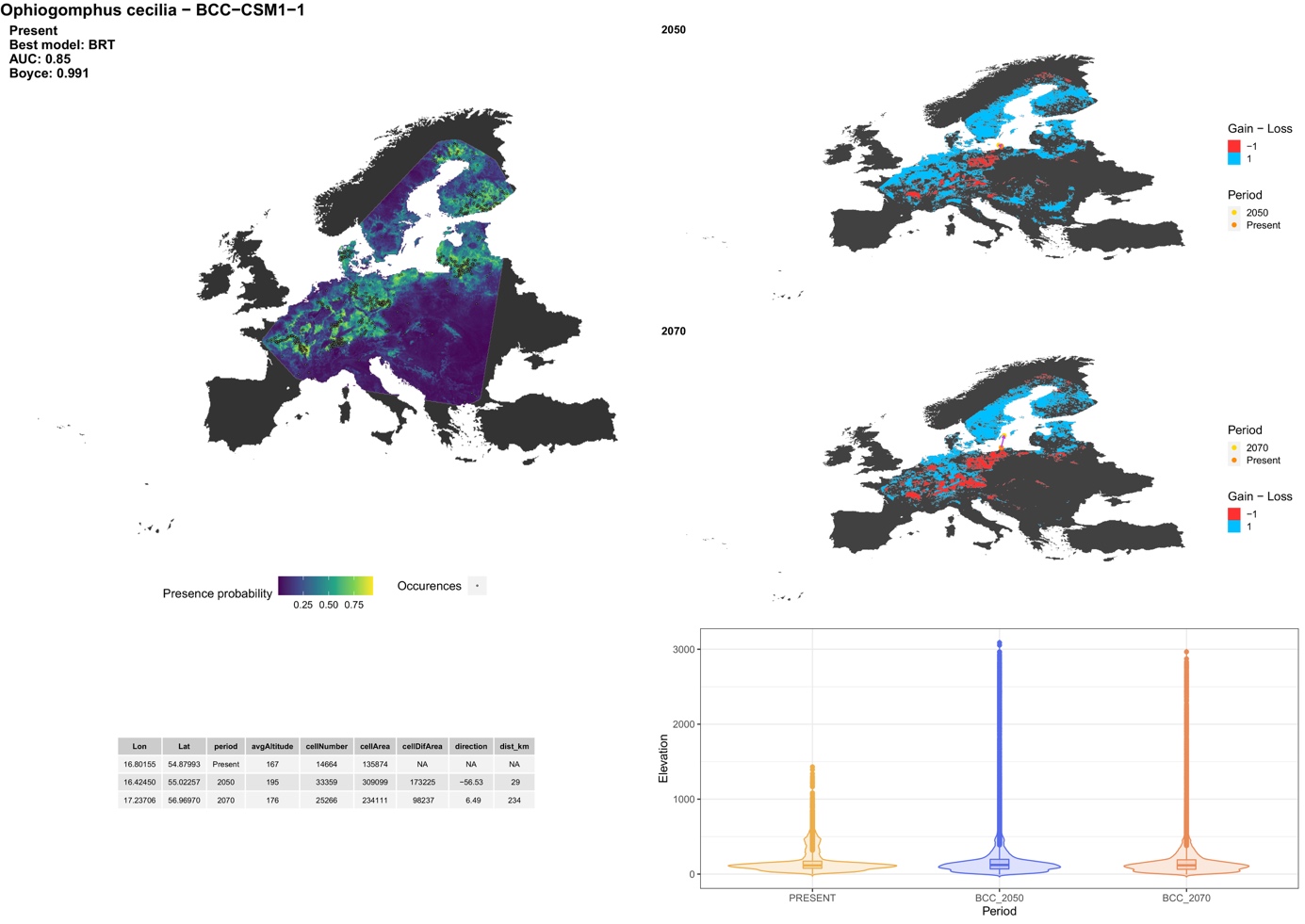

##### *Stylurus flavipes* (Charpentier, 1825)

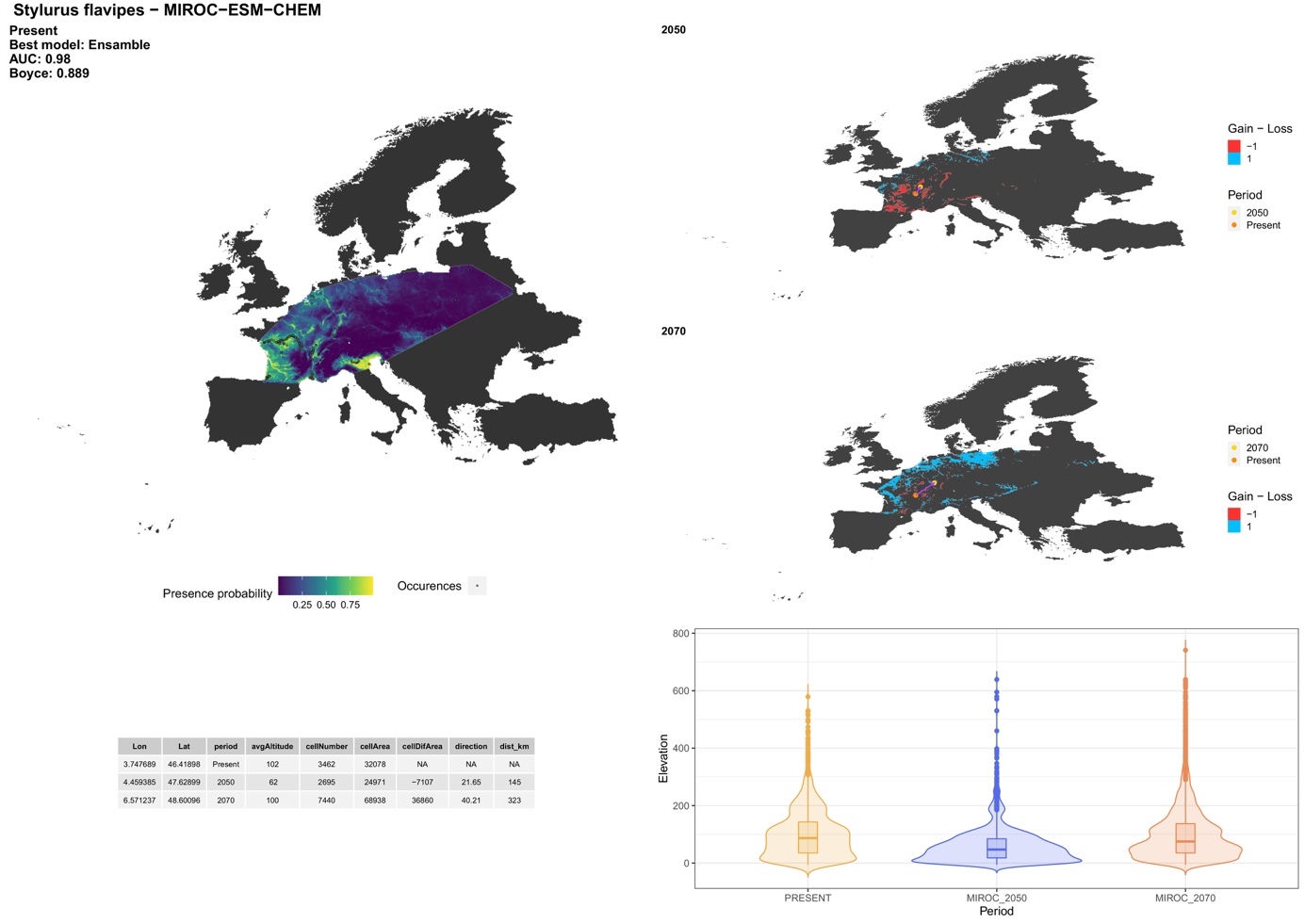

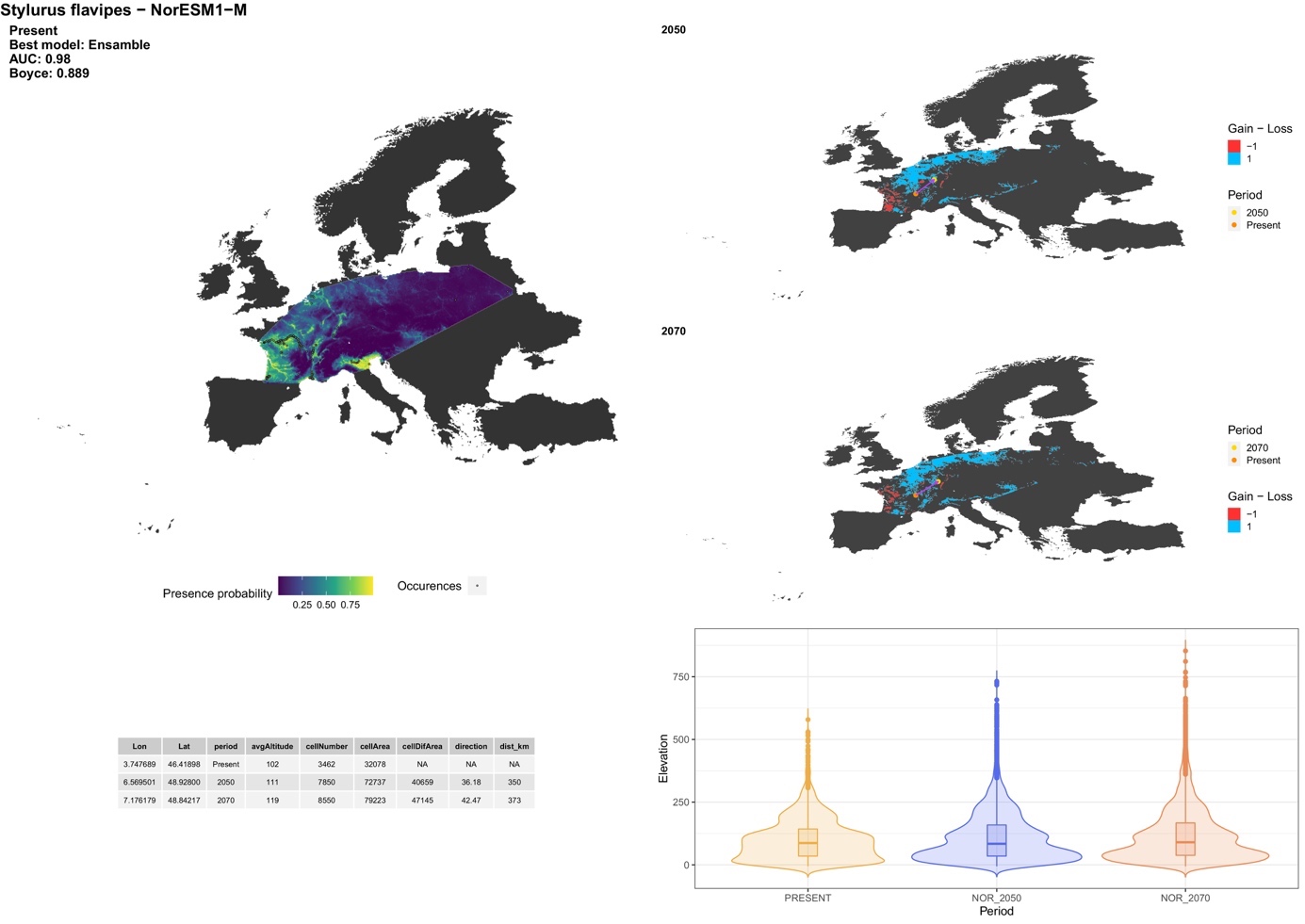

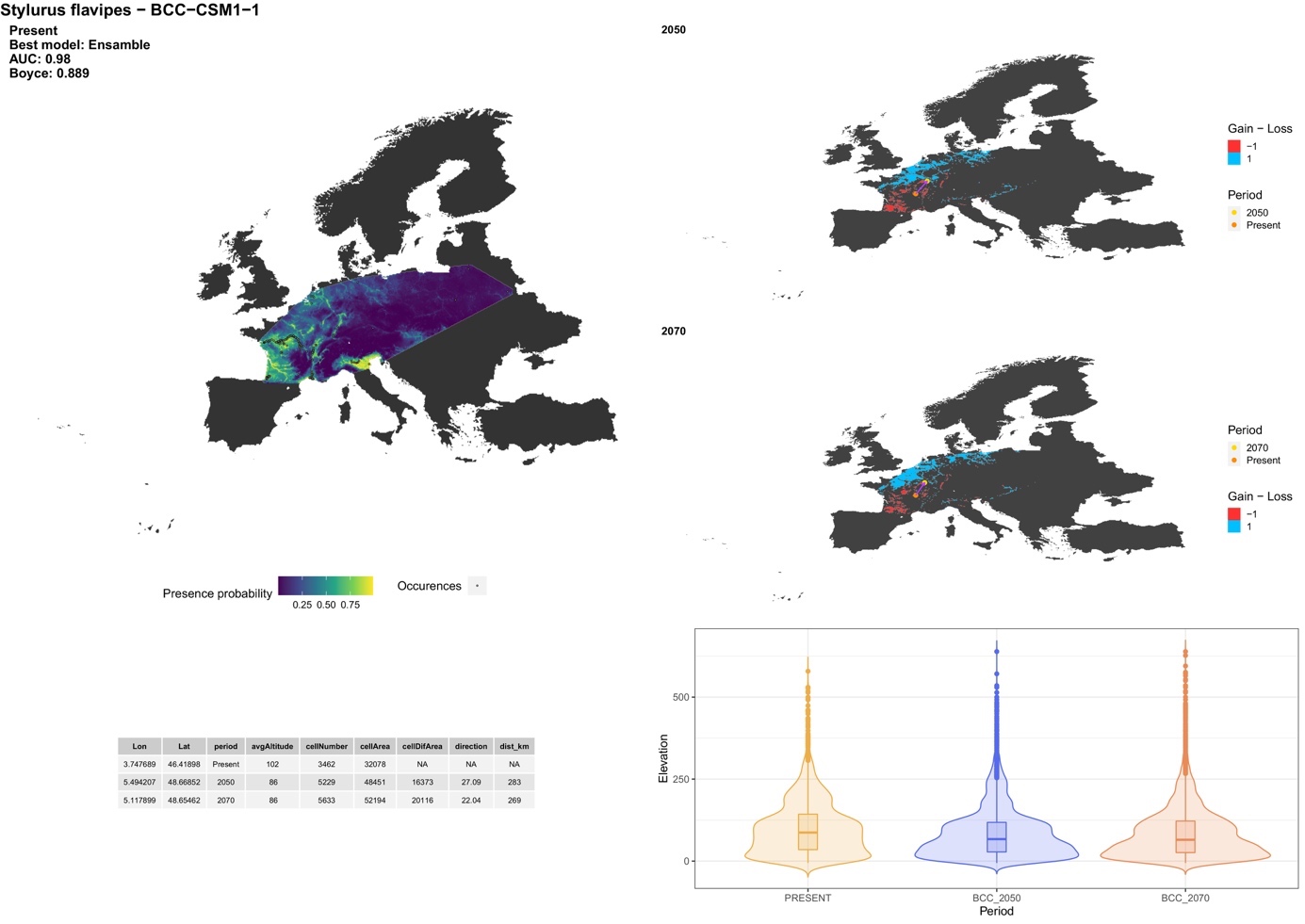

##### **Family: Cordulegastridae**

##### *Cordulegaster bidentata* Selys, 1843

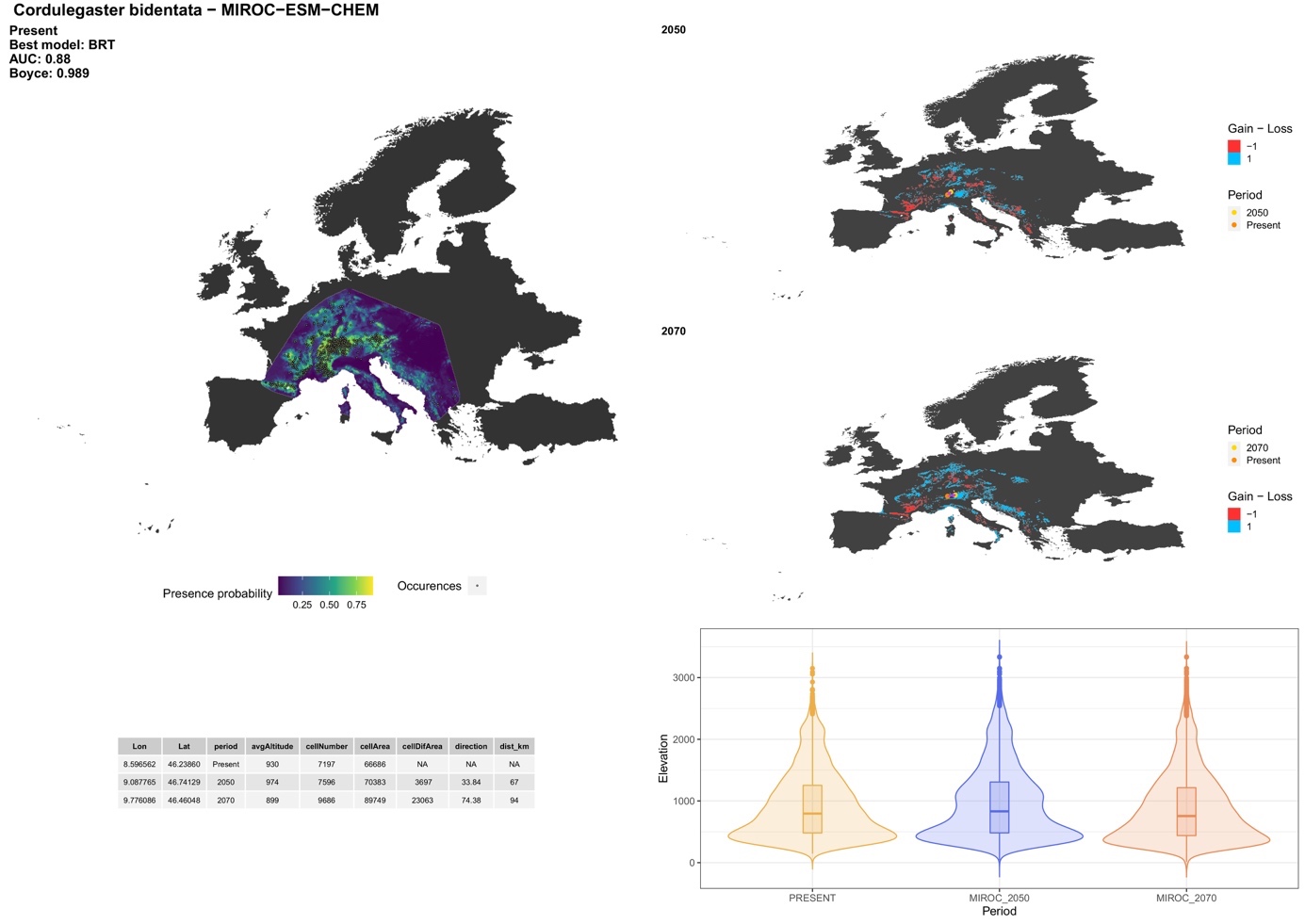

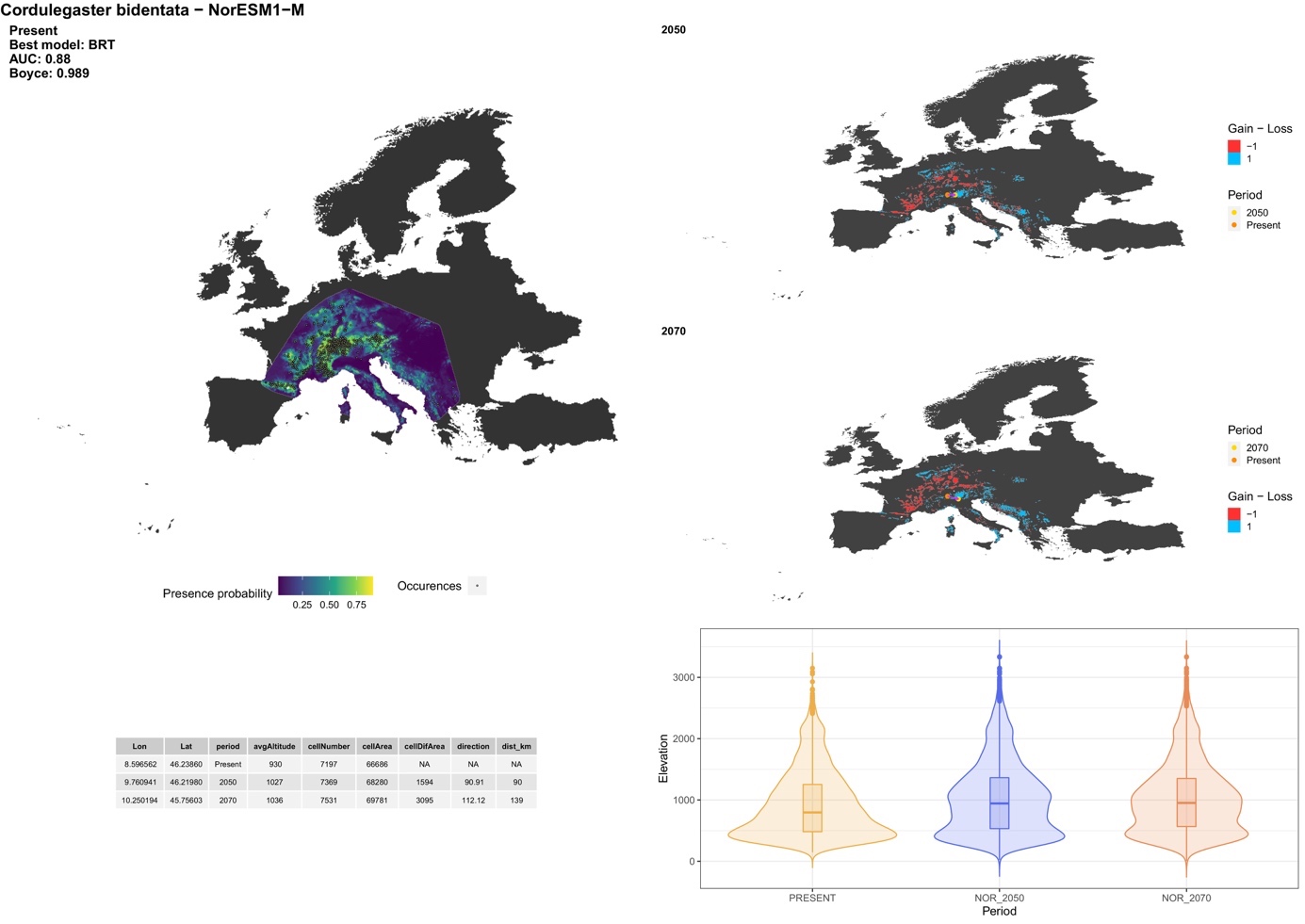

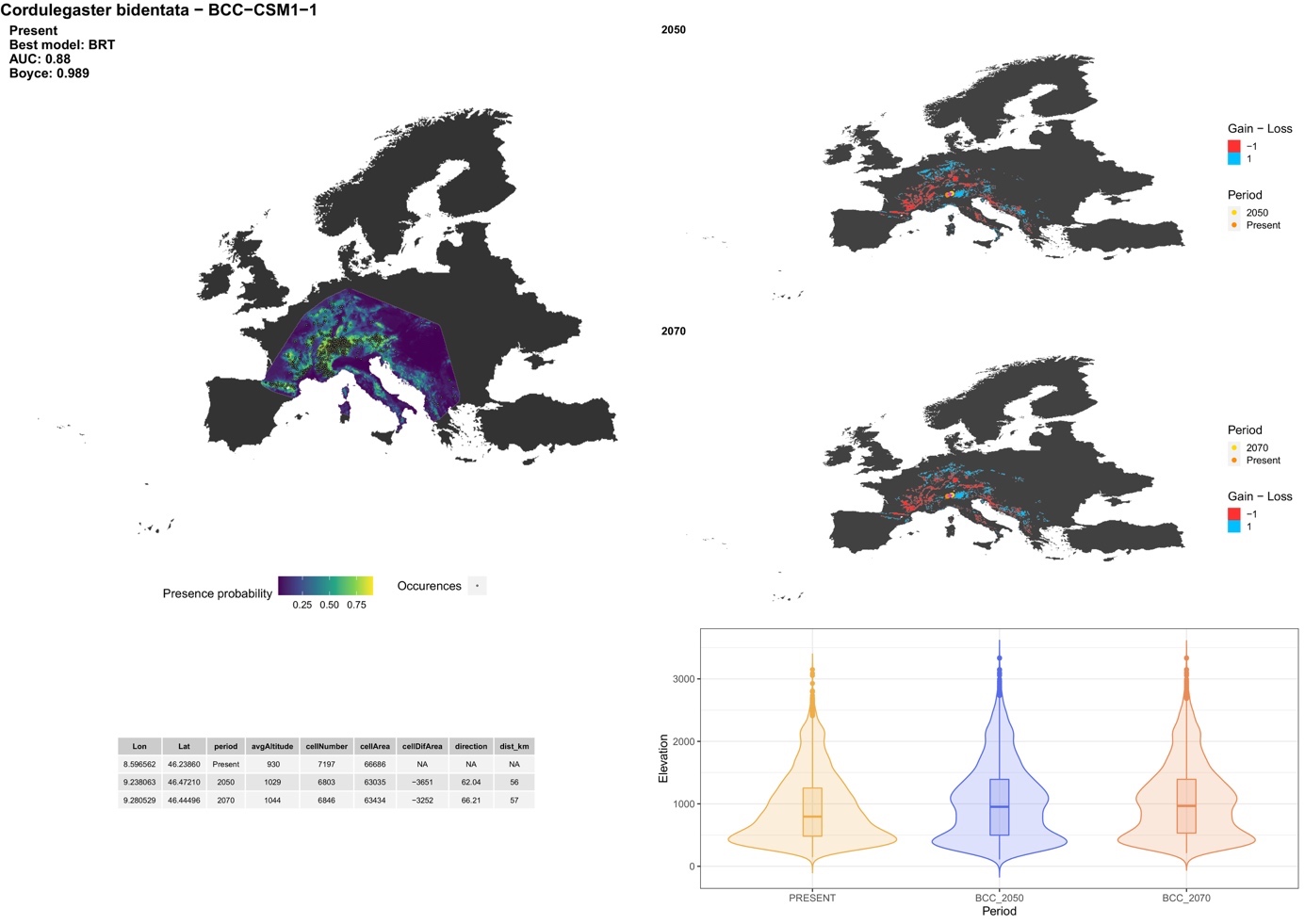

##### *Cordulegaster boltonii* (Donovan, 1807)

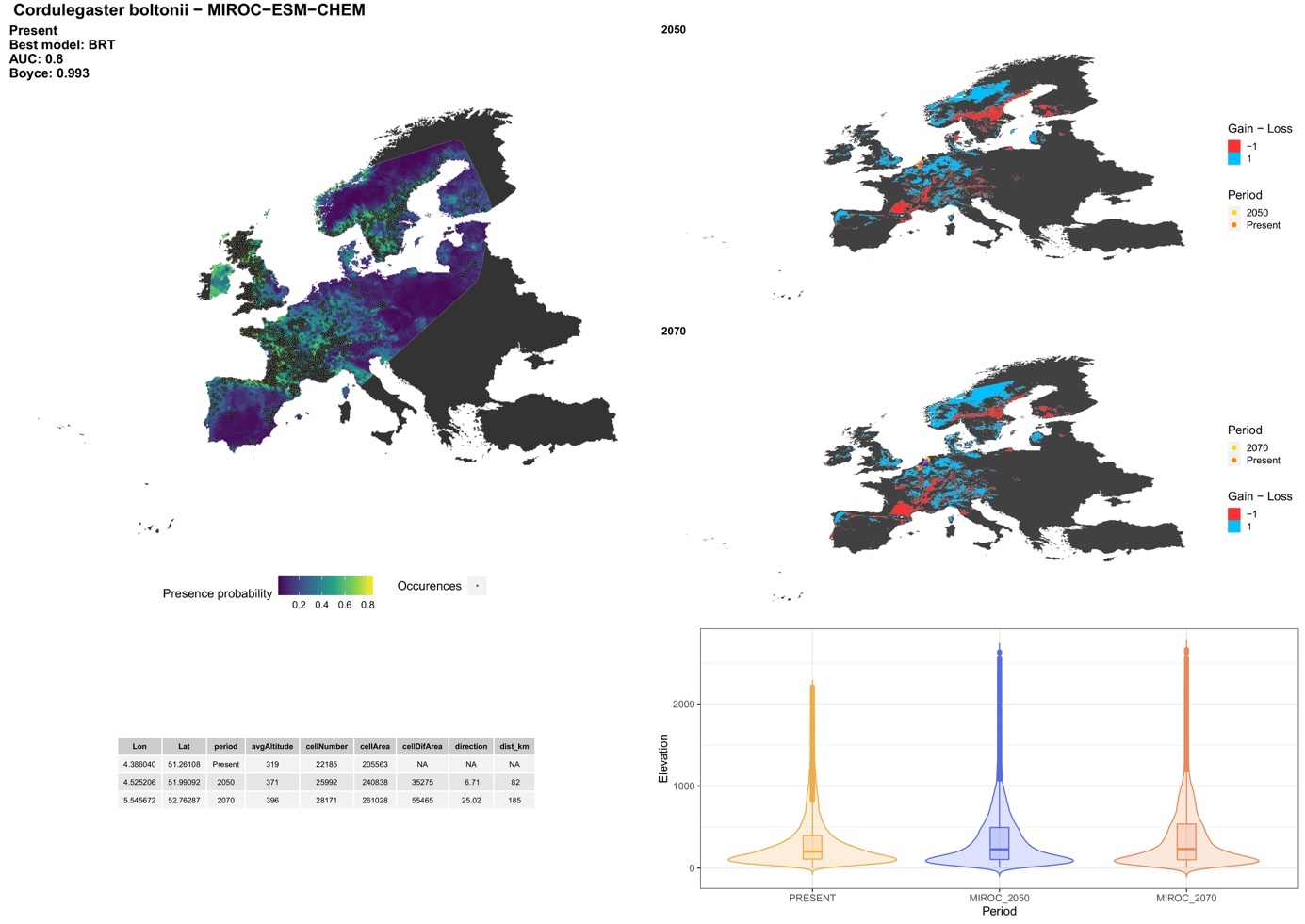

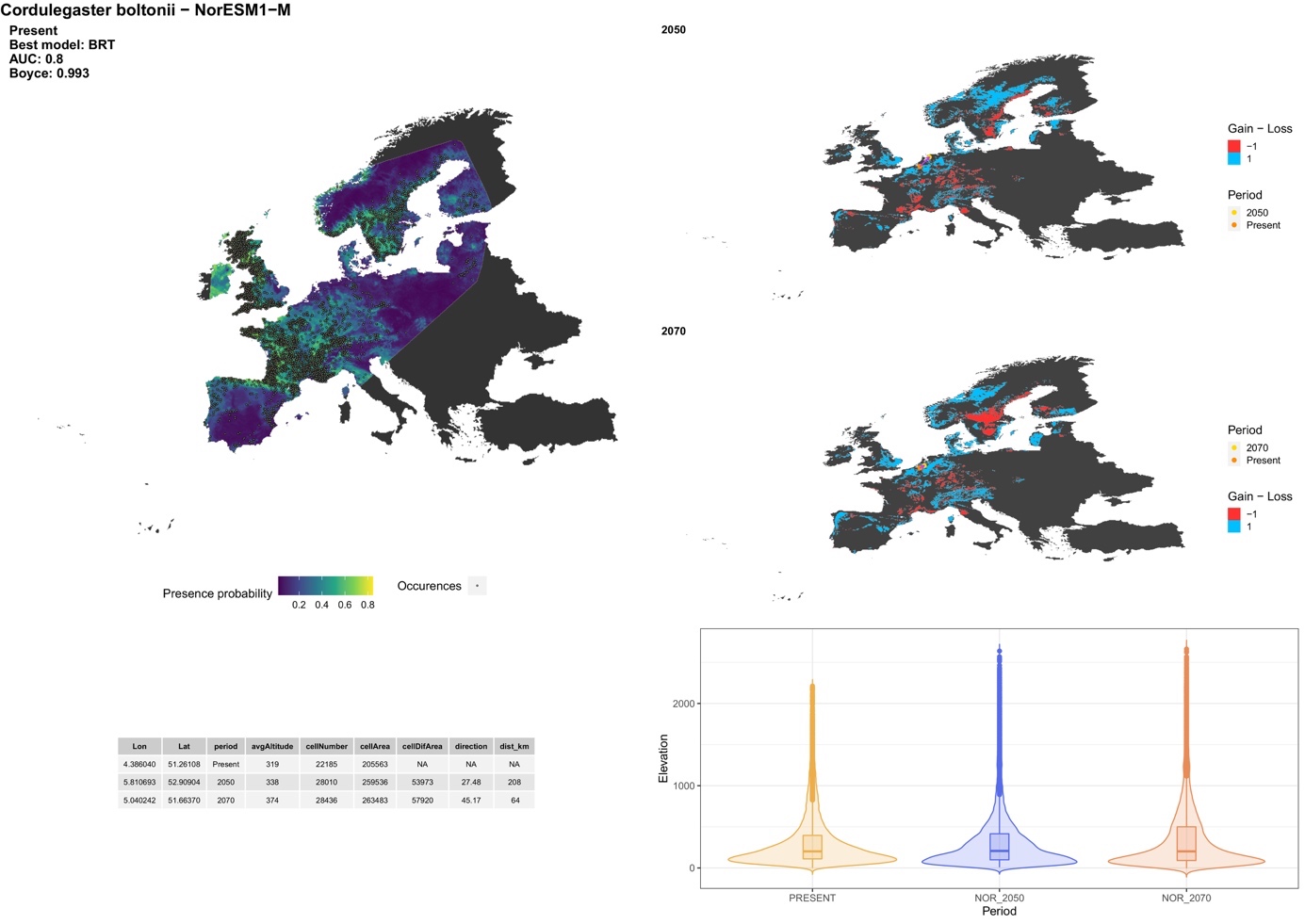

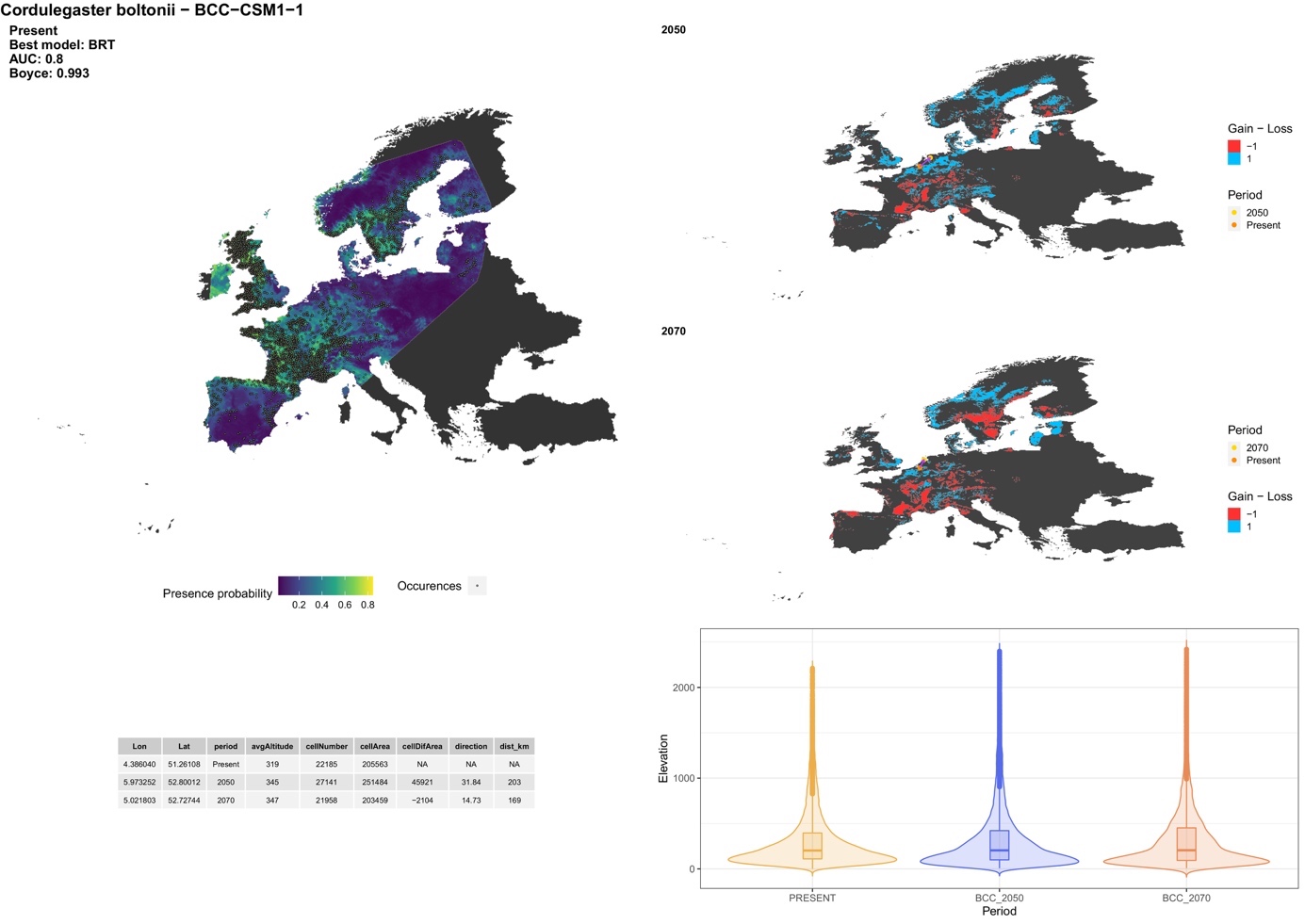

##### **Family: Macromiidae**

##### *Macromia splendens* (Pictet, 1843)

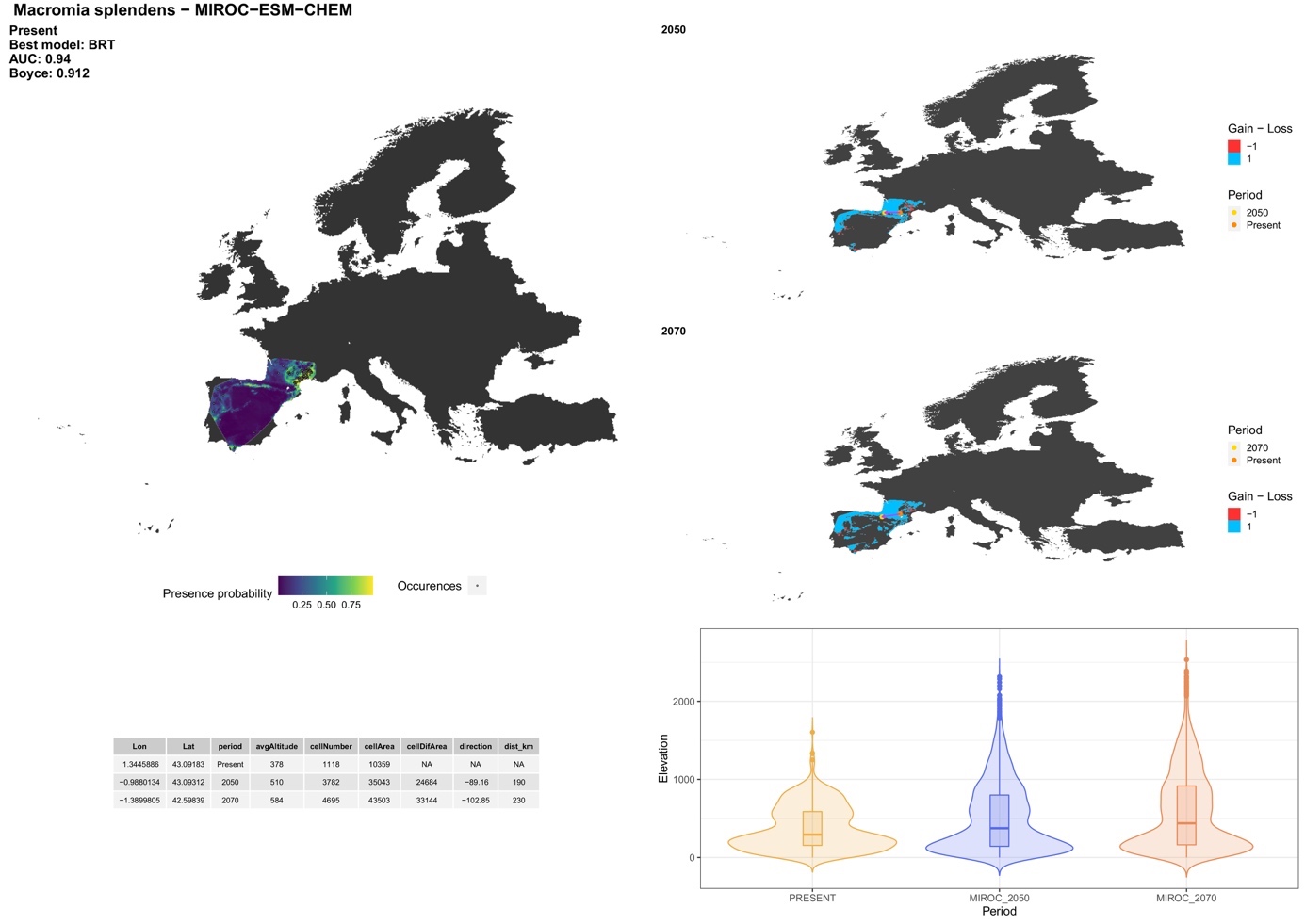

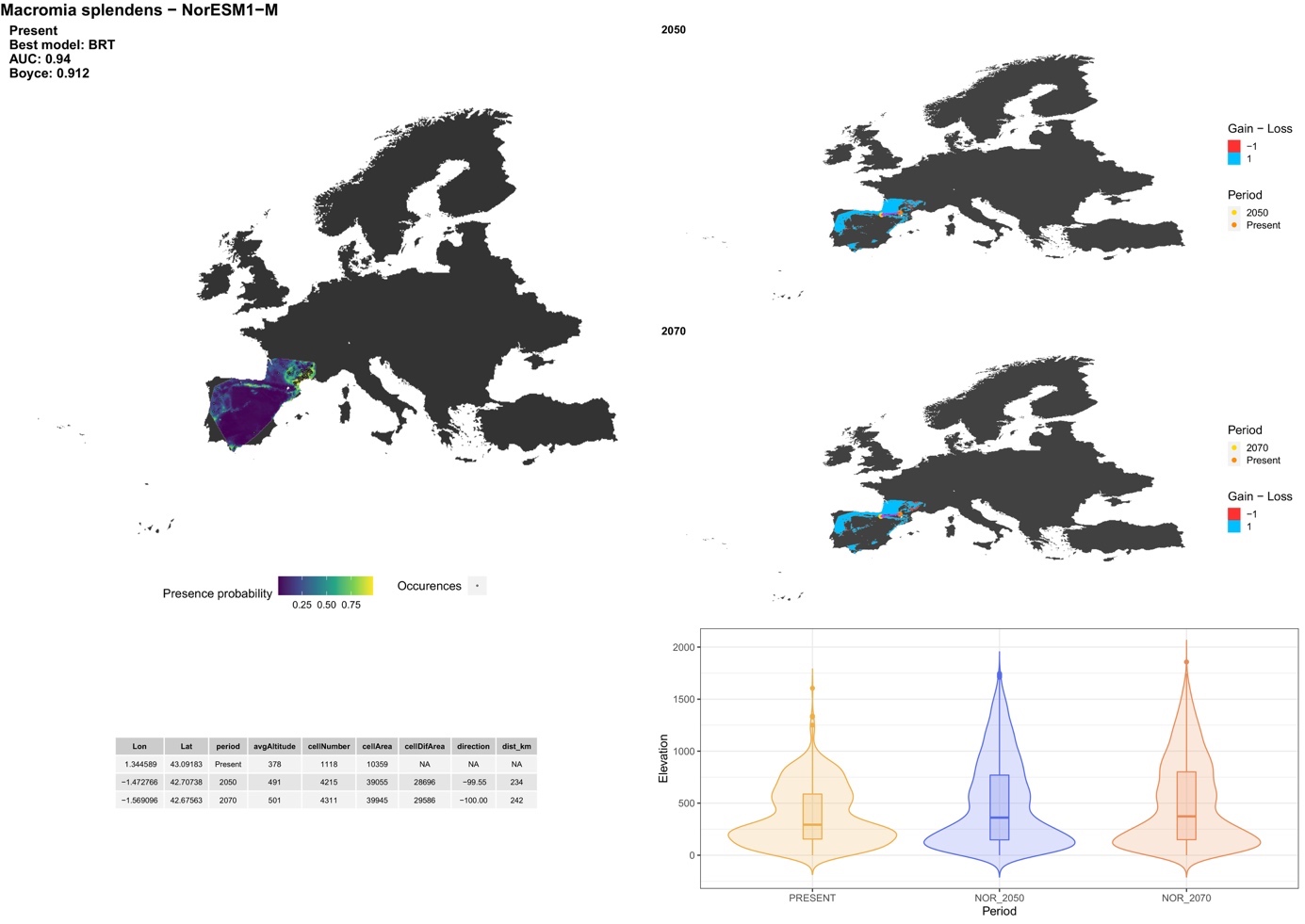

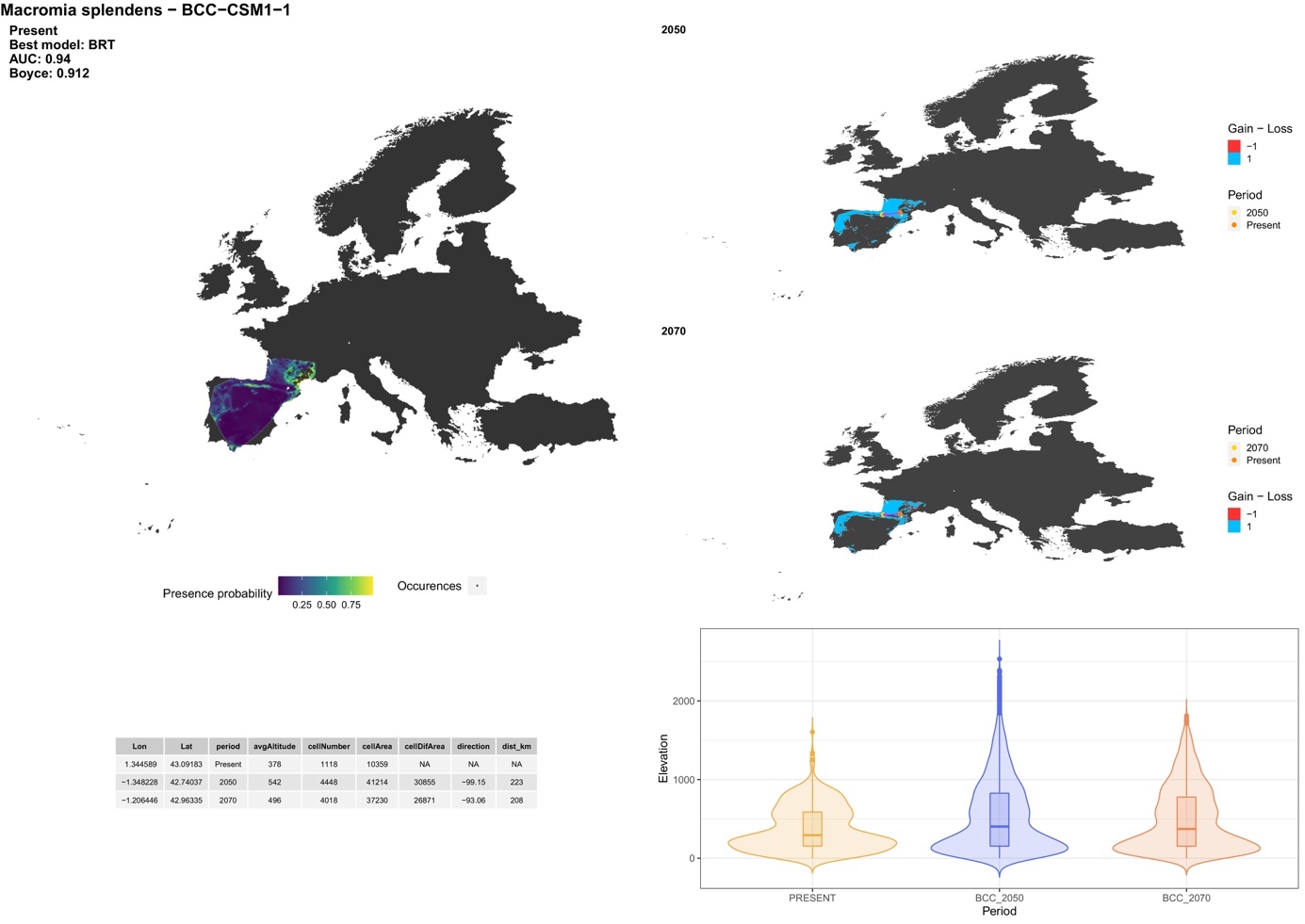

##### **Family: Corduliidae**

##### *Cordulia aenea* (Linnaeus, 1758)

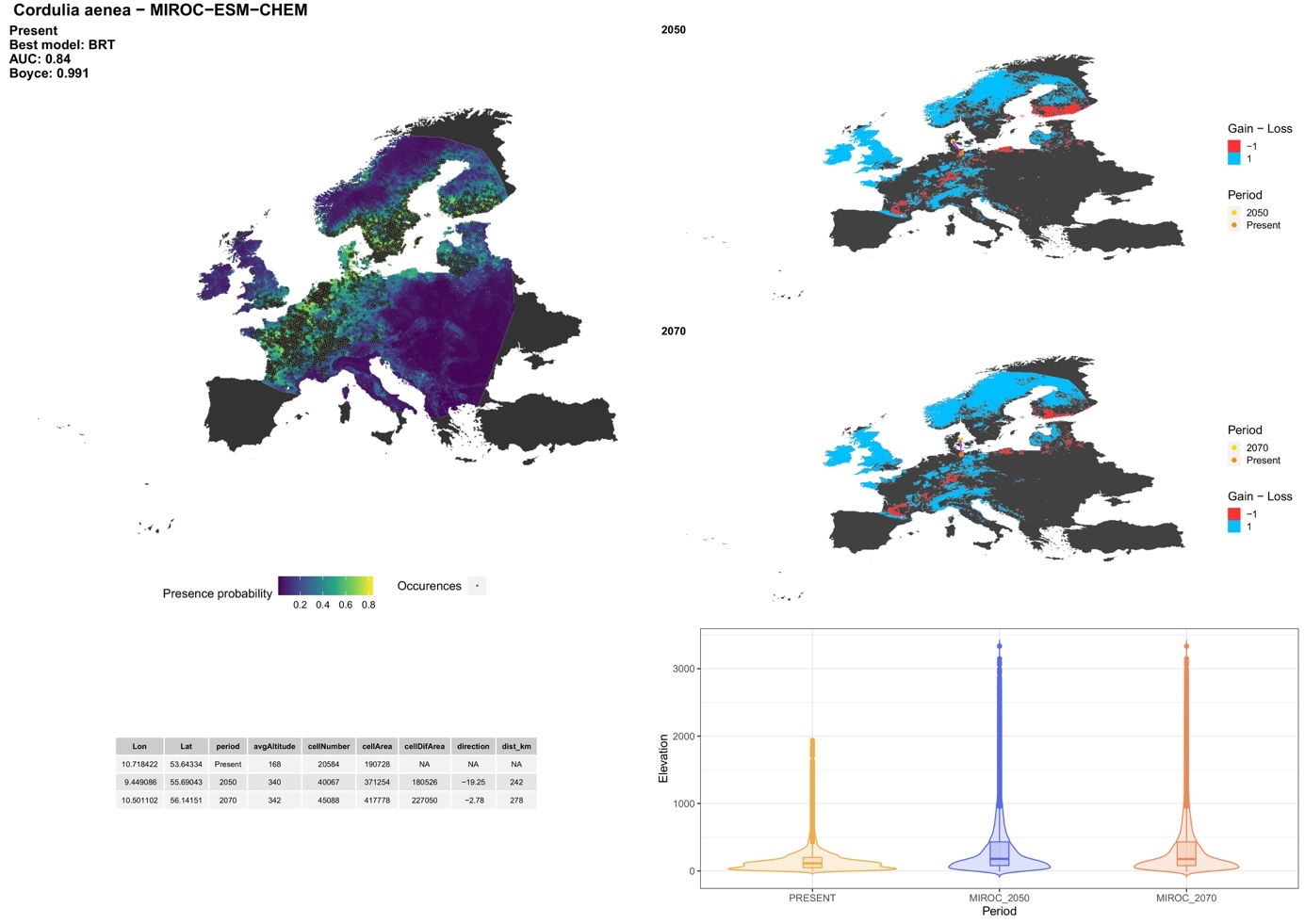

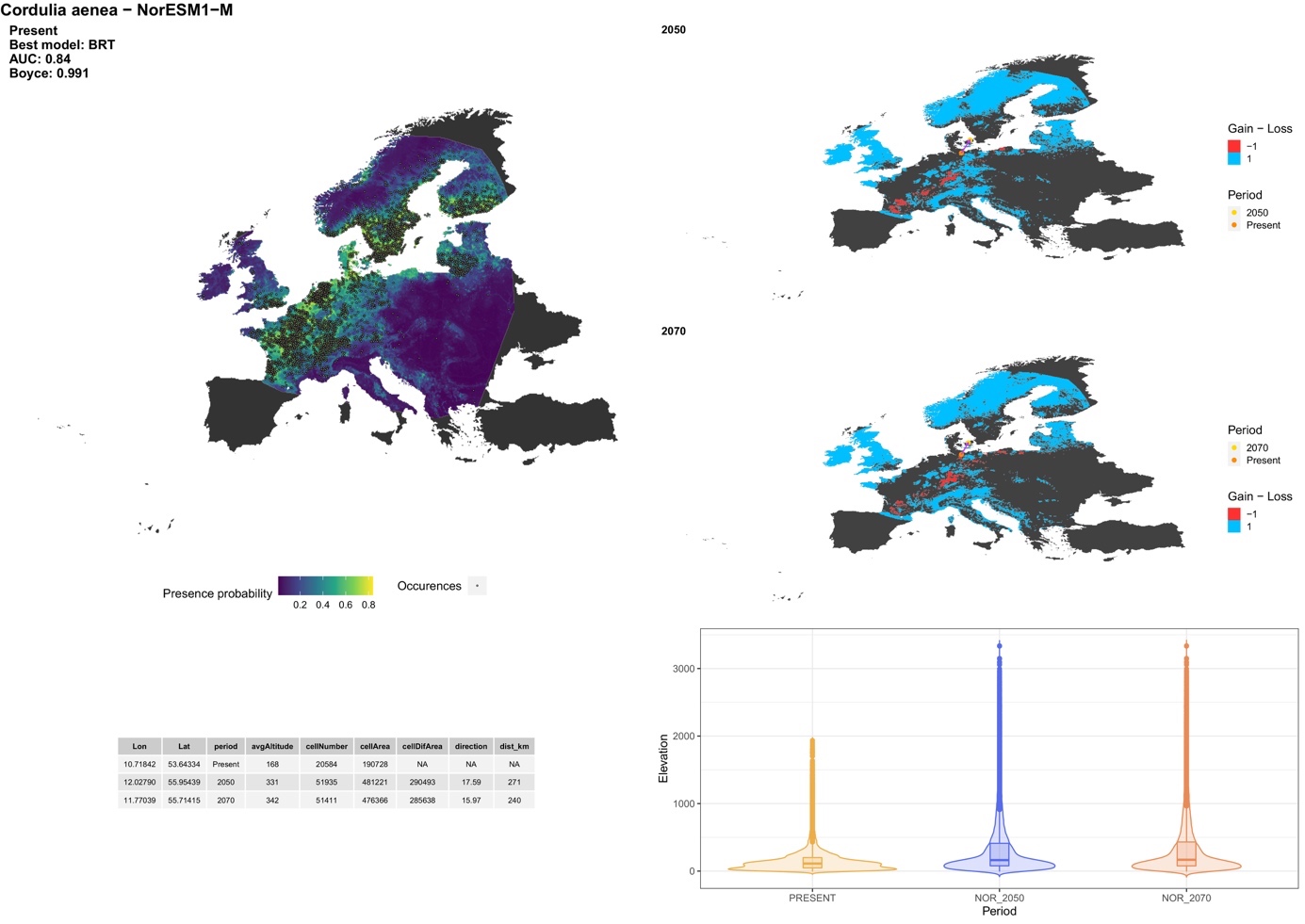

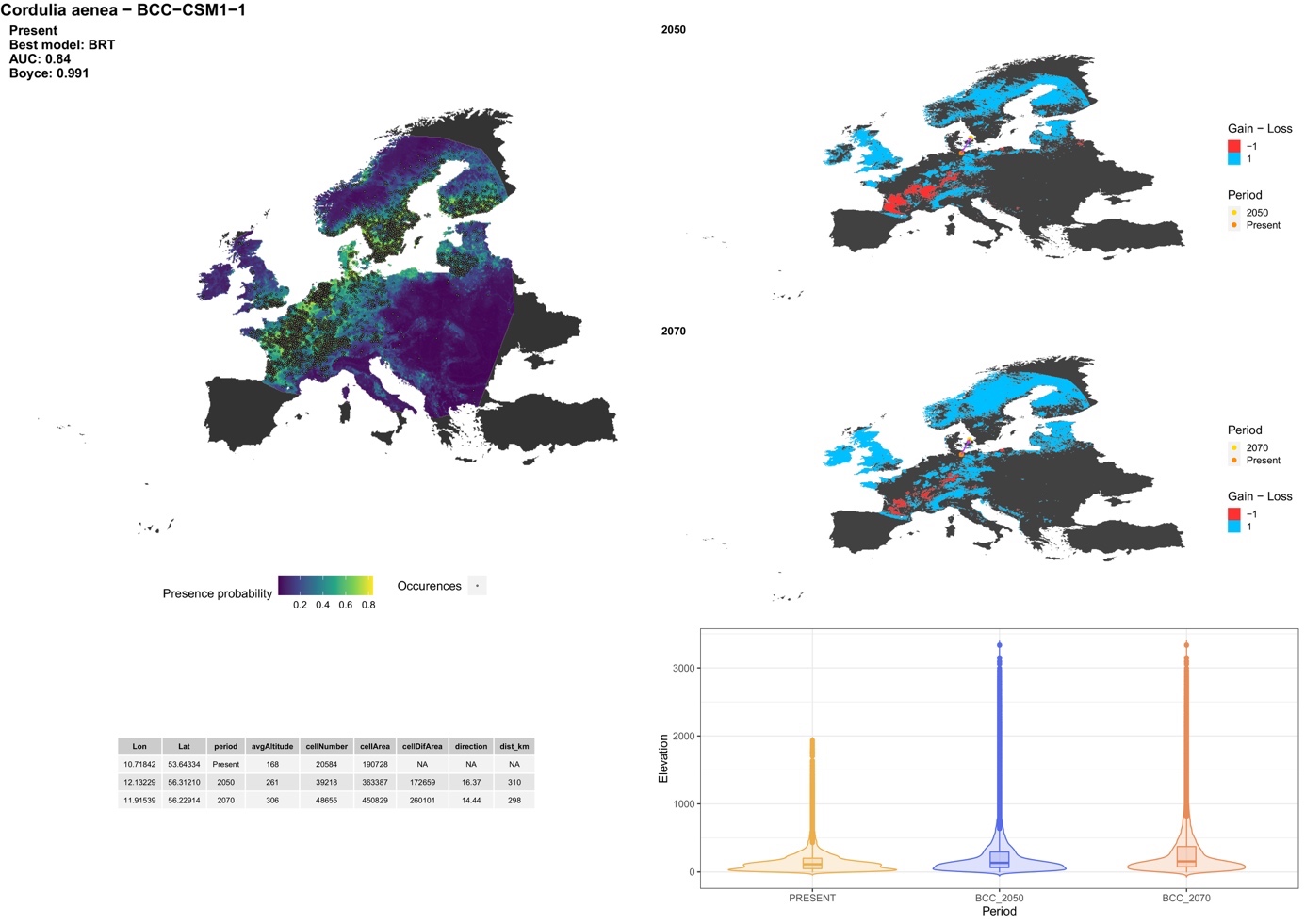

##### *Epitheca bimaculate* (Charpentier, 1825)

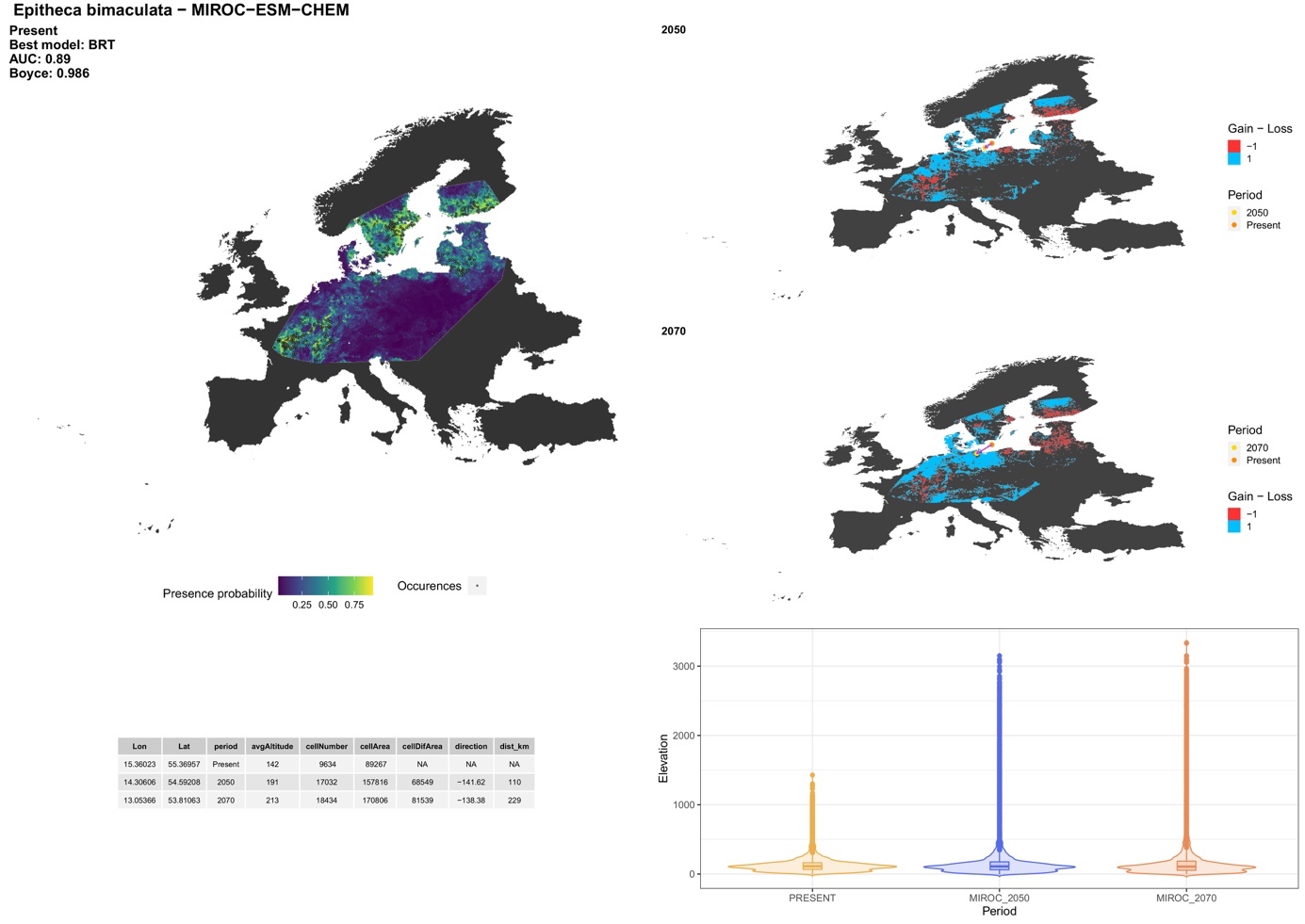

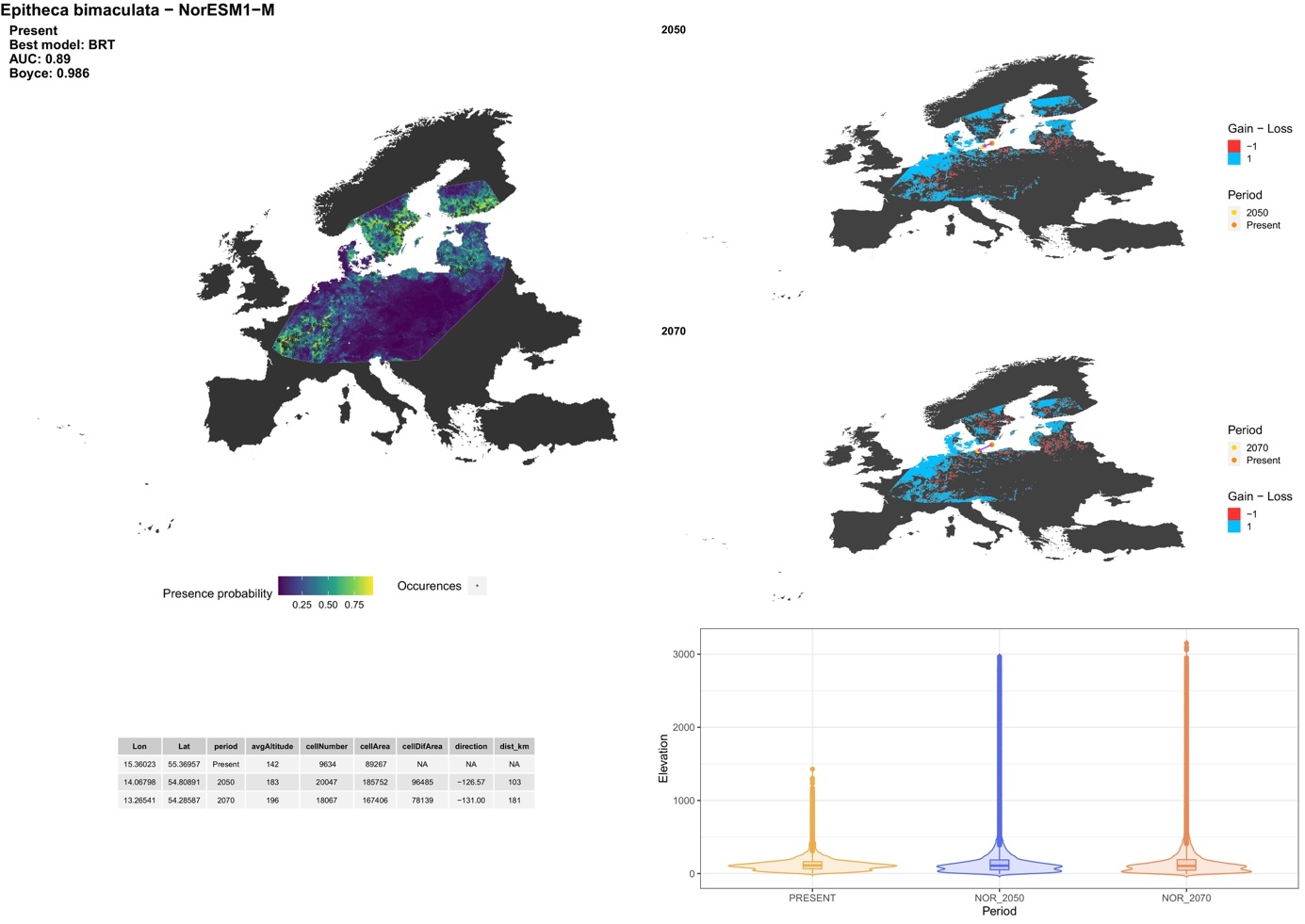

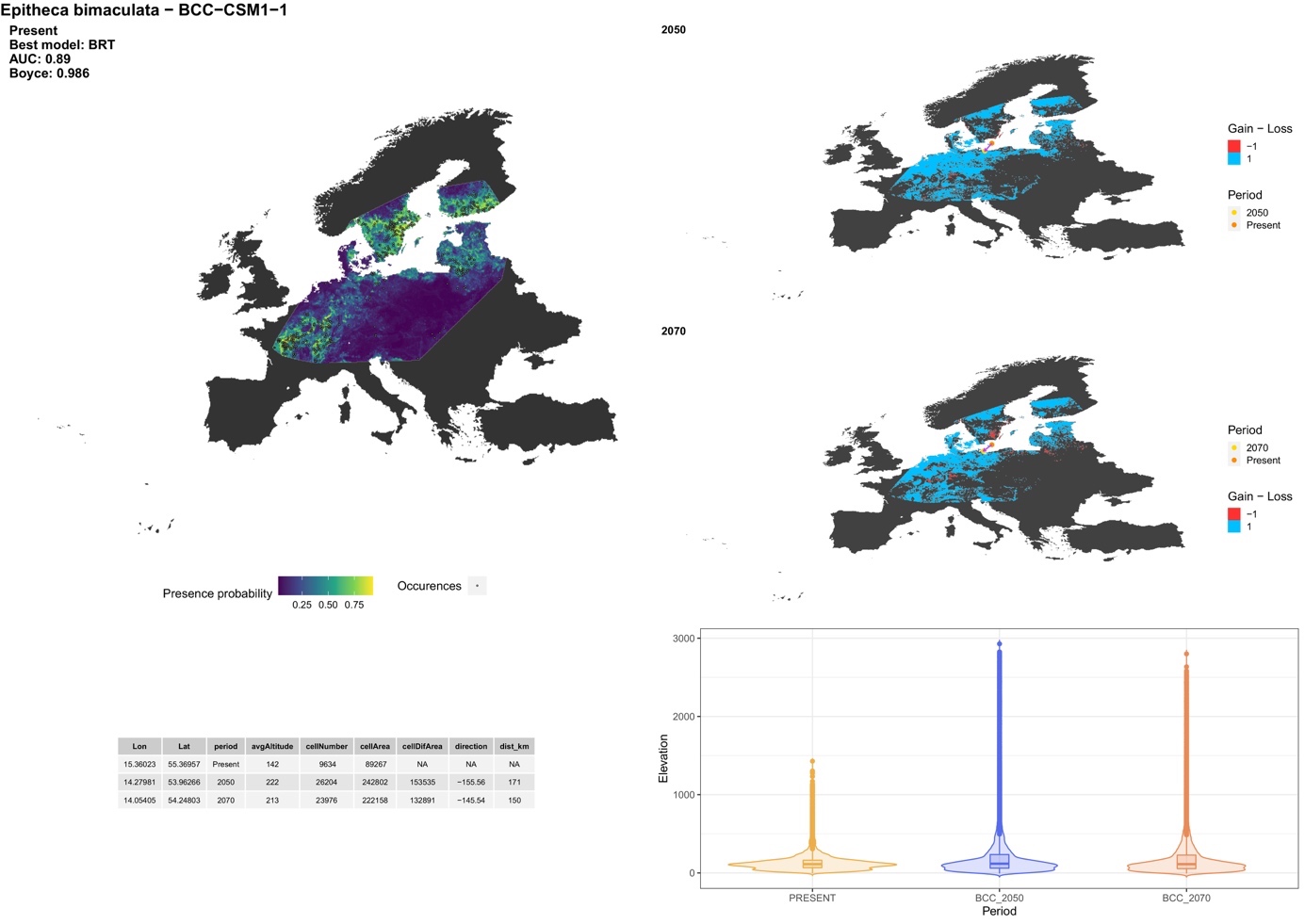

##### *Somatochlora alpestris* (Selys, 1840)

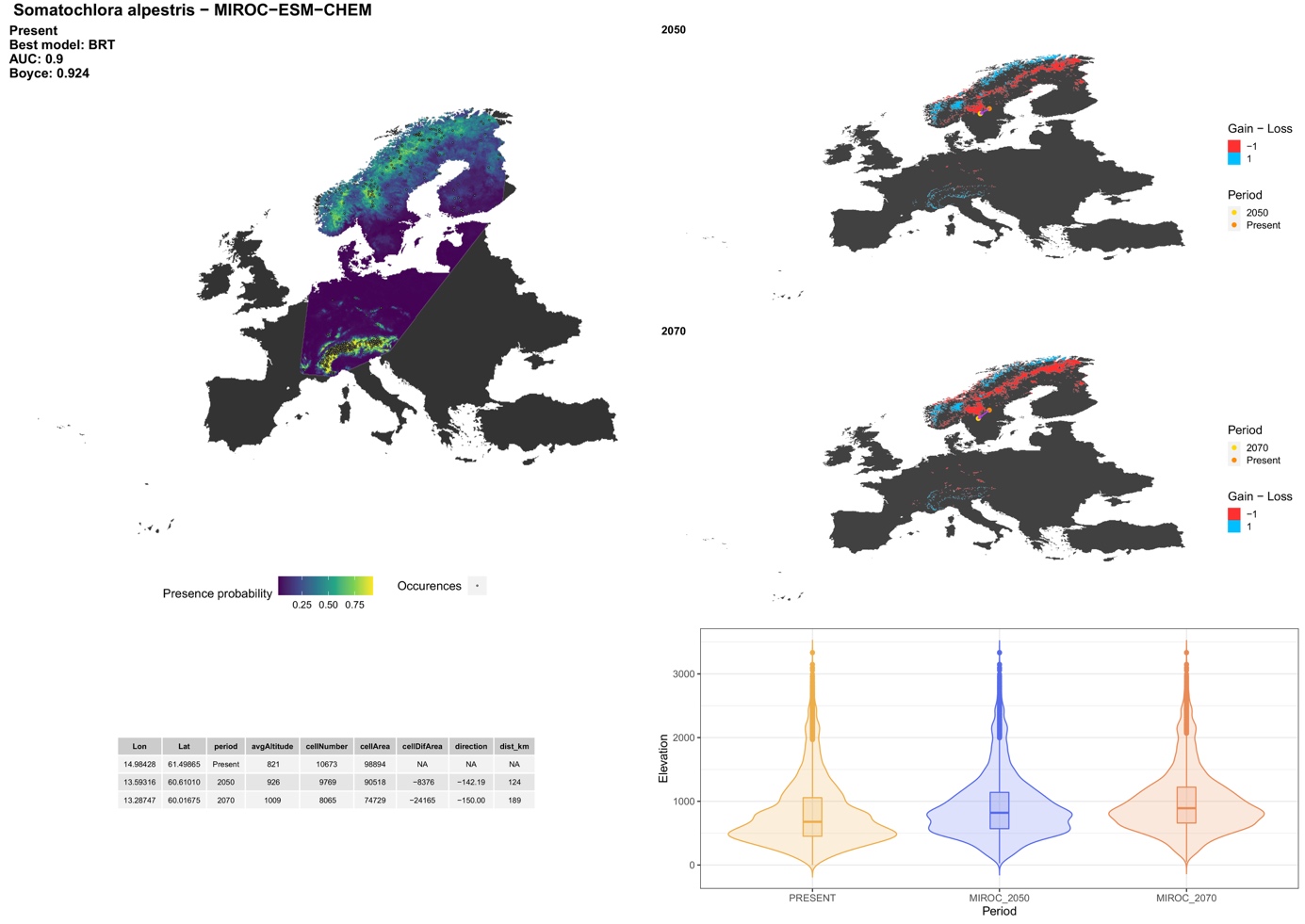

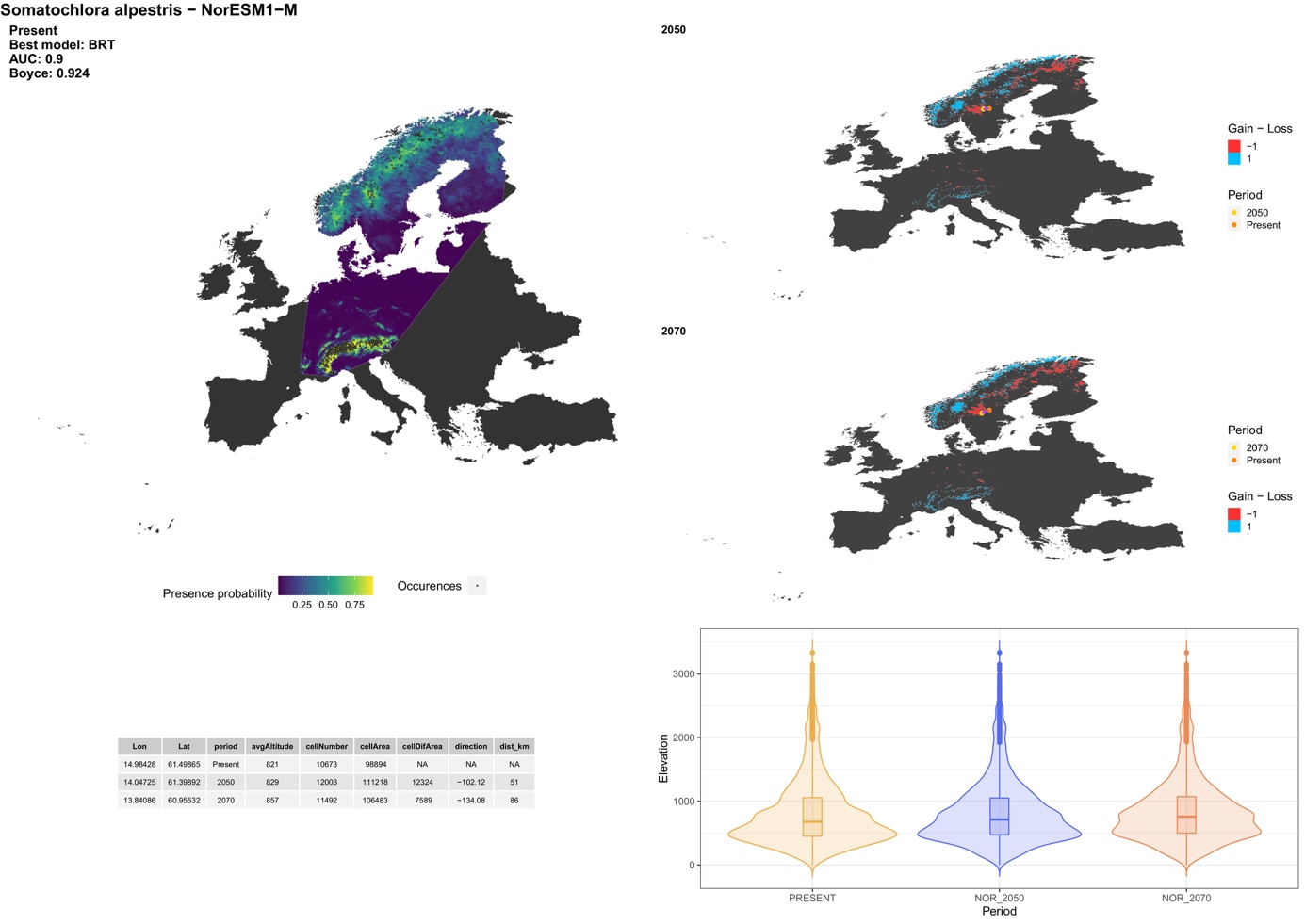

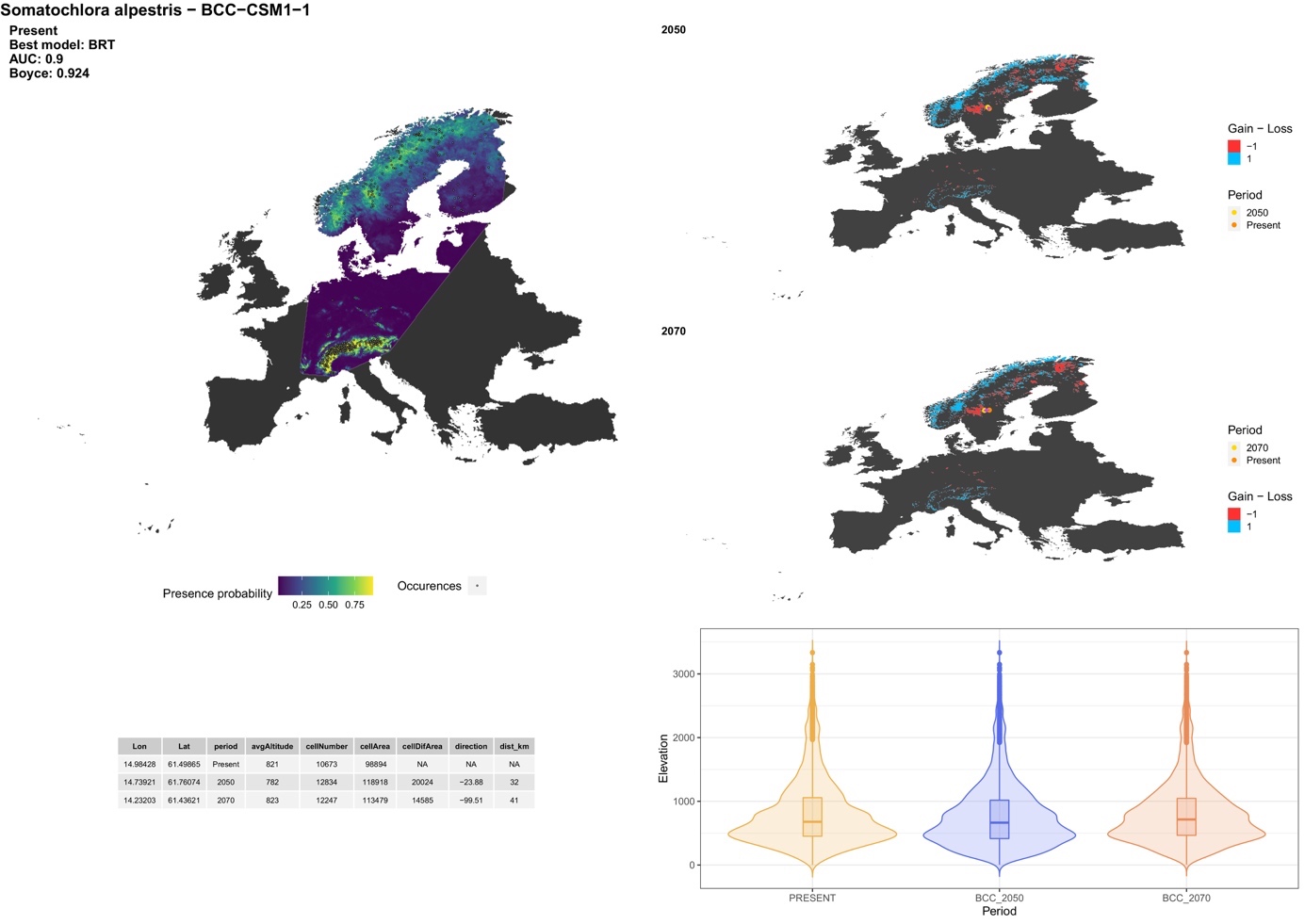

##### *Somatochlora arctica* (Zetterstedt, 1840)

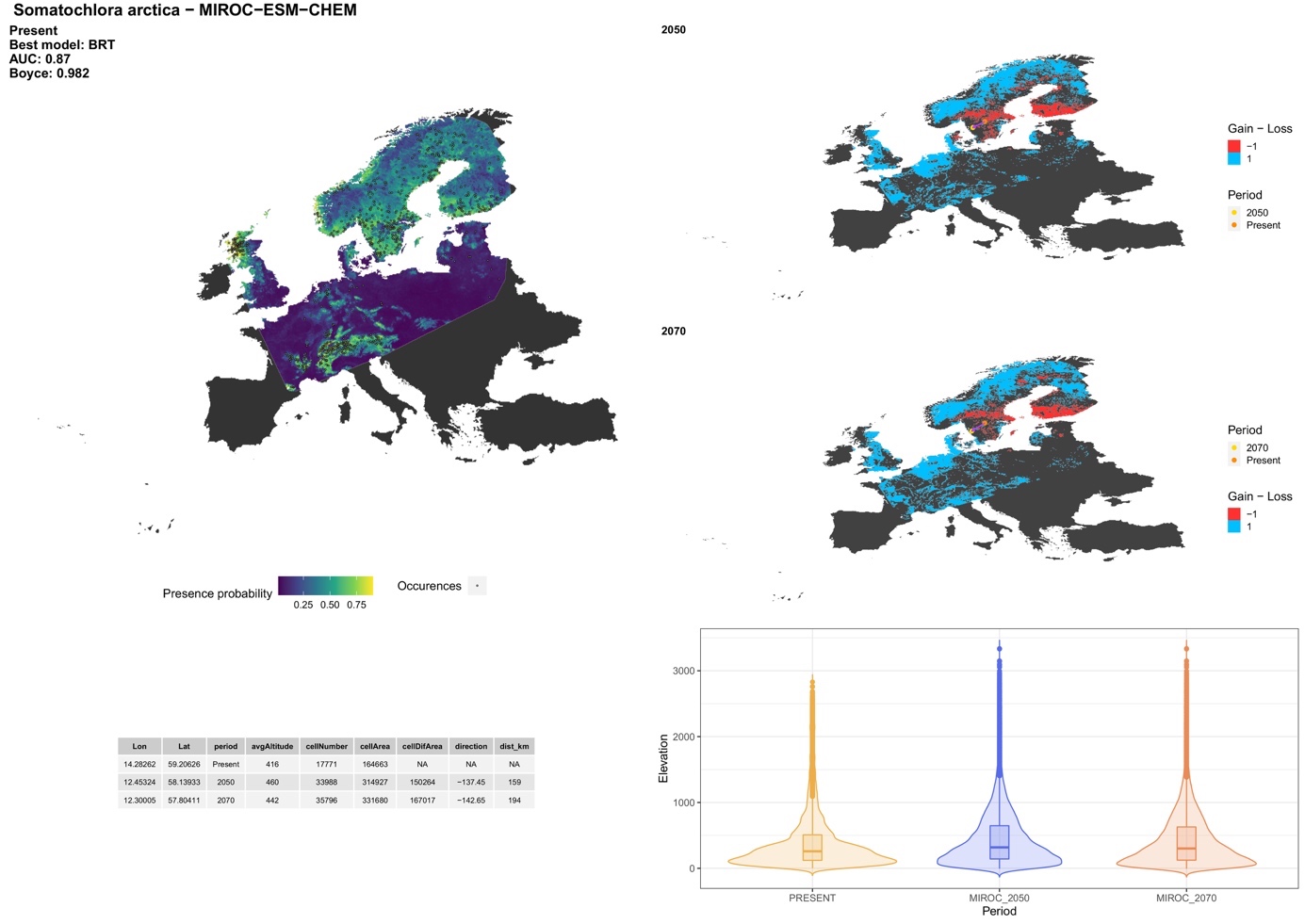

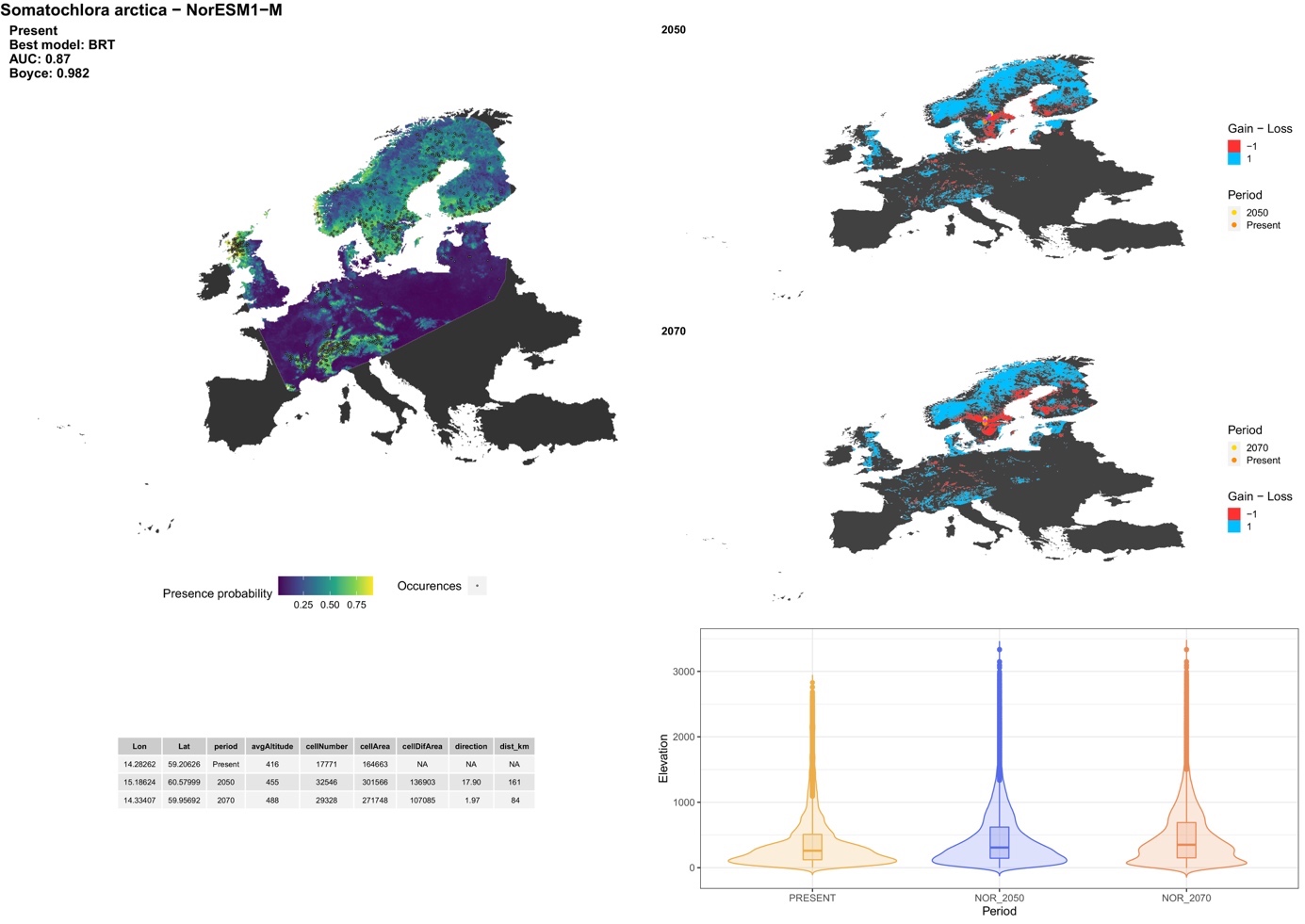

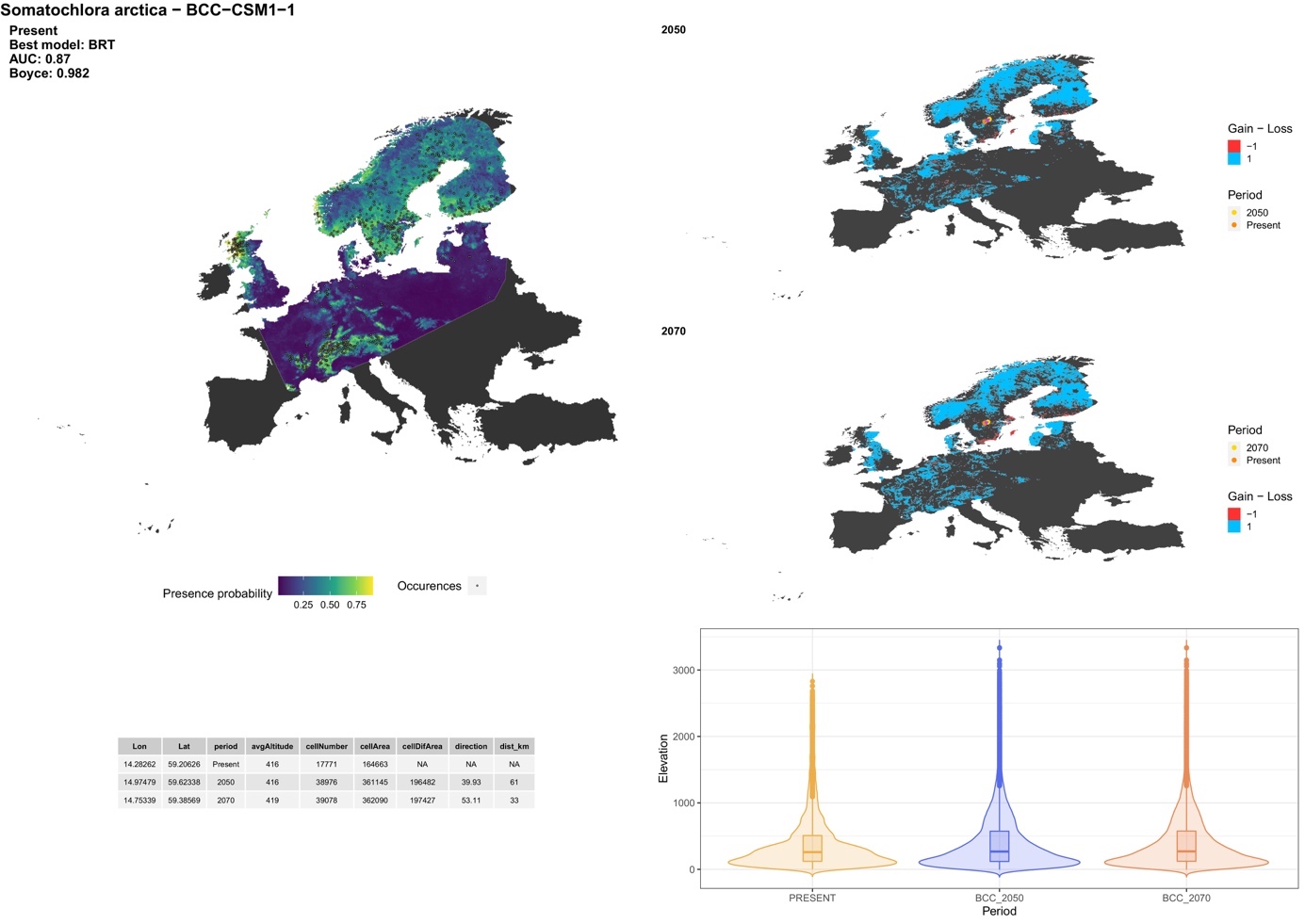

##### *Somatochlora flavomaculata* (Vander Linden, 1825)

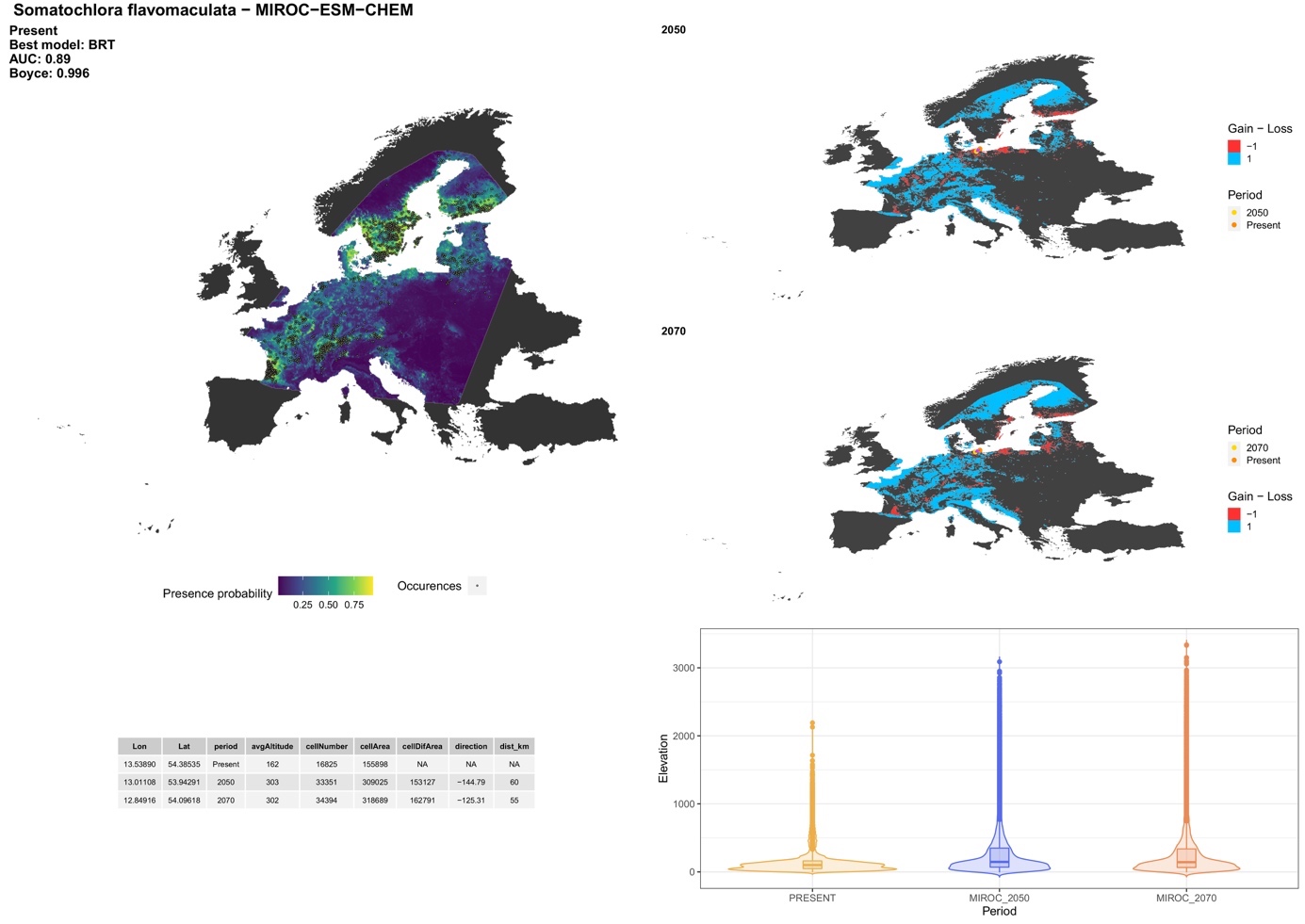

##### *Somatochlora meridionalis* Nielsen, 1935

##### *Somatochlora metallica* (Vander Linden, 1825)

##### **Family: Libellulidae**

##### *Brachythemis impartita* (Karsch, 1890)

##### *Crocothemis erythraea* (Brullé, 1832)

##### *Diplacodes lefebvrii* Rambur, 1842

##### *Leucorrhinia albifrons* (Burmeister, 1839)

##### *Leucorrhinia caudalis* (Charpentier, 1840)

##### *Leucorrhinia dubia* (Vander Linden, 1825)

##### *Leucorrhinia pectoralis* (Charpentier, 1825)

##### *Leucorrhinia rubicunda* (Linnaeus, 1758)

##### *Libellula depressa* Linnaeus, 1758

##### *Libellula fulva* Müller, 1764

##### *Libellula quadrimaculata* Linnaeus, 1758

##### *Orthetrum albistylum* (Selys, 1848)

##### *Orthetrum brunneum* (Fonscolombe, 1837)

##### *Orthetrum cancellatum* (Linnaeus, 1758)

##### *Orthetrum chrysostigma* (Burmeister, 1839)

##### *Orthetrum coerulescens* (Fabricius, 1798)

##### *Orthetrum taeniolatum* (Schneider, 1845)

##### *Orthetrum trinacria* (Selys, 1841)

##### *Selysiothemis nigra* (Vander Linden, 1825)

##### *Sympetrum danae* (Sulzer, 1776)

##### *Sympetrum depressiusculum* (Selys, 1841)

##### *Sympetrum flaveolum* (Linnaeus, 1758)

##### *Sympetrum fonscolombii* (Selys, 1840)

##### *Sympetrum meridionale* (Selys, 1841)

##### *Sympetrum pedemontanum* (Müller in Allioni, 1766)

##### *Sympetrum sanguineum* (Müller, 1764)

##### *Sympetrum sinaiticum* Dumont, 1977

##### *Sympetrum striolatum* (Charpentier, 1840)

##### *Sympetrum vulgatum* (Linnaeus, 1758)

##### *Trithemis annulata* (Palisot de Beauvois, 1807)

##### *Trithemis arteriosa* (Burmeister, 1839)

##### *Trithemis kirbyi* Selys, 1891

### **Zygoptera**

##### **Family: Calopterygidae**

##### *Calopteryx haemorrhoidalis* (Vander Linden, 1825)

##### *Calopteryx splendens* (Harris, 1780)

##### *Calopteryx virgo* (Linnaeus, 1758)

##### *Calopteryx xanthostoma* (Charpentier, 1825)

##### **Family: Coenagrionidae**

##### *Ceriagrion tenellum* (De Villers, 1789)

##### *Coenagrion armatum* (Charpentier, 1840)

##### *Coenagrion caerulescens* (Fonscolombe, 1838)

##### *Coenagrion hastulatum* (Charpentier, 1825)

##### *Coenagrion johanssoni* Wallengren, 1894

##### *Coenagrion lunulatum* (Charpentier, 1840)

##### *Coenagrion mercuriale* (Charpentier, 1840)

##### *Coenagrion ornatum* (Selys, 1850)

##### *Coenagrion puella* (Linnaeus, 1758)

##### *Coenagrion pulchellum* (Vander Linden, 1825)

##### *Coenagrion scitulum* (Rambur, 1842)

##### *Enallagma cyathigerum* (Charpentier, 1840)

##### *Erythromma lindenii* (Selys, 1840)

##### *Erythromma najas* (Hansemann, 1823)

##### *Erythromma viridulum* (Charpentier, 1840)

##### *Ischnura elegans* (Vander Linden, 1820)

##### *Ischnura genei* (Rambur, 1842)

##### *Ischnura graellsii* (Rambur, 1842)

##### *Ischnura pumilio* (Charpentier, 1825)

##### *Nehalennia speciosa* (Charpentier, 1840)

##### *Pyrrhosoma nymphula* (Sulzer, 1776)

##### **Family: Euphaeidae**

##### *Epallage fatime* (Charpentier, 1840)

##### **Family: Lestidae**

##### *Chalcolestes parvidens* Artobolevsky, 1929

##### *Chalcolestes viridis* (Vander Linden, 1825)

##### *Lestes barbarous* (Fabricius, 1798)

##### *Lestes dryas* Kirby, 1890

##### *Lestes macrostigma* (Eversmann, 1836)

##### *Lestes sponsa* (Hansemann, 1823)

##### *Lestes virens* (Charpentier, 1825)

##### *Sympecma fusca* (Vander Linden, 1820)

##### *Sympecma paedisca* (Brauer, 1877)

##### **Family: Platycnemididae**

##### *Platycnemis acutipennis* Selys, 1841

##### *Platycnemis latipes* Rambur, 1842

##### *Platycnemis pennipes* (Pallas, 1771)

##### **Family: *Incertae sedis***

##### *Oxygastra curtisii* (Dale, 1834)
