## Supplementary material for "Climate change will redefine taxonomic, functional, and phylogenetic diversity patterns of Odonata in space and time": S8_Alpha and Beta_v2

**Supplementary material S8**. Quantification of alpha diversity per different climate scenarios (BCC-CSM1-1; MIROC-ESM-CHEM; NorESM1-M) and time periods (current; 2050; 2070). For future scenarios, the cold-colour palette indicates the species loss, whereas the warm-colour palette indicates the spacies gain.

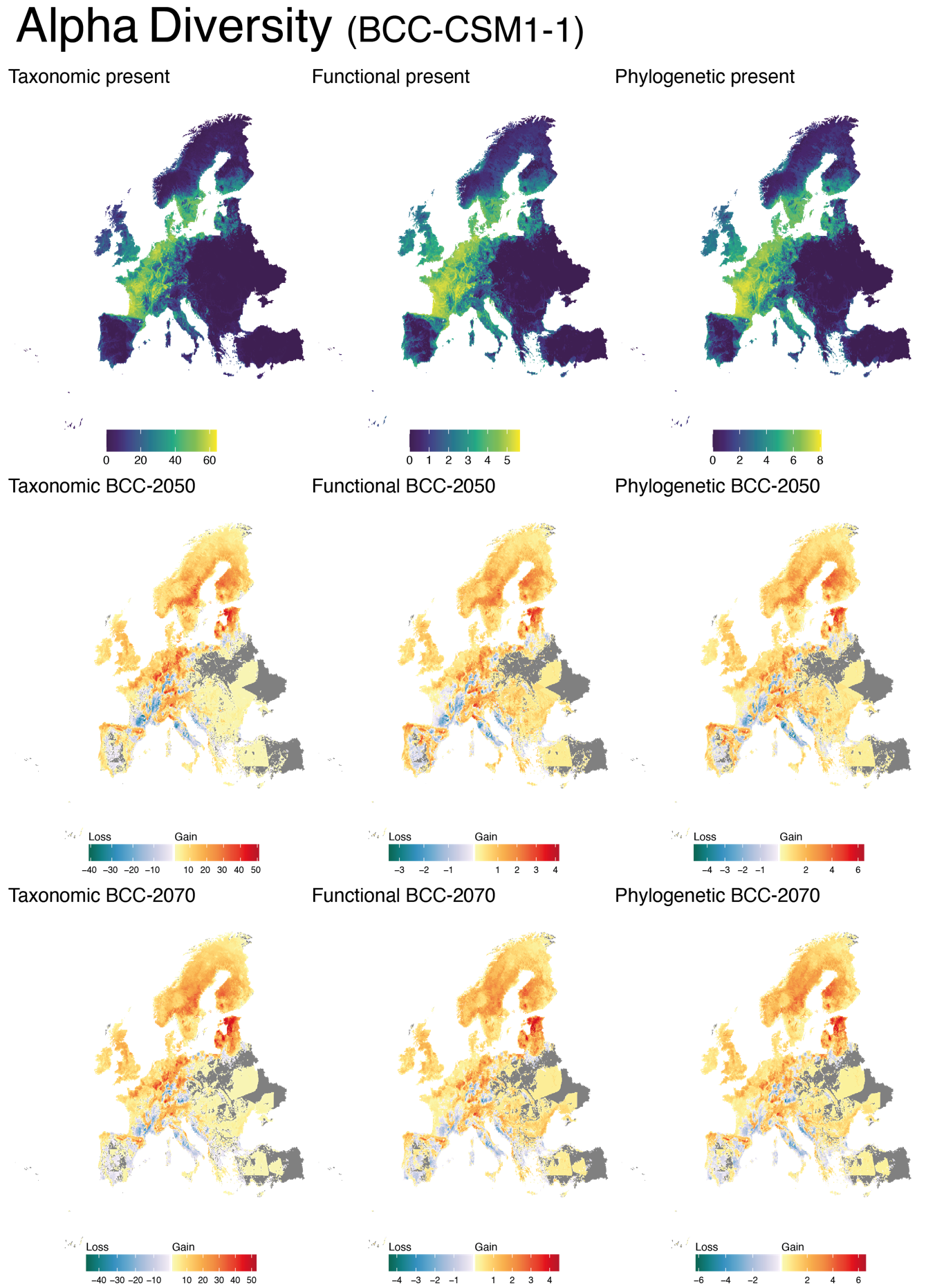

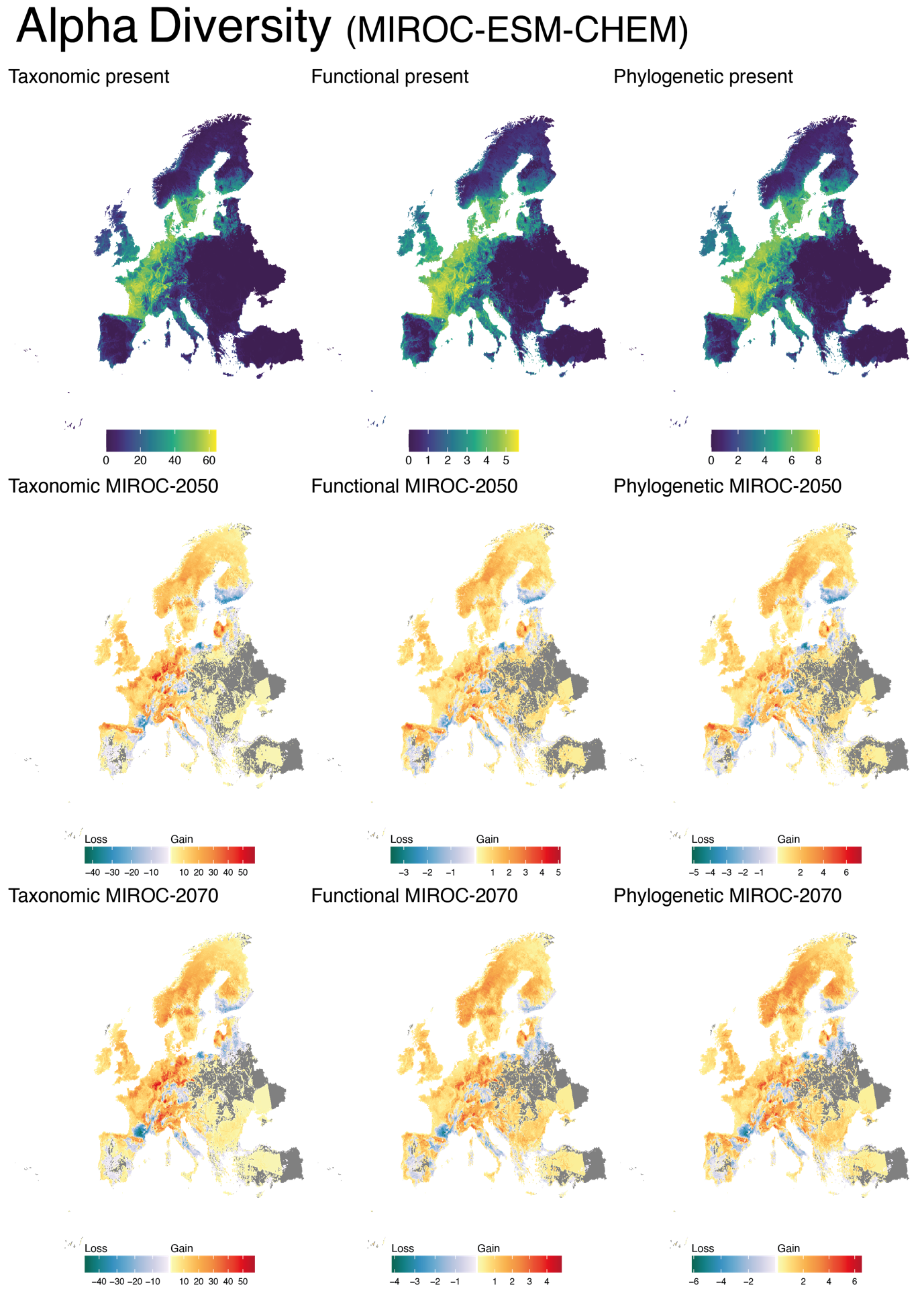

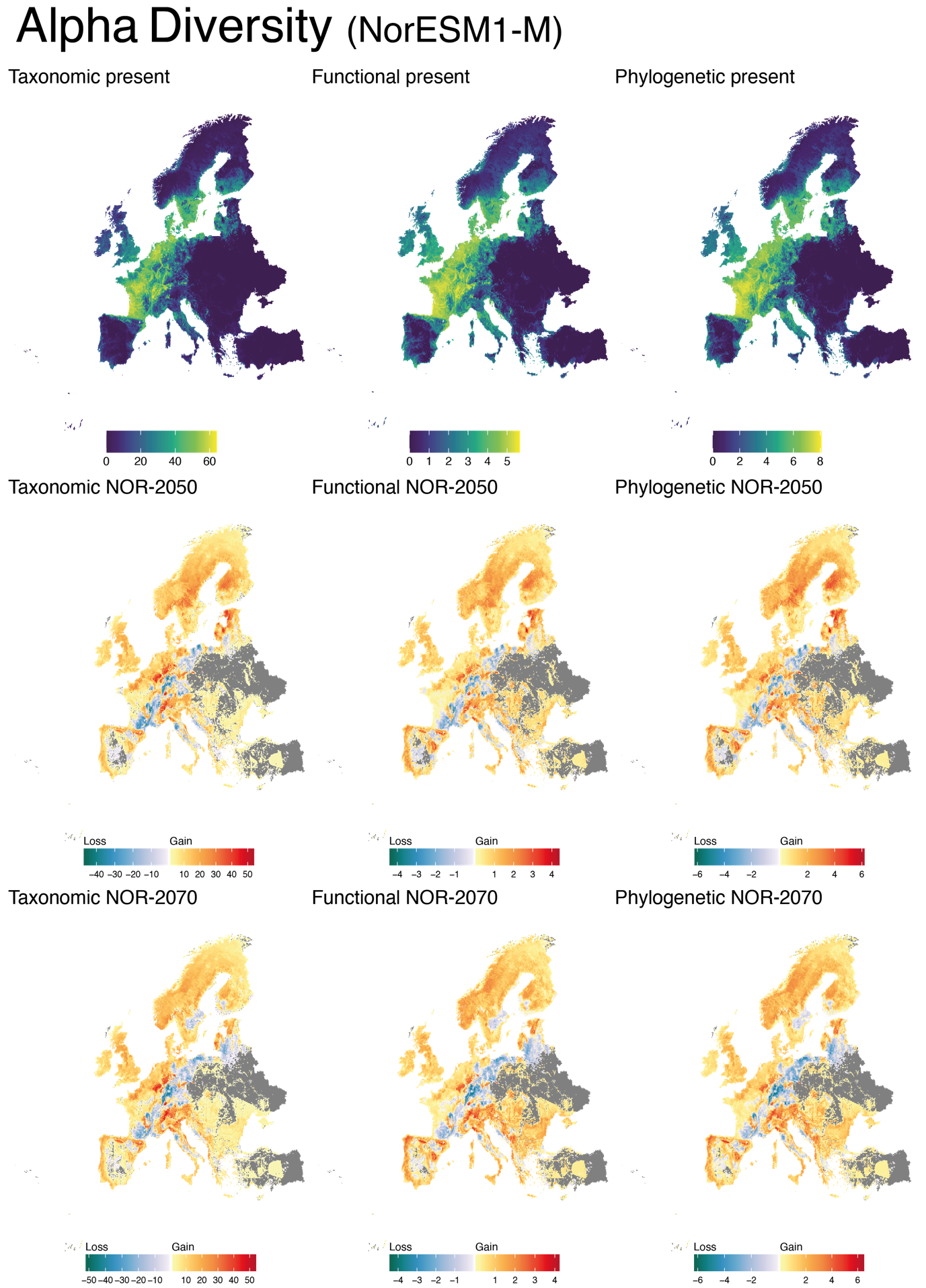

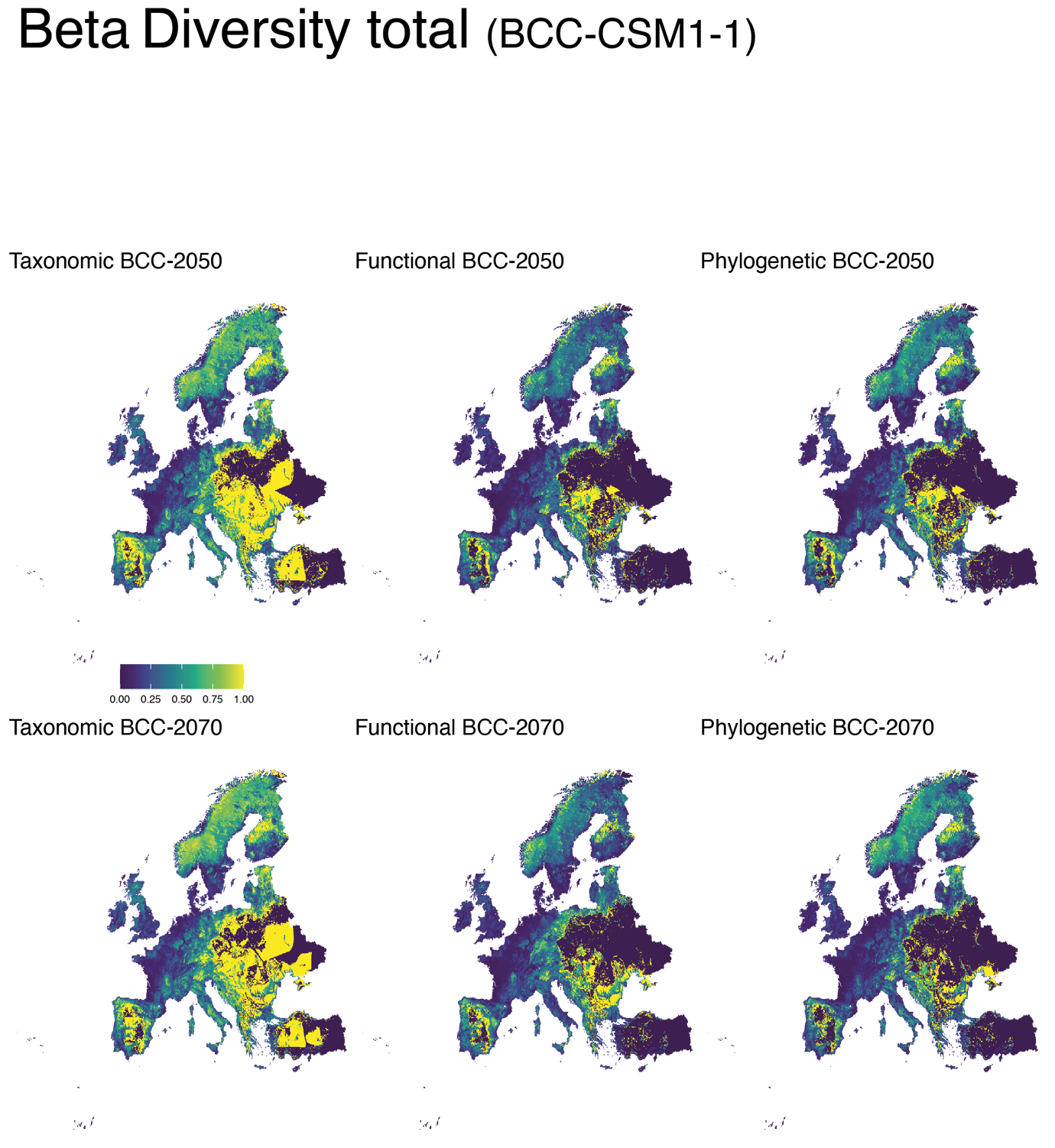

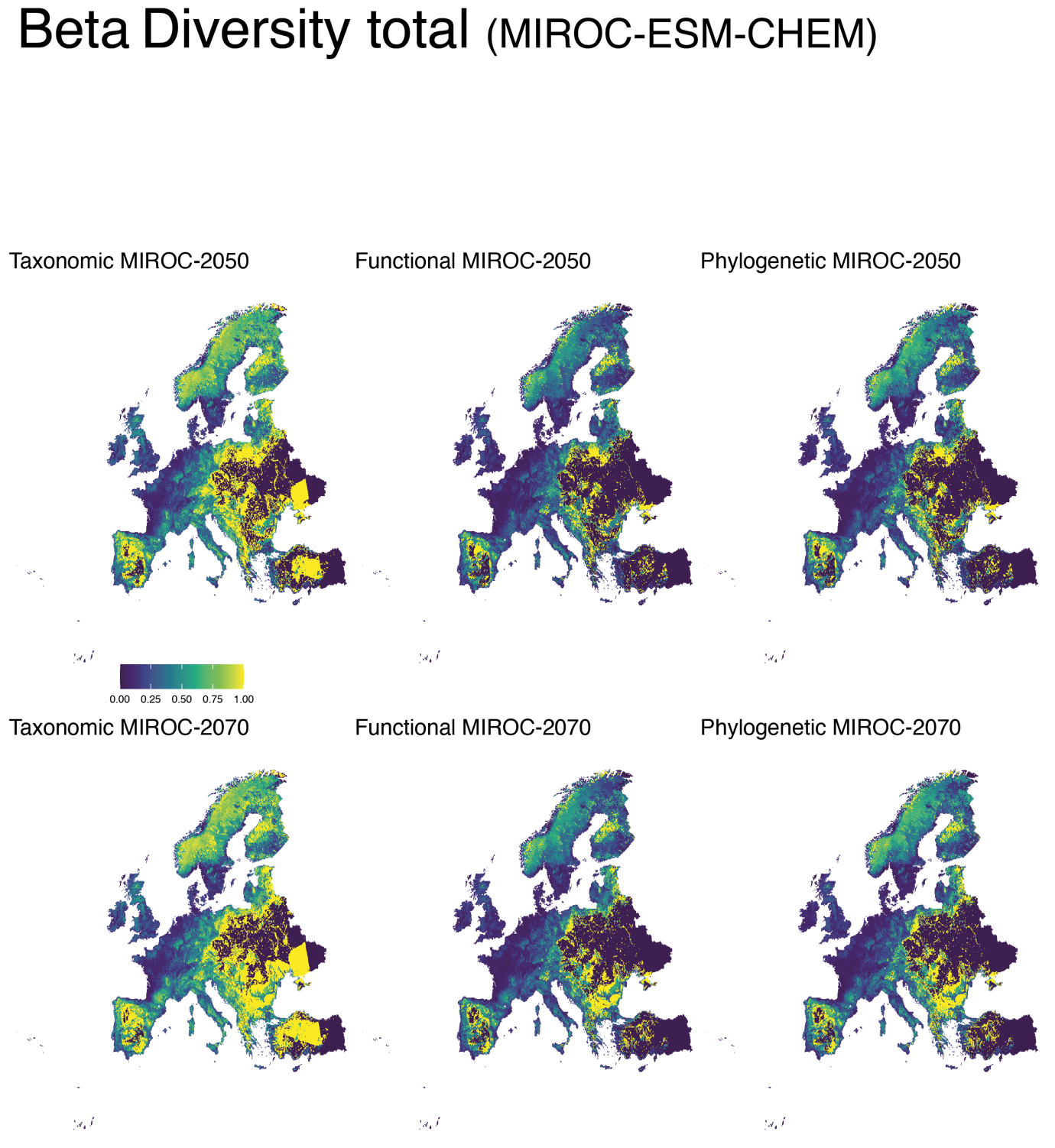

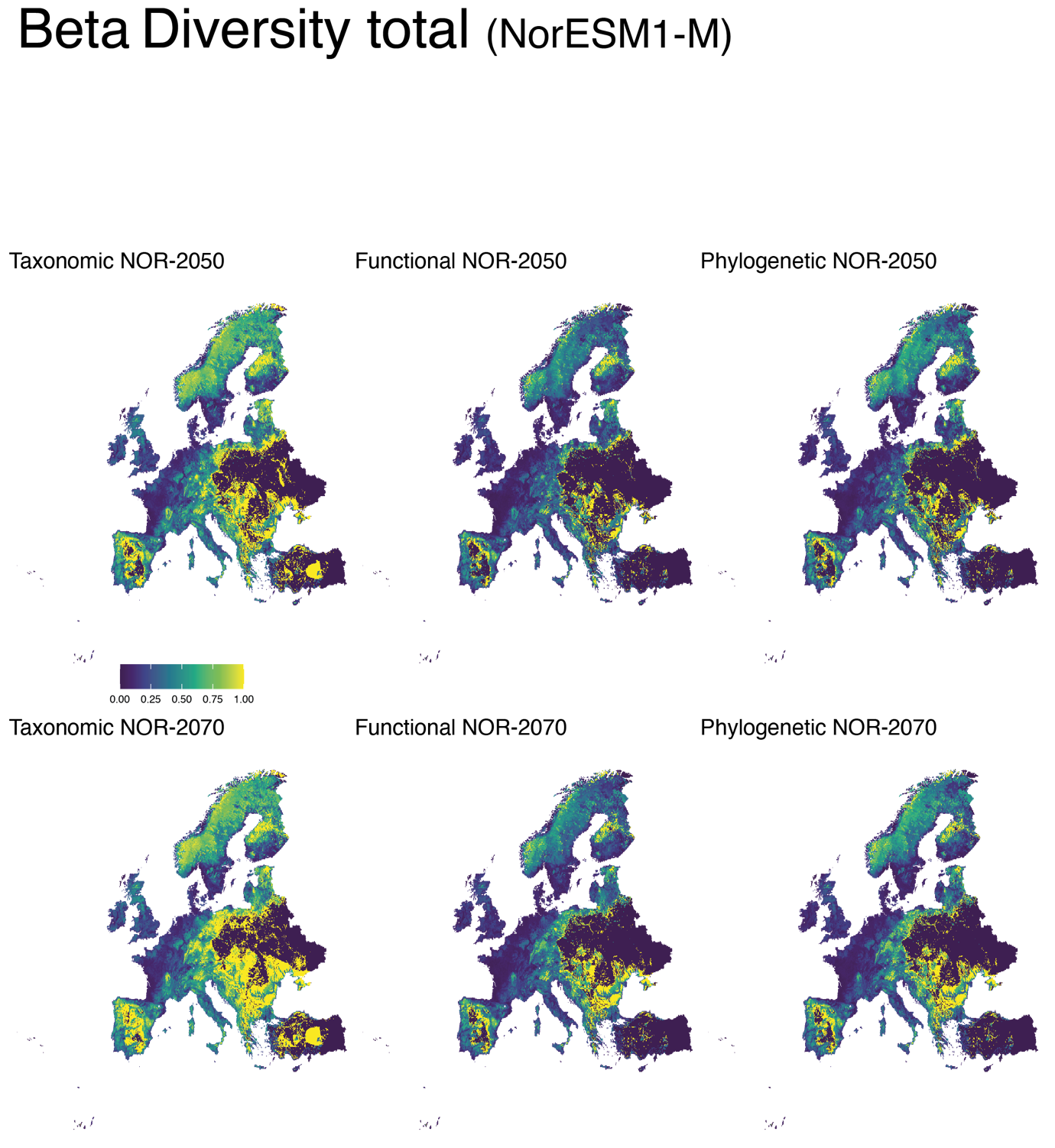

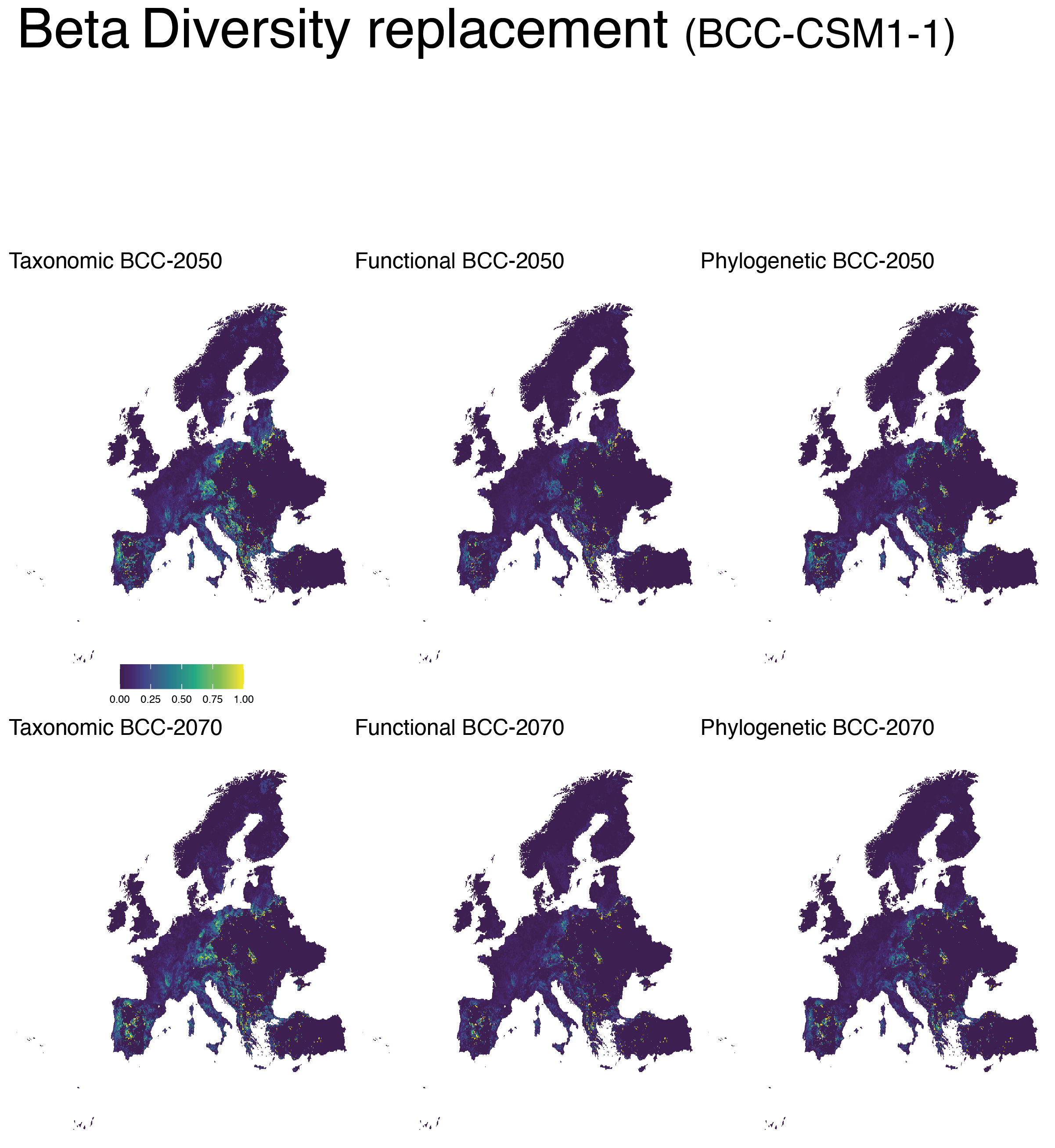

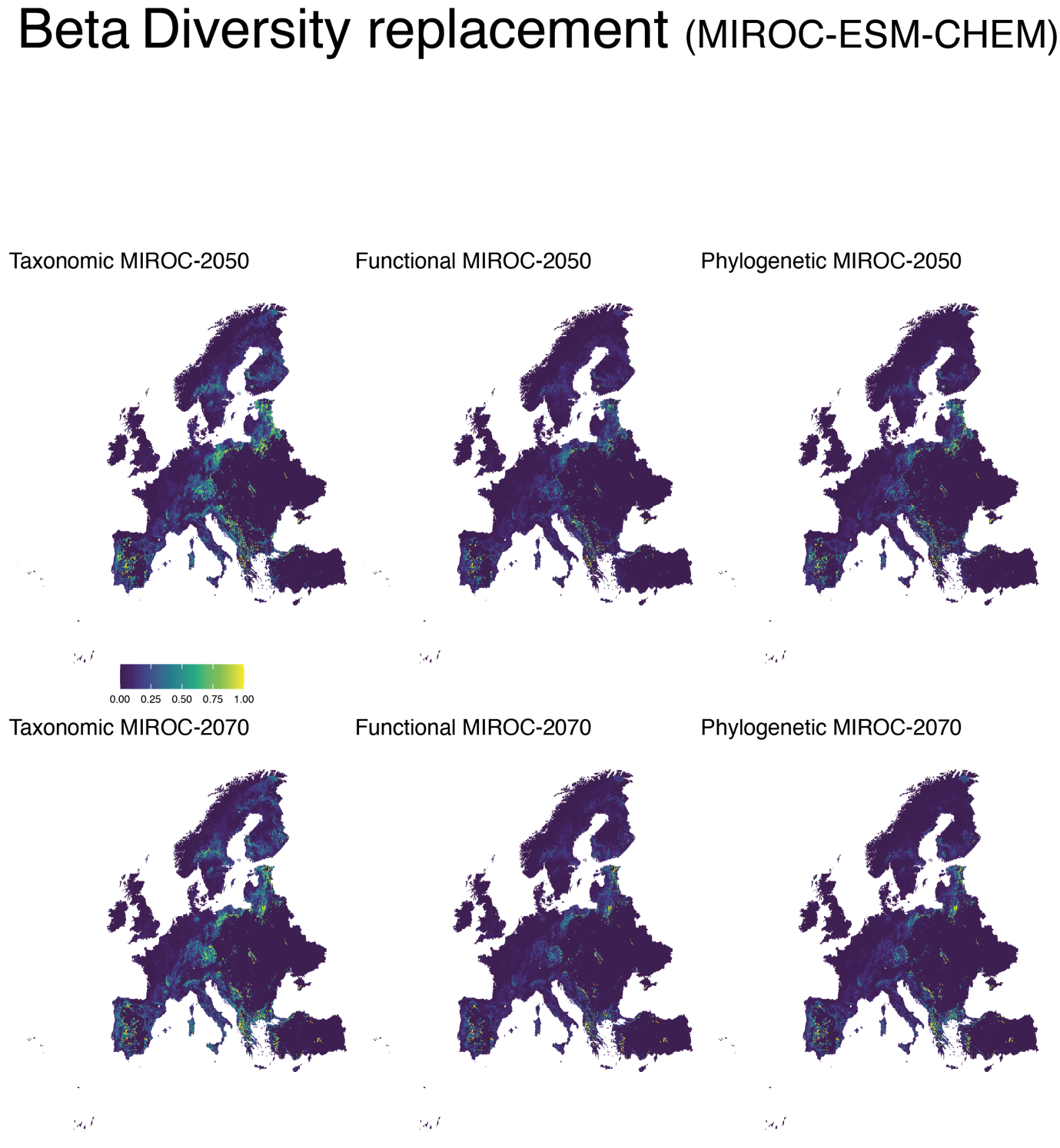

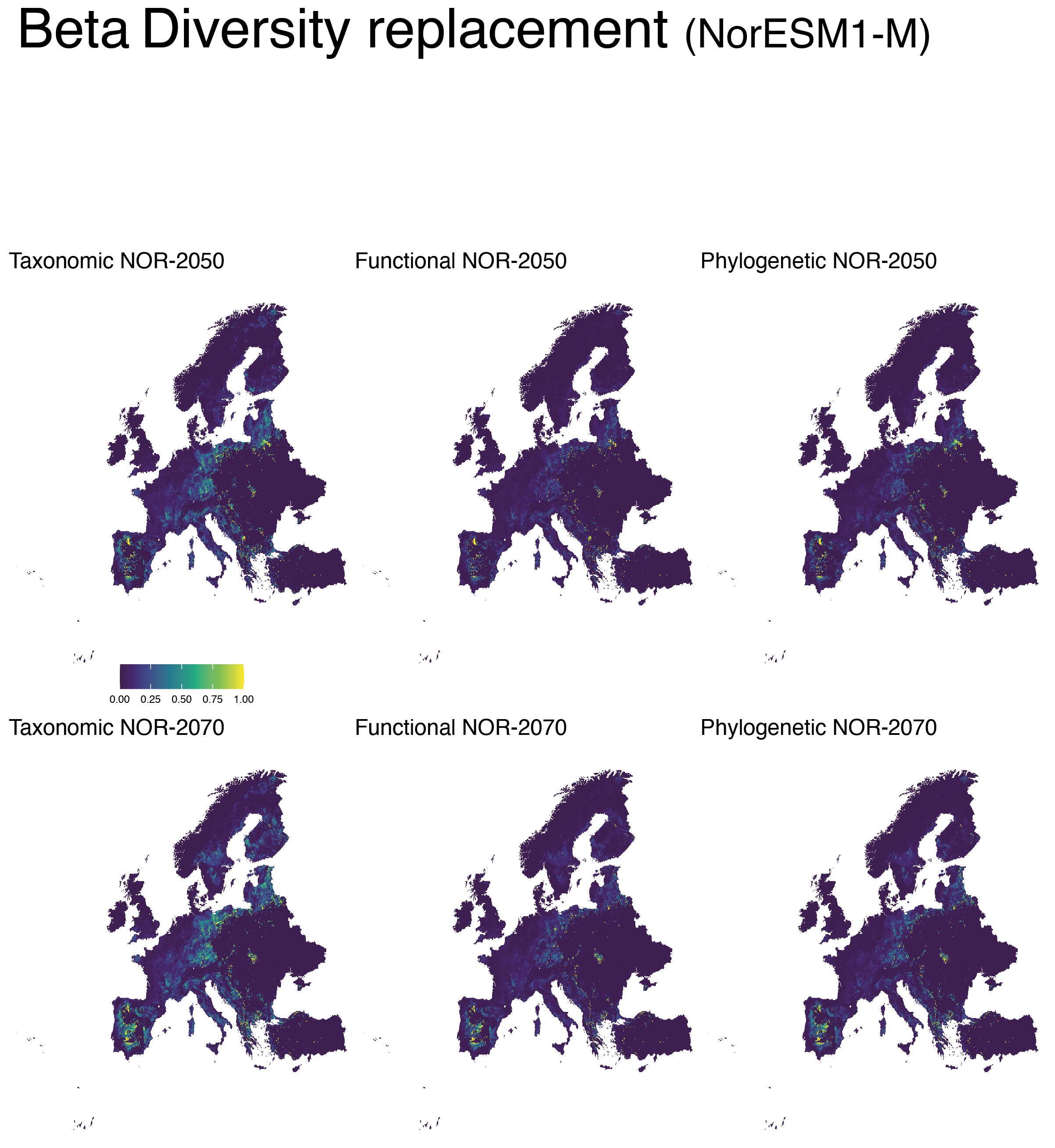

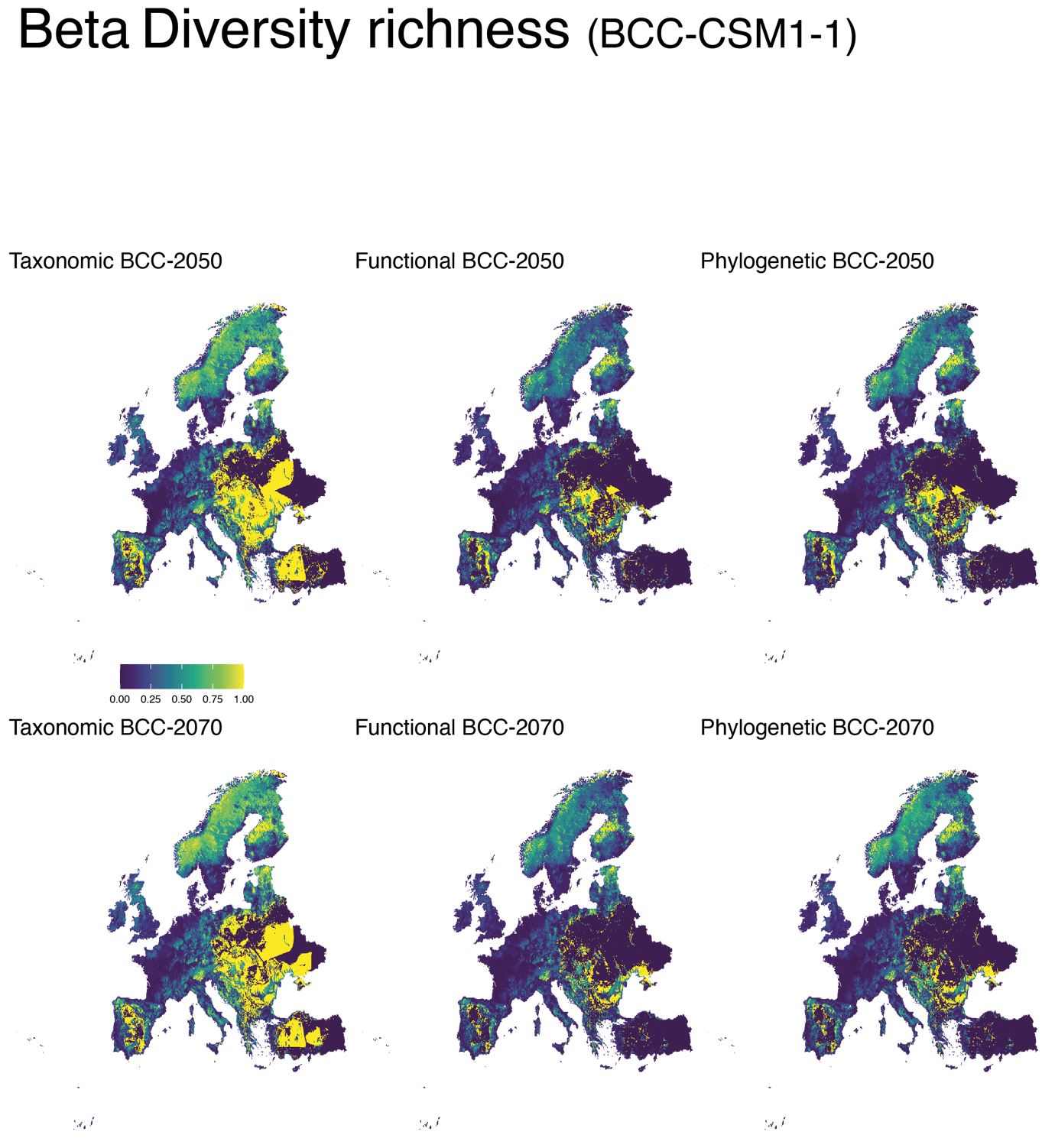

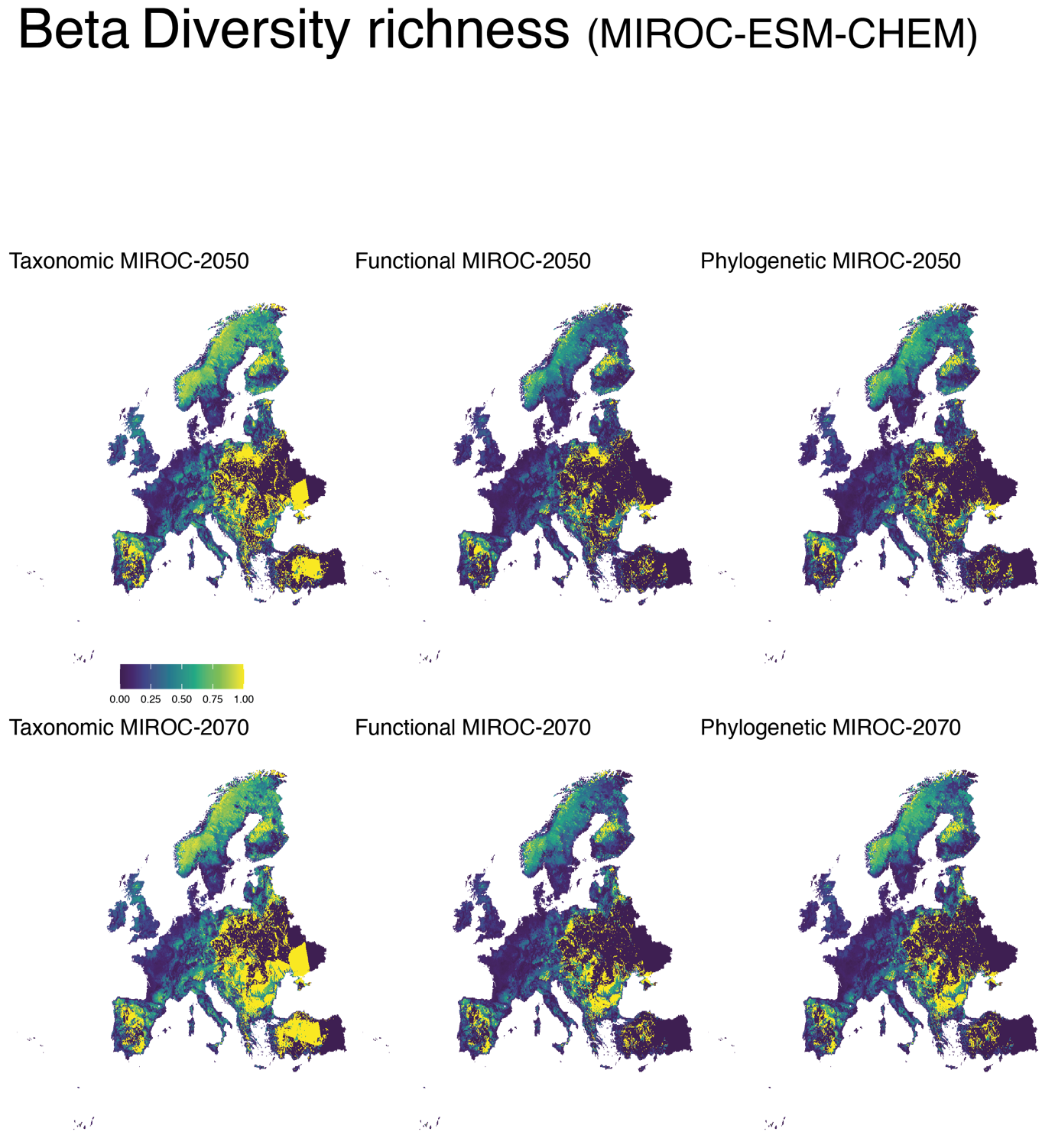

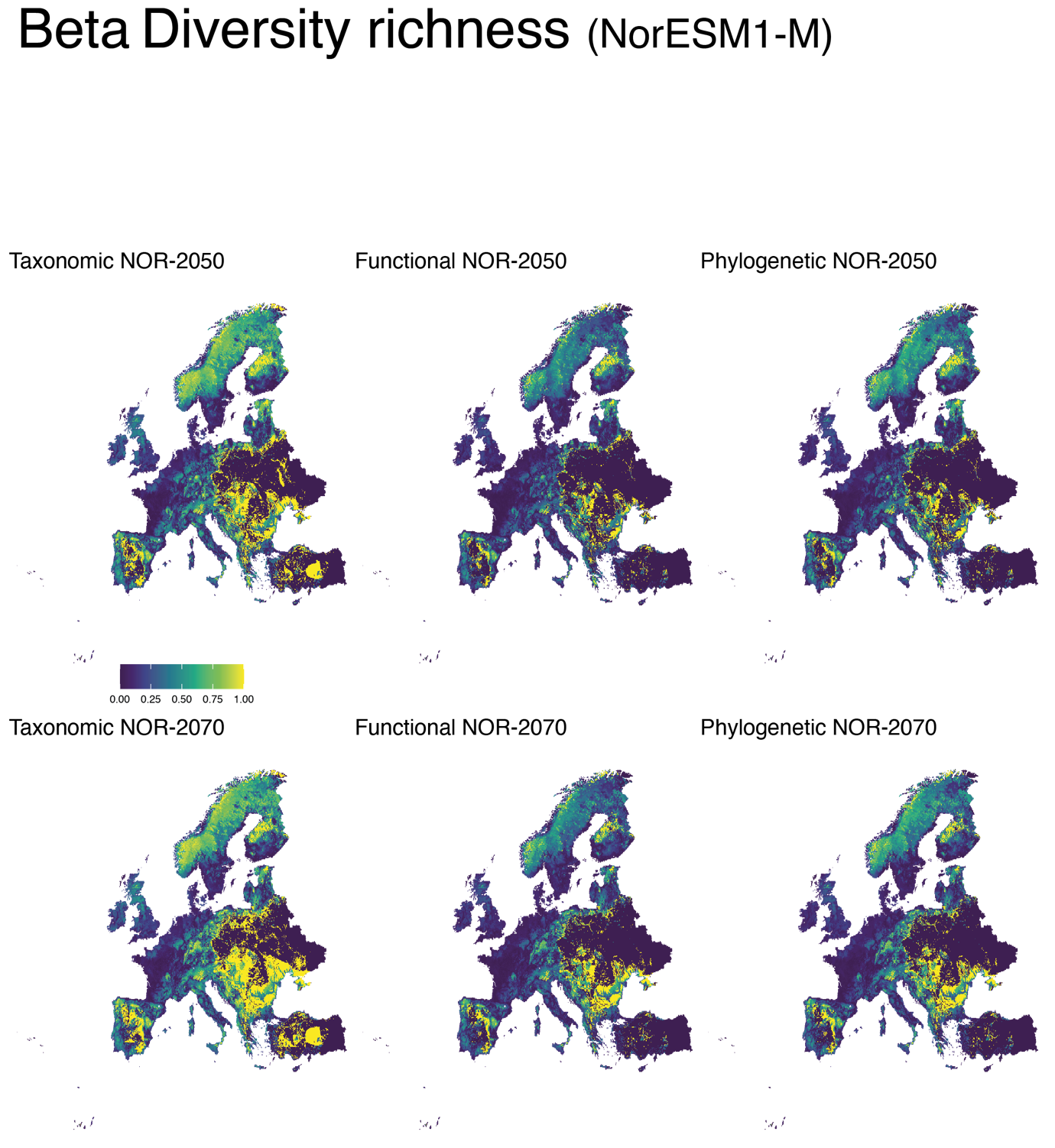
