## Supplementary material for "Climate change will redefine taxonomic, functional, and phylogenetic diversity patterns of Odonata in space and time": S10_Ancestral_character_recostruction

**Supplementary material S10** Reconstruction of ancestral character states for European Odonata per different climate scenarios (BCC-CSM1-1; MIROC-ESM-CHEM; NorESM1-M), time periods (current; 2050; 2070), and three selected characters (variation in habitat suitability; altitudinal shift; centroid shift).

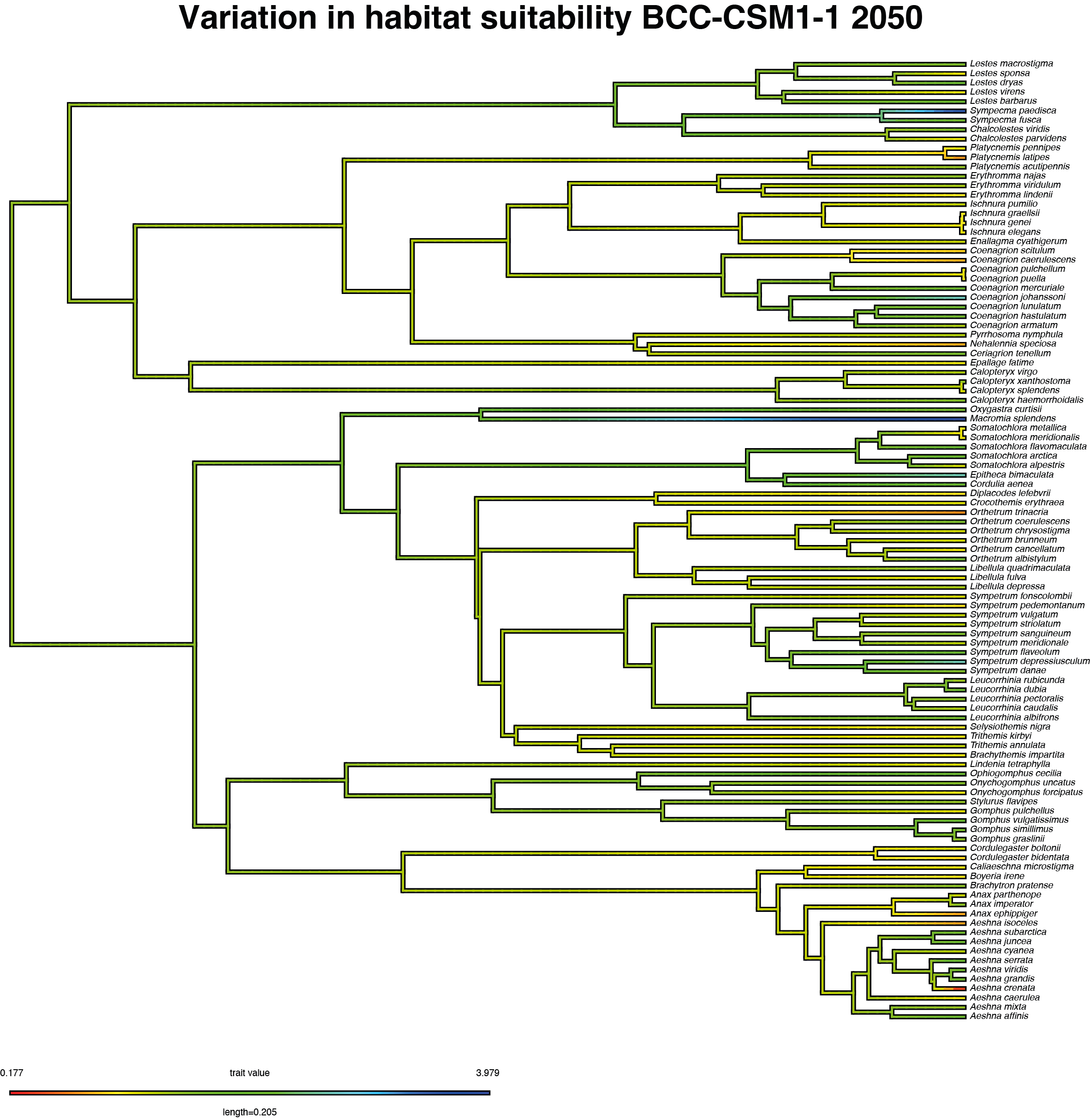

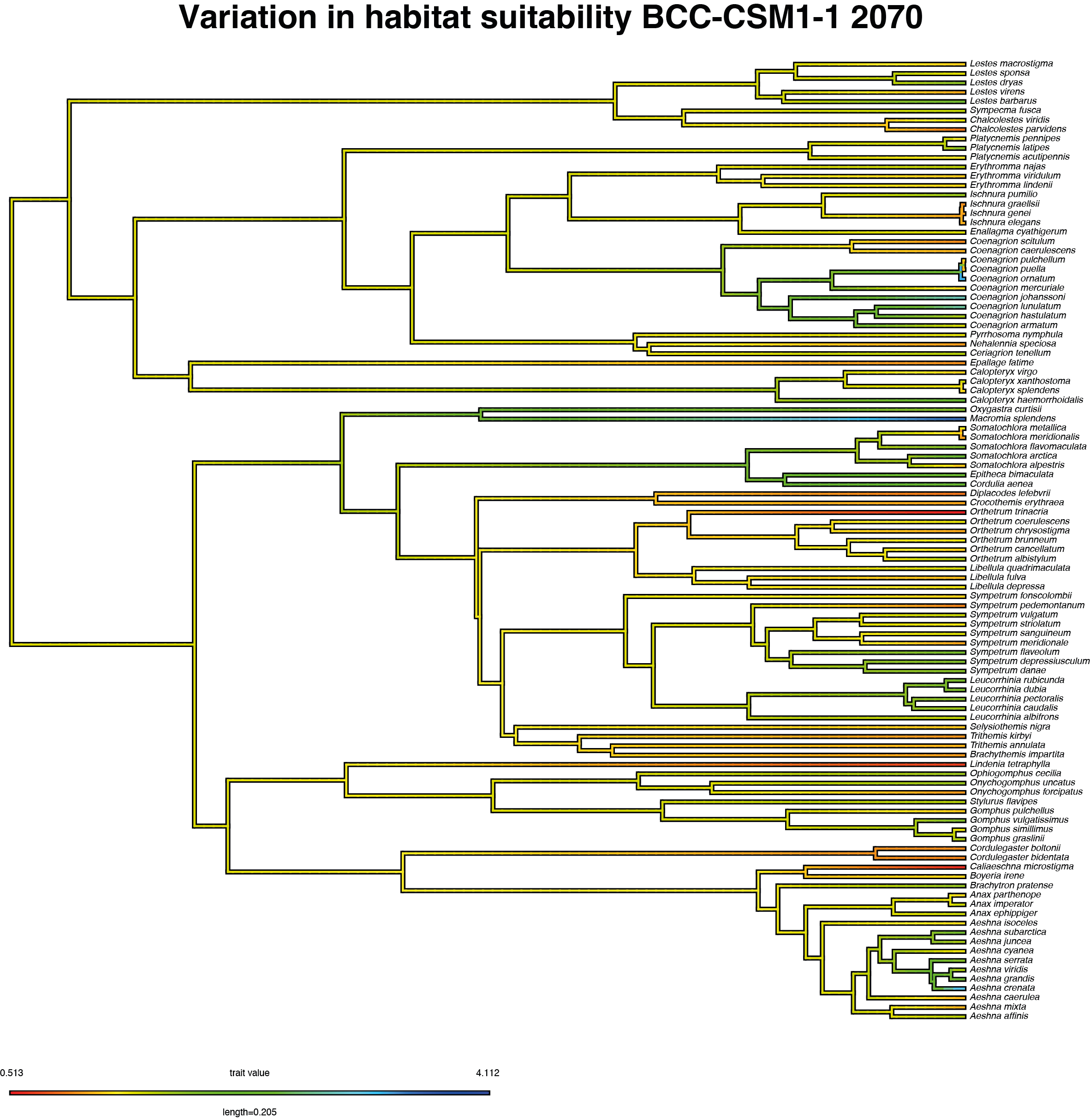

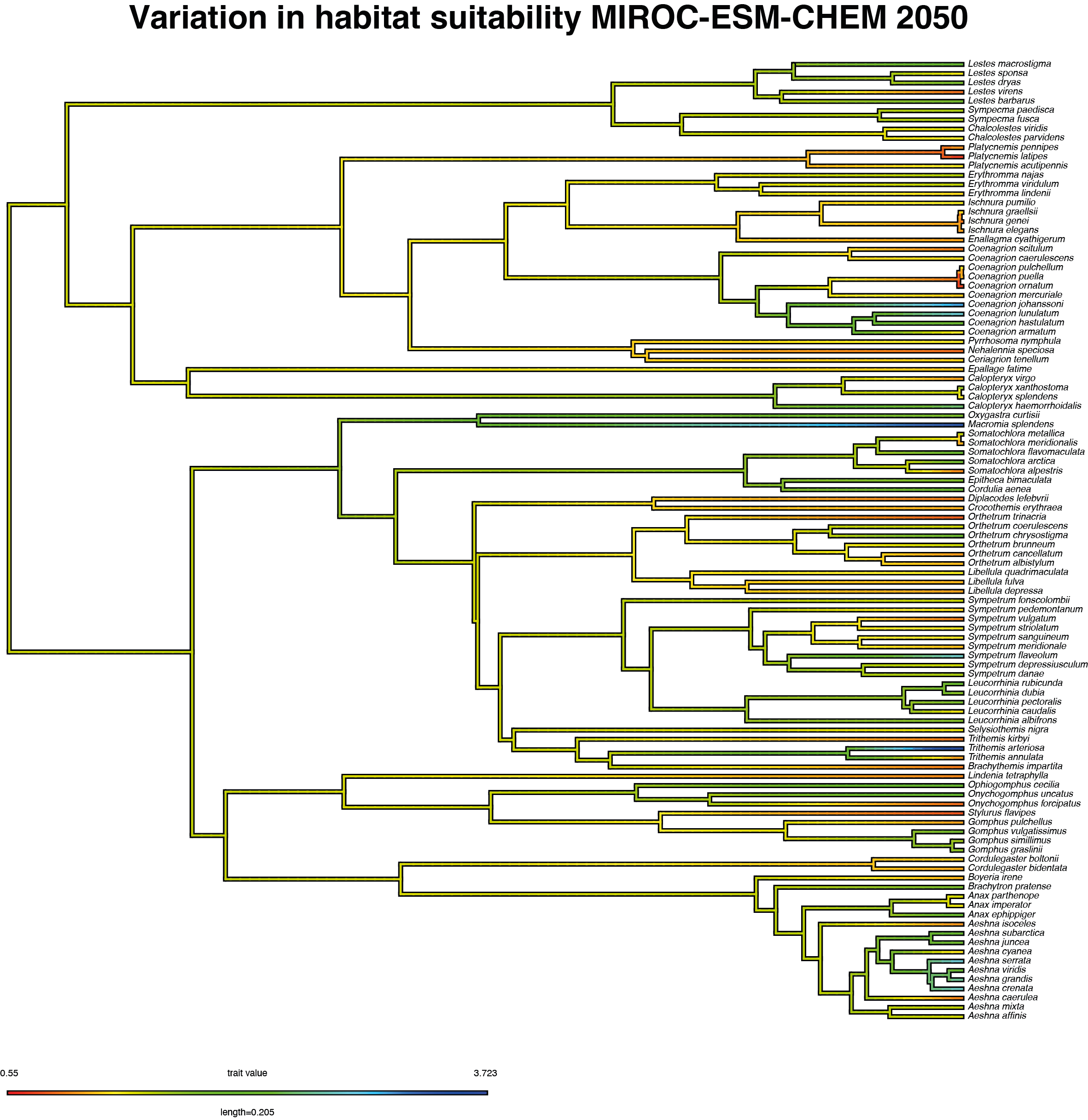

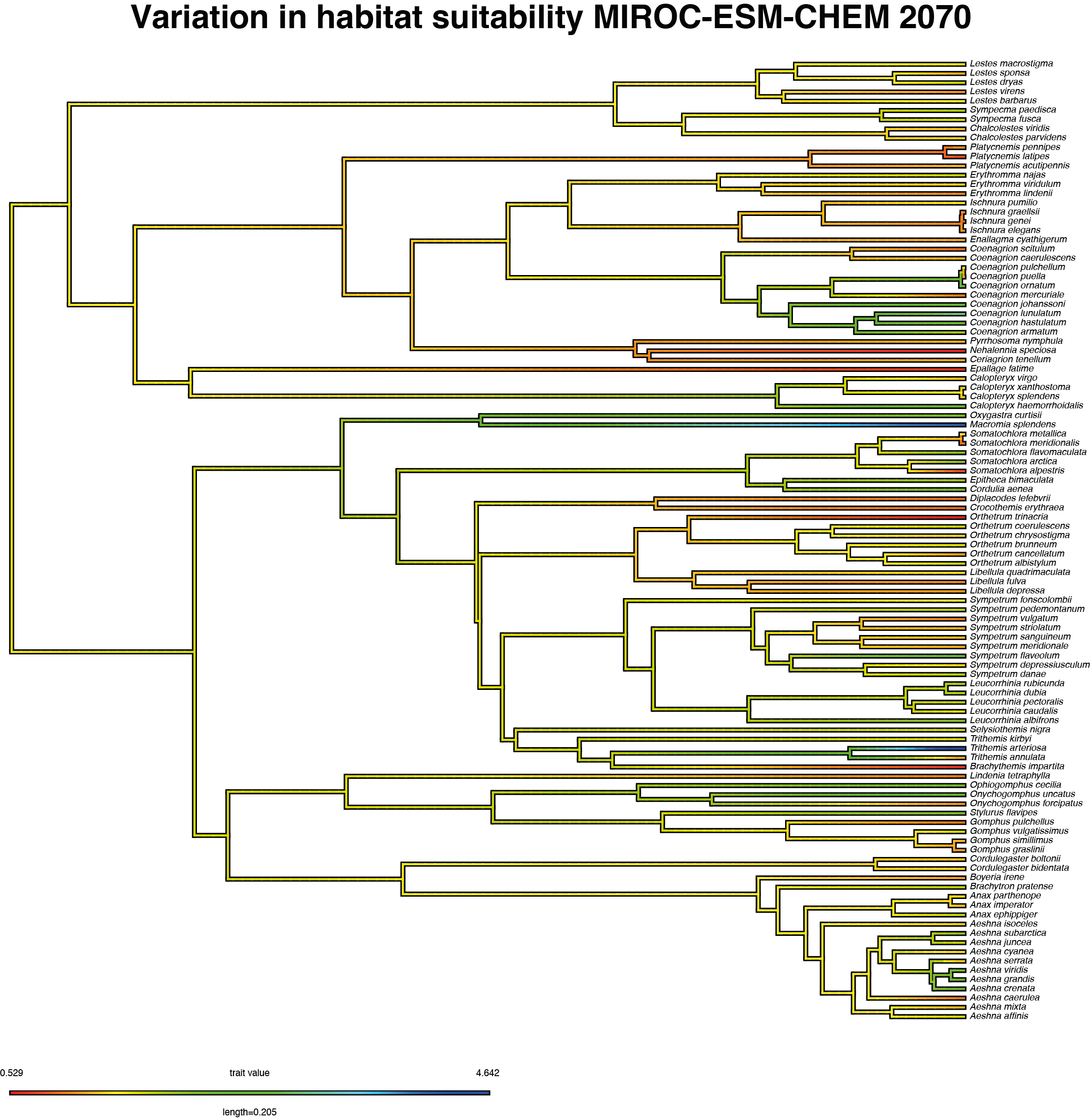

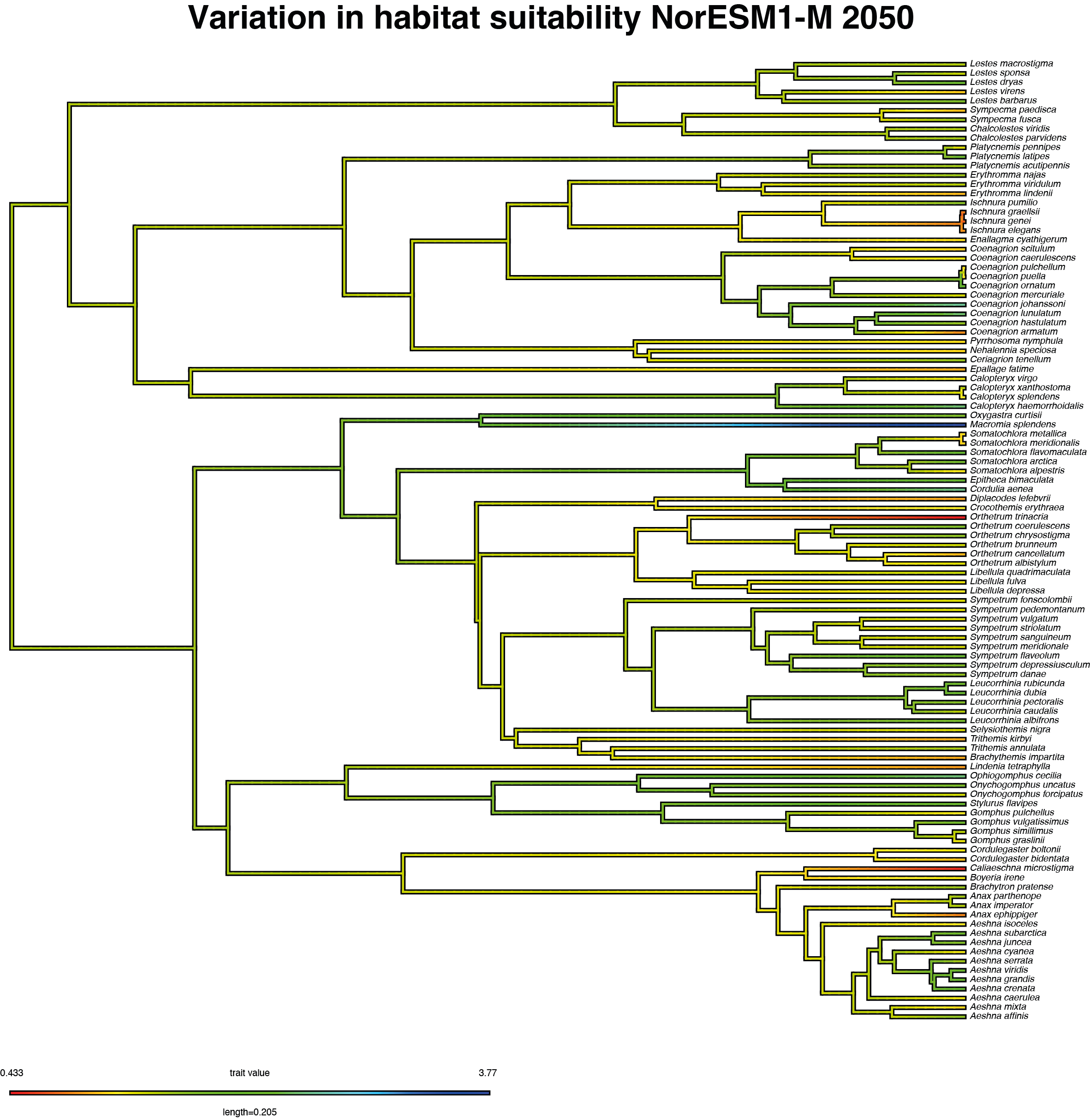

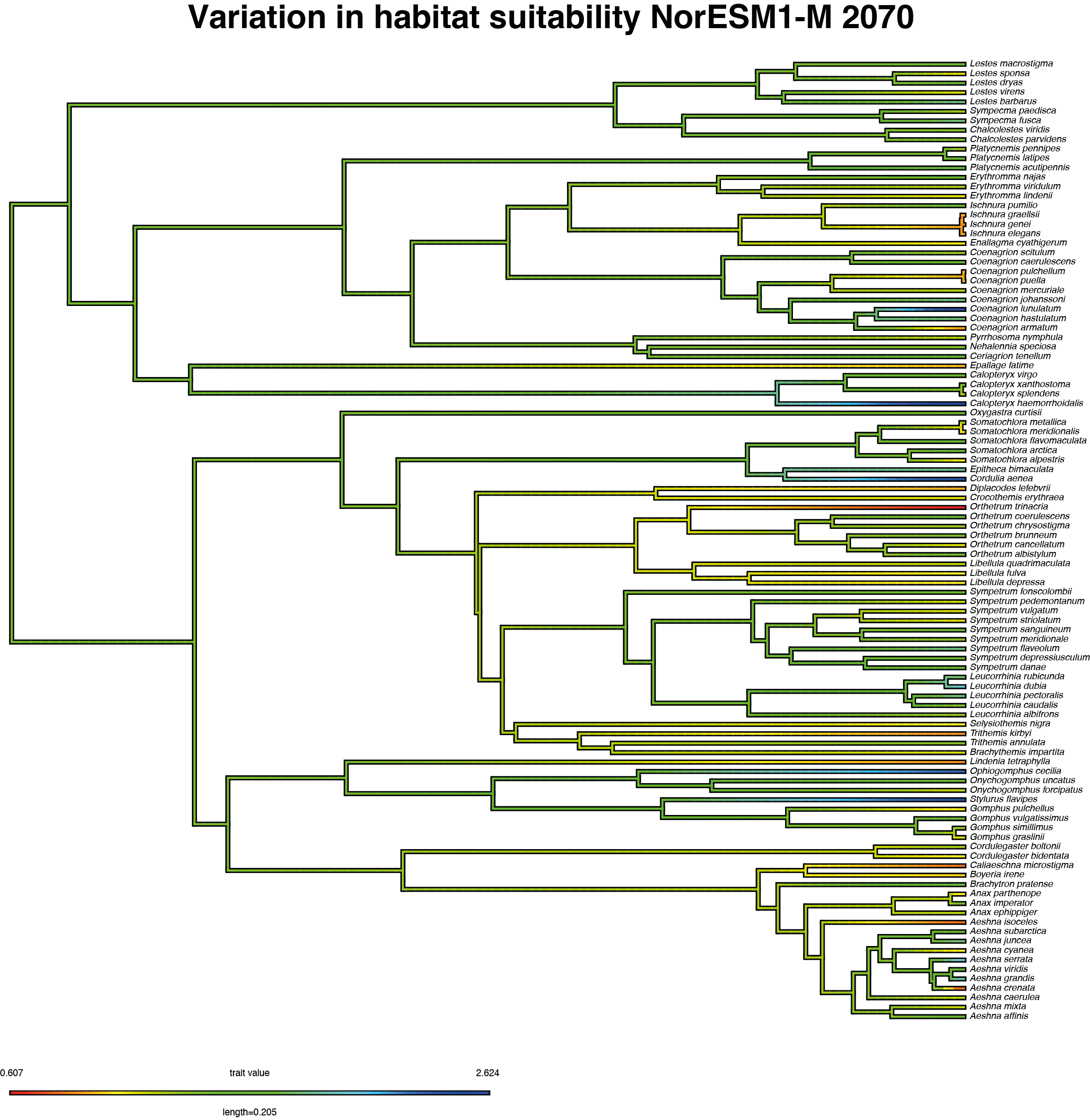

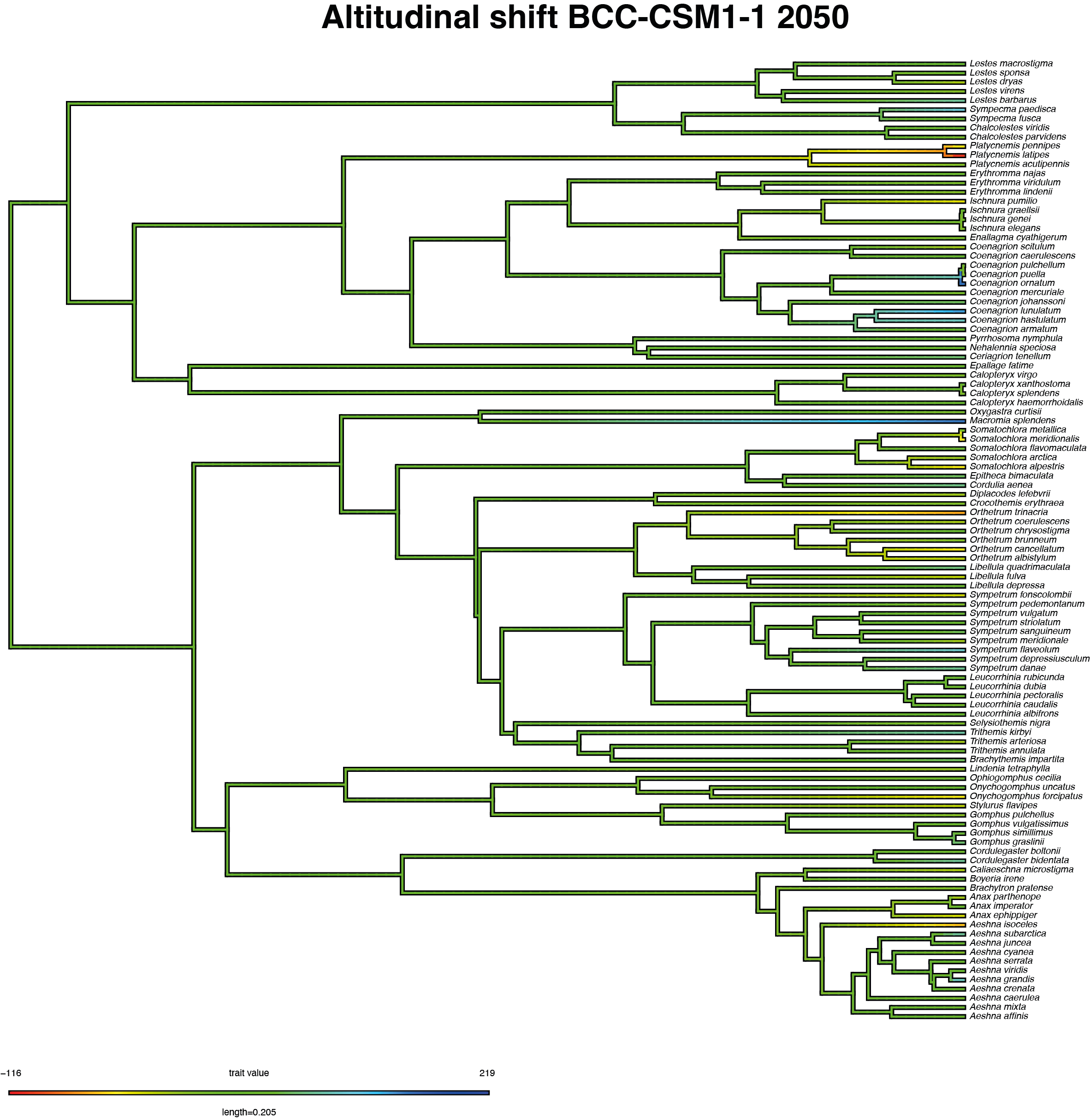

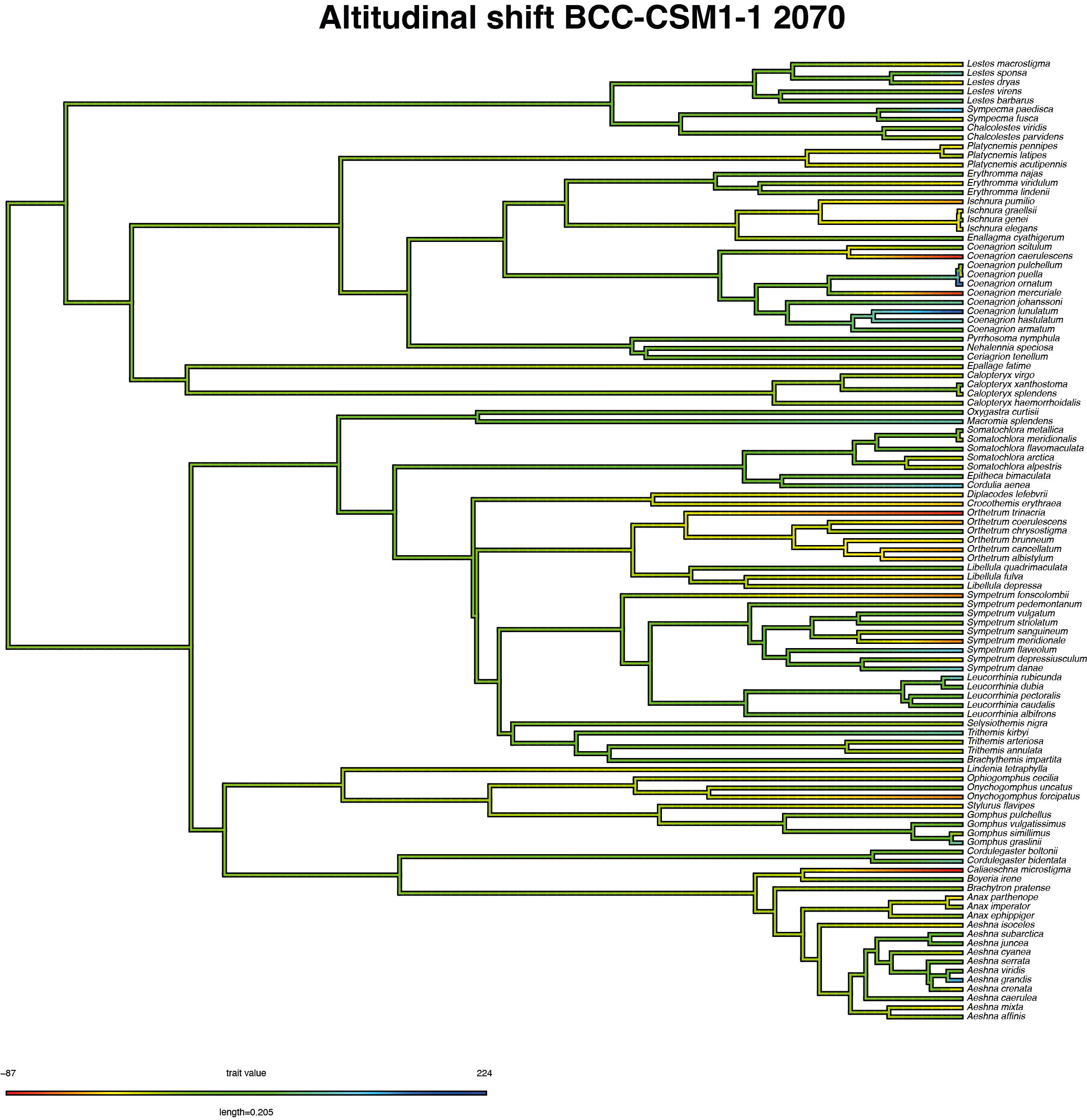

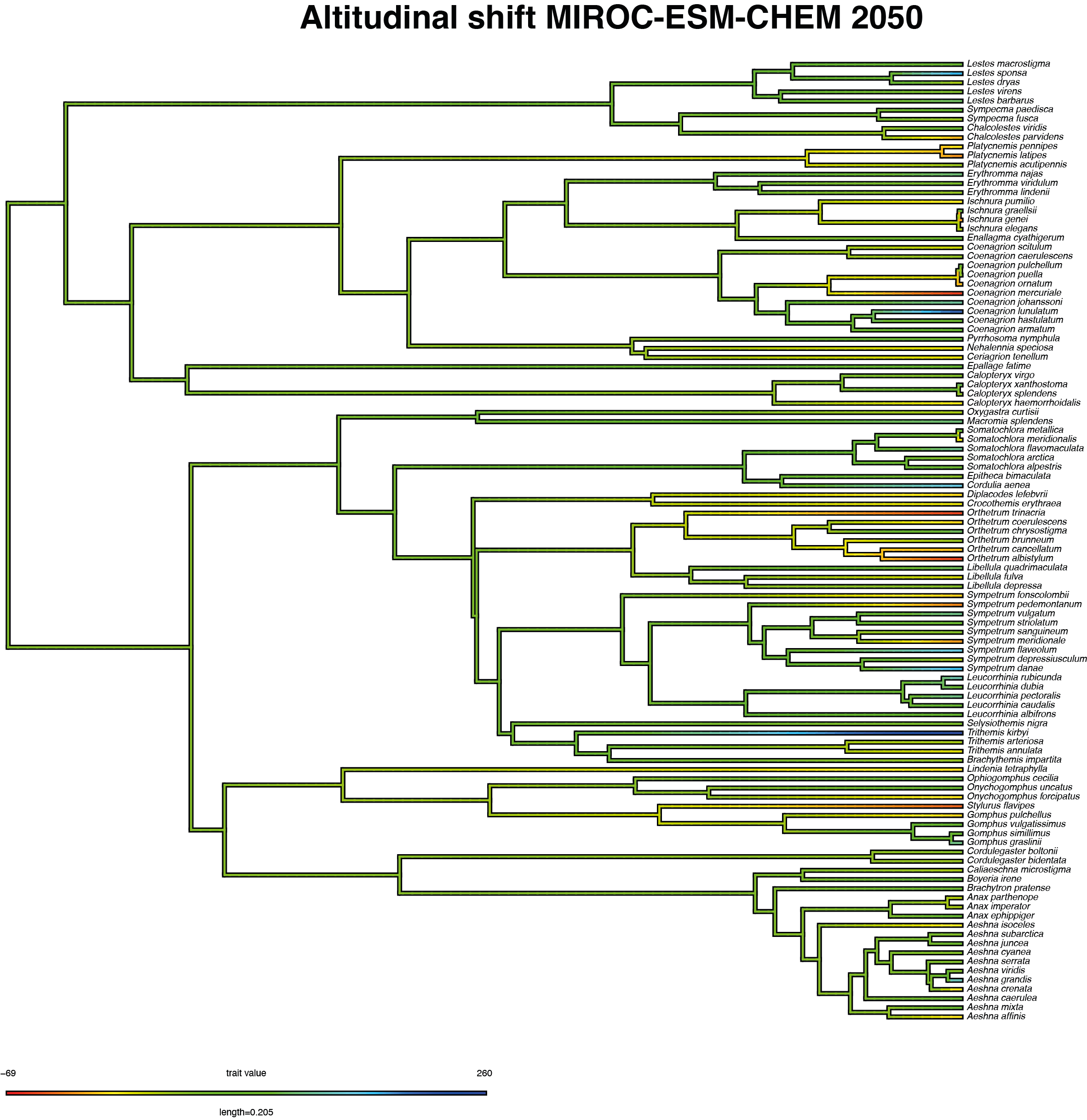

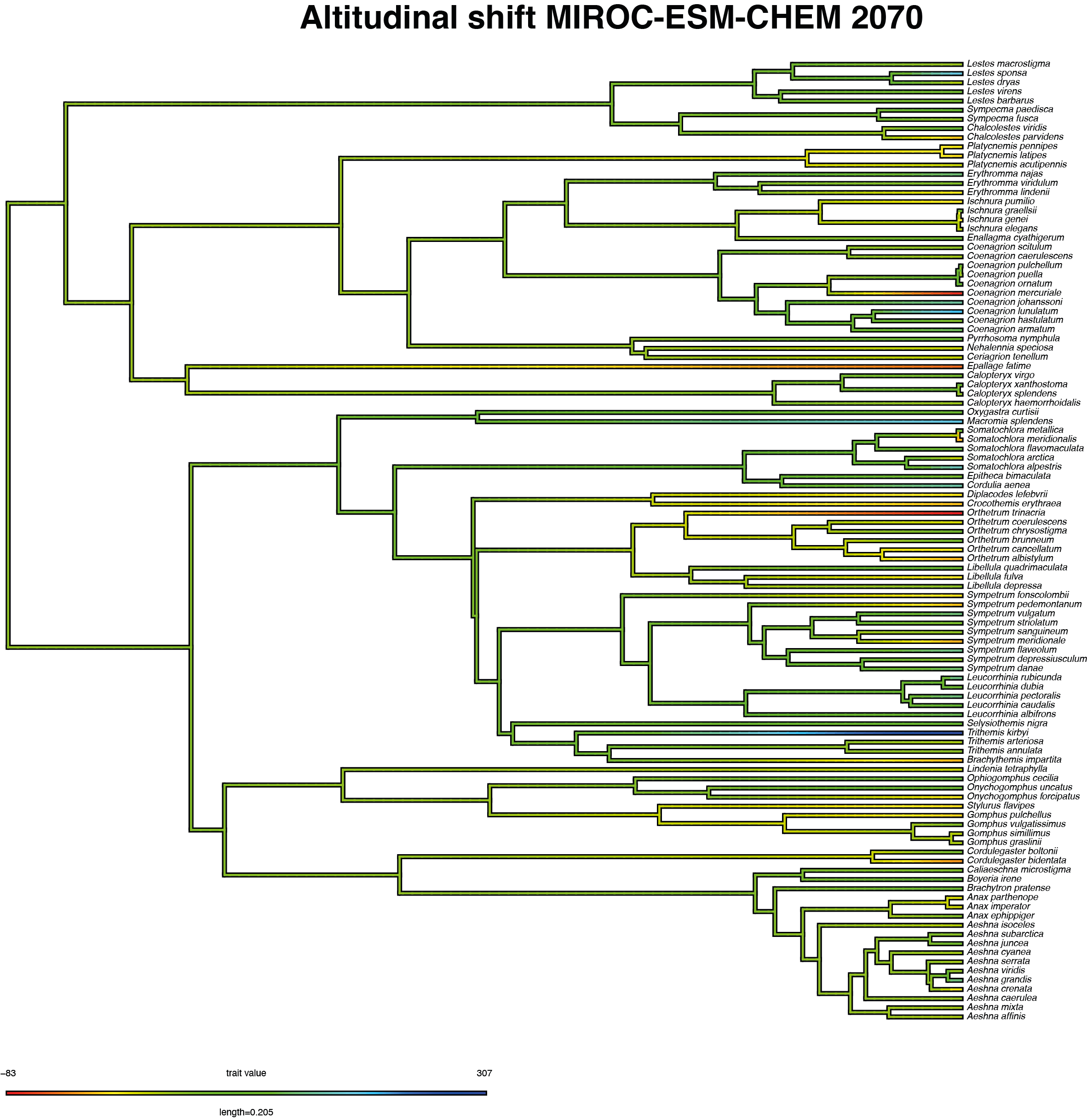

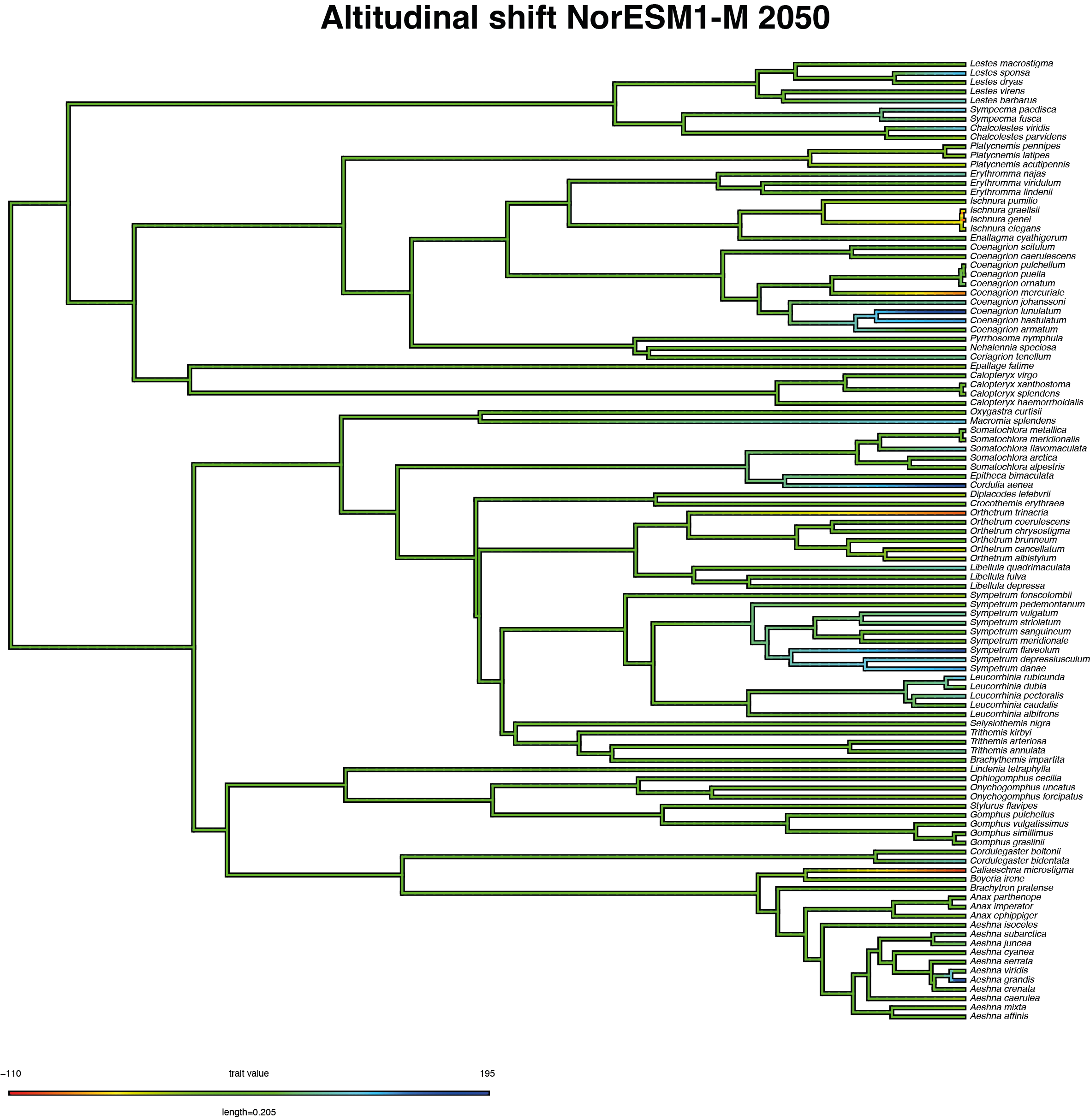

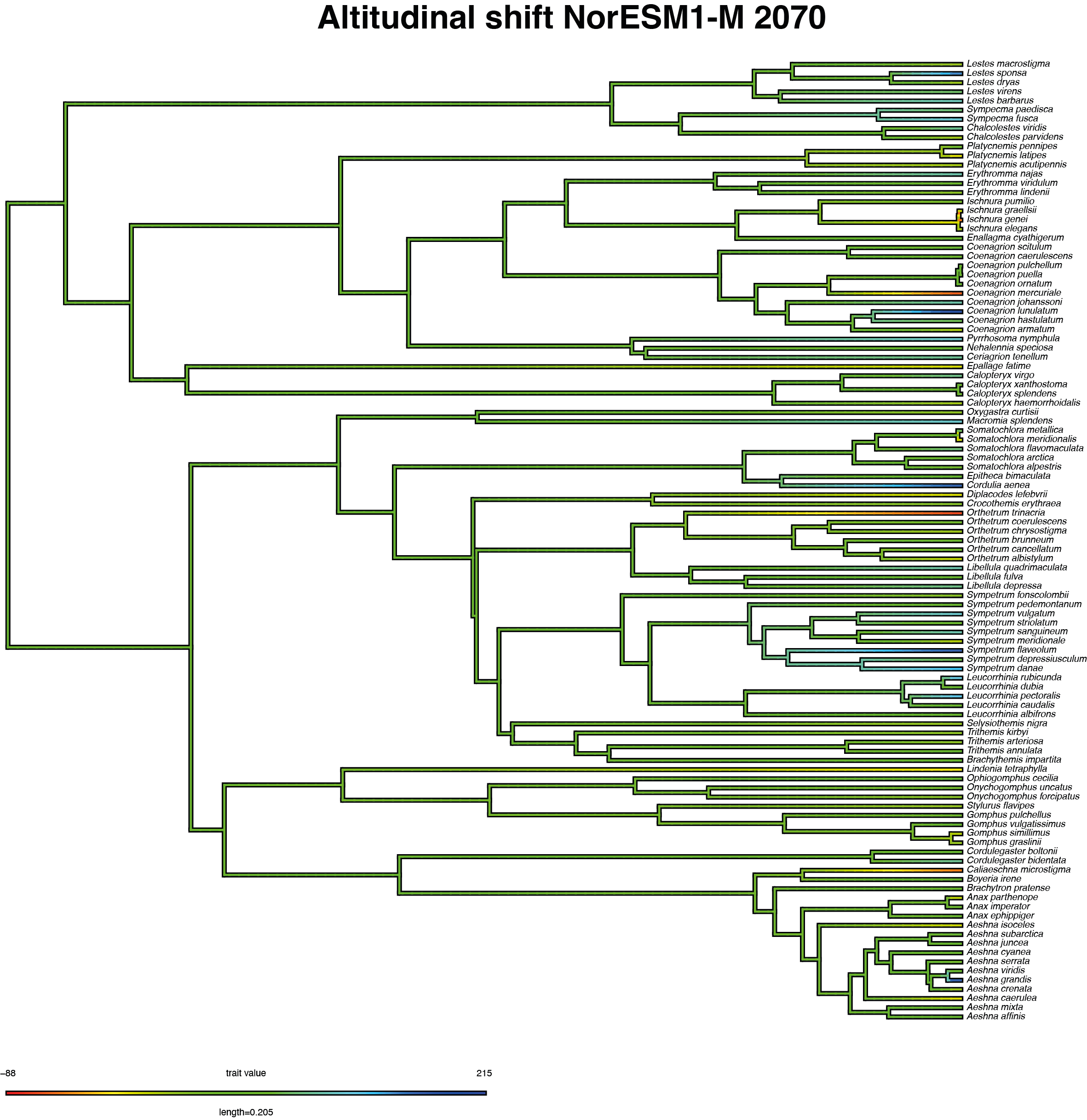

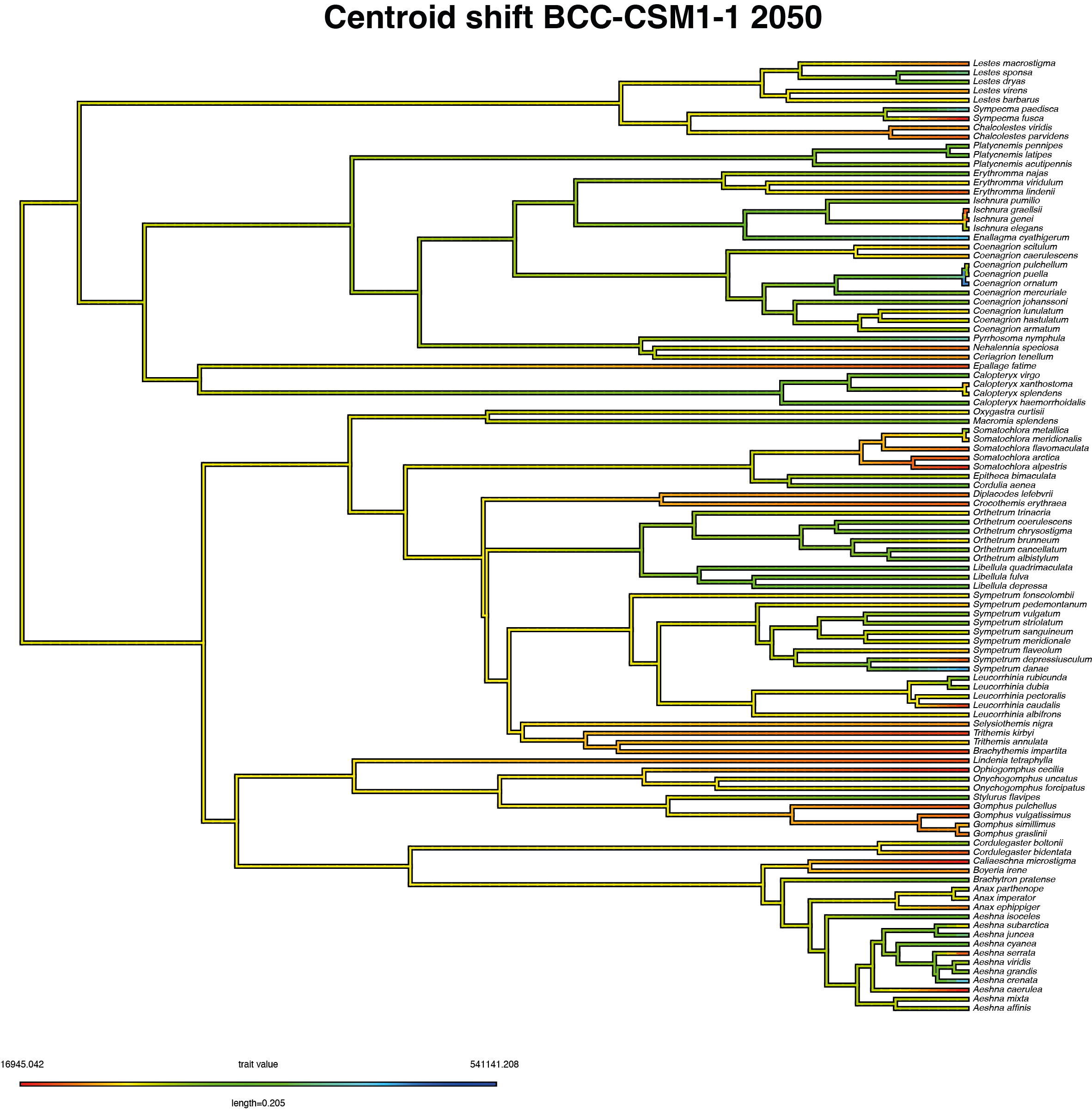

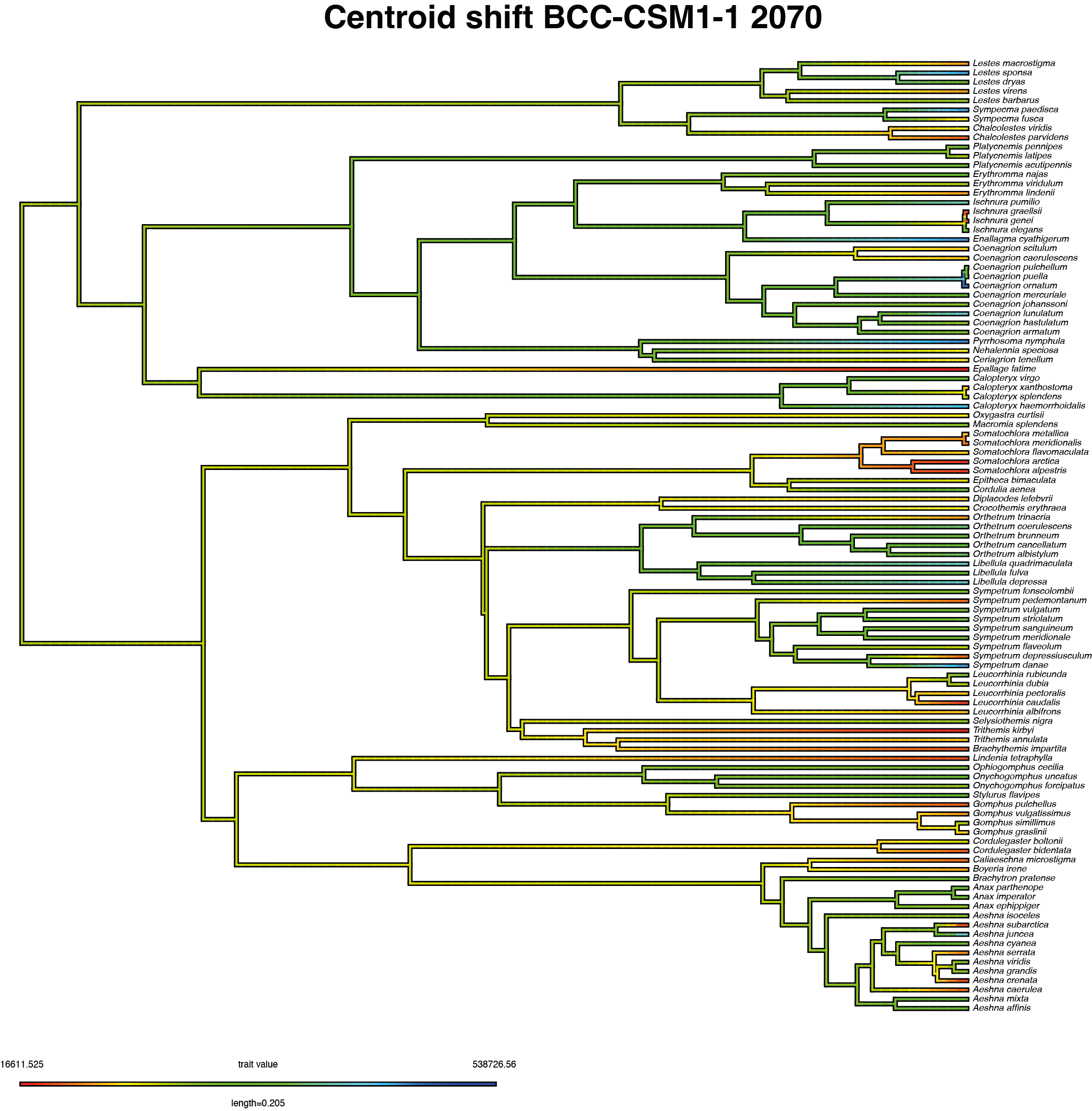

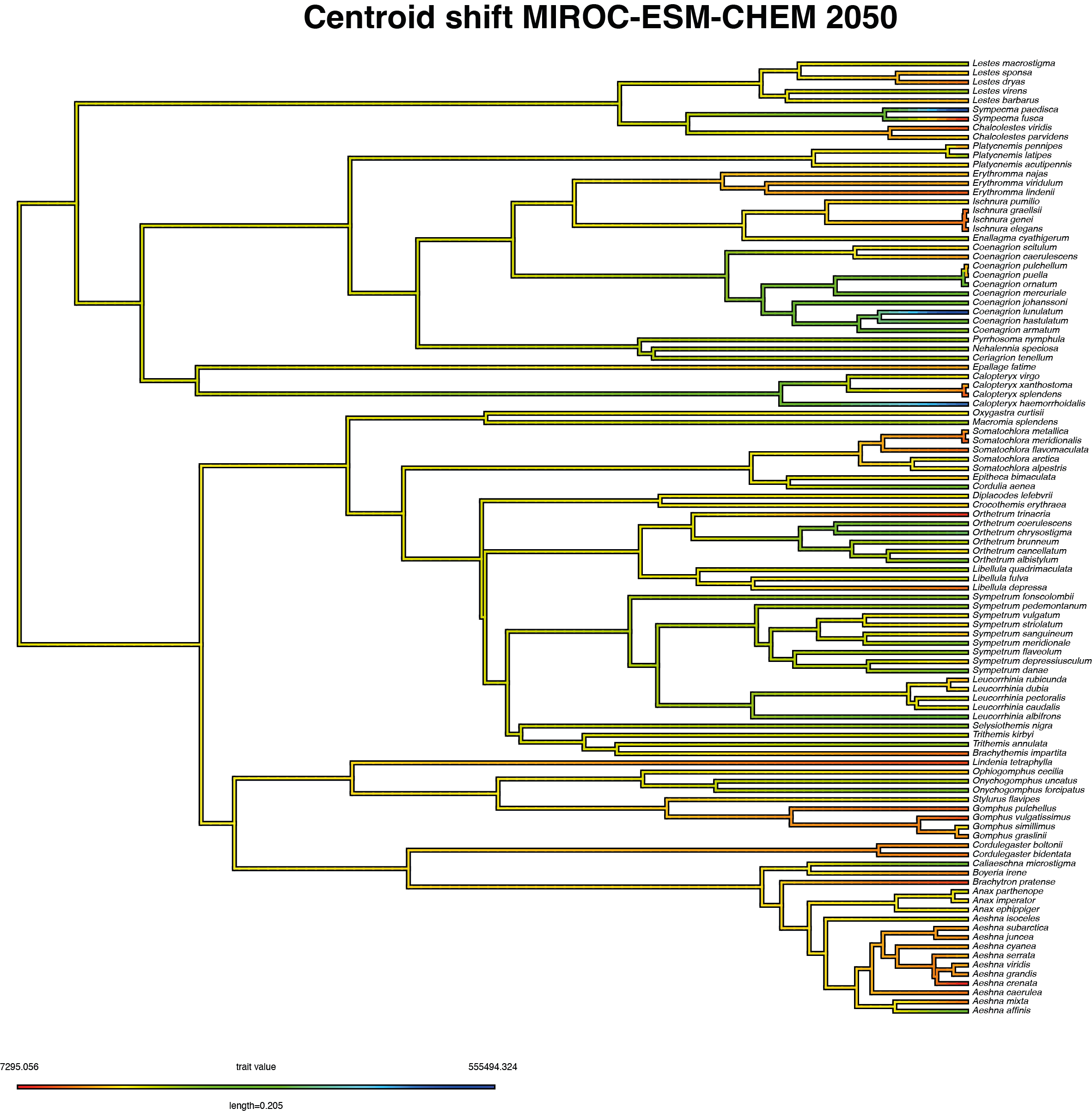

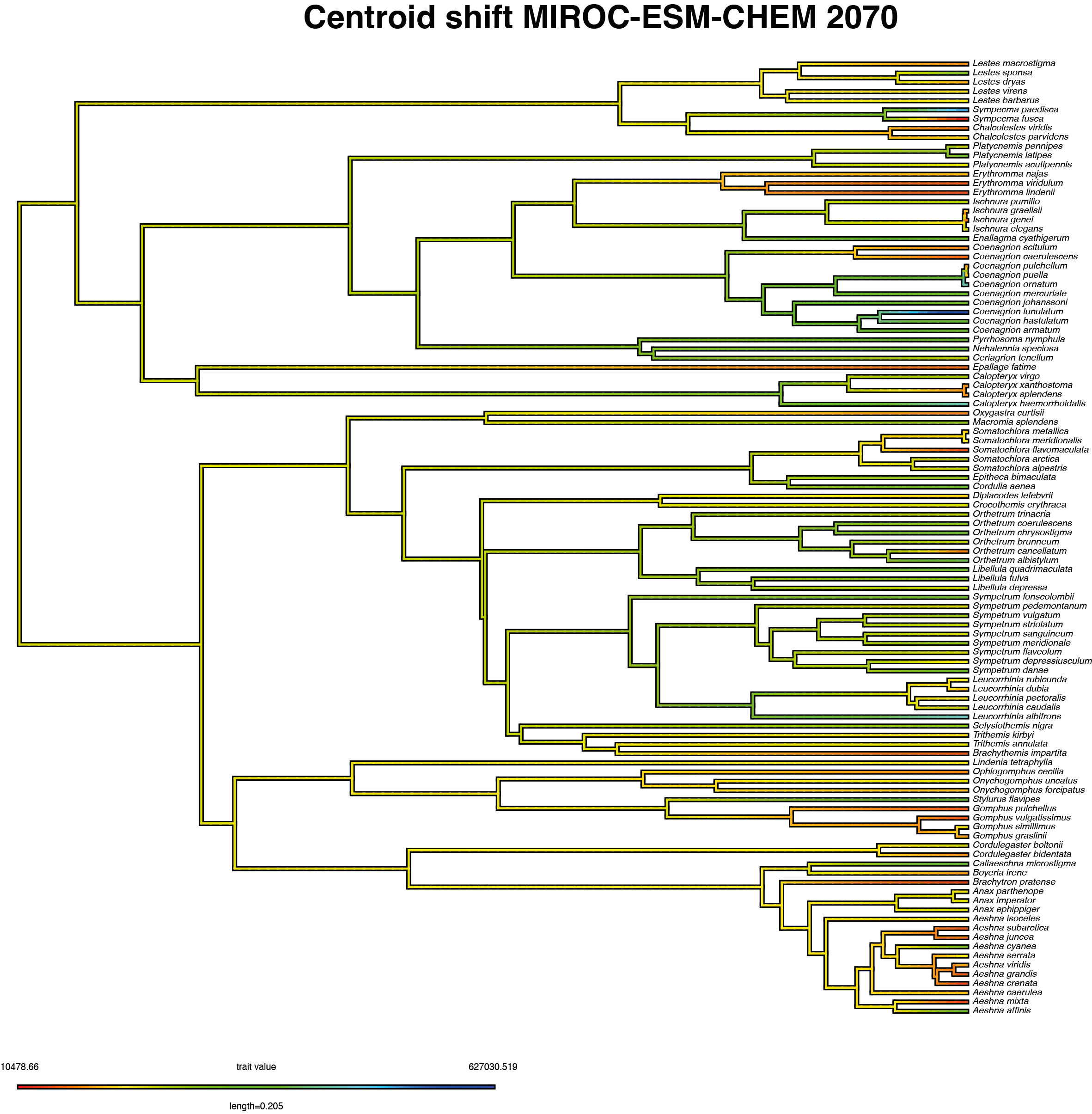

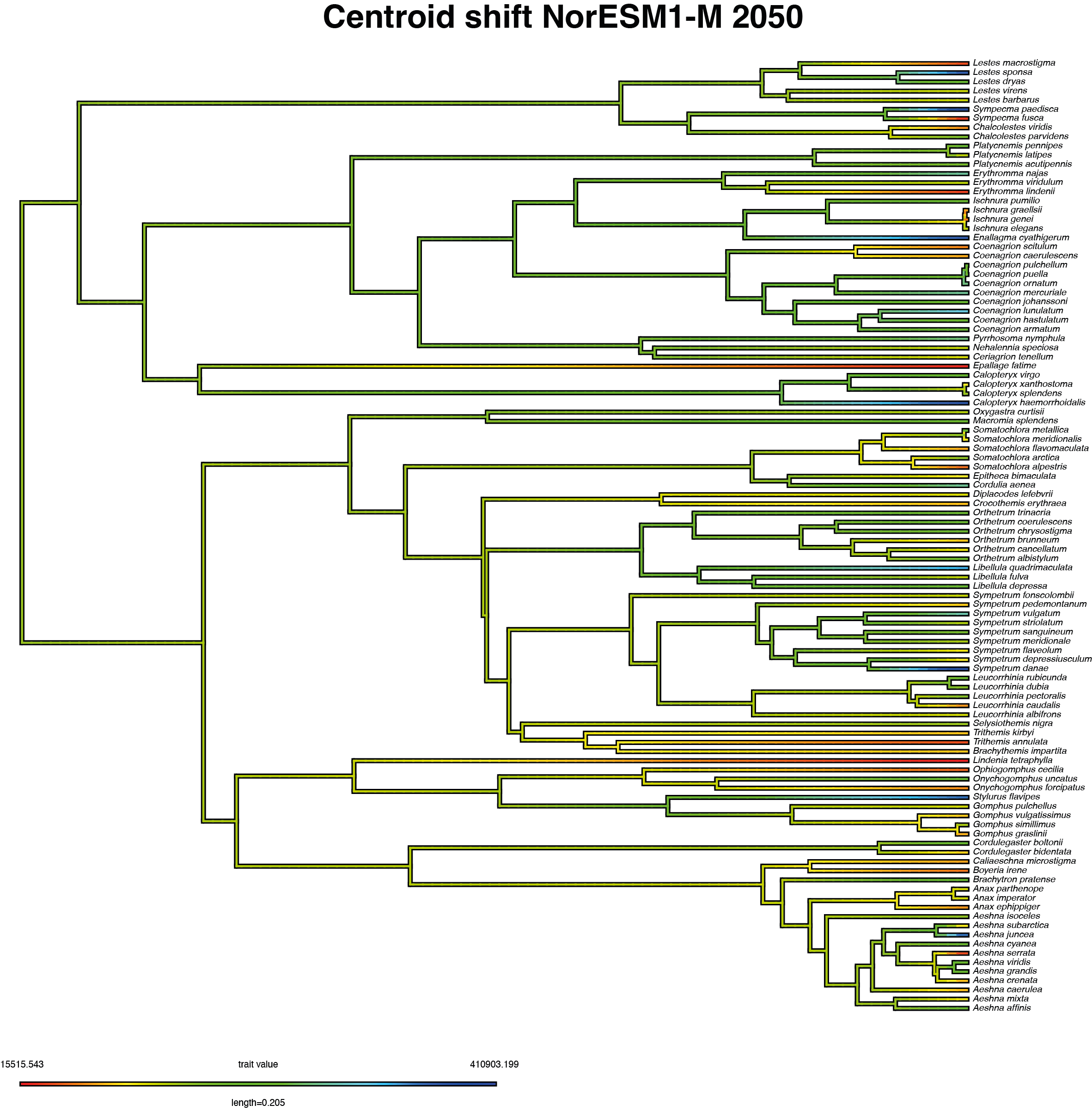

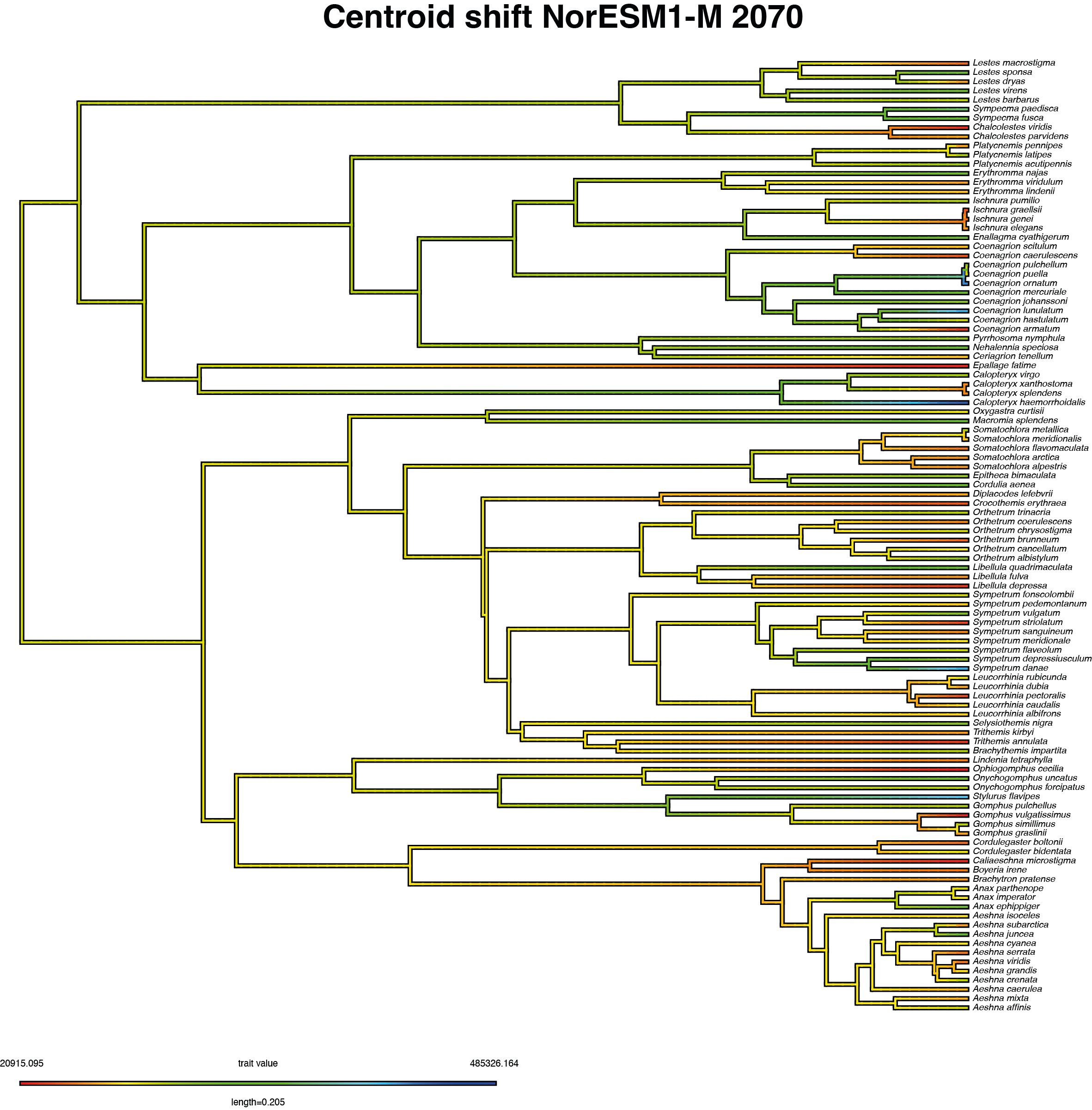
